# HPRC2: A human pangenome reference with near-complete coverage of common genetic variation

**DOI:** 10.64898/2026.07.21.739710

**Authors:** Julian K. Lucas, Prajna Hebbar, Wen-Wei Liao, Juan F. Macias-Velasco, Adam M. Novak, Mobin Asri, Jennifer R. Balacco, Andrew P. Blair, Davide Bolognini, Jana Ebler, Joshua M.V. Gardner, Margarita Geleta, Cristian Groza, Andrea Guarracino, Peter Heringer, Glenn Hickey, Sergey Koren, Shuangjia Lu, Maximillian G. Marin, Christopher Markovic, Mira Mastoras, Capucine Mayoud, Brandy McNulty, Julian M. Menendez, Anna Minkina, Saswat K. Mohanty, Jean Monlong, Katherine M. Munson, Keisuke K. Oshima, David Porubsky, T. Rhyker Ranallo-Benavidez, Alessandro Raveane, William E. Seligmann, Ruhollah Shemirani, Yoshihiko Suzuki, Jack A.S. Tierney, Ivo Violich, DongAhn Yoo, Xiaoyu Zhuo, Derek Albracht, Ivan A. Alexandrov, Jamie Allen, Alawi A. Alsheikh-Ali, Casey Andrews, Dmitry Antipov, Lucinda Antonacci-Fulton, Alexander Arguello, Marcelo Ayllon, Edward A. Belter, Halle D. Bender, Katherine E. Bonini, Silvia Buonaiuto, Shuo Cao, Ann M. Mc Cartney, Pi-Chuan Chang, Xian Chang, Jitender Cheema, Claudio Ciofi, Hiram Clawson, Sarah Cody, Vincenza Colonna, Holland C. Conwell, Mark Diekhans, Maria Angela Diroma, Zheng Dong, Danilo Dubocanin, Jordan M. Eizenga, Parsa Eskandar, Eddie Ferro, Sarah M. Ford, Willard W. Ford, Adam Frankish, Mallory A. Freeberg, Qichen Fu, Shenghan Gao, Yan Gao, Gage H. Garcia, Obed A. Garcia, John E. Garza, Mohammadmersad Ghorbani, Tina A. Graves-Lindsay, Bida Gu, Leanne Haggerty, Nancy F. Hansen, Yue Hao, Todd L. Hillaker, S. Nakib Hossain, Neng Huang, Sarah E. Hunt, Toby Hunt, Nafiseh Jafarzadeh, Nivesh Jain, Maryam Jehangir, Juan Jiang, Juhyun Kim, Bonhwang Koo, Milinn Kremitzki, Daofeng Li, Ronghan Li, Jiadong Lin, Tianjie Liu, Ryan Lorig-Roach, Hailey Loucks, Jane E. Loveland, Jianguo Lu, Walfred Ma, Franco L. Marsico, Jack A. Medico, Younes Mokrab, Shabir Moosa, Avelina Moreno-Ochando, Shinichi Morishita, Jonathan M. Mudge, Njagi Mwaniki, Nasna Nassir, Chiara Natali, Shloka Negi, Lingbin Ni, Faith Okamoto, Chie Owa, Sadye Paez, Clelia Peano, Brandon D. Pickett, Laura Pignata, Timofey Prodanov, Anandi Radhakrishnan, Brian J. Raney, Andreas Rechtsteiner, Luyao Ren, Fedor Ryabov, Samuel Sacco, Farnaz Salehi, Aarushi Sehgal, Mahsa Shabani, Shadi Shahatit, Vikram S. Shivakumar, Swati Sinha, Linnéa Smeds, Steven J. Solar, Marco Sollitto, Nicole Soranzo, Marie-Marthe Suner, Arda Söylev, Chad Tomlinson, Francesca Floriana Tricomi, Matteo Tommaso Ungaro, Rahul Varki, Brian P. Walenz, Charles Wang, Lisa E. Wang, Aaron M. Wenger, Conor V. Whelan, Zilan Xin, Zheng Xu, Wenjin Zhang, Ying Zhou, Giulia Zunino, Nicolas Altemose, Floris P. Barthel, Christina Boucher, Guillaume Bourque, Andrew Carroll, Monika Cechova, Mark J.P. Chaisson, Haoyu Cheng, Robert Cook-Deegan, Daniel Doerr, Richard Durbin, Anna-Sophie Fiston-Lavier, Giulio Formenti, Stephanie M. Fullerton, Robert S. Fulton, Shilpa Garg, Nanibaa’ A. Garrison, Richard E. Green, Carol W. Greider, Melissa Gymrek, Maximilian Haeussler, Mohammad Amiruddin Hashmi, David Haussler, Alexander G. Ioannidis, Charles H. Langley, Ben Langmead, Heather A. Lawson, Glennis A. Logsdon, Kateryna D. Makova, Fergal J. Martin, Matthew W. Mitchell, Pilar N. Ossorio, Nadia Pisanti, Pjotr Prins, Mikko Rautiainen, Arang Rhie, Michael C. Schatz, Laura B. Scheinfeldt, Kishwar Shafin, Jouni Sirén, Andrew B. Stergachis, Ahmad Abou Tayoun, Mohammed Uddin, Flavia Villani, Mitchell R. Vollger, Kai Ye, Evan E. Eichler, Erik Garrison, Ira M. Hall, Erich D. Jarvis, Eimear E. Kenny, Heng Li, Jonathan LoTempio, Tobias Marschall, Karen H. Miga, Adam M. Phillippy, Ting Wang, Benedict Paten

## Abstract

A pangenome reference overcomes the inherent limitation of any individual reference genome by integrating the variation present in a population. We present the Human Pangenome Reference Consortium’s (HPRC) Release 2 (HPRC2), an openly available, second phase pangenome that is an approximately fivefold expansion in genome number over HPRC Release 1 (HPRC1) and measurable improvement in genome completeness, contiguity, and accuracy. Selecting samples with a principled algorithm prioritising common variant coverage, HPRC2 contributes 460 haplotypes that together capture over 99% of common variation observed in the All of Us Research Program v8 cohort. Combining high-coverage long and ultra-long reads with modern assemblers and polishers, we produce thousands of telomere-to-telomere (T2T) chromosomes, and relative to HPRC1 halve the number of structurally unreliable regions as well as individual base errors per haplotype. We complement the assemblies with whole genome multiple alignments and gene annotations, and derive formal pangenome coordinate systems for addressing off-reference variation, demonstrating that individual human genomes contain more than one hundred thousand variants not succinctly described with respect to existing reference genomes. We also present the first matched long-read backed pantranscriptome and panepigenome at this scale, provide continuous local-ancestry estimates spanning every genome, and outline a host of new tools and applications that leverage the pangenome resource for improved genomics analysis.

## Main

The initial publication of a draft human reference genome^1^ launched a generation of biomedical research grounded in a single linear reference. That reference matured through successive revision and careful curation^2^, but remains notably incomplete. Nearly two decades later, the first gap-free T2T haploid assembly, T2T-CHM13, resolved an additional 8% of the human genome^3^. With major improvements in genome assembly^4,5^, it is now joined by the first complete diploid T2T genome assemblies^6–8^.

Relative to a linear reference, a pangenome reference aspires to be a reference for a population, capturing a set of segregating variants not found solely in any one genome. A pangenome can be viewed as consisting of three key components: haplotype-resolved genome assemblies from multiple individuals, alignments that identify regions shared among those assemblies and regions where they differ, and annotations that describe both shared and variable genes and other functional elements^9^. These elements can be expressed as a variation graph whose topology encodes the alignment, whose paths encode the assemblies, and on which annotations can be expressed^10–12^. The first Human Pangenome Reference Consortium (HPRC) release (HPRC1)^13^ delivered such a resource at a meaningful scale, building on earlier efforts^14^.

Alongside HPRC1, population-scale long-read structural variant catalogs now span hundreds to thousands of individuals^15–17^, and regional pangenome efforts are capturing additional population-specific variation^18,19^. Outside of humans, comparable approaches in primates^20^, livestock^21^, and plants^22^, reflect a broader shift toward pangenomes as a framework for comprehensively understanding genetic and functional variation^23^. HPRC1 and these related resources have catalyzed a generation of pangenome-aware methods, which now include high-performance short-^24^ and long-read aligners^25^, best-in-class variant callers^26,27^, and methods for imputation-based haplotype reconstruction^28,29^ that extract latent information from existing, predominantly short-read datasets, as well as methods tailored towards functional pangenomics^30,31^.

Here we present HPRC Release 2 (HPRC2): our second draft of the human pangenome, as part of a planned three-phase project to construct a publicly shared^32^, freely available global pangenome reference. These data are shared openly as a public domain resource. Of note, HPRC2 includes contributions from the Human Pangenome Project (HPP), an international effort to build an integrated human pangenome resource, to which HPRC contributes as a partner. Taken together, these improvements provide broader, more uniform, and deeper access to human genetic variation relative to HPRC1. Along with developments of pangenomic tooling demonstrated here and elsewhere, this represents a meaningful step towards an updated reference resource that better represents human genetic diversity.

### Capturing greater than 99% of common variation

For HPRC2, we targeted an approximately fivefold expansion in the total number of included genomes, selecting additional samples predominantly from the 1000 Genomes Project (1000G)^33^. To maximize the capture of common genetic variation, we developed an iterative greedy algorithm, MaxVar, that utilized the pre-existing 1000G short-read based genotype data to rank the selection of individuals for inclusion. MaxVar operates by maximizing the coverage of common biallelic variants across iterations (**Fig. 1a**), where coverage is defined as the presence of at least one individual in the reference panel who carries the alternate allele. Here, and in what follows, common is defined as an allele frequency (AF) ≥ 1%.

**Figure 1:**
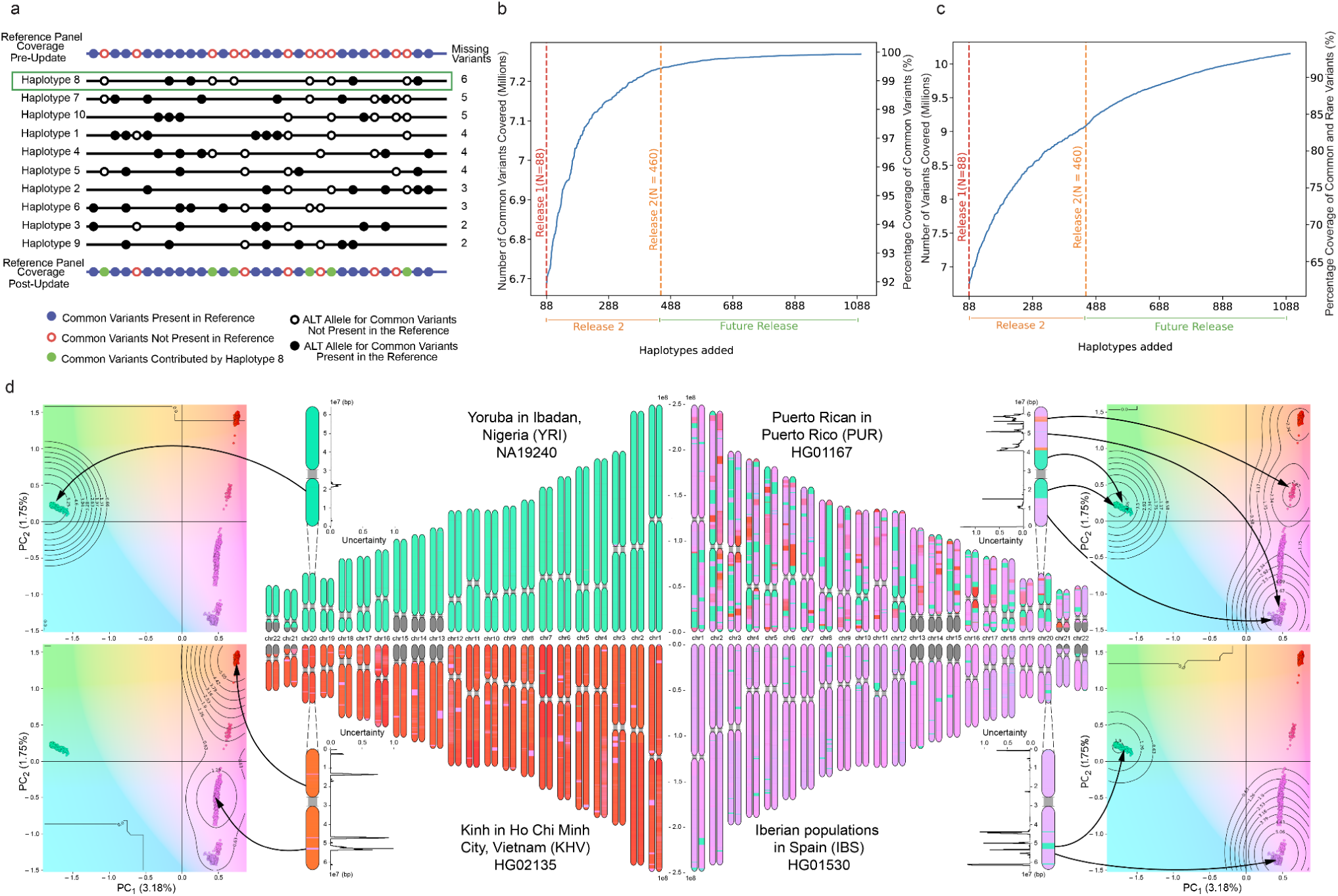
Selecting sufficient samples to capture common variation and providing local ancestry labeling. **a.** Schematic of single MaxVar iteration. The top track displays common variant coverage in the reference panel prior to iteration, with solid blue circles denoting variants already covered in HPRC1 and open red circles denoting variants absent from that reference. Haplotypes in the selection pool are ranked in descending order by the number of uncovered common variants they carry – that is, their marginal contribution to reference coverage upon addition. Open and solid black circles represent alternate alleles for common variants absent from and present in the HRPC1 reference, respectively. Haplotype 8, ranked first with six potential novel contributions, is selected and added for sequencing to the new HPRC2 reference. The bottom track shows the updated coverage following the iteration, with the six newly covered variants highlighted as solid green circles. **b.** MaxVar coverage of common small variants in the AoU Research Program cohort (N≈440,000). The x-axis represents the cumulative number of haplotypes added to the reference panel, and the dual y-axes indicate the absolute number (left) and the percentage (right) of common small variants covered. Vertical dashed lines mark HPRC1 (N=88 haplotypes) and HPRC2 (N=460 haplotypes). Coverage to the right of the orange dashed line reflects projected estimates generated by continuing to run MaxVar beyond HPRC2. **c.** As in Fig. 1b, but instead showing MaxVar coverage of common and rare small variants (AF ≥ 0.1%) in the AoU Research Program cohort. **d.** Genome-wide continuous local ancestry coordinates inferred by PCLAI. PCLAI regresses per-window genomic variation onto a continuous reference PCA space, producing ancestry coordinates per genomic window in PC1-PC2 space together with a per-window uncertainty score. For four 1000G individuals (YRI NA19240, KHV HG02135, PUR HG01167, and IBS HG01530) chromosome ideograms paint each window with coordinates using an interpolated CIELAB colormap derived from PCA position enabling visualization of continuous variation by PCLAI along each chromosome. PCA panels display the reference space (each dot is a reference in 1000G) and contours that illustrate the genome-wide distribution of predicted window locations for the individual, while accompanying uncertainty summaries for a haplotype from Chromosome 20 shows confidence across the windows with higher uncertainty typically concentrated in windows at recombination breakpoints between regions painted with different genetic ancestry.

To evaluate MaxVar’s performance, we quantified common small variant (small variant being either a SNP, or short insertion or deletion (indel) less than 50 bases in length) coverage using short-read whole-genome sequencing data from the All of Us Research Program v8 (AoU; N≈440,000). We identified 7,274,300 common small variants in AoU, of which 6,686,383 (91.92%) were covered by HPRC1 (88 haplotypes; **Fig. 1b; Supplementary Fig. 1**). Incorporation of additional haplotypes using MaxVar leads to a steep initial increase in common small variant coverage, followed by diminishing returns as the remaining uncovered variants become increasingly rare or population-specific. HPRC2 (460 haplotypes total) expands coverage to 7,233,162 (99.43%) of these common small variants, representing a substantial improvement in the representation of this large U.S.-based biobank. Extrapolating along the observed trajectory suggests that the common variant space in AoU approaches saturation with continued reference expansion, with inclusion of ≥99.9% of common small variants projected to occur once 932 haplotypes are included. We further characterized coverage at the individual level by quantifying, for each AoU participant, the number and proportion of common variants absent from our pangenome reference for which the individual carries at least one copy of the alternate allele. The distribution of this residual missingness is shown in **Supplementary Fig. 2**, highlighting heterogeneity across individuals and genetic ancestry. We repeated the variant coverage trajectory analysis in AoU but instead using an AF cutoff of ≥0.1%, which therefore includes both rare (but not ultra-rare) and common small variants (**Fig. 1c**). The HPRC2 includes 83% of such variants, increasing from 62% in HPRC1. Future pangenomes well beyond a thousand haplotypes will be needed to achieve >99% coverage of such variants.

To provide chromosomal segment level ancestry (local ancestry) estimates for the selected genomes, we applied Point Cloud Local Ancestry Inference (PCLAI)^34^, in which continuous genetic ancestry coordinates are computed per haplotype window along each chromosome of each sample by regression into a reference PCA space (PC1-PC2). That PCA space is here spanned by the 1000G samples that were not used in the HPRC2 dataset (**Fig. 1d**). Across the autosomes, each window is assigned a coordinate pair via PCLAI and painted on the chromosome ideograms using an interpolated CIELAB colormap from the PCA space. Accompanying per-window uncertainty scores summarize confidence in each local ancestry estimate.

To support categorical analyses of local ancestry when needed, a four class discretization of this continuous PCA space was generated. Each of these four classes map closely with existing 1000G continental ancestries. Indeed, **Supplementary Fig. 3** shows strong concordance between these discretized PCLAI estimates and results from a standard unsupervised genetic ancestry clustering method (ADMIXTURE)^35,36^.

The left side of **Fig. 1d** illustrates PCLAI for the Yoruba in Ibadan, Nigeria (YRI) individual NA19240 and for the Kinh in Ho Chi Minh City, Vietnam (KHV) individual HG02135. PCLAI produces a quasi-uniform coordinate profile across chromosomes with windows projecting to a single region of the reference PCA space. This pattern is accompanied by a low uncertainty distribution, consistent with stable local ancestry positioning across the genome. By contrast, the Puerto Rican in Puerto Rico (PUR) individual HG01167 on the right exhibits a mosaic-like pattern of local coordinates across chromosomes. Windows alternate between multiple regions of the PCA reference space, resulting in frequent shifts in the color-encoded chromosome painting. The uncertainty distribution broadens relative to the less heterogeneously labeled genomes, with elevated uncertainty concentrated in windows that fall between dense reference point clouds (recombination breakpoints between haplotypes). Finally, the Iberian populations in Spain (IBS) individual HG01530 has haplotypes concentrated in a PCA region with other European (EUR) 1000G superpopulation samples, but with sporadic haplotypes shifting toward a PCA region shared with many African (AFR) 1000G superpopulation samples.

### Embedding ELSI in HPRC2

Embedded within HPRC is an ethical, legal, and social implications (ELSI) component to consider, among others issues relating to population descriptors and representation (**Supplementary Table 1** gives an overview). We sought to prioritize samples based on direct measures of genetic variation rather than proxies such as racial, ethnic, national, or continental categories^37^. Regarding labelling, each sample keeps its unchanged primary label (e.g. ASW, MXL), meaning the descriptors that were originally assigned by 1000G. We aggregate primary labels into secondary labels (e.g. “continental” or “superpopulation” groups) only where an aggregate analysis is necessary to answer specific scientific questions. In such cases, secondary labels are mapped back to primary labels for transparency.

The PCLAI-derived secondary labels described in the preceding section represent inferred shared genetic ancestry, not identity. Following US National Academy of Sciences guidelines, we therefore refer to the discretized PCA regions as European-like (EUR-like), South Asian-like (SAS-like), African-like (AFR-like), and Indigenous American–East Asian-like (AMR/EAS-like), reflecting genetic similarity to these 1000G sample groups, as aggregated under secondary labels presented in Byrska-Bishop et al^38^. A direct mapping between 1000G participants and these secondary labels is provided in the Supplementary Materials. The actions taken and resulting tools, including ad hoc approaches to labeling developed as the analyses reported in this manuscript were finalized, are summarized in **Supplementary Tables 2,3,4**.

### Assembling thousands of complete, T2T chromosomes

To assemble the 230 genomes included in HPRC2, we produced Pacific Biosciences (PacBio) HiFi, Oxford Nanopore Technologies (ONT), and Illumina Hi-C sequencing data (**Fig. 2a**) from lymphoblastoid cell lines (LCLs) of the samples selected by MaxVar (171 total). HPRC1 samples (47 total) were rebasecalled for PacBio and additionally sequenced with ONT when necessary to create a uniform dataset. Importantly, the HPRC worked with two HPP partner sites, the University of Tokyo^39^ and Human Technopole, to incorporate data and assemblies generated parallel to the HPRC production centers into HPRC2. In total 14 samples were sequenced by HPP collaborating institutions and integrated into the HPRC2 dataset through harmonization of sequencing coverage, basecalling, and quality-control procedures across the entire HPRC2 dataset. This represents a successful proof of concept and provides a model for data integration across multisite and international collaboration on combined pangenomic references like those in the HPP.

**Figure 2.**
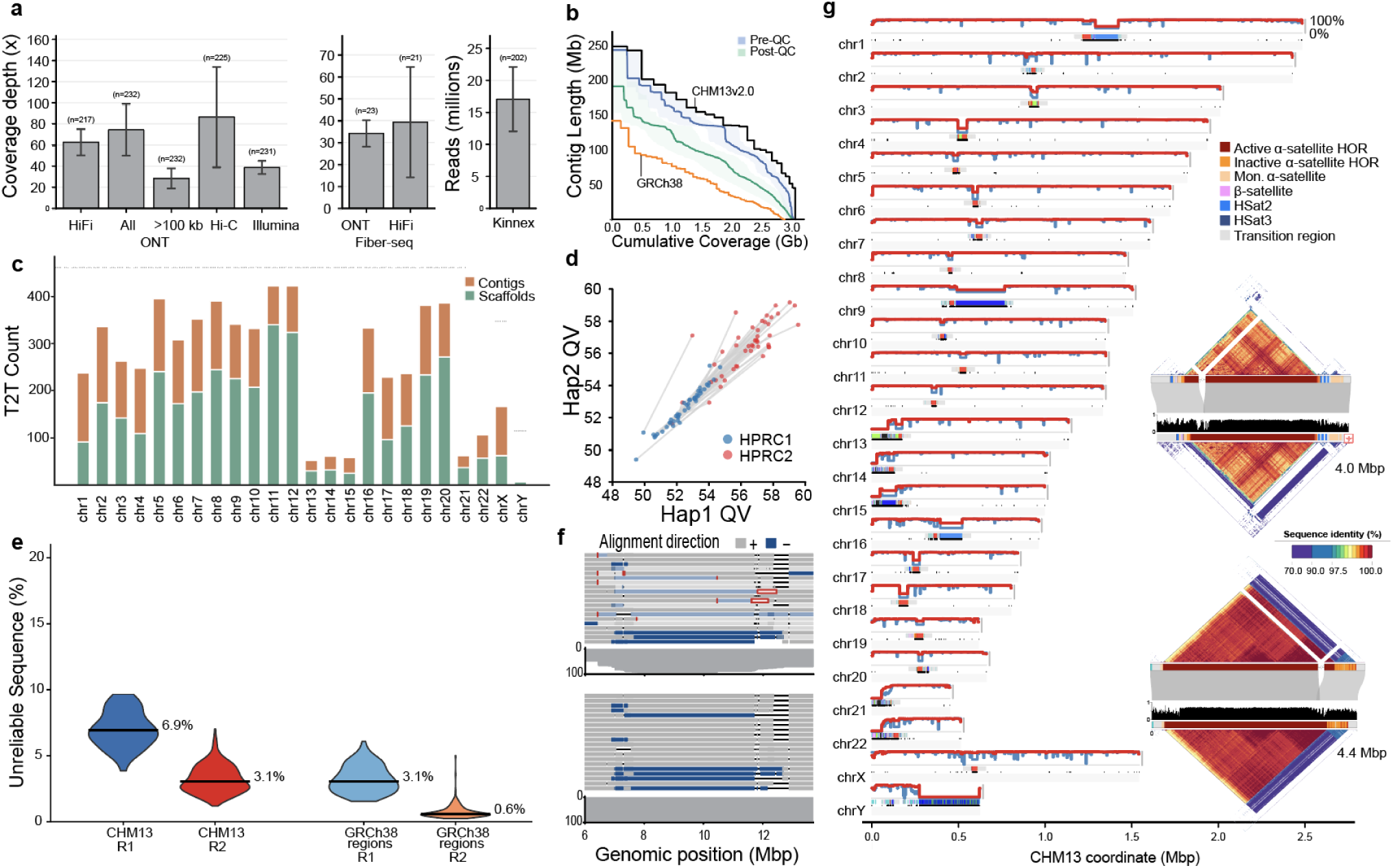
| Sequencing and assembly of 230 individuals create well-resolved regions that were previously missing. **a**, Sequencing coverage for data released in HPRC2. Sequencing coverage was calculated as raw bp divided by an estimated genome size of 3.1Gbp; error bars represent the standard deviation of coverage. **b**, NGx plot showing HPRC2 assembly contiguity for the assemblies (blue) and the assemblies after splitting sequences at regions that fail QC. Solid lines represent the median and bands representing the 10th-90th percentile range show variability across the assemblies. T2T-CHM13 v2.0 and GRCh38 split on gaps are included for reference. **c**, Count of T2T representation in HPRC2. Reference-level samples assembled outside of the HPRC2 effort (CHM13, HG002, HG06807) were excluded, leaving 115 male and 115 female samples. Maximum count lines (dotted) are placed at 460 for autosomes, 315 for chrX, and 115 for chrY. **d**, Estimated assembly base-level error (QV) from Illumina k-mers for HPRC1 assemblies and matched assemblies in HPRC2. **e**, Unreliability in HPRC2 assemblies as a percent of current reference genomes. Genomic windows of the reference are deemed unreliable for each assembly if there are either no sequences spanning the region, or if the assembly representation of that region has structural errors predicted. **f**, Assembly representation of recurrent inversions and microdeletions at 8p23 in HPRC1 (top) and HPRC2 (bottom). Grey boxes represent alignment on the same strand as T2T-CHM13, blue boxes represent inversions. Light grey and blue boxes indicate regions with multiple contigs or scaffolds in the assembly that represent the region. **g**, Genome view of unreliable regions in HPRC1 and HPRC2. (Inset) Example chromosome breaks in HPRC1 assemblies which are well assembled in HPRC2.

HPRC2 also includes functional data from the source LCLs to enable characterization of the transcriptional and epigenetic landscape of these genomes and their variations. Since the first release, PacBio and ONT sequencing data now include predictions of 5-methylcytosine (5mC)^40–42^, and we additionally generated long-read transcripts for 206 samples (PacBio Kinnex^43^) and long-read chromatin accessibility for 38 samples (Fiber-seq^44^). By pairing deep, high-quality sequencing datasets and assemblies with methylation, long-read RNA, and chromatin accessibility data, HPRC2 represents a resource for functional interpretation of the newly resolved regions. Sample information can be found in **Supplementary Table 5** and per-sample sequencing data and coverage are summarized in **Supplementary Table 6**.

Chromosome-scale, haplotype-resolved assemblies were produced with Hifiasm^5,45^ using HiFi and ONT Ultra-Long (UL) inputs for assembly graph construction with either Illumina sequencing of the parents (termed trio) (126 out of 230) or Hi-C sequencing (104 out of 230) used for haplotype phasing. We observed no differences in contiguity or completeness between the trio- and Hi-C-based assemblies but a small, statistically significant difference in base-level correctness (QV trio=57.09 vs. Hi-C=56.04; **Supplementary Fig. 4**), further confirming that parental data are no longer required for haplotype-resolved assembly^46^. Assemblies were polished with DeepPolisher^47^ to improve base-level quality, then extraneous sequences such as EBV and read adapter sequences were removed, and mitochondrial contigs were re-assembled with a dedicated pipeline. Details can be found in the Methods (and **Supplementary Fig. 5**). HPP samples were assembled by their source institutions and harmonized with HPRC assemblies using the HPRC post-assembly workflows. HPP assemblies had slightly higher contiguity and predicted error sequence, but were similar to HPRC assemblies in completeness metrics (**Supplementary Figs. 6, 7**). An experimental set of Verkko^4^ assemblies were also generated for all HPRC samples to facilitate analyses of Chromosome Y^48^ and the short arms of acrocentric chromosomes^49^, which Verkko is specially designed to scaffold using Hi-C data (**Supplementary Table 7**). These supplementary assemblies are described in a companion effort.

Structural quality was assessed using a combination of HMM-Flagger, run separately with HiFi and ONT sequencing reads^50^, and NucFlag, run with HiFi reads^51^. In order to reduce one-off false positives, erroneous regions were defined as any region with an assembly error predicted by two of the three different structural QC evidence streams. Area Under N50 (AUN) analysis shows that HPRC2 assemblies approach the contiguity of T2T-CHM13 (a theoretical maximum), and the gap-free and flagged error-free regions of the HPRC2 assemblies both have higher AUN than those of the Genome Reference Consortium’s (GRC) GRCh38 reference (**Fig. 2b**). Thus, the validated regions of each individual HPRC2 assembly are more continuous, on average, than GRCh38.

HPRC1 relied primarily on HiFi for assembly, and ONT UL for validation. With improved assembly methods that combine HiFi and ONT UL during the assembly process, automated creation of T2T chromosomes is now possible^5,52^. HPRC2 includes 3,666 chromosomes (35%) assembled T2T without gaps (termed T2T contigs) and an additional 2,490 chromosomes (24%) assembled T2T with one or a few unresolved gaps (T2T scaffolds) (**Fig. 2c**). Acrocentric chromosomes and chrY show markedly lower T2T counts principally due to their long satellite arrays while over 68% of other chromosomes are assembled as either T2T contigs or scaffolds. Base-level quality (QV) estimated using k-mers from Illumina reads show low error rates, with an average QV of 55 (1 error every ∼350,000 bases), a marked improvement compared to HPRC1 with a predicted QV of 52.5 by the same analysis (1 error every ∼175,000 bases) (**Fig. 2d, Supplementary Table 8**). Data from the HG002 reference sample, for which a complete, nearly-perfect T2T assembly already exists^6^, was downsampled to match HPRC2 coverage metrics and included as a control. The Illumina k-mer-based QV estimate of the HG002 control assembly matches well to the alignment-based measurement against the HG002 reference (k-mer QV=54.2, alignment-based QV=51).

To assess the number of high quality local reference haplotypes at each region across the genome, we projected assemblies onto CHM13 (**Supplementary Figs. 8,9**). To aggregate across assemblies we tiled T2T-CHM13 into 500kb syntenic windows (satellite arrays such as rDNA and alpha satellites which were larger than 500kb and regions which lacked clear synteny to T2T-CHM13 were expanded to form larger windows with clear bounding alignments) and counted the number of haplotypes that cover the genomic windows without assembly gaps or error flags. As compared to calculating the sum of QC flagged bases from individual tools, this approach increases the fraction of the genome tallied as unreliable (because a flag or break in a window marks the whole interval in the given haplotype), but better reflects the fact that assembly gaps and modestly sized error flags can create large regions unusable for analysis. Genome wide, HPRC2 has 3.1% of regions per haplotype estimated to be unreliable as compared to 6.9% in HPRC1. Subsetting to regions present and well assembled in GRCh38, only 0.59% of regions across all assemblies are estimated to be unreliable (**Fig. 2e**). As a result, HPRC2 accurately and contiguously assembled hundreds of diverse haplotypes across the genome including in regions with well known structural diversity. However, the most difficult regions – namely, acrocentric p-arms and the repeat-heavy q-arm of chromosome Y (all of which are absent from GRCh38) – are still typically not assembled accurately in HPRC2. These regions will be the target of further efforts in a future release. Also notably, analysis of LCL telomere lengths using long-read data suggests that LCLs do not accurately represent telomere length in primary tissue (**Supplemental Note 1**, **Supplemental Fig. 10**), indicating that further work is needed to create a more representative reference of telomere lengths across tissues.

Prior analysis of HPRC1 (and other assemblies with HiFi but not ONT-UL) found that assembly gaps cluster near regions of large, highly identical, repeats and that polymorphic inversions flanked by those repeats were frequently misoriented^53^. ONT-UL data can help span these large repeats, and their inclusion into assembly generation improves resolution of these complicated genomic regions. Chromosome 8p23.1 showcases these assembly improvements; the region harbors one of the longest known inversion polymorphisms in the human genome (∼4.1 Mbp; **Fig. 2f**), resulting in assembly gaps in nearly half of all HPRC1 haplotypes (45.7%) in this region. In contrast, all HPRC2 assemblies are gapless and predicted to be accurately assembled. Importantly, of the 94 haplotypes in HPRC1, only 17 of them had detectable inversions in the HPRC1 assemblies, due to assembly gaps obscuring the inversion, while 35 of them had detectable inversions in the HPRC2 assemblies. This is the result of including ONT UL data in HPRC2 assemblies, which proved crucial for assembling the large blocks of segmental duplications (SDs) flanking the 8p23.1 inversion. We further validated the correct representation of the 8p23.1 inversion using Strand-seq data^54^ in 29 out of sampled 30 (97%) assembled haplotypes (**Supplementary Fig. 11**).

Similarly, centromeres were a region that were consistently broken in HPRC1 (**Figure 2g, inset**), with many of the breaks and misassemblies associated with the centromere dip region (CDR)^55,56^. Assessing centromere active arrays, 15% are predicted to be contiguous and accurately assembled in HPRC1 while in contrast 66% of centromeres (6,946 total) in HPRC2 assemblies are similarly estimated to be complete and accurate ^57^. Given the substantial expansion in samples, this provides thousands of centromere complete arrays across the set for analysis.

### An Expanded Map of Haplotype Resolved Variation

To relate the assembled genomes to each other we constructed pangenome alignments using the Minigraph^58^, Minigraph-Cactus (MC)^59^ and new Implicit Pangenome Graph (IMPG)/Pangenome Graph Builder (PGGB) pipelines^60,61^. In each case these generalized multiple sequence alignments, which represent all forms of variation, are encoded as variation graphs^10^. The Minigraph graph is constructed starting from a chosen linear reference (we separately used both GRCh38 and T2T-CHM13) by iteratively adding the additional haplotypes (including alternative references) to construct a pangenome graph at the resolution of structural variants (i.e. missing small variants). These graphs are useful for visualization and mapping structurally variable loci. The MC graphs are a base-level resolution refinement of the Minigraph alignments, built to include both small variants as well as structural variants and largely excluding non-reference centromeric and satellite (censat) sequences that are not reliably comparable by the approach. We refer to the non-reference sequence included in the MC graphs as alignable, and note that this approximately corresponds with the euchromatic portion of the added pangenome haplotypes. By providing a complete, base-level map of all the alignable sequences these graphs are useful for analyses requiring a comprehensive set of common variation and are amenable to mapping based analyses. The IMPG alignments represent an alternative, reference free representation of a large subset of the possible pairwise alignments between the haplotypes, facilitating extraction of relevant subsets of pairwise alignments between the genomes without explicitly materializing the variation graph. The IMPG alignments cover all sequences, including censat sequences. This makes them useful for analyzing variation in regions where a single multiple-sequence alignment is too uncertain but a collection of independent pairwise alignments can usefully relate the sequences. To convert the IMPG pairwise alignment set into a variation graph, the PGGB graph pipeline was run on the set of pairwise alignments. Given the previously demonstrated similarity between the MC and PGGB alignments across much of the genome outside of censats and acrocentric short-arm sequences^13^, we focus on reporting the GRCh38-based MC results apart from where noted.

Starting from the GRCh38 autosomes, we charted pangenome growth with the addition of extra haplotypes (**Fig. 3a, Supplementary Figs. 12, 13**). In comparison to HPRC1 alignments, the amount of sequence observed in at least two haplotypes increased by 42 Mb and the amount of sequence occurring in at least one haplotype increased by 128 Mb. In total we catalog 271 Mb of non-reference alignable sequence, with approximately 106 Mb of such sequence deemed common, that is present at ≥1% AF. This is a slight reduction in the estimate relative to HPRC1^13^, likely due to reduced estimation error and changes in alignment parameters. Agreeing with the MaxVar analysis, this apparent saturation suggests almost all common variants are represented in the HPRC2 graphs, and that the HPRC2 graphs represent a considerable expansion in the representation of rarer polymorphisms. In fact, an intrinsic analysis using incidence-based sample coverage^62^ shows that with 98.77% probability, any randomly chosen variant from a new sample is already contained in the graph (**Supplementary Fig. 14**). To define the structure of the core pangenome we examined multi-MUMs, which are maximally-sized exact substring matches across a pangenome that are unique within each genome, and thus represent a set of conserved sequences shared among all haplotypes^63,64^. Multi-MUM coverage measures the proportion of reference bases contained in conserved multi-MUM syntenic blocks. Considering the complete HPRC2 set, >70% of the pangenome is syntenic with T2T-CHM13 (**Supplementary Fig. 15**), compared to ∼85% in HPRC1, with this core measure of overall synteny steadily decreasing as the sample size grows due to the capture of structural variants (SVs).

**Figure 3.**
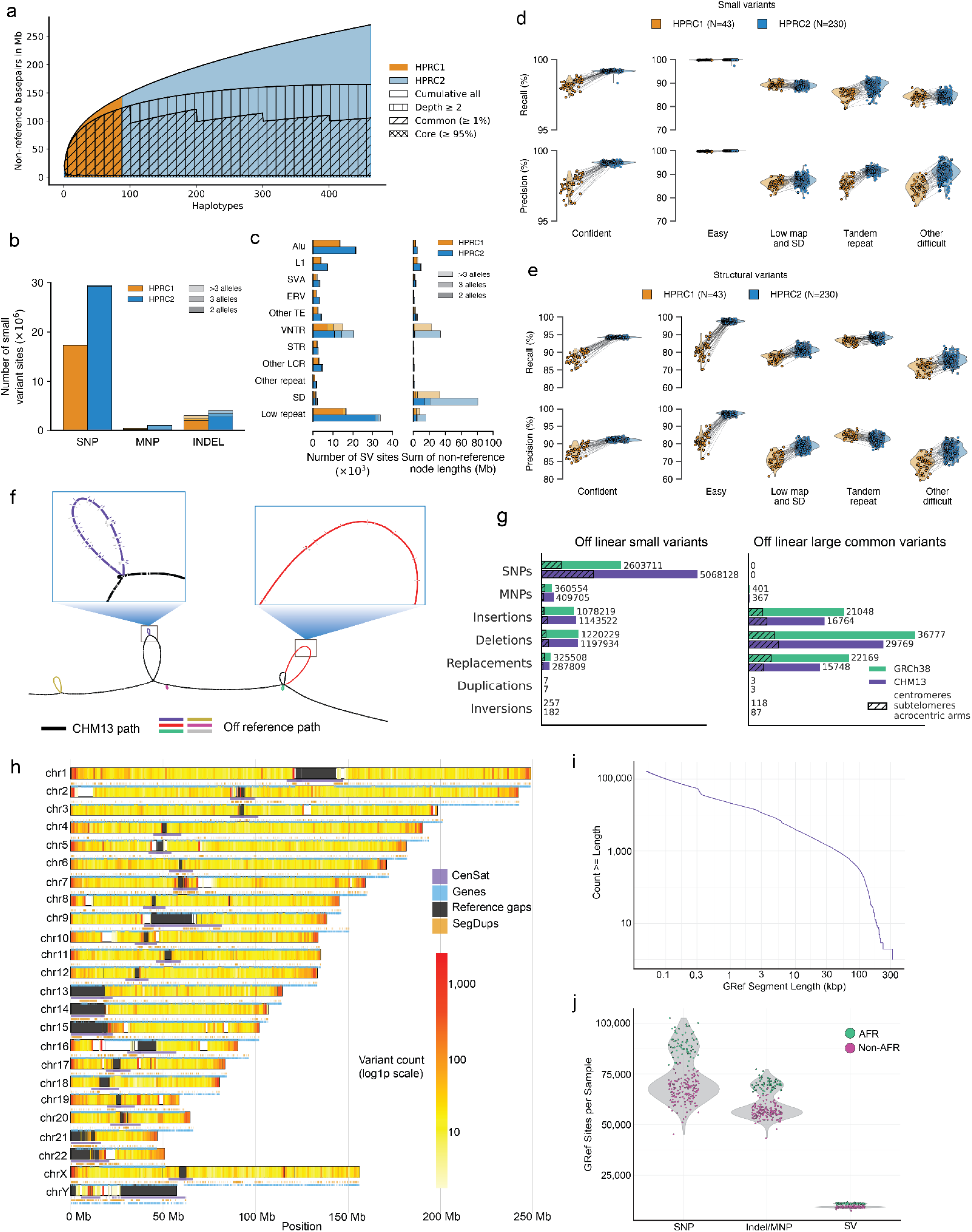
- An integrative map of common variation. **a.** Growth plot for autosomal non-reference base pairs in the HPRC2 graph, starting from the HPRC1 haplotypes. The seamless transition between the two groups implies that the HPRC1 and HPRC2 populations display a similar degree of genomic variability. **b.** Autosomal small variant sites in the HPRC1 and HPRC2 graphs, stratified by variant type and the number of alleles per site. MNP, multinucleotide polymorphism. **c.** Autosomal SV sites in the HPRC1 and HPRC2 graphs, stratified by repeat class and the number of alleles per site. The left plot shows the number of SV sites, and the right plot shows the sum of non-reference node lengths. Other TE, a site involving mixed classes of transposable elements (TEs). VNTR, variable-number tandem repeat, a tandem repeat with unit motif length ≥7 bp. STR, short tandem repeat, a tandem repeat with unit motif length ≤6 bp. Other LCR, low-complexity regions involving mixed VNTR and STR categories or low-complexity regions without a clear VNTR or STR pattern. Other repeat, a site involving mixed repeat classes. SD, segmental duplication. Low repeat, a site in which only a small fraction of the longest allele is annotated as repeat sequence. **d-e.** Precision and recall of autosomal small variants (d) and SVs (e) in the HPRC1 and HPRC2 graphs relative to per-sample joint ground truth callsets. Joint ground truth callsets were derived from merged callsets by retaining variants supported by at least two of 14 callers. Comparisons were restricted to dipcall confident regions and stratified by Genome in a Bottle (GIAB) v3.6 genomic context. Each point represents one sample, and matched samples between HPRC1 and HPRC2 are connected by lines. Low map, low mappability. SD, segmental duplication. **f.** Example of graph reference coordinates. The HLA-DRB1 region of T2T-CHM13 (black) in the T2T-CHM13-based HPRC2 MC graph, subset to four samples (NA19043, HG02056, HG01099, and NA20827) for illustration. GRef contigs, each representing an underlying haplotype sequence from an assembly, are shown in colours, with small nested variants highlighted in the inset. **g**. Left: Off-reference small variants identified by Pantree. Right: Off-reference SVs identified by Pantree. Results shown for both the GRCh38- and T2T-CHM13-based MC graphs. Variant edges with both end vertices not landing on the linear reference genome were extracted and classified into the following classes based on the tree reference allele length (r) and alternate allele length (a): SNPs (r=a=1), MNPs (r=a>1), insertions (r=0), deletions (a=0) and replacements (r≠a, a>0 and r>0). A variant is large if a or r is 50 bp or longer. Furthermore, a duplication results from a cycle and an inversion involves an edge connecting the forward and reverse strands in the oriented reference tree. Subtelomeres refer to 500-kb regions from the telomere of each chromosome arm and acrocentric arms correspond to the entire short arms of chromosomes 13, 14, 15, 21 and 22. **h.** Locations of GRef contigs projected onto GRCh38 from the HPRC2 MC graph. Regions clipped by MC (>10-kb intervals with no alignment during graph construction) are shown in black. Censat, gene, and SD annotations are also shown. **i.** Length distribution of GRef contigs in the GRCh38-based MC graph. **j.** Off-reference sites of variation per sample as represented in the GRef VCF, with samples stratified into AFR and non-AFR populations (based on 1000G recruitment site) to show differences in variant counts.

We compared autosomal variant site composition between the HPRC1 and HPRC2 graphs (**Fig. 3b-c**; see **Supplementary Figs. 16, 17** for corresponding T2T-CHM13 analyses). As expected from the larger number of samples represented in the HPRC2 graph, it contains more small variant sites across all variant types (34.49 million total small variant sites vs. 20.78 million in the HPRC1 graph), as well as more SV sites (109.16 thousand total SV sites vs. 65.90 thousand) and greater total non-reference node length across all SV classes (189.92 Mb vs. 90.78 Mb). In SD regions, however, the distribution of non-reference node length across allele-count classes differed between HPRC1 and HPRC2, with a larger biallelic contribution and a modestly smaller relative contribution from sites with more than three alleles in HPRC2. This difference may reflect the higher quality of the assemblies and/or alignments used in HPRC2, which might reduce spurious splitting of similar alleles into multiple groups in complex duplicated regions.

To establish concordance with existing variant calling methods, we benchmarked autosomal small variants and SVs in the HPRC1 and HPRC2 graphs against per-sample joint ground truth callsets. These callsets, jointly comprising small variants and SVs, were constructed by merging outputs from 14 variant callers (4 assembly-based and 10 PacBio HiFi read-based, all operating relative to the GRCh38 linear reference genome) and retaining variants supported by at least two callers (**Methods**). For small variants, the HPRC2 graph showed substantially higher overall recall and precision than the HPRC1 graph in confident regions (99.16% F1 vs. 97.87% F1), with the clearest gains observed in tandem repeat regions, whereas performance in easy regions was already near saturation in HPRC1 and remained similarly high in HPRC2 (**Fig. 3d**; see **Supplementary Fig. 18a**) for corresponding T2T-CHM13 analyses). These gains again likely reflect the assembly improvements as well as improvements in the alignment pipeline. For SVs, the HPRC2 graph also showed higher overall recall and precision than the HPRC1 graph in confident regions (92.64% F1 vs. 88.29% F1) and across most genomic contexts, with especially clear improvements in easy, low mappability, and SD regions (**Fig. 3e**; see **Supplementary Fig. 18b**) for corresponding T2T-CHM13 analyses). Only precision in tandem repeat regions was lower in HPRC2, indicating an increased number of false positives in this context. This may reflect the larger number of samples represented in the HPRC2 graph, which could increase sequence complexity in already-complex tandem repeat regions and make SV representation and comparison more challenging.

#### Mapping pangenome coordinates

Graphs give us an alternate window into the structural variation in the human pangenome. The novel sequence component of this variation, when viewed in the context of a linear reference, is predominately contributed by SV insertions. Although these inserted sequences can be effectively genotyped^28,29^, nested variation within SVs can become lost when projecting back to a linear reference. This is because, although nested variation within inserted sequence is visible within the graph, it cannot be directly expressed in a reference context, becoming hidden when projected to a reference missing the insertion. We address this by introducing the notion of pangenome reference coordinates, exploring two complementary strategies. The first, termed Pantredie^65^, constructs a minimum spanning tree over the graph to determine edges that can be indexed to refer succinctly to off-reference variants in standard variants formats, such as VCF^66^. The second, termed Graph Reference (GRef) coordinates, was inspired by rGFA^58^ as well as the alternate contigs of the Genome Reference Consortium’s assemblies^67^, and serves to augment a chosen linear reference with a set of contigs representing novel insertion sequences found in the graph (**Fig. 3f**). The result is a vertex-disjoint path cover of the graph, giving a set of contigs, each corresponding to a substring of an underlying haplotype assembly, that, with the starting reference, span the graph.

Using Pantree we count 2.6 million SNPs, 2.3 million indels and 80 thousand SVs that are off reference **(Fig. 3g**; the GRef approach gives similar numbers, see **Supplementary Fig. 19**). This represents a significant fraction (7.1% of SNPs, 23.0% of indels and 52.2% of SVs) of all variants within the alignable pangenome. Using the GRef coordinates, we analysed the location of non-reference contigs across the genome (**Fig. 3h**) as well as their length distribution (**Fig. 3i**) and allele-frequencies allele frequency distribution **(Supplementary Fig. 19**). We observe 162,807 such non-reference alignable off-reference sequences ≥ 50 bp in length with a cumulative length of 231 Mb, and, notably 879 such sequences ≥ 50 kb in length. As expected, their locations and frequencies mirror the expected distribution of SVs within the genome. Quantifying the number of off-reference variants per sample in the pangenome, using the GRef method and `vg deconstruct`, we find a median 79,088 SNPs (1.85% of SNPs per sample), 57,435 indels and MNPs (4.16%) and 9,517 SVs (11.65%) (**Fig. 3j**).

#### 12% of the human genome is segmentally duplicated in HPRC2

Because SDs are recognized as some of the most challenging regions of the genome to accurately assemble, we specifically assessed and validated SD content for 459 haploid genome assemblies from HPRC2 (>1 kb length and 90% sequence identity; Methods), as described previously^15,68,69^. Compared to previous phased genome assembly efforts^13,15^, we find that HPRC2 genome assemblies recover more SDs with greater contiguity (**Supplementary Fig. 20**). For example, we estimate 171.6 Mb of SDs per haplotype (**Fig. 4a, Supplementary Fig. 21a**) compared to 168.1 Mb (Human Genome Structural Variant Consortium Release 3 (HGSVC3)^15^) and 166.9 Mb (HPRC1). We validated the presence of novel SDs using short-read sequencing depth, excluding assembly errors as previously described^50,57,70,71^. We find that 94.6% of SDs are validated, with 93.5% and 95.4% validated for inter- and intrachromosomal SDs, respectively (**Supplementary Figs. 21, 22**). A small fraction (13.2%) of interchromosomal duplications that were >98% identical failed to validate in a subset of genome assemblies (**Supplementary Fig. 22**) suggesting the presence of residual artefactual SDs in phased genomes (12.5 Mb/haplotype across 46 assemblies). Consistent with earlier reports, 1000G AFR samples harbor significantly more and distinct repertoire of SDs than 1000G samples collected elsewhere (p=0.00068; **Supplementary Fig. 21b, Fig. 4b**)^15,69^. Restricting our analysis to validated SDs, we estimate that a remarkable 12% (382.7 Mb) of the human pangenome is segmentally duplicated. While a large proportion (63.0% or 241.1 Mb) is shared by more than 90% of human haplotypes, the results suggest an extraordinary degree of copy number and structural variation per genome, including novel SDs at new locations depending on the individual (Lin et al., companion manuscript) (**Fig. 4b, Supplementary Fig. 21c**). Only a small proportion of SDs (42.5 Mb) appears fixed in the human population (**Fig. 4b**), including novel protein-coding genes that show constraint in the human population (Ren et al., companion manuscript).

**Figure 4.**
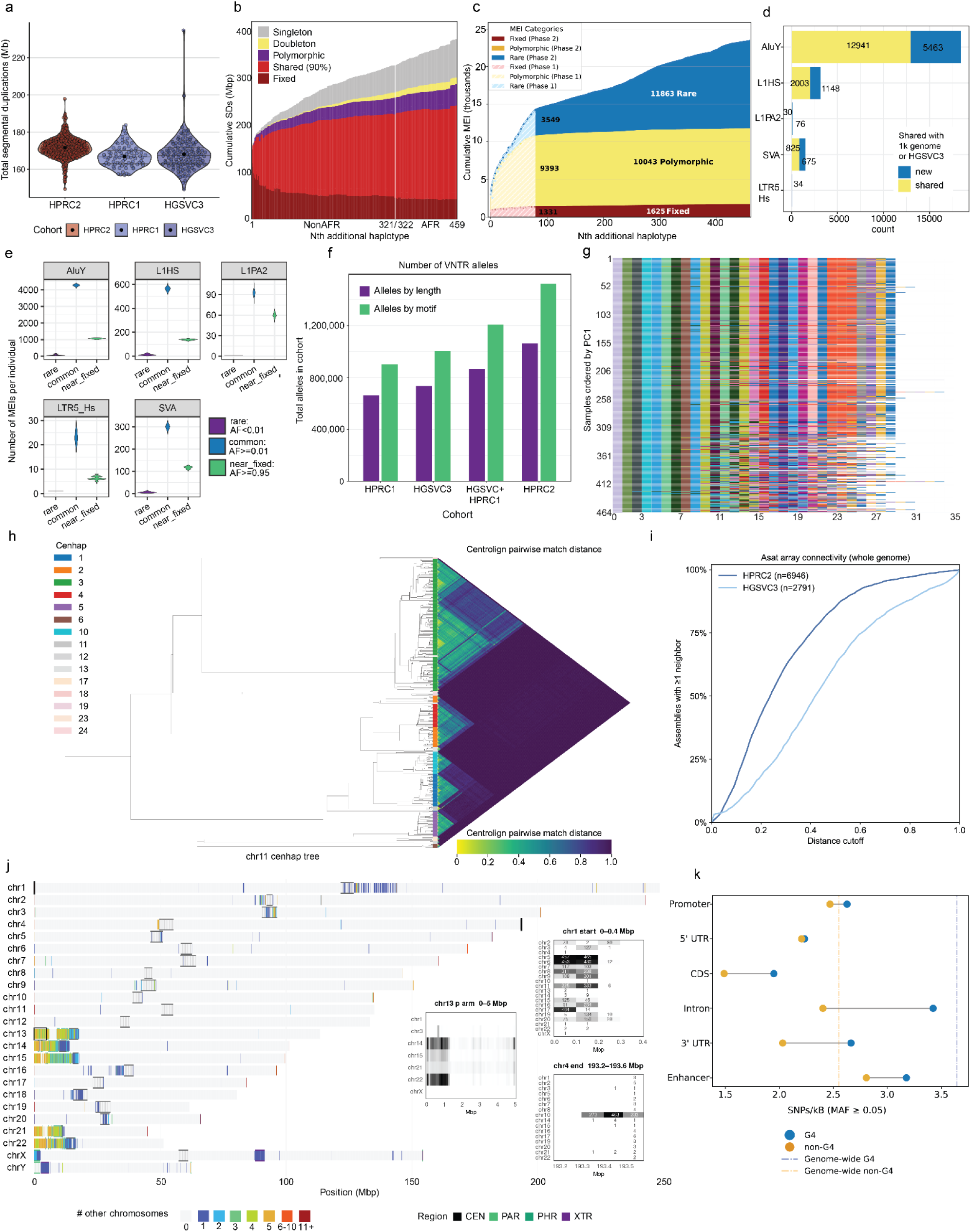
- Completing the map of common structural variation. **a.** Comparison of total number of SD bases per haplotype assemblies from HPRC2, HGSVC3^15^ and HPRC1^13^. **b.** Cumulative SD bases across HPRC2. The x-axis indicates haplotype assemblies (n=459), including T2T-CHM13 and excluding samples without short-read sequencing data to validate SD calls, ordered by samples from non-AFR 1000G populations followed by samples from AFR 1000G populations. The y-axis shows the cumulative sum of maximum length SDs. The quantification was performed excluding acrocentric p-arms and two subtelomeric 500 kbp end sequences. The singleton, doubleton, polymorphic, and shared categories are assigned by the allele counts/frequency (AC/AF); singleton: AC=1, doubleton: AC=2, polymorphic: AC>2, and shared: AF>0.9. Fixed SDs, indicated by dark red, represent SDs that are fixed by both allele frequency (AF=1) and length (standard deviation/mean < 0.1). **c.** MEI AF distribution in the human population, stratifying near_fixed (“core” - ≥0.95), common (≥0.01 and <0.95) and rare (<0.01) MEIs. **d.** Total number of polymorphic MEIs identified by HPRC2. The number of new MEIs were colored by pink while those identified before by the 1000G or HGSVC3 were colored by teal. **e.** Violin plots representing numbers of polymorphic MEIs in each diploid genome, stratified by AF between rare (<0.01), common (≥0.01), and near fixed (≥0.95). **f.** Number of alleles at variable tandem repeat loci for HPRC1 (N=95), HGSVC3 (N=130), nonredundant HPRC1 and HGSVC3 (N=218), and HPRC2 (N=464). Loci invariant in each cohort are excluded. The alleles by length only consider total copy number of all repeated motifs; alleles by motif consider distinct allele combinations. **g.** Waterfall plot of the *PLIN4* coding VNTR (chr19:4510838-4513560) on HPRC2 genomes, sorting the haplotypes by PCLAI principal component 1, which principally stratifies AFR-like from non-AFR-like genomes; the 1000G AFR populations exhibits the highest allele length and compositional diversity while the 1000G EAS populations exhibits the lowest. **h.** Centromere haplotype tree for HPRC2 chr11, with colors indicating the centromere haplotype assignment of each haplotype. Heatmap contains the Centrolign pairwise match distance for the alpha satellites between each sample pair. **i.** Relatedness of alpha satellite arrays. All pairs of alpha satellite arrays passing QC for each chromosome were aligned against each other using Centrolign, allowing calculation of a pairwise match distance metric (methods). Curve represents the percent of assemblies in each study (HPRC in dark blue and HGSVC in light blue) with greater than one neighbor assembly within the distance cutoff on the x axis. **j.** Inter-chromosomal matches across the T2T-CHM13 genome. Left: karyogram of T2T-CHM13 v2.0. Each 100 kb window is coloured by the number of other chromosomes from which at least one HPRC haplotype contig reaches that window: grey, none; blue to dark red, 1 to 11+. Overlaid tracks mark centromeres (CEN, black), pseudoautosomal regions (PAR, orange), acrocentric pseudo-homologous regions (PHR, green) and the X-transposed region (XTR, blue). Black boxes on the karyogram mark the three regions shown to the right as grey-scale heat maps: chr1p tip (0–0.4 Mbp, top right), chr13 short arm (0–5 Mbp, middle) and chr4q tip (193.2–193.6 Mbp, bottom right). In each heat map, cells show the number of haplotypes supporting the other-chromosome matches (row) in each window (column); darker cells mean more haplotypes. **k.** The orange and blue dots show the SNP frequency per kb at G4 and non-G4 sites, respectively, for each of the functional categories shown on the y-axis. The vertical orange and blue dashed lines indicate the genome-wide SNP frequency per kb for G4 and non-G4 sites, respectively. The horizontal black lines connecting the orange and blue dots, highlight the magnitude of SNP frequency difference between G4 and non-G4 sites.

#### Cataloging polymorphic mobile element insertions

We annotated mobile element insertions (MEIs) within the set of SV indels. We considered all polymorphic MEIs found in at least one haplotype, but not fixed in the human population^72^. Growth analysis with increasing cohort size indicates that we capture almost all common MEIs (AF ≥ 1%) and many rarer MEIs (Fig. 4c). In total, we identified 23,531 such MEIs, including 18,597 Alu, 3,260 L1, 1,500 SVA, and 77 LTR elements; ∼30% of these MEIs are described for the first time in HPRC2 (**Fig. 4d**). Analysis at the superpopulation level confirms in this large set that the AFR superpopulation harbors the greatest diversity with the highest number of rare MEIs (**Supplementary Fig. 23**). Analyzing AF by family, the modal class of Alu, L1HS, and SVA insertions were singletons (34% of all events), suggesting a recent mutational origin (**Supplementary Fig. 24**) while, as expected, most L1PA2 insertions are ancient and fixed^73^. Although most (52.4%) individual MEIs in the pangenome are rare, >98% of polymorphic MEIs per individual are common, with an average of 5,246 per individual (4,268 AluY, 563 L1HS, 92 L1PA2, 300 SVA, and 23 LTR5), relative to an average of 82 rare MEIs per individual (63 AluY, 12 L1HS, 1 L1PA2, 6 SVA, and 1 LTR5) (**Fig. 4e**). Furthermore, a substantial minority (26%) of common MEIs are nearly fixed (AF ≥ 0.95) (mean 1384 per individual: 1062 AluY, 139 L1HS, 60 L1PA2, 117 SVA, and 6 LTR5), with the exception of older L1PA2 elements, where the large majority (∼65%) of common MEIs are nearly fixed (Jiang et al., companion manuscript).

#### Assembly of tandem repeats provides sequence information beyond copy number estimates

Tandem repeat (TR) sequences account for the largest class of SV by count per genome^15,74^. Comparing variable-number tandem repeat (VNTR) loci in HPRC2 with earlier cohorts **(Fig. 4f**), the total number of alleles observed increased with cohort size, increasing 60% over the HPRC1 and 68% when considering motif composition, indicating that previous studies did not reflect saturation of VNTRs in populations. In a companion study we used vamos^75^ to generate a catalog of sequence-resolved variation for 4.2 million TR sequences measuring repeat copy-number (length) variation as well as inter-repeat (motif) polymorphisms^76^. Highlighting the value of having sequence assemblies rather than just copy number estimates from k-mer histograms or read pileups, we found a 14% increase of loci that showed population differentiation (ANOVA test, **Supplementary Fig. 25**) when measured by motif composition compared to copy-number variation alone (247,570/215,835). This includes the *PLIN4* VNTR associated with skeletal muscular disorder^77^ that has rare expansions^16^ (**Fig. 4g**).

#### Representing centromere diversity

Centromeres have been excluded from genomic analysis due to the difficulties with assembling and aligning them, stemming from their length, structural diversity, high repeat content and absence of meiotic recombination^3,56^. The alpha satellite component of the centromere, which houses the binding site of the kinetochore during cell division, is particularly challenging to align due to its high level of local homogeneity^78^. Using Centrolign, a new tool specialized for aligning alpha satellite arrays, we constructed pairwise divergence estimates of alpha satellite array sequences (Eizenga, et al., companion manuscript).

We find high evolutionary turnover of these regions (for example, **Fig. 4h** shows the evolutionary relationship between haplotypes in centromere 11), such that only 42.5% of haplotypes have at least one neighbor sample within 20% sequence divergence in HPRC2, compared with 18.4% in HGSVC (**Fig. 4i**). This indicates that despite reaching saturation of common variation for much of the rest of the genome, improved assembly (34% of arrays were not predicted complete and accurate and included in this analysis) and more samples will be needed to adequately capture the diversity in the centromere. In a companion paper, we use Centrolign to develop the first variation graphs representing centromere pangenomes (Eizenga, et al., companion manuscript), a step toward their incorporation into the whole genome pangenome graphs.

#### Mapping subtelomeric interchromosomal rearrangements

Tracks of homology between chromosomes are rare and concentrate at a small set of well-known shared regions^79^. Using the IMPG alignments we tracked evidence of strong homology between chromosomes (Methods), providing a haplotype-resolved view of these events. As expected, the most widespread signal is on the acrocentric short arms (chr13, 14, 15, 21, 22), which share sequence almost exclusively among themselves (**Fig. 4j, chr13 p-arm inset**)^80^. We also observe the anticipated blocks of homology at the pseudoautosomal regions (PAR) between the X and Y chromosomes, and between the X-transposed region (XTR) onto the Y chromosome. However, the most extreme windows are subtelomeric, regions long known to be hot spots of interchromosomal recombination and segmental duplication^81,82^: the chr1 p-arm tip matches many other chromosomes (**Fig. 4j, top-right inset**), while the chr4 q arm tip is a near-exclusive match to chr10 q arm through the shared D4Z4 macrosatellite, which is linked to facioscapulohumeral muscular dystrophy^83^ (**Fig. 4j, bottom-right inset**). In a companion manuscript we demonstrate that these subtelomeric interchromosomal matches are associated with the exchange of telomere repeat arrays within the human germline^84^. In a second companion manuscript, we extend this analysis from telomere-repeat arrays into the adjacent subtelomeric sequence, providing a population-scale, haplotype-resolved map of polymorphic interchromosomal sharing. Pseudo-homolog regions occur on 41 of 48 chromosome arms and, after excluding the acrocentric short arms and sex-chromosome pseudoautosomal regions, encompass 3.51 Mb of subtelomeric sequence organized into 15 interchromosomal communities. Their sequence similarity is associated with three-dimensional proximity, and complete multigenerational pedigree assemblies identify putative recombination between non-homologous chromosome ends, supporting recurrent ectopic exchange as a mechanism maintaining this compartment^85^.

#### Predicted G-quadruplex secondary structures have consistently higher SNP rates

The expanded assembly collection across near complete genomes allows us to interrogate with less bias different categories of DNA sequence. For example, existing variant databases are primarily derived from short-read sequencing data, which are prone to errors in regions predicted to form non-B DNA structures^86^. Using a non-B DNA annotation pipeline, we found all haplotypes have a total non-B DNA motif content that lies in-between what is found on the previous human reference genome, hg38 (8.89%), and the present T2T reference genome T2T-CHM13 (11.50%, **Supplementary Fig. 26** and **Supplementary Table 9**), likely reflecting inter-individual variation and differing completeness of the assemblies. Using SNPs derived from the PGGB graph, we observed 377,395 SNPs present in a predicted G-quadruplex (G4), a type of non-B DNA, a substantial fraction of which are not present in gnomAD^87^ (6.2%) or did not map to the GRCh38 reference (19.3%) (fractions for all predicted non-B types are found in **Supplementary Fig. 27**). Interestingly, we observe consistently higher SNP densities in G4-forming regions relative to non-G4 regions genome-wide, and across functional elements, including promoters, genes, and enhancers, with coding sequences showing the lowest SNP occurrence rates and intronic regions the highest (**Fig. 4k**). The higher densities at G4 versus non-G4 regions in introns (which mostly evolve neutrally) likely reflects differences in mutation rates. However, the differences between G4s and non-G4 regions of non-intronic functional elements were much smaller (in particular for promoters and 5′ UTRs), suggesting selection operating at G4 sites^88^.

### Annotating the HPRC2 pantranscriptome

To generate a comprehensive annotation of all genes and transcripts within each haploid assembly of the HPRC2 cohort, we applied two complementary pipelines: CAT2 (an updated version of the Comparative Annotation Toolkit^89^), and the Ensembl pipeline^13^. To benchmark annotation quality, we compared transcripts called by the Ensembl and CAT2 pipelines across five biotype categories. The two pipelines showed strong concordance, particularly for well-characterized biotypes (**Supplementary Fig. 28a,b,c**). Across all assemblies, gene counts were highly consistent, with protein-coding gene totals clustering tightly around ∼20,000-20,200 for X-only containing haplotypes and ∼19,200-19,500 for Y-only containing haplotypes (**Fig. 5a**). Pseudogene and noncoding gene counts showed similarly narrow distributions, with T2T-CHM13 falling within the higher end of the observed range and GRCh38 occasionally appearing as an outlier, particularly for Y-containing categories. All but two haplotypes captured all expected single-copy protein-coding genes and >98% of noncoding genes, consistent with the high completeness. Each individual harbored an average of 101 (median 104; range 81-216) nonsense and frameshift mutations in the canonical transcript of protein-coding genes, consistent with prior estimates of Loss-of-Function (LoF) burden per human genome^87,90^. This supports that the assemblies and gene annotations capture coding variation at expected sensitivity without inflated artifactual calls.

**Figure 5:**
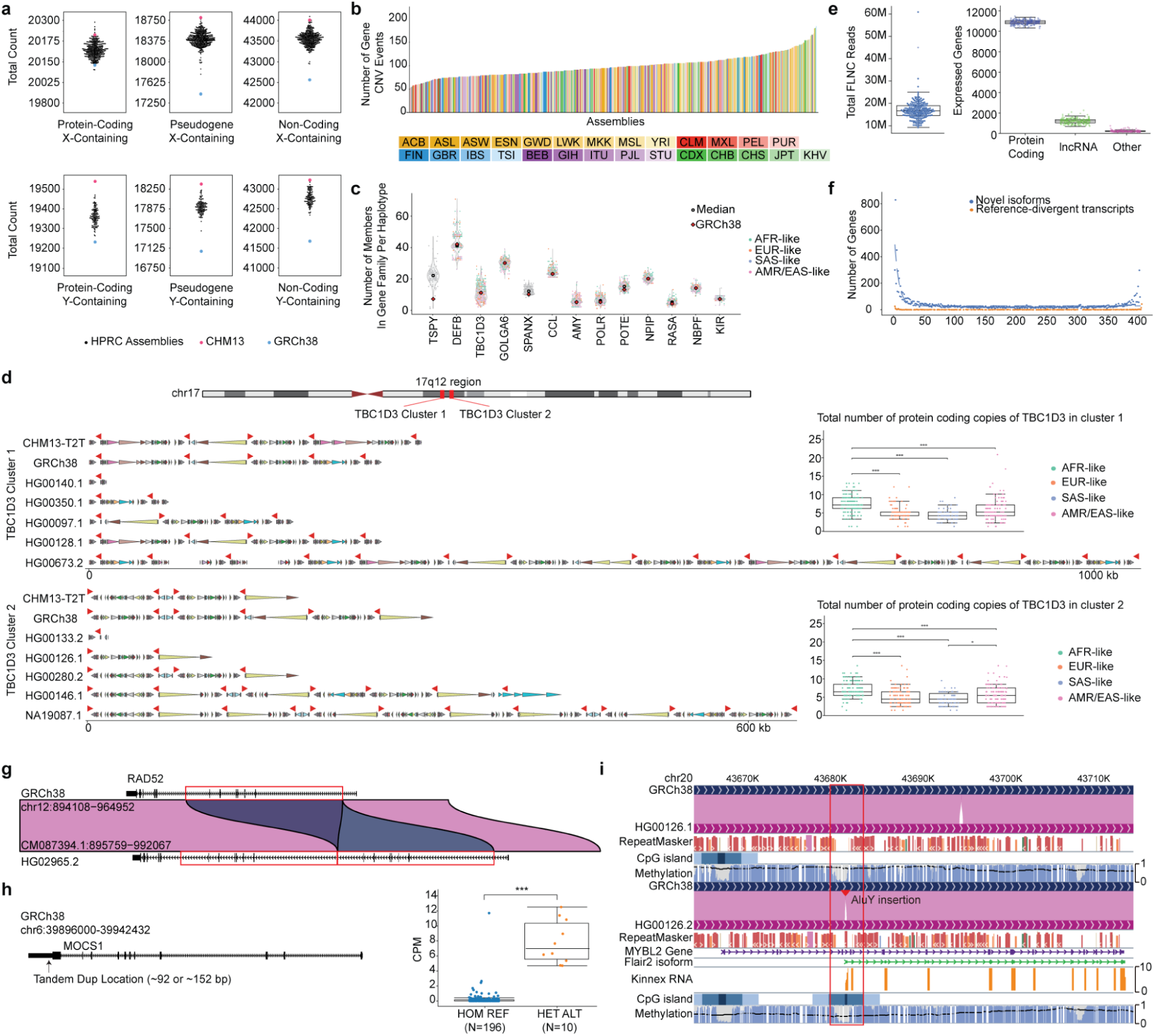
A pantranscriptome comprehensively capturing common haplotype resolved gene variations. **a**. Protein coding gene, pseudogene, and non-coding RNA counts, based on the CAT2 annotations, stratified by assemblies that contain an X chromosome vs those that contain a Y chromosome. Counts in GRCh38 (blue dot) and T2T-CHM13 (pink dot) shown. **b**. CNV events across each individual haplotype assembly coloured by population labels. For each gene family (see **Methods**), a deviation from the median count was considered as a variation and the summation of these variants (including both duplications and deletions) across all families for a haplotype is shown here. We are using a qualitative colour map to colour haplotypes where each colour family (yellow, red, purple, green, and blue) represent individual 1000G superpopulations and each shade within a colour family represents its subpopulation. **c**. Counts of individual genes across each of the haplotype assemblies for the most copy-number variable gene families. Each assembly is coloured by the PLCAI derived local ancestry for the locus of the gene family. **d**. Ideogram (top) shows the location of the two *TBC1D3* expansion blocks, Cluster 1 and Cluster 2 on chromosome 17q12. Below, haplotype-resolved representations of Cluster 1 (upper) and Cluster 2 (lower) are shown for the GRCh38 and T2T-CHM13 references alongside, for each cluster, the shortest confirmed configuration, the longest confirmed configuration, and the three most common configurations observed across the 437 QC-passing HPRC2 haplotypes. Red triangles denote individual *TBC1D3* protein-coding copies; coloured arrows below illustrate segmental duplication content annotated with DupMasker^97^. Box plots (right) show the total number of protein-coding *TBC1D3* copies per haplotype in each cluster, stratified by PLCAI-derived local ancestry. Scale bars indicate 1,000 kb (Cluster 1) and 600 kb (Cluster 2). **e**. Total number of reads, and gene-level expression across the 204 individuals with matched Kinnex data. **f**. The prevalence of genes having a novel isoform (blue) or a reference-divergent transcript (orange) across all of the 408 haplotype assemblies with matched Kinnex data. **g**. Reference-divergent transcript at *RAD52* in HG02965 (haplotype 2). Alignment of the HG02965.2 assembly (CM087394.1:895759–992067) to GRCh38 (chr12:894108–964952) reveals a genomic duplication (red boxes) spanning 6 exons in an internal region of *RAD52* that encodes the N-terminal single-stranded DNA binding domain. The resulting transcript carries a duplicated copy of this domain, altering the protein architecture relative to the reference isoform. **h**. Reference-divergent transcript in *MOCS1* 3’ UTR associated with elevated expression. Left: gene model of *MOCS1* (GRCh38, chr6:39896000–39942432) showing the location of a small tandem duplication (∼92 or ∼152 bp) within the 3’ UTR. Right: *MOCS1* expression (CPM) in individuals homozygous for the reference allele (HOM REF, N=196) versus those heterozygous for the duplication (HET ALT, N=10). **i**. HPRC epigenome browser view of the *MYBL2* locus on chromosome 20 in GRCh38 and the two haplotypes of HG00126, with tracks for sequence alignment, RepeatMasker, CpG islands, and PacBio methylation. HG00126.2 carries an intronic AluY insertion (red arrowhead, boxed) absent from GRCh38 and HG00126.1, which creates a hypomethylated CpG island and initiates a novel *MYBL2* isoform (green, FLAIR2^98^) supported by Kinnex long-read RNA-seq (orange).

We identified gene copy number variants (CNVs) across the HPRC cohort by comparing the copy number of gene families against the median gene count of that family across the entire HPRC set and the GRCh38 and T2T-CHM13 assemblies. Comparing against the median copy number of the set avoids the reference bias of comparing to the copy number of a single reference. Each phased haplotype harbored on average 126 gene duplication or deletion events, with a distribution ranging from approximately 50 to over 150 events per assembly (**Fig. 5b** (CAT2)**, Supplementary Fig. 29a** (Ensembl)).

Many of the most copy-number variable gene families were previously identified as CNV-enriched in HPRC1, but the broader and more complete set of HPRC2 provides finer resolution of individual-specific counts and population-specific patterns. Among the most copy-number variable multi-exon gene families (**Fig. 5c** (CAT2)**, Supplementary Fig. 29b** (Ensembl)), *TSPY* exhibited the widest range. Several other highly variable families, including *TBC1D3*^91^, *AMY*^92^, *KIR*^93^, and *NPIP*^94^, have been previously characterized as copy-number variable in population genomic studies, and our data recapitulate and extend those findings with haplotype-resolved resolution across a diverse cohort.

For several of these families, the GRCh38 reference copy number diverges substantially from the population median, falling near or below the lower quartile of the observed distribution, particularly for *TSPY*, *DEFB*, and *NBPF*, further illustrating the limitation of a single linear reference in representing multicopy loci. To formally test for genetic ancestry associated differences in copy number, we applied the PCLAI local ancestry assignments described above to each gene locus in each haplotype. Quantifying these patterns across haplotypes, *TBC1D3* was by far the most ancestry-stratified family (*p = 9 × 10^-23^, ε^2^= 0.23*), followed by *CCL* (*ε^2^ = 0.17*); other families reached statistical significance but with small effect sizes, indicating that ancestry explains only a minor fraction of their copy-number variance (**Methods**).

Among the copy-number variable gene families, *TBC1D3* was of particular interest given its human lineage-specific expansion, and implicated role in cortical neurogenesis^91^. The Guitart et al. study characterized *TBC1D3* architecture using 69 human haplotypes, of which 46 were drawn from the HPRC1 assemblies; here, the 437 haploid assemblies passing QC in the *TBC1D3* locus in HPRC2 provide a near order-of-magnitude increase in haplotype sampling and enable finer resolution of the locus’s structural and population-level diversity. Consistent with previous reports, we observed that *TBC1D3* copies are dispersed across the two arms of Chromosome 17, but most copies map to one of two blocks at locus Chromosome 17q12, hereafter referred to as *TBC1D3* Cluster 1 and Cluster 2 (**Fig. 5d**).

Across the assemblies, we were able to resolve 89 unique structural configurations of Cluster 1 and 130 unique configurations of Cluster 2, reflecting the extreme architectural diversity of the locus; for each cluster, **Fig. 5d** shows the two references alongside the shortest and longest confirmed configurations and the three most common arrangements. Haplotype length spanned from compact configurations comparable to or smaller than GRCh38 and T2T-CHM13 (e.g., HG00140.1, HG00350.1, HG00133.2) to dramatically expanded arrangements harboring more than a dozen additional paralog copies (e.g., HG00673.2, NA19087.1). That range recapitulates the hypervariability described by Guitart et al., who reported haplotypes differing by as many as 20 *TBC1D3* copies and ∼1 Mb in length, and extends it across the broader and more ancestrally diverse HPRC2 cohort. We also note that transcription at this locus is restricted to a subset of paralogs^91^ (**Supplementary Fig. 30**) regardless of copy number, and whether copy number variation has functional consequences in humans is not established. The haplotype-resolved architecture provided here offers a population-scale resource for revisiting rearrangements at this locus.

#### A matched long-read pantranscriptome resource

For 206 individuals in HPRC2, we also generated long-read transcript data from the paired lymphoblastoid cell lines to further characterize the transcriptome across the pangenome. Each sample was sequenced to a median of 16.6 million total reads (median read length 1.7 kb), enabling the identification of ∼11,000 protein coding genes and ∼2,000 lncRNAs and other biotypes per assembly with an expression level of ≥1 count per million (CPM) (**Fig. 5e**).

To assess the prevalence of novel transcriptomic features across the pangenome, we quantified the frequency at which genes harbored novel isoforms using haplotype-specific long-read transcript alignments (**Fig. 5f**). Novel isoforms, defined as transcripts not present in the reference GENCODEv48 annotation, were widespread, with 759 genes across all biotypes consistently gaining at least one previously unannotated isoform across the majority of assemblies. This set includes 74,897 unique isoforms that appear in >1% of haplotypes.

To systematically characterize expressed sequence variation relative to the human reference genome, we developed the Reference Divergent Transcript (RDT) detector pipeline (**Methods**), a multi-step workflow that identifies full-length RNA transcripts whose sequences diverge from GRCh38, as assessed by reduced sequence identity and alignment score to the standard reference. Applying RDT-detector to the matched long-read transcript data and phased haplotype assemblies, we identified 68,681 unique reference-divergent transcript (RDT) sequences distributed across 578 genes. Across haplotypes, the median number of RDT-genes detected per haplotype was 53, reflecting widespread but variable divergence from GRCh38 across the cohort. The incidence of individual RDTs and novel isoforms ranged from those supported by a single haplotype to those present across all haplotypes, indicating that reference divergence at these loci spans from rare, potentially private variation to near-universal differences from GRCh38 (**Fig. 5f**).

Two RDT examples illustrate how matched haplotype-resolved transcript and genome data reveal functional variation invisible to reference-based approaches. In HG02965.2, a 25.7 kb segmental duplication at *RAD52* produces a transcript encoding a duplicated copy of the N-terminal single-stranded DNA binding domain (**Fig. 5g, Supplementary Fig. 31**), altering the protein domain architecture of a gene central to homologous recombination. Full-length transcript sequences from PacBio Kinnex long-read RNA-seq confirm expression of the novel isoform harboring a complete in-frame ORF. At *MOCS1*, a small tandem duplication (∼92 to ∼152 bp) within the 3’ UTR defines a distinct RDT isoform, and individuals heterozygous for this variant show markedly elevated *MOCS1* expression relative to homozygous reference individuals (**Fig. 5h, Supplementary Fig. 32**), implicating the duplication in post-transcriptional regulation. Neither variant is represented in GRCh38, and both would be missed or misassembled by reference-only transcript reconstruction. Together, these examples demonstrate that pairing a pantranscriptome with matched personal assemblies identifies both structural and regulatory variation that a single linear reference cannot capture.

We annotated TE sequences within each haplotype-resolved transcript model to determine how they contribute to reference-divergent transcriptomes (**Supplementary Fig. 33**; Jiang et al., companion manuscript). TE-associated transcripts were abundant, accounting for ∼44.6% of assembled transcripts, but nearly all involved fixed or reference-annotated TE sequences, reflecting the known contribution of ancient TEs to human transcript structure^95,96^. We therefore focused on the smaller but uniquely pangenome-resolved class of isoforms associated with polymorphic TE insertions absent from GRCh38. These included both MEI-associated transcripts from recent canonical mobilization events and transcripts associated with other polymorphic TE-containing alleles, likely reflecting older or structurally complex TE variation. Each haplotype harbored 2-48 polymorphic TE-related transcripts, including 0-26 associated with MEIs, representing non-reference isoforms that would be missed or misassembled by reference-only transcript reconstruction. At *MYBL2*, an intronic AluY insertion in HG00126.2 created a novel hypomethylated CpG island and initiated a previously unannotated transcript (**Fig. 5i**), illustrating how pangenome-resolved transcript discovery connects non-reference TE sequence to new regulatory elements and isoform diversity.

### A Panepigenome

Long-read sequencing recovers haplotype sequence and base modifications from the same molecules. Here we present a haplotype-resolved epigenomic catalog defined across the sequence diversity captured by HPRC2, rather than projected onto a single linear reference.

#### Non-reference CpG methylation across the pangenome

HPRC2 contains 17.6 million CpGs absent from T2T-CHM13, a 51.9% increase over the 33.8 million in the linear reference, compared with an 18.1% increase across the other 15 dinucleotides combined (**Fig. 6a, Supplementary Fig. 34a,b**). CpGs are gained through point mutation, GC-biased gene conversion at recombination hotspots^99^, and structural variation, including transposable element insertions and segmental duplications^100^. CpGs are lost, in part, by CpG-specific 5-methylcytosine deamination^101^, driven by their role as the principal substrate for DNA methylation. This regulatory-mutational coupling generates high inter-individual variability in CpG presence and accounts for their disproportionate accumulation in the pangenome. HPRC2 adds 6.47 million non-reference, common CpGs, with growth curves indicating saturation (**Fig. 6a**). In contrast, rare and ultra-rare CpGs (AF<0.01) continue to accumulate approximately linearly with the number of sampled genomes (**Fig. 6a**), reaching 10.9 million in HPRC2. Insertions contributed 5.69 million, SNPs 5.75 million, and inversions 629,206. A further 6.35 million could not be cleanly attributed to a single simple variant call (**Supplementary Table 10**). Of the 17.6 million non-reference CpGs with respect to T2T-CHM13, satellites contributed the single largest fraction (5.88M), followed by TEs (4.20M), CpG islands (CGIs) (3.34M), segmental duplications (1.78M), and promoters (1.41M); a further ∼4.7M were not attributable to any of these annotations (**Fig. 6b**; per-class enrichment over T2T-CHM13, **Supplementary Table 10**). Categories can overlap: 519,430 sites fell in both a CGI and a promoter, polymorphic regulatory CpGs with potential consequences for gene expression.

**Figure 6.**
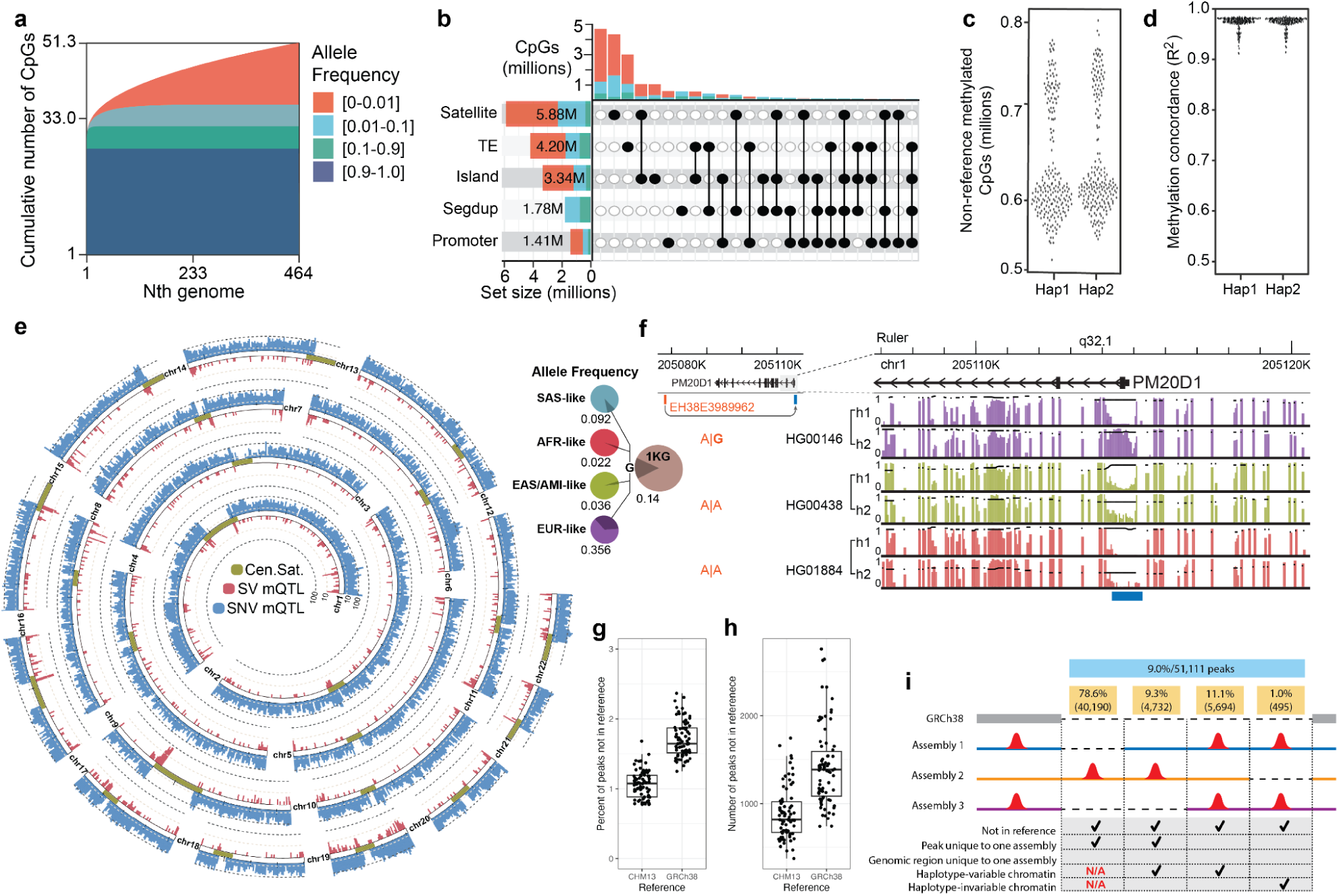
A haplotype-resolved panepigenome expands CpG and regulatory sequence space and links local ancestry-associated genetic variation to methylation and regulatory activity. **a**. Growth curve showing cumulative CpG discovery across HPRC2 haplotypes stratified by CpG frequency. CpGs were stratified by allele-frequency bins of 0 to 0.01, 0.01 to 0.1, 0.1 to 0.9, and 0.9 to 1.0. **b**. Genomic annotation of non-reference CpGs added by the panepigenome. Non-reference CpGs were intersected with assembly-specific annotations for satellites, transposable elements, CpG islands, segmental duplications, promoters, CpG shores, CpG shelves, and other annotated genomic contexts. **c**. Distribution of non-reference CpG count with high-confidence DNA methylation estimates captured by Panmethyl across HPRC2 individuals. Because each donor’s assembly is included in the graph, this is an idealized estimate. **d.** Per-haplotype concordance (R²) between Panmethyl on-graph methylation estimates and pb-CpG-tools assembly-based estimates at CpGs callable by both pipelines (Mean R² = 0.97). **e**. Genome-wide localization of promoter mQTLs with SNP or SV lead variants in T2T-CHM13 coordinates, shown relative to centromere and satellite annotations. Enrichment of mQTL intervals in centromeric and satellite-associated regions was evaluated against chromosome-aware shuffled interval sets. **f**. mQTL lead variant allele frequency stratified by PCLAI local ancestry and browser view of the PM20D1 region showing haplotype-resolved methylation, and genotype at the associated promoter mQTL lead variant. The lead variant rs9438393 falls within ENCODE enhancer EH38E3989962, approximately 36.6 kb from the PM20D1 promoter, and is associated with PM20D1 promoter methylation. Tracks show six representative haplotypes from three individuals, with genotype at the lead variant indicated at left. **g-h**. Percent and total number of FIRE peaks per haplotype assembly in genomic regions with no homology to common reference genomes (CHM13 and GRCh38). **i**. Locations of non-reference FIRE peaks relative to other haplotypes. Categories include peaks that are found only in one haplotype assembly on a genomic region also unique to that assembly, haplotype-unique peaks on non-unique genomic regions, non-unique peaks which are accessible in a subset of haplotype assemblies containing that genomic region, and non-unique peaks which are actuated in all assemblies which contain this genomic region.

We measured DNA methylation using PacBio and ONT long-read sequencing of the source LCLs (**Supplementary Note 2, Supplementary Fig. 35, Supplementary Fig. 36**). Estimates from the two platforms agreed closely over 1-kb intervals containing ≥10 CpGs at ≥10× coverage (mean deviation 1.55% ± 1.59%, **Supplementary Fig. 37**). Class-stratified methylation profiles of non-reference CpGs matched expected patterns from their reference counterparts (**Supplementary Fig. 34c**). A companion study of transposable elements (J. Juan., X. Zhuo, et al. companion manuscript) examines methylation at the polymorphic and broadly hypermethylated Alu, L1, SVA, and LTR5 families in depth (**Supplementary Fig. 34f-e**), revealing class-specific patterns. On-graph methylation calling^102–104^ recovers non-reference CpG methylation in samples without a matched assembly; in HPRC2 Panmethyl^102^ yielded 627,109 such CpGs per sample on average (**Fig. 6c**) at R² = 0.97 against assembly-based calls (**Fig. 6d**), making methylation characterization at common structural variants tractable at population scale.

#### Mapping methylation quantitative trait loci (mQTLs) across the pangenome

We next performed an mQTL analysis of promoter regions, identifying 80,854 promoter mQTLs (**Supplementary Table 11)**; 10.2% lay within structurally complex genomic regions difficult to interrogate with standard short-read, linear-reference functional genomics workflows (**Fig. 6e**). Although only 2,201 had SV lead variants, these were 1.71-fold enriched in CenSat regions (empirical P = 2.0 × 10^-4^), whereas single-nucleotide, indel, and small complex lead variants were depleted there, indicating that CpG methylation variation within centromeric and satellite-associated sequence is potentially more likely to be modulated by SVs and likely reflecting the higher background rate.

Promoter methylation is a key epigenetic regulator of gene expression, and promoter mQTLs identify inherited variants that shape this regulatory state; we asked how the lead variants underlying these associations are distributed across local genetic ancestries. At each lead-variant site, each haplotype was assigned a local ancestry by PCLAI and discretized into four local ancestry clusters. Of the 60,665 biallelic lead variants, most occur in all four clusters (75.7%), yet many differ significantly in allele frequency between them (**Supplementary Note 3, Supplementary Fig. 38**). The PM20D1 promoter mQTL illustrates this. The mQTL lead variant rs9438393 is ∼16-fold more frequent on EUR-like than AFR-like backgrounds (per-haplotype frequency 0.356 versus 0.022), and the distribution of per-haplotype PM20D1 promoter methylation states differs significantly across the four local-ancestry clusters (empirical P = 0.0099; **Fig. 6f; Supplementary Fig. 38h,i**). The variant lies within an ENCODE candidate enhancer linked to the PM20D1 promoter^105^, is a GTEx eQTL for PM20D1^106^, and is a GWAS association for monocyte count^107^, an immune-cell trait consistent with the lymphoblastoid context assayed here. Together these annotations describe a functional, trait-associated regulatory variant whose frequency, and the promoter methylation it marks, vary across local ancestries. These methylation patterns are browsable in the HPRC Panepigenome Browser ^108^.

#### A paired long-read chromatin epigenome map of the pangenome

For 38 of the assemblies we generated long-read chromatin epigenomic data from the paired LCLs to further characterize the accessible chromatin landscape across the pangenome (**Supplementary Table 6**). Each sample was subjected to single-molecule chromatin fiber sequencing (Fiber-seq), which uses a nonspecific N6-adenine methyltransferase (Hia5) to stencil protein occupancy footprints along DNA molecules via m6A-modified bases^44^. 20 of the HPRC Fiber-seq-treated samples were subsequently subjected to PacBio sequencing (median Gb = 108.1, N50 = 23,218 bp), 23 to ONT Simplex sequencing (median Gb = 106.9, N50 = 63,951 bp), and five were subjected to both (**Supplementary Fig. 39**). In addition, we performed both PacBio and ONT sequencing on HG002 LCLs for benchmarking purposes. Overall, we observed consistent chromatin stenciling patterns across both the PacBio and ONT samples, although the protein footprinting resolution was reduced with ONT sequencing because ONT Simplex captures adenine methylation along only one strand (**Supplementary Fig. 39, Supplementary Table 12**).

Fiber-seq data from each sample was then mapped to its paired diploid assembly, and Fiber-seq inferred regulatory elements (FIREs) were identified, resulting in 151,963 accessible chromatin elements identified per haplotype on average. Using a pangenome graph of all assemblies with Fiber-seq data and the T2T-CHM13 and GRCh38 reference genomes (Methods), we identified a consensus set of 568,430 accessible chromatin peaks across the 39 samples. Importantly, because these chromatin maps are grounded in the pangenome, they permitted the discovery of 51,111 accessible chromatin elements located within genomic loci absent from GRCh38 (9.0% of all consensus accessible chromatin peaks, **Fig. 6g-h**).

The discovery of these new accessible chromatin elements highlights that traditional short-read epigenetic maps grounded within GRCh38 are missing a significant portion of the regulatory landscape of a cell. Interestingly, most non-reference peaks exist in a single assembly and are enriched for being within segmental duplications (64%, 32,782 of 51,111, **Fig. 6i**). In companion studies, we demonstrate how these single-molecule chromatin maps can be used to resolve the unique chromatin architecture of telomeres^84^ and segmental duplications (Ren et al., in preparation), as well as how genetic variation modulates chromatin accessibility across the pangenome (Minkina, Mah-Som et al., in preparation).

### Pangenome applications

A key application of the pangenome is typing SVs that are otherwise difficult to detect with short-read sequencing. In a companion study, we applied PanGenie (v4.2.1)^109^ genotyping with the HPRC2 T2T-CHM13-based MC graph across 3,202 1000G short-read genomes, and show that we confidently genotype a median of 27,428 SVs per sample, thousands more SVs than alignment-based linear-reference short-read SV callers (**Fig. 7a, Methods, Supplementary Note 4**). Gains in total SVs genotyped per sample are substantial relative to the GRCh38-based HPRC1 pangenome (median number of SVs per sample increased by 9,339) and more modest relative to the recent T2T-CHM13-based HGSVC3 + HPRC1 pangenome^15^ (median increased by 2,742), indicating saturation of common SVs. Stratifying by AF, HPRC2 captures hundreds of additional rare SVs (AF < 1%) per sample compared to the HGSVC3 and HPRC1 PanGenie sets (**Fig. 7b**). This illustrates that continued expansion of pangenomes for SV imputation will increasingly capture rarer alleles.

**Figure 7:**
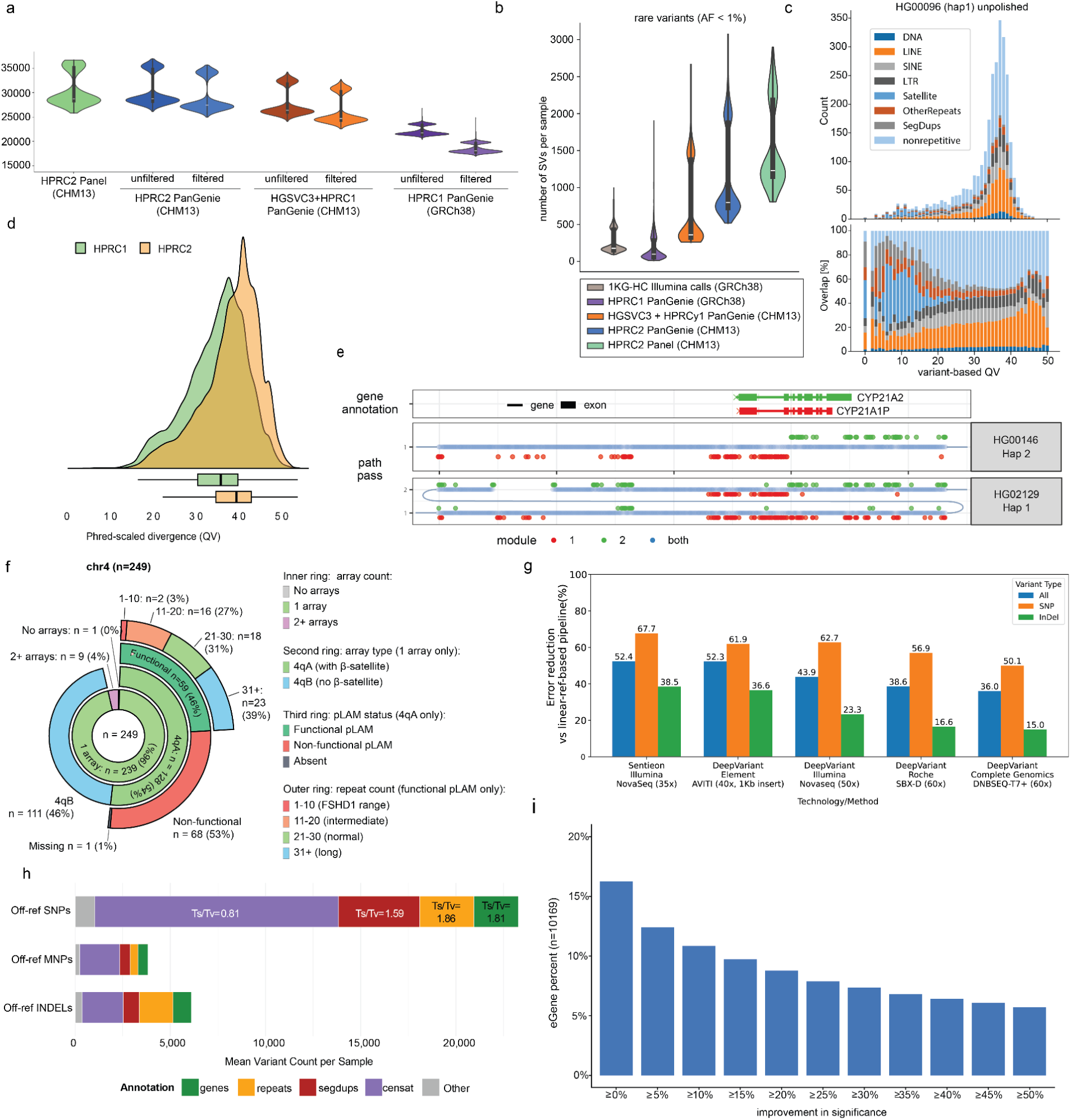
HPRC2 enables discovery in samples compared to it. **a**. Numbers of total SVs per sample across PanGenie-based genotypes for all 3,202 1000G samples computed based on three different Minigraph-Cactus graphs: the HPRC2 graph (CHM13-based, 462 haplotypes), the HGSVC3+HPRC1 graph (CHM13-based, 214 haplotypes) and the HPRC1 graph (GRCh38-based, 88 haplotypes). Numbers are shown for the respective unfiltered genotypes and filtered sets excluding sites not reliably genotypable by PanGenie. Additionally, the numbers of SVs per sample in the HPRC2 panel is shown, including variant calls directly derived from the 462 haplotype assemblies. **b**. Number of rare SVs (allele frequency < 1%) per sample observed across four 1000G callsets as well as the HPRC2 panel. The four 1000G callsets analyzed contained a traditional alignment-based callset produced from short-read data of all 3,202 individuals (grey) as well as the PanGenie genotypes computed for all 3,202 individuals based on the HPRC1 (GRCh38-based, purple), HGSVC3 + HPRC1 (CHM13-based, orange) and HPRC2 (CHM13-based, blue) graphs. Results for the HPRC2 panel are shown in green and include variants called from the Minigraph-Cactus graph from assemblies of 462 haplotypes. **c**. top: Variant-based QVs computed across 1 Mbp windows along the consensus haplotype generated from phased PanGenie genotypes for HG00096. Each bar is annotated by fractions of overlaps with BISER and RepeatMasker annotations across windows reported with a certain QV. bottom: Histogram with variant-based QVs computed across 1 Mbp windows along the consensus haplotype generated from phased PanGenie genotypes for HG00096, with bars normalized by their height. Each bar represents the fractions of overlaps with BISER and RepeatMasker annotations across windows reported with a certain QV. **d**. Haplotype availability across 273 challenging medically relevant genes ^110^, 35 HLA and 10 KIR genes for 462 HPRC2 haplotypes. Availability is calculated as phred-scaled sequence divergence (QV) between assembly haplotypes and closest independent haplotypes from the HPRC releases 1 (green) and 2 (orange). **e**. Two haplotypes are “painted” by their traversal through the RCCX pangenome. Dots are nodes ordered by their position in the pangenome (x-axis). Informative nodes (red/green) are specific to a module (pseudogene/gene). HG02129 has two modules, hence its path passes through the pangenome twice (y-axis). The second module of HG02129 and the first and only of HG00146 carry a fusion pseudogene-gene, marked by a switch from a red segment to a green segment, within the body of the CYP21A2 gene (upper panel). **f**. Sunburst chart showing the hierarchical classification of 249 chromosome 4 haplotypes across four concentric rings (innermost to outermost): (1) number of D4Z4 arrays detected (0, 1, or 2+); (2) for single-array haplotypes, allelic subtype: 4qA (terminal β-satellite present) or 4qB (absent); (3) for 4qA haplotypes, pLAM status: functional (ATTAAA), non-functional (ATCAAA), or absent; and (4) for functional-pLAM haplotypes, repeat-unit count binned by size: 1–10 (FSHD1 contraction range), 11–20 (intermediate), 21–30 (normal), and 31+ (long). Of 239 single-array haplotypes, 128 were 4qA, 59 of which carried a functional pLAM; 2 of these fell within the FSHD1 contraction range (≤10 repeat units). **g**. Percentage of variant-calling errors fixed by integrating the HPRC2 pangenome into the vg-DeepVariant and Sentieon pipelines relative to their linear-reference-based equivalents. Variant calling was performed on HG002 reads from four sequencing platforms (Illumina NovaSeq, Roche SBX-D, Element AVITI, and Complete Genomics DNBSEQ-T7+) mapped to GRCh38. In the baseline pipelines, neither the mapper nor the variant caller used the HPRC2 pangenome, except for SBX-D and DNBSEQ-T7+ data, for which only the mapper (BWA-MEM) was linear-reference-based. The calls were benchmarked against the GIAB assembly-based benchmark (GIAB-v5.0-GRCh38). Error reductions are shown separately for SNPs (orange), InDels (green), and all variants combined (blue). **h**. Mean number of SVs genotyped per sample (n=9) with vg call relative to T2T-CHM13 on the v2.1 HPRC Minigraph-Cactus graph, stratified by HPRC annotation. Small off-reference GRef variants nested in these insertions are also shown. (**i**) Proportion of 10,169 eGenes with improved association strength using the merged 1000G and pangenome variant callset (28.4 million variants) relative to the 1000G callset alone (19.2 million variants). For 10,169 eGenes detected using the merged callset, improvement was defined as the percent increase in -log10(beta-approximated p-value) for the lead variant from the merged callset compared with the lead variant from the 1000G alone callset. Larger -log10(p) indicates stronger association. Bars show the proportion of eGenes exceeding each percent-improvement threshold.

The use of PanGenie for imputation of all forms of variation paves the way toward more complete genome inference at scale. To demonstrate this we phased the PanGenie genotypes and reconstructed draft haplotype resolved assemblies of all 1000G samples, resulting in median k-mer based QVs of 43 (∼1 error per twenty thousand bases). Expectedly, however, genomic sequences overlapping satellites and some segmental duplications remain challenging to reconstruct based on short read genotypes (**Fig. 7c**) with our current alignments.

Beyond genome-wide genotyping, the pangenome enables targeted analysis of loci of medical interest. Indeed, HPRC2 de novo assemblies better cover human diversity across highly polymorphic and complex genes: for 318 challenging medically relevant loci^110^, median sequence divergence between local haplotypes and the closest independent haplotypes increased from QV 35.8 (HPRC1) to 39.5 (HPRC2). Similarly, the fraction of underrepresented haplotypes (no other haplotype with QV ≥ 30) decreased from 23.5% to 13.1% (**Fig. 7d**). Using ctyper^111^ to type these regions in held out short-read samples gave concordant results **(Supplementary Note 5; Supplementary Fig. 40**).

The more complete representation allows for more complete genome inference and a clearer delineation between common and pathogenic variation. For example, the segmentally duplicated RCCX module, which contains the CYP21A2 gene and CYP21A1P pseudogene, is frequently subject to deletions and interlocus gene conversion that can cause the recessive condition congenital adrenal hyperplasia (CAH). The standard genetic test to identify pathogenic alleles is convoluted and incomplete, but recently a specialized pangenomic method has been developed, implemented in the tool Parakit, that enables full characterization of the alleles from long-read sequencing^112,113^. Updating this method to use HPRC2, the local pangenome for the challenging RCCX region is more complete, with 51% more informative markers compared to HPRC1 (**Supplementary Fig. 41**, **Supplementary Note 6**), increasing Parakit’s ability to detect fusions and gene conversions. As in the HGSVC3 study^15^, a majority of haplotypes carried two modules (80.3%) and a minority carried one or three modules (10.0% and 9.0%), although we also found three haplotypes (0.6%) carrying four modules. Parakit further identified 7 candidate risk-carrying haplotypes (out of 462) in HPRC2 that carry a fusion overlapping the CYP21A2 gene or a gene conversion resulting in LOF and a known pathogenic variant (**Fig. 7e**).

Complex loci with extensive copy number variation are difficult both to resolve and to display, because near-identical repeat units align ambiguously to a linear reference. The D4Z4 macrosatellite on chromosome 4q35 is a clinically important example: contraction to ≤10 units on “permissive” haplotypes causes facioscapulohumeral muscular dystrophy type 1 (FSHD1), one of the most commonly inherited myopathies^114,115^. Using KaryoScope, an alignment-free, k-mer-based annotation tool developed in a companion study^116^, we resolved and visualized D4Z4 arrays across the q arms of the 249 T2T chromosome four haplotypes in HPRC2 (**Fig. 7f; Supplementary Fig. 42a-b; Supplementary Note 7**).

Of the 239 haplotypes carrying a single D4Z4 array, 128 (53.6%) were 4qA – the permissive allele, marked by a terminal β-satellite and carrying the polyadenylation signal required for pathogenic DUX4 expression – and 111 (46.4%) were 4qB, which carries neither^83,115^. BLAST-based analysis confirmed a functional polyadenylation signal (ATTAAA) in 59 of 128 4qA haplotypes (46.1%), of which two fell within the FSHD1 pathogenic range (≤10 units), carried by two of 154 individuals – a carrier frequency (∼1.3%) consistent with the incomplete penetrance observed at the upper end of the pathogenic range^117^ (**Supplementary Fig. 42c**).

#### Pangenome based read mapping improves small variant genotyping

In addition to structural characterization, pangenomes are increasingly used by the best performing read mapping and small variant calling pipelines^24,26,27^. To demonstrate this across a range of different sequencing technologies, we used Sentieon^118^ and DeepVariant^27^ based best practice pipelines, each using the HPRC2 MC GRCh38 based pangenome graph or, alternatively, the GRCh38 linear reference (**Supplementary Notes 8,9**). In all cases the output variant calls were expressed against GRCh38, the difference being in which inputs were provided for mapping and variant calling. Assessing against the most complete T2T-assembly-based benchmark available (GIAB-v5.0), for every tested sequencing technology, overall point variant calling accuracy was improved by the use of the pangenome (**Fig. 7g, Supplementary Tables 13 and 14**), with reductions in errors genome-wide ranging from 52.4% to 36% overall, with larger gains for SNPs of between 67.7% and 50.1%.

Our mapping-based genotyping workflows were limited to making calls with respect to a linear reference. However, pangenome graphs facilitate exploration of variants that occur within regions not represented in conventional references. To demonstrate this, we created a new pipeline to generate and utilize GRef coordinates, allowing variants to be expressed against the sequences of T2T-CHM13 and additionally an alignment disjoint set of non-reference haplotypes that cover the pangenome. As a proof of concept, we called variants for nine GIAB samples using Illumina short-read sequencing data. We explored two different variant calling approaches to determine consistency, one calling against the graph itself using vg call^119^, and one produced by projecting read mappings to the GRef sequences and calling with FreeBayes^120^. An average of 32,221 small variants and 726 SVs were called off reference per sample using vg call (**Fig. 7h**), with 81% also being called by FreeBayes (**Supplementary Fig. 43**). While this is a minority of the total off reference variation observed per haplotype in HPRC2 assemblies (**Fig. 3g**), this both demonstrates that substantial variation can be called off reference and that further technology development, likely involving long reads^121^, will be needed to fully ascertain such variants in new samples.

#### Pangenome-based variant calling improves trait association

The comprehensive map of genome variation provided by HPRC2 has the potential to improve trait association studies through the inclusion of difficult-to-detect variants that are not well ascertained by short-read genome sequencing studies that rely on alignment of reads to the GRCh38 reference. As part of a companion paper, we used the genetics of gene expression variation as a model for the gains that might be expected from the application of pangenomic methods to future genome-wide association studies (GWAS). We performed expression quantitative trait locus (eQTL) mapping in a set of 430 samples from 1000G that have both short-read WGS^38^ and RNA-seq data^122^, and measured the added value of supplementing 1000G variant calls with additional pangenome-specific variants identified by three methods that use HPRC2: PanGenie^29^, danbing-tk^123^, and a new alignment-based method (EdgeDepth^124^) for associating pangenome graph features with traits.

Notably, pangenome-specific variants account for 20.3% of lead markers at the 10,169 eQTLs found using the merged “HPRC2+1000G” callset, demonstrating that HPRC2 enables the discovery of many putative functional variants that would otherwise be missed. Consistent with this, the HPRC2+1000G callset identifies a much larger fraction of eQTLs caused by multiallelic SVs (2.5%) and intermediate sized indels (10-49 bp) (2.9%) than the 1000G callset alone (0.6% and 1.3%, respectively), and a larger fraction of variants in difficult genomic regions (28.1% vs 21.7%), greatly expanding the spectrum of regulatory variation relative to prior short-read eQTL studies^125–129^.

Perhaps more importantly, the addition of pangenome-specific variants improves trait association significance at a large number of genes, even when controlling for the number of tests (**Fig. 7i**): for example, 893 eGenes (8.8% of total) had a ≥20% improvement in -log10 p-value in the merged variant callset relative to 1000G alone, 581 (5.7%) had a >50% improvement. These gains are primarily due to the EdgeDepth method (78.1% of loci), showing that aligning reads to the HPRC2 pangenome graph is a powerful approach. Given that eQTLs – like GWAS hits – are typically caused by regulatory variants that alter gene expression, these results suggest that application of pangenomic methods to GWAS will improve trait association power and causal variant prediction, and thus increase the potential for novel discoveries.

## Discussion

HPRC2 extends the human pangenome reference to 460 haplotypes and, by optimized sample selection, covers over 99% of common variants observed in the All of Us Research Program v8 cohort^130^. It represents substantially improved assemblies relative to earlier efforts, three complementary pangenome alignment representations, the first alignment based pangenome coordinate systems, and provides matched pantranscriptome and panepigenome layers. The assemblies underpinning HPRC2 integrate high-coverage HiFi^131^ with ultra-long ONT^132^, combined trio or Hi-C phasing, and a long-read polishing pipeline^47^. Relative to HPRC1, the fraction of regions estimated to be structurally unreliable and the number of base errors halves. HPRC2 assemblies also have excellent contiguity with a majority (59%) of chromosomes being represented with T2T contigs or scaffolds, and 93% of chromosomes assembled with 3 or fewer gaps or breaks (**Supplementary Fig. 44**).

However, several issues remain with reaching the goal of truly complete, T2T assemblies; 34% of the active alpha-satellite centromeric arrays either were not contiguously assembled or contain flagged errors, and relatively few acrocentric short arms (157 total, 6.8% of total) were assembled contiguously with no estimated structural QC; similarly a small fraction of segmentally duplicated regions cannot yet be validated against short-read depth, indicating residual assembly artifacts (12.5 Mbp per haplotype).

At the base level, accuracy estimates indicate that the assemblies are correct at the one error in a hundred thousand scale, principally being held back by the limitations of current long-read technologies to resolve longer homopolymer lengths^47^. Continued improvements in long-read accuracy, polishing and assembly will therefore be needed before assembly quality ceases to be a limiting factor, although recent innovations in error correction for ONT ultra-long reads appear promising in reducing errors and improving contiguity, particularly in regions only spanned by those reads^133,134^.

Coverage of common variation in HPRC2 is approaching saturation for the populations sampled. The MaxVar trajectory shows steep early gains followed by diminishing returns as the variants added become increasingly rare or population-specific. Given this, we estimate that outside of hard to assemble regions (defined as centromeres, satellites on the q-arm of chrY, and acrocentric short-arm sequences; 90% of the T2T-CHM13 genome lies outside these regions), we are assembling haplotypes that include 99% of all common variants for major populations, like those represented by All of Us, with 99.6% completeness and correctness (as judged by an absence of flagged errors). This is a substantial improvement over HPRC1 and comparable prior efforts^13,15,18^. Looking ahead, with further assembly improvements and for the populations sampled, we can expect to reach completeness of all common variation across the entire genome, with the potential exception of the rDNA arrays, which remain refractory to current assembly methods^6^.

While an important achievement, the anticipated completion of this map of common variation will still leave significant gaps. As expected, rarer variation is and will be less well covered, for example, at an AF ≥ 0.1% we estimate recovery of slightly over 80% of such AoU variants in HPRC2. Given that rare variants in the 0.1% to 1% range often have stronger effect sizes than common variants ^135^, and that contemporary biobanks have the scale to test associations to this variation ^130,136^, it is desirable to expand the number of genomes sampled to address this potentially important missing variation. The pangenome will allow the genetic context of this rarer variation to be better understood, and should assist in its accurate genotyping in samples compared to it. Our analysis indicates it will require on the order of thousands of samples to reach 99% coverage for variants down to 0.1%. Hypervariable regions, such as large repeat arrays, contain an excess of rarer variants, meaning that deeper sampling will be necessary to saturate haplotype representation there, as our analysis of alpha-satellite array completeness indicates. Lastly, these estimates of coverage do not include human populations that are not well represented by 1000G and All of Us, in particular Oceania and East, Central, and Southern Africa^137,138^; engaging with and extending pangenome efforts to such populations should be an important area of future collaborative work.

Projects that seek to recruit individuals because their genomic information may include variation not present in existing resources bears risk. Accordingly, ELSI considerations of future expansion of pangenomic efforts to include additional population groups will be a key concern moving forward.

Recruitment done well, like in the 1000G, can include and empower communities and build trust in research. Done less well, such projects can lead to rejection of genetics, genomics, and the research enterprise. For instance, the Human Genome Diversity Project (HGDP)^139^ aimed to discover human genetic variation by sampling demographically diverse human groups around the world, including Indigenous Peoples^140^. The project was rejected by many Indigenous communities, who viewed it as extractive, disrespectful, and antithetical to their values and interests^141,142^. That backlash diminished the HGDP’s opportunities to achieve its aims.

Furthermore, expanding to include new groups will require the resource to accommodate additional population labeling practices. HPRC2 builds on work across multiple projects with different labeling practices and was evaluated against All of Us, which uses yet another population labeling scheme. Our awareness of the labeling issues and future collaboration will help keep future drafts on firm ethical footing with respect to their use.

Learning from these successes and failures, we aim to create a resource that represents global human genome variation. As said elsewhere, “[t]he lack of representation of diverse ancestral backgrounds in genomic research is well-known” and creates both technical and ethical problems for translating genome research into effective medical tests and treatments^143^.

To relate the HPRC2 assemblies to one another we report three complementary alignment representations, Minigraph^58^, MC^59^ and IMPG/PGGB^60^, each suited to different downstream tasks. Minigraph offers a compact, SV-resolution representation for visualisation and read placement; MC supplies base-resolution alignments in the alignable, predominantly euchromatic portion, of the genome and is the basis for the provided pangenome coordinates; and IMPG/PGGB efficiently represents a large set of pairwise alignments, including in centromeric and acrocentric regions where a single multiple-sequence alignment is currently so unreliable as to be omitted from both the Minigraph and MC alignments.

Two pangenome coordinate systems, Pantree^65^ and Graph Reference (GRef), are introduced to make off-reference variation addressable within standard file formats. Pantree indexes a minimum spanning tree over the graph; GRef extends a chosen linear reference with a vertex-disjoint set of contigs covering non-reference insertions. As such they provide a non-redundant substrate onto which variants can be expressed, building on similar prior models^58,67^. They complement the use of the coordinates of the underlying haplotypes in the pangenome, which are standardized by the pansn format^144^. Together they permit nested variation within insertions that is otherwise lost when projecting back to a single linear reference to be expressed in variant formats (e.g. VCF). The per-sample numbers demonstrate the importance of these non-reference coordinates: a median of 143,275 variants (2.6% of all variants per sample) are non-reference under GRef, indicating that off-reference content is a significant, hitherto under appreciated feature of human genomes that has been eclipsed by reference bias.

Similarly, in a companion effort using PacBio HiFi Fiber-seq and Illumina datasets, 2.8% and 2.1% of somatic variants, respectively, were detected on HPRC2 haplotype paths absent from GRCh38 using a new pangenome driven tool^145^. While the tooling to support off-reference coordinate systems is early, we anticipate near term work to build and mature support for these systems such that future variant calling pipelines are able to address this additional variation while requiring minimal extensions of existing formats or downstream tools.

Several analyses indicate that pangenome-aware pipelines now exceed linear-reference pipelines for standard variant-calling tasks in terms of accuracy while not being substantially computationally more intensive^26,27^. Using DeepVariant and Sentieon based pipelines, point-variant errors fall consistently across a broad range of sequencing technologies relative to GRCh38 alone, with a 36-52.4% reduction in small variant errors relative to closely comparable linear reference based pipelines. Complementing this, the new version of PanGenie using HPRC2 is able to genotype a median of ∼27,428 SVs per short-read dataset, a ∼1.5-fold increase over previous versions using the HPRC1 resource and comparable in number to long-read based discovery approaches^15,146^. In the future, the combination of these approaches should improve the accuracy and completeness of genome inference; demonstrating this potential, we show in a companion article^109^ that additional polishing of our haplotype reconstructions of 1000G samples can reach a median QV46 (∼1 difference per 40,000 bases).

A key promise of improved variant calling is that it will enable the discovery of new variants and genes that underlie human traits. Future biobank-scale studies will be required to address this, however, there is promising evidence from eQTL analyses presented here, where application of pangenome-based methods identified numerous trait associated SVs and indels that were missed by standard GRCh38-based methods, and substantially strengthened association significance for a nontrivial fraction of eGenes (conservatively, >5%)^124^.

HPRC2 is the first pangenome to include matched long-read transcriptomic, methylation and chromatin accessibility data at this scale. Using the long-read RNA-seq data, two independent annotation pipelines applied to each HPRC2 haplotype recover highly consistent, near-complete gene annotations; with respect to existing GENCODE annotations the median haplotype contains >99% of known protein-coding genes and >98% of non-coding genes. This provides fuller insight into the pervasive impact of structural variation on the transcriptome, with the median annotation cataloging 126 gene duplication or deletion events, such that 14% of GENCODE protein coding genes and 19% of non-coding genes are either duplicated or deleted in at least one haplotype in HPRC2. Underlying the gene duplications, we find that 12% of the human genome is segmentally duplicated in HPRC2, expanding substantially our catalog of these events from earlier studies ^13,15,18^.

The matched long-read transcript data also identifies tens of thousands of GRCh38 reference-divergent transcript (RDT) sequences distributed across hundreds of genes. This catalog represents transcripts that are poorly represented by the existing reference. Given that these RDTs were identified only from sequencing of LCLs, future work is needed to create comprehensive RDT libraries across cell types and genetic backgrounds and ultimately create a comprehensive pantranscriptome that addresses reference bias in tasks such as gene and transcript expression quantification.

The long-read Fiber-seq datasets collected across 38 lymphoblastoid cell lines with paired HPRC assemblies identified 51,111 accessible chromatin candidate cis-regulatory elements (cCREs) that are present within genomic loci absent from GRCh38, representing 9% of all of the consensus cCREs identified across these samples. Furthermore, as part of companion manuscripts we show that these previously unanotated cCREs include clinically relevant gene enhancers (Minkina, Mah-Som, et al., in preparation), demonstrating that our existing catalogs of GRCh38-grounded cCREs are incomplete, even within lymphoblastoid cells, which are well represented in large cCRE catalogs such as ENCODE^147^. Further work extending these epigenetic measurements across cell types is required to fully resolve how genetic variation across the pangenome is impacting human health and disease at a population level.

These findings extend beyond chromatin accessibility. The pangenome reveals 17.6 million CpGs not represented in any single linear reference, accounting for 34% of all CpGs observed across the cohort. These CpGs are concentrated in satellites, transposable elements, segmental duplications, and CpG islands and can now be incorporated into population-scale analyses through on-graph methylation calling^102,103^. Promoter mQTL mapping against the HPRC2 graph identified 80,854 associations, with structural-variant lead variants enriched 1.71-fold in centromeric and satellite sequence. Together, these findings indicate that the greatest gains from pangenome-aware methylation analysis occur in repetitive and structurally variable regions that remain poorly accessible to short-read, linear-reference workflows.

HPRC2 should prove useful for many analyses, and we encourage community development upon it. It is best understood not as a finished product but as the second phase of a planned three-phase project to construct a stable, shared global pangenome reference^148^. In HPRC2 global collaboration expanded the representation of 1000G samples of Japanese individuals in Tokyo (JPT)^39^ and Toscani in Italia (TSI). In phase three, coordination with HGSVC3, international pangenome efforts and further expanded recruitment of newly consented samples by HPRC will allow the construction of a global human pangenome at a substantially larger scale.

Improvements in assembly, reaching at least T2T scaffolds for almost all chromosome haplotypes and further reducing errors, should result in near uniform representation of all classes of variation genome-wide. New alignment methodologies should provide useful representation of satellites, expanding the proportion of the genome usefully represented in the variation graph. The continued maturation of tools will allow the resource to be integrated into many genomic and transcriptomic workflows - many as black box applications that simply improve inference for critical tasks. The result will be a more complete, less reference biased view of all variation across human populations and new discoveries across the entirety of the genome.

## Methods

### Data Availability

Data is released openly in the public domain in accordance with HPRC data use best practices^149^ which aim to promote the ethical, legal, and fair use of all HPRC data and apply to all HPRC data users. They articulate best practices concerning: i) future scientific publications, ii) intellectual property, iii) re-identification and harm, iv) fairness and beneficence, v) population descriptor use, and vi) disclaimers. Sequencing data and assemblies are uploaded to International Nucleotide Sequence Database Collaboration (INSDC) repositories. Sequencing data are available in Sequence Read Archive BioProjects PRJNA701308 and PRJNA731524. Assemblies are available in Genbank Bioproject PRJNA730822. Sequencing data, assemblies, assembly annotations, and pangenome alignments are also available through AnVIL (https://anvilproject.org) and the AWS Open Data Program (https://registry.opendata.aws). Metadata and download URIs available through the HPRC Data Explorer on https://humanpangenome.org/. Assemblies and select annotations are available through a UCSC Genome Browser Hub (https://hgdownload.gi.ucsc.edu/hubs/HPRC/index.html) as well as the Ensembl Genome Browser^150^ (https://projects.ensembl.org/hprc/). Assembly and epigenomic data is available in the HPRC Epigenome Browser (https://epigenome.humanpangenome.org/).

### Sample Selection and local ancestry labelling

#### MaxVar Algorithm

For samples with existing sequence data, in each iteration, MaxVar enumerates all common variants present in the selection pool, excludes those already covered by the existing reference data set of individuals, and ranks candidate individuals by the number of additional variants they would contribute upon inclusion. The top-ranked individual is added to the reference, and the algorithm proceeds to the next iteration with updated coverage estimates. This greedy selection process prioritizes individuals that maximize marginal gains in coverage, yielding a rapid increase in the fraction of common variants represented in the panel.

#### Point Cloud Local Ancestry Inference (PCLAI)

Point cloud local ancestry inference (PCLAI)^34^ training and inference was performed in GRCh38 coordinates using 1000G^33^ samples. PCLAI represents each haplotype as a sequence of fixed genomic windows and predicts, for each window, a continuous coordinate in a reference two-dimensional PCA space and a prediction confidence score. The reference PCA was spanned by 1000G samples with low recent admixture, defined as individuals with at least 95% assignment probability to a single cluster after unsupervised ADMIXTURE^35,36^ clustering at K=7. In this analysis, each PCLAI segment corresponded to a non-overlapping window of 1,000 SNPs. Prior to training, the VCF was downsampled to retain variants with MAF≥0.001 and LD-thinned with PLINK 2^151^ using a 50 kb window, a step size of 25 SNPs, and r^2=0.99.

For HPRC2 pangenome samples, local ancestry painting was performed using variants derived from MC alignments. The MC wave VCF was first converted to a biallelic SNP representation and intersected with the set of variants used for PCLAI training. To avoid rephasing of the wave VCF, missing sites were imputed with Minimac4^152^. PCLAI was then run on autosomes in GRCh38 coordinates and the outputs were represented as BED tracks. Each BED row corresponds to a haplotype-specific genomic window and encodes the chromosome interval, sample, haplotype, chromosome-window identifier, the inferred coordinate, confidence score, and an associated RGB color from the interpolated CIELAB space. The confidence score ranges from 0 (lowest confidence) to 1000 (highest confidence) and it was computed as: each window’s uncertainty (dispersion of results after 100 stochastic forward passes with dropout layers kept active, computed as the trace of the empirical covariance matrix) was divided by the chromosome-specific 99th percentile of uncertainty values and truncated at 1. The result was inverted and multiplied by 1,000 to produce a confidence score. Windows with confidence scores below 141 were filtered from the released BED tracks. The cutoff was calibrated empirically and is approximately equivalent to removing windows whose uncertainty 85.9% of the chromosome-specific 99th percentile uncertainty value. To project PCLAI inferences from GRCh38 onto pangenome assemblies, the BED tracks were converted to T2T-CHM13 and native pangenomic assembly coordinate systems using the LiftOver tool from the UCSC Genome Browser. Because coordinate conversion was harder at some particular difficult loci, such as acrocentric short arms and centromeric regions, there are some missing intervals in the released pangenomic PCLAI tracks, besides other sources of missingness such as absence of source calls in the MC VCF or low-confidence filtering. The finalized PCLAI BED tracks for HPRC2 are available at https://github.com/human-pangenomics/hprc_intermediate_assembly/tree/main/data_tables/annotation/pclai in GRCh38, T2T-CHM13, and assembly coordinate systems.

For analyses requiring categorical genetic ancestry labels, continuous PCLAI coordinates were discretized after inference. As described in *Benchmark: discretized PCLAI vs. ADMIXTURE,* 15-nearest-neighbors (KNN) classifier using Euclidean distance was trained in the same reference PCA space with 1000G labels. Because of class imbalance, samples from East Asian-like and Indigenous American-like 1000G populations were grouped into a single class, AMR/EAS-like (**Supplementary Fig. 3a**). Each inferred haplotype-window coordinate was then assigned to the corresponding KNN-defined region, resulting in discrete labels used for

downstream analyses. For further guidance on imputation and clustering of PCLAI results, refer to the PCLAI Manual at https://github.com/AI-sandbox/hprc-pclai.

To enable a direct comparison with discrete genetic ancestry inference methods, we discretized PCLAI’s^34^ continuous per-window coordinates by fitting a L_2_ distance 15-nearest-neighbors (KNN) classifier in the reference PCA space and using its decision boundaries to partition the space into four data-driven regions. Each genomic window was assigned to one of these four KNN regions based on its predicted coordinates. We then computed, for each sample, the genome-wide percentage of windows assigned to each KNN region, resulting in a per-sample four-component profile. Across the benchmark test set of 100 samples, we compare these discrete PCLAI profiles to optimized ADMIXTURE^35,36^ (K=4). Because ADMIXTURE cluster indices and KNN region indices are not inherently aligned, we first determined a one-to-one mapping between the four ADMIXTURE clusters and the four KNN regions using dominant (argmax) assignments in the test set (i.e., matching each ADMIXTURE cluster to the KNN region that most frequently co-occurred as the dominant label across samples). Under this mapping, the resulting dominant-label assignments were highly concordant between the two methods. The confusion matrix of ADMIXTURE (K=4) argmax cluster labels versus discretized PCLAI (KNN-region) argmax labels showed perfect agreement (**Supplementary Fig. 3**). These results support the use of discretization as an optional post-processing step when categorical labels are required.

### Sequencing Data

#### Sample Source

Lymphoblastoid cell lines (LCLs) used for sequencing from the 1000 Genomes Project (**Supplementary Table 5**) were obtained from the NHGRI Sample Repository for Human Genetic Research at the Coriell Institute for Medical Research. HG06807 LCLs from the Human Pangenome Reference Collection were also obtained from the NHGRI Sample Repository at Coriell. HG002 (GM24385) and HG005 (GM24631) LCLs were obtained from the NIGMS Human Genetic Cell Repository at Coriell.

#### Cell Line Production

All new HPRC R2 LCLs, as well as HPP LCLs sequenced by Human Technopole, were derived from the original expansion culture lot to ensure the lowest possible number of passages and to reduce overall culturing time. Cells used for PacBio HiFi, Nanopore Ultra-long, Omni-C (i.e. Hi-C), Kinnex RNAseq, and Fiber-seq production as well as g-banded karyotyping and Illumina Omni2.5 microarray were expanded to a total culture size of 4×10^8^ cells, resulting in a total of five passages post-cell line establishment or receipt at Coriell. Cells were split into production-specific sized vials: HiFi (2×10^7^ cells), Nanopore (5×10^7^ cells), Omni-C (5×10^6^ cells), Kinnex RNAseq (5×10^6^ cells), and Fiber-seq (1×10^7^ cells). Cells for Fiber-seq were stored in 65% RPMI-1640, 30% FBS, and 5% DMSO and frozen as viable cultures until ready for processing. All other cells were washed in PBS and flash frozen as dry cell pellets. Fiber-seq cells were split into two reactions (5×10^6^ cells) and prepared for DNA extraction at Coriell following established methods ^44^, including nuclei isolation and chromatin stenciling with Hia5 and SAM, and were flash frozen prior to sequencing.

#### Cell Line Karyotyping and Microarray QC

G-banded karyotype analysis was performed on 5×10^6^ live cells harvested at a minimum of passage 5 (post-establishment). For all LCLs, 20 metaphase cells were counted, and a minimum of five metaphase cells were analyzed and karyotyped. Chromosome analysis was performed at a resolution of 400 bands or greater. A pass/fail criteria was applied before LCLs proceeded to sequencing. LCLs with normal karyotypes (46,XX or 46,XY) or lines with benign polymorphisms that are frequently seen in apparently healthy individuals were classified as passes. LCLs were classified as a fail if two or more cells harbored the same chromosomal abnormality. Cell line specific karyotypes can be found in **Supplementary Table 15** for all lines that passed our filtering criteria. DNA used for microarray was isolated from frozen cell pellets (3×10^6^-7×10^6^ cells) using the Maxwell RSC Cultured Cells DNA Kit on a Maxwell RSC 48 instrument (Promega). DNA was genotyped at the Children’s Hospital of Philadelphia’s Center for Applied Genomics using the Infinium Omni2.5-8 v1.3 BeadChip (Illumina) on an iScan System instrument (Illumina).

#### PacBio HiFi Sequencing

PacBio sequencing was performed across three centers: Washington University in St. Louis (WUSTL), Rockefeller University, and the University of Washington (UW). Throughout this manuscript, these centers are referred to by their abbreviations WUSTL, Rockefeller, and UW, respectively.

##### DNA Extraction

High-molecular-weight DNA was isolated from frozen cell pellets using a kit designed to produce highly intact DNA: at WUSTL & Rockefeller, the Qiagen MagAttract HMW DNA kit (Qiagen p/n 67563); at UW, the MagAttract HMW or NEB Monarch HMW DNA Extraction Kit for Cells & Blood (New England Biolabs p/n T3050L). Extracted DNA was checked for quantity with a fluorometric method (Qubit dsDNA ThermoFisher p/n Q32854) and length was assayed using a FEMTO Pulse instrument (Agilent p/ns M5330AA and FP-1002-0275). Samples with sufficient mass (≥ 5 µg) and length (mode ≥ 20 kbp) proceeded to sequencing library preparation.

##### Sequel II library preparation and sequencing

Extracted high-molecular-weight DNA was diluted to < 50 ng/µl and sheared using the Megaruptor 3 instrument twice to achieve mode fragment lengths ∼ 22 kbp; depending on initial QC and Hydropore lot, typical shear settings were 30/31, 29/31, or 28/30. Samples processed in project year 2 and the first half of year 3 were prepped using the SMRTbell Express Template prep kit 2.0 (PacBio p/n 100-938-900); samples from the second half of year 3 used the SMRTbell prep kit 3.0 (PacBio p/n 102-141-700). Sample barcoding was achieved using Barcoded Overhang Adapter kits 8A/8B (PacBio p/ns 101-628-400/101-628-500) for the first half of year 3, and transitioned to SMRTbell adapter index plate 96A (PacBio p/n 102-009-200) in the second half of year 3. Manufacturer’s protocols for the respective kits were generally followed with the following modification: after adapter or barcoded adapter ligation and exonuclease treatments, samples were size-selected using a PippinHT instrument on a 0.75% agarose cartridge using a 15000 bp high-pass setting (Sage Science p/ns HTP0001, HPE7510).

After final library QC using Qubit and FEMTO Pulse, libraries were prepared for sequencing on Sequel II (WUSTL/UW) or IIe (Rockefeller) instruments using the latest chemistries for long-insert HiFi libraries: for year 2 samples, v2 or v2.2; for year 3, v2.2 or 3.2. Movies were acquired with Adaptive Loading and the longest possible movie times, i.e. 30 hrs for year 2 and the first half of year 3, 40 hrs for the second half of year 3. Where possible, full subreads files were exported to enable DeepConsensus polishing after ccs consensus analysis.

##### Revio library preparation and sequencing

Extracted high-molecular-weight DNA was diluted to < 50 ng/µl and sheared using the Megaruptor 3 instrument twice to achieve mode fragment lengths ∼ 22 kbp; depending on initial QC and Hydropore lot, typical shear settings were 28/31 or 29/31. Samples were processed using SMRTbell prep kit 3.0, SMRTbell adapter index plate 96A and PippinHT size selection as described above.

After final library QC, libraries were prepared for sequencing on Revio instruments using software version 12.0 (Year 4), 13.0 (Years 4 & 5), or 13.1 (Year 5), the release chemistry (p/n 102-739-100 & 102-202-200), and the longest possible movie times (24 or 30 hrs.) The Revio instrument runs DeepConsensus onboard on all data sets.

#### DeepConsensus Basecalling on Sequel II Data

PacBio HiFi data generated on Sequel II and IIe platforms were rebasecalled with DeepConsensus^153^ when subreads were available (604 out of 680 files). Reads.bam files were prepared for DeepConsensus with a WDL (https://github.com/human-pangenomics/hpp_production_workflows/data_processing/wdl/tasks/readsbam4dc.wdl). Briefly, the reads.bam files were subset with Samtools to retain reads with predicted read quality over 0.88. Multiplexed or barcoded libraries were demultiplexed and barcodes were removed with lima (v2.7.1). Resultant reads.bam along with subreads.bam files were input to DeepConsensus (v1.2) for generation of fastq HiFi (>Q20) data.

#### PacBio Kinnex Sequencing

Total RNA was extracted from flash-frozen cell pellets using a commercial kit: Qiagen RNeasy Mini kit (Qiagen p/n 74104) using harmonized protocols across UW, WUSTL, and Rockefeller. RNA quality assessment followed equivalent criteria at all centers, although instrumentation differed slightly; all samples were evaluated with the RNA 6000 Nano kit on an Agilent 2100 Bioanalyzer (Agilent p/n 5067-1511), and Rockefeller additionally assessed RNA quality using an Agilent Fragment Analyzer using an RNA kit (Part Number: DNF-471-0500). RIN scores were very high with almost all samples > 9.0. 250-300 ng of each RNA sample was used as input and processed with the Iso-Seq express 2.0 and Kinnex full-length RNA kits according to manufacturer’s recommendations (PacBio p/n 103-071-500 & 103-072-000). Briefly, full-length cDNAs were generated using a poly-dT RT primer and a template switch oligo to add directional universal priming sites to each molecule, then the cDNA was amplified and a sample barcode added using PCR. After cleanup and QC, equal molar amounts of eight different sample cDNAs were pooled into each Kinnex pool, then split into eight separate PCR reactions to add directional sequence handles. The eight monomers were repooled and ligated using the directional handles and added SMRTbell adapters into 8-mer Kinnex arrays. An exonuclease treatment removed imperfect SMRTbell templates and the resulting libraries were prepared for sequencing on a Revio instrument using the release chemistry (PacBio p/n 102-739-100 & 102-202-200) on SMRT Link version 13.0 or 13.1 and 24- or 30- hour movies. All sequencing centers used a standardized PacBio Kinnex workflow targeting approximately 10 million full-length non-concatemer (FLNC) reads per sample. Variability in sequencing output between runs resulted in some samples exceeding the target FLNC yield, producing deeper transcriptome coverage and increased detection of low-abundance transcript isoforms. These differences in sequencing depth contributed to modest center-associated batch effects. Downstream analyses included normalization and batch correction to account for sequencing-depth differences and minimize center-associated technical effects. Raw sequencing reads were processed in SMRT Link using the “Read Segmentation and Iso-Seq” pipeline which splits each concatenated read into its component monomers, demultiplexes the reads by the cDNA sample barcodes, and filters out artifacts by searching for the expected primer sequences and poly-A tail. The resulting FLNC reads were used for all downstream analyses.

#### Fiber-seq: PacBio

Fiber-seq lysates prepared at Coriell were treated in the same way as standard HiFi preps processed in Year 5 as described above, with the exception that the extracted DNA tended to have carryover of sodium dodecyl sulfate (SDS, used as a stop solution for the Fiber-seq tagging). If DNA samples appeared sudsy during shear, 1/100 volumes of 1% Triton X-100 in water was added to the post-shear bead wash to remove residual SDS before prep. After sequencing, Fiber-seq samples were additionally processed with the fibertools pipeline to call m6A marks and open chromatin regions^154^.

#### Oxford Nanopore R9 Sequencing

High molecular weight (HMW) DNA was extracted from 16 million cells using the Circulomics Nanobind CBB Big DNA Kit (NB-900-001-01) and UHMW DNA Auxiliary Kit (NB-900-101-01) according to the manufacturer’s instructions (Document ID: EXT-CLU-001), with modifications to accommodate increased input. Reagent volumes were scaled accordingly, and DNA was eluted in a final volume of 2,250 µL. All steps were performed in 5 mL tubes. Libraries were prepared using the Circulomics UL Library Prep Kit (NB-900-601-01) in combination with the Oxford Nanopore Technologies Ultra-Long DNA Sequencing Kit SQK-ULK001 following the manufacturer’s protocol with proportional scaling of reagents. Libraries were eluted in 675 µL to enable three loading events across three flow cells. Sequencing was performed on a PromethION 48 using R9.4.1 flow cells (FLO-PRO002). Three libraries were loaded per flow cell, with washing (ONT Wash Kit, EXP-WSH004) and reloading every 24 h for a total runtime of 72 h. ONT sequencing data generated on R9.4.1 pore flowcells on PromethION and GridION instruments were basecalled using Guppy (v6.4.6 and v6.5.7) in super-accuracy mode with simultaneous 5-methylcytosine (5mC) modification calling at CpG sites (dna_r9.4.1_450bps_modbases_5mc_cg_sup_prom configuration). A subset of 5 runs were basecalled with Guppy (v6.3.7 and v6.5.7) using an extended model that additionally called 5-hydroxymethylcytosine (5hmC) at CpG sites (dna_r9.4.1_450bps_modbases_5hmc_5mc_cg_sup_prom configuration). Basecalling produced unaligned BAM files with embedded base modification tags.

#### Oxford Nanopore R10 Sequencing

For early R10 sequencing, 50 million cell pellets were split into three aliquots and processed using the NEB HMW DNA Extraction Kit for Tissues (T3060), following the Oxford Nanopore Technologies genomic DNA extraction protocol (v114_revL_27Nov2022). DNA was eluted in 2,250 µL elution buffer (EEB) to accommodate increased input. Libraries were prepared using the Oxford Nanopore Technologies Ultra-Long DNA Sequencing Kit SQK-ULK114 with reagent volumes scaled threefold and processed in 5 mL tubes. During precipitation, DNA was pelleted at 2,000 × g for 3 min (twice) in place of the precipitation star. Libraries were eluted in 675–800 µL to support 3–4 loading events across three flow cells. Sequencing was performed on the PromethION 48 using R10.4.1 flow cells (FLO-PRO114M), with flow cells washed and reloaded every 24 h for total run times of 72–96 h. R10.4.1 pore flowcell data generated on PromethION instruments were basecalled using Dorado (v0.6.0) with the super-accuracy model dna_r10.4.1_e8.2_400bps_sup@v4.3.0. Simultaneous methylation calling for 5-methylcytosine and 5-hydroxymethylcytosine at CpG sites (5mCG_5hmCG) was performed during basecalling. Basecalling produced unaligned BAM files with embedded base modification tags.

#### Oxford Nanopore R10 Sequencing (Updated Protocol)

HMW DNA was extracted from ∼20 million cells using the NEB HMW DNA Extraction Kit for Tissues (T3060), following the ONT genomic DNA extraction protocol with modifications to preserve ultra-high molecular weight DNA. Proteinase K and RNase A incubation times (10–30 min) were adjusted based on sample homogeneity. DNA was eluted in 200 µL EEB, incubated on beads overnight, and adjusted to a final volume of 750 µL. Extractions were performed at least 3 days prior to library preparation and stored at 4 °C. Libraries were prepared using the Oxford Nanopore Technologies Ultra-Long DNA Sequencing Kit (SQK-ULK114) without scaling reagent volumes to maximize read length. Libraries were eluted in 1,020 µL to enable up to 15 loading events (five loads per flow cell). Each load consisted of 67.5 µL library, 7.5 µL Loading Solution UL, and 75 µL Sequencing Buffer UL. Additional reagents from the ONT Ultra-Long Auxiliary Kit (EXP-ULA001) were used as needed. Libraries were prepared up to one week prior to sequencing and stored at 4 °C to allow homogenization. Sequencing was performed on the PromethION 48 using R10.4.1 flow cells (FLO-PRO114M), with 2–3 flow cells per sample. Libraries were loaded sequentially (4–5 loads per flow cell), with washing and reloading every 24 h for total run times of 96–120 h. Sequencing runs performed after January 2024 incorporated ONT E8.2.1 adapter chemistry.

#### Fiber-seq: Oxford Nanopore

Fiber-seq lysates were prepared and provided by Coriell. HMW DNA was extracted using the NEB HMW DNA Extraction Kit for Tissues (T3060), following the ONT genomic DNA protocol with modifications, including standard-bore pipette mixing to reduce viscosity. DNA was eluted in 750 µL EEB.

Libraries were prepared using the Oxford Nanopore Technologies Ultra-Long DNA Sequencing Kit SQK-ULK114 and eluted in 340 µL to enable up to five loading events per sample.

Sequencing was performed on the PromethION 48 using R10.4.1 flow cells, with sequential loading (five loads per flow cell) and wash/reload cycles every 24 h for 120 h. Fiber-seq ONT data were basecalled using Dorado (v1.1.1) with the super-accuracy model sup@v5.2.0.

Simultaneous modification basecalling was performed for

5-methylcytosine/5-hydroxymethylcytosine (5mCG_5hmCG) and N6-methyladenine (6mA) to support Fiber-seq chromatin accessibility analysis, producing unaligned BAM files with embedded base modification tags.

#### Illumina Omni-C Sequencing

Omni-C libraries were prepared from cell lines using the Dovetail™ Omni-C™ Kit (PN 21005G) according to the manufacturer’s protocol with minor modifications. Chromatin was fixed in nuclei, digested in situ with DNase I, and captured on chromatin beads. Fragment ends were repaired, ligated to biotinylated bridge adapters, and proximity-ligated. After crosslink reversal and DNA purification, excess biotin was removed. Libraries were constructed using the NEB Ultra II DNA Library Prep Kit with Illumina-compatible adapters, and biotinylated fragments were enriched on streptavidin beads. Libraries were split into two indexed replicates prior to PCR to maintain complexity and sequenced on an Illumina NovaSeq 6000.

#### Illumina Whole Genome Sequencing

The HPRC utilized high coverage Illumina data from the 1000 Genomes project ^38^ when available for sample data as well as for parental data for samples which are the child in a trio pedigree. For samples without Illumina WGS data available (NA21309, HG01123, HG02486, HG02559, HG03471. Parents of NA21309: NA21307/NA21307 were also sequenced)), the HPRC extracted DNA, created Illumina TruSeq PCR-free libraries as described previously. Libraries were sequenced to a targeted depth of 30X using an Illumina NovaSeq 6000 sequencer.

### Assembly

#### Assembly Creation

Assemblies were generated with Hifiasm v0.19.7 or v0.19.9 with PacBio HiFi reads as input with ultralong integration. In cases where both CCS and DeepConsensus basecalled data was available for PacBio reads, DeepConsensus was used. Assemblies were created with a WDL pipeline that incorporates preprocessing (https://github.com/human-pangenomics/hpp_production_workflows/blob/master/assembly/wdl/workflows/hic_hifiasm_assembly_cutadapt_multistep.wdl or https://github.com/human-pangenomics/hpp_production_workflows/blob/master/assembly/wdl/workflows/trio_hifiasm_assembly_cutadapt_multistep.wdl). PacBio Hifi reads containing adapter sequences were removed with Cutadapt with the --discard-trimmed option. When parental Illumina data was available for trio phasing, yak k-mer databases were created for the the parental and sample Illumina data for input to Hifiasm. Samples without parental Illumina data were phased with Hi-C. Hifiasm was called with --ul-cut 50000 --dual-scaf --telo-m CCCTAA. Assemblies from male samples with Hi-C phasing had sex chromosomes re-assigned with yak’s sexchr and groupxy.pl commands. chromosomes Y and X were assigned to haplotype 1 and 2, respectively.

#### Assembly Polishing With DeepPolisher

In order to correct base-level errors in the assemblies, we applied a polishing pipeline based around DeepPolisher as described in ^47^. Briefly, DeepPolisher is an encoder-only transformer model that uses HiFi read alignments to an assembly to identify errors and suggest polishing edits. To produce the input HiFi read alignments we used the PHARAOH pipeline, which takes in ONT UL information to correct the phasing of reads in long stretches of false homozygosity in the assemblies, enabling DeepPolisher to make corrections in those regions.

We ran the PHARAOH pipeline using default parameters as described in ^47^, and DeepPolisher using docker container google/deepconsensus:polisher_v0.0.8_12122023 with model “checkpoint-665” and default parameters. As detailed in, the pipeline applies a conservative set of GQ filters to the polishing edits vcf before applying them to the assemblies (GQ 20 for 1 bp insertions, GQ 12 for 1 bp deletions, and GQ 5 for all other variants). For each sample, we only used input HiFi reads with DeepConsensus v1.2, or HiFi Revio reads (which uses DeepConsensus v1.1) in order to match the input data the DeepPolisher model was trained on. Coverage did not drop below 30x total for any assembly despite this filtering. The DeepPolisher pipeline added an average of 12,806 edits per assembly (**Supplementary Table 8**).

#### Assembly Cleanup

Assemblies were prepared for upload to Genbank with a WDL that removes contamination and re-assembles the mitochondrial contig. Briefly, foreign contamination from exogenous DNA such as EBV and residual adapter sequences was identified with FCS-GX^155^ and removed from assemblies. Mitochondrial DNA breaks Hifiasm’s diploid coverage assumptions creating multiple assembled contigs that do not reflect the true sequence. Mitochondrial assemblies not integrated into the rest of the genome as NUMTs were identified with BLAST and removed. Mitochondrial reads were then identified and input to a modified version of MitoHifi^156^. A representative mitochondrial assembly was appended to haplotype 2 for Hi-C phased assemblies, or the maternal haplotype for trio phased assemblies. Assembly metadata is available in **Supplementary Table 16**.

#### Initial Assembly QC

High-level assembly quality control metrics were assessed with multiple externally available tools in an automated workflow https://github.com/human-pangenomics/hpp_production_workflows/blob/master/QC/wdl/workflows/assembly_qc.wdl (commit d73431d). Non-assembly inputs required for the workflow are available as a Zenodo archive (10.5281/zenodo.20752648). Component parts, and pertinent details of the workflow are described below.

Contiguity (N50 and AUN) metrics were calculated with calN50.js (r4)^157^. T2T sequences were identified by aligning assembled sequences against a reference genome (CHM13v2.0) and determining if the sequence represents the full target chromosome and has telomeric repeats on both ends. Alignment was performed with MashMap^158^ v3.1.3 with --perc_identity 95 --noSplit options against CHM13v2.0. Telomeric repeats and assembly gaps (runs of N’s) were identified with the telo (-s 10) and gap (-l 2) commands from seqtk^159^ v1.4. Alignments that were entirely in the regions of either the pseudoautosomal regions in the sex chromosomes or the p-arms in acrocentric chromosomes (p-arm telomere up to and including the rDNA arrays) were discarded. T2T status was assigned when assembled sequences had both telomeric repeats within 10kbp of both ends of the sequence and alignment blocks that cover >95% of the reference chromosome’s length. T2T sequences with gaps were classified as T2T scaffolds, and T2T sequences without gaps were classified as T2T contigs.

Single copy gene presence and multicopy gene collapses were assessed with Compleasm^160^ v0.2.6 and the asmgene command in paftools.js in minimap2 v2.23^161^. Compleasm was run using the primates odb10 BUSCO lineage with modifications to remove genes missing or fragmented in CHM13 and to create sex-chromosome specific lineages. A database for assessing chrY containing haplotypes was then created by further removing genes found on chrX, and a database for assessing chrX was created by further removing genes found on chrY. The lineages and lists of removed genes are available on Zenodo (10.5281/zenodo.20752648). Asmgene was run with cDNA from Ensembl genes (Homo_sapiens.GRCh38.cdna.all.fa) mapped against the assembly compared against precomputed transcripts mapped to GRCh38 (hs38.paf).

Large misjoins in the assemblies were identified by aligning each assembly to CHM13v2.0 with minigraph v0.21^58^ using -xasm followed by running paftools.js misjoin (minimap2 v2.28) on the resultant PAF file. Interchromosomal misjoins (mappings where a sequence maps to two chromosomes with both ends not landing in centromeres, acrocentric p-arms, or PAR regions) were selected from the output files.

K-mer-based quality value (QV) scores were computed with yak v0.1^162^ and Merqury^163^ (Meryl v1.4) with a default k-mer size of 31. Illumina WGS reads were used as input except in the case of HG02109 where no Illumina data was available and Illumina Hi-C reads were used. Trio-phased assemblies had phasing accuracy (switch and hamming error rates) assessed with yak trioeval using k-mer databases built from paternal and maternal reads. Switch estimates were also obtained with Merqury similarly. QC results are available in **Supplementary Table 17**.

#### Assembly Quality Control: HMM-Flagger

HMM-Flagger is a reference-free, read-mapping-based tool for evaluating phased diploid assemblies. It takes long read alignments to an assembly as input and applies a hidden Markov model (HMM) to detect coverage anomalies along the assembly and classify different types of large structural errors. The error categories include false duplications (Dup), collapses (Col), and erroneous (Err) blocks.

We ran HMM-Flagger (v1.2) on 231 HPRC2 diploid assemblies using both PacBio HiFi and ONT reads (with either R9 or R10 chemistry). Long reads were mapped to each diploid assembly using minimap2. For HiFi and ONT R9.4.1 reads, we used the parameters -k 25 -ax lr:hqae --cs --eqx -L -I8g, whereas for ONT R10.4 reads we used -k 15 -ax map-ont --cs --eqx -L -I8g. Minimap2 (v2.28) was run using the Docker image mobinasri/long_read_aligner:v1.1.0. Reads were mapped simultaneously to both assembly haplotypes (the parameter -I8g was specified for this purpose).

The resulting BAM files were then processed with the correct_bam program (from the docker image mobinasri/secphase:v0.4.4) using the parameters --primaryOnly --minReadLen 5000 --minAlignment 5000 --maxDiv 0.1, which filter out reads or alignments shorter than 5 kb, secondary alignments, and alignments with gap-compressed divergence greater than 0.1.

HMM-Flagger was run using the Docker image mobinasri/flagger:v1.2.0. We used the HMM-Flagger presets --hifi for HiFi reads, --ont-r9 for ONT R9.4.1 reads, and --ont-r10 for ONT R10.4 reads. To account for potential platform-specific coverage biases in peri/centromeric satellite regions, we provided CenSat annotations to HMM-Flagger. These annotations were generated with the following workflow:

https://github.com/kmiga/alphaAnnotation/blob/main/cenSatAnnotation/centromereAnnotation.w dl

The paths to CenSat annotations were specified in a JSON file and provided to bam2cov, a program that generates coverage files out of BAM files for running HMM-Flagger (More details in https://github.com/mobinasri/flagger). The --runBiasDetection option was enabled in bam2cov to identify and mark biased regions in the coverage files passed to HMM-Flagger.

For mapping reads and running HMM-Flagger we used the this WDL, which encapsulates the commands and parameters explained earlier: https://github.com/mobinasri/flagger/blob/main/wdls/workflows/hmm_flagger_end_to_end_with_mapping.wdl

An example of a json file for running the WDL with HiFi reads is available here (Please note that the HMM-Flagger version in this JSON is v1.1 however HMM-Flagger was later rerun with v1.2 for the finalized BED files):

https://github.com/human-pangenomics/hprc_intermediate_assembly/blob/main/assembly_qc/batch2/hmm_flagger/hifi/runs_toil_slurm/hmm_flagger_end_to_end_with_mapping_input_jsons/HG00706_hmm_flagger_end_to_end_with_mapping.json

And here is an example for ONT reads:

https://github.com/human-pangenomics/hprc_intermediate_assembly/blob/main/assembly_qc/batch2/hmm_flagger/ont/runs_toil_slurm/hmm_flagger_end_to_end_with_mapping_input_jsons/HG00706_hmm_flagger_end_to_end_with_mapping.json

For each diploid assembly, HMM-Flagger outputs two main BED files. The first contains the inferred errors (labeled Err, Dup, and Col) directly derived from the posterior probabilities of the HMM. The second BED file, referred to as the “conservative” set, has higher specificity and is generated by filtering the initial BED file using self-homology mappings of assembly contigs against themselves. In cases of false duplication, we expect multiple contigs to map to the falsely duplicated region, whereas collapsed regions are expected to lack mappings from other contigs. These two principles are used to filter the original error calls as the last part of the HMM-Flagger pipeline. We used the conservative BED file for all HMM-Flagger-based analyses in this study.

#### Assembly Quality Control: NucFlag

To provide an orthogonal method of assembly error detection, we ran NucFlag v0.3.3^57^ using the same PacBio HiFi read alignments used in HMM-Flagger’s validation workflow. NucFlag was run with the following command: nucflag -i {input.bam_file} -d {output.plot_dir} -o {output.misassemblies} -t {threads} -p {processes} --chrom_sizes {input.chrom_sizes} -c {input.config} --output_cov_dir {output.cov_dir}. The full config file used in this analysis is available at

https://github.com/koisland/Snakemake-NucFlag-HPRC/blob/main/config/nucflag.toml.

Briefly, NucFlag v0.3.3 identifies candidate assembly errors within genome assemblies based on inconsistencies in the genome assembly and native long-read sequencing data. To do this, it uses peak/valley detection in the first- and second-highest read pileup nucleotide frequency of aligned PacBio HiFi reads to detect misjoins (where two contigs are erroneously joined together) and collapses in sequence (where two or more highly identical sequences are collapsed into a single representative sequence). NucFlag considers any region with a drop in read depth of >95% of the mean coverage or zero supporting reads as a candidate misjoin (MISJOIN). Regions with a nucleotide frequency 250% times the mean coverage are considered candidate collapses (COLLAPSE), and if intersecting a heterozygous site, a collapse with variation (COLLAPSE_VAR). Additionally, regions with a peak in coverage and a nucleotide frequency ratio greater than 20% of the mean coverage are considered candidate small collapses (COLLAPSE_OTHER); regions below this threshold are potential heterozygous sites and typically omitted (HET). NucFlag outputs a list of candidate assembly errors and their predicted error class as a BED file, and it produces a plot to show the read depth across the region and the precise location of the assembly errors to aid in visualization.

#### Comparison of centromeres in HPRC Release 1 and 2

To compare the contiguity and accuracy of centromeres between the HPRC Release 1 and 2 genome assemblies, we first selected 40 samples shared between the two releases, of which half are male and half are female and span global populations (**Supplementary Table 18**). We then ran CenMAP v1.2.0^57^ (commit `80d553f9e88172d4fc77ce889c3637041dbc567`) on these 40 samples with default settings to detect completely assembled centromeric α-satellite HOR arrays with the following commands: cenmap -c config.yaml -j 30 --workflow-profile workflow/profiles/lpc/

We identified centromeres that were accurately assembled in the Release 1 and 2 datasets by intersecting the coordinates of the active α-satellite HOR array with NucFlag’s error calls, excluding HETs, using BEDtools^164^ (v2.31.1) and a custom script, https://github.com/koisland/HPRC_R1vR2_centromeres/blob/main/scripts/nucflag/filter_arrays.py. Any centromere without a NucFlag error call was considered accurately assembled, while those with a NucFlag error call were considered to be misassembled.

We then used a custom script, https://github.com/koisland/HPRC_R1vR2_centromeres/blob/main/scripts/complete_counts/count_complete_cens.py, to determine the number of complete and accurately assembled centromeres, where completeness was defined as having no breaks in contiguity across the α-satellite HOR array, and accuracy was defined as having no intersection with NucFlag’s non-HET calls. This script is run with the following command: count_complete_cens.py -a Release1_AS-HOR_lengths.bed -b Release2_AS-HOR_lengths.bed -o {output.png}

### 8p23.1 Assembly to Reference Alignment

We have aligned assemblies for 47 samples generated as part of both HPRC release 1 and 2 using minimap2 (v2.24 and v2.28) to the human reference genome T2T-CHM13 with the following parameters: -x asm20 --secondary=no -s 25000 -K 8G. The resulting PAF files were loaded using the SVbyEye^165^ function readPaf. Then PAF alignments for each haplotype were subsetted to the region of interest (chr8:6000000-13700000) using SVbyEye function subsetPafAlignments. Next the directionality of each assembled haplotype was synchronized such that there are always directly oriented alignments at the region boundaries. We have extracted only matched bases between query and reference genome by parsing the CIGAR strings for each alignment. Last we have removed small contigs whose alignments were completely embedded within other larger contigs. Such alignments were then used for the final visualisation of the 8p23.1 region.

#### Assembly QC Integration

To generate integrated structural QC bed tracks, three distinct QC bed files were used: Flagger-HMM predictions from HiFi alignments, Flagger-HMM predictions from ONT alignments, and NucFlag predictions from HiFi alignments. Bed intervals from the three source bed file were intersected with a 100kb merge distance. Merged regions with predicted assembly issues from more than one source were labeled as “error” and interpreted as probable assembly errors. Regions with predicted assembly issues from only one source were labeled as “qc_flag” and not included in estimates of assembly errors.

#### Assembly QC Projection

Assembly sequences were projected onto the CHM13 v2.0 reference using minigraph v0.21-r606 with the -xasm preset. The resulting PAF alignments were filtered, merged, bridged across large alignment gaps, and then extended to sequence termini using custom scripts. This process created projection chains for determining which regions on CHM13 are covered by an assembly and which have QC errors.

Alignments with identity lower than 70% were removed (except in chrY where a cutoff of 40% was used). Alignment blocks shorter than 50 kb were filtered out unless they occurred within the terminal 200 kb of a query sequence. Remaining alignment blocks on the same strand within 200kb on both the query and reference were merged. For each assembled sequence (query) the alignment with the largest total query coverage was retained. To prevent false dropouts in large diverged sequences, alignments were interpolated when possible using flanking alignments. Similarly, alignments were extended to the query/reference ends when they fell within 10% of the reference chromosome end.

Coverage on CHM13 was calculated from the final projection; regions without any assembly projections were deemed “no_coverage”. Intersected structural QC regions in the assembly’s coordinate system were then projected onto CHM13 using the projection represented as a chain file. The projected QC were combined with no_coverage regions to define unreliable regions.

To assess the number of high quality local reference haplotypes at each region across the genome, we tiled T2T-CHM13 into 500kb syntenic windows (satellite arrays such as rDNA and alpha satellites which were larger than 500kb and regions which lacked clear synteny to T2T-CHM13 were expanded to form larger windows with clear bounding alignments) and counted the number of haplotypes that cover the genomic windows without assembly gaps or error flags. As compared to calculating the sum of QC flagged bases from individual tools, this approach increases the fraction of the genome tallied as unreliable (because a flag or break in a window marks the whole interval in the given haplotype), but better reflects the fact that assembly gaps and modestly sized error flags can create large regions unusable for analysis

In order to calculate the number of assembly sequences needed to represent each chromosome in CHM13, the final projections were counted when they contributed unique coverage on a chromosome. Only assemblies whose projections cover over greater than 90% of the chromosome after interpolation and extension were counted. For HPRC2 assemblies, the count was increased by the number of N gaps.

#### Assembly Contiguity

NG50 plots with T2T-CHM13v2.0 and GRCh38 no-alt analysis (GCA_000001405.15_GRCh38_no_alt_analysis_set) included as references. For T2T-CHM13v2.0, chrY was excluded; for GRCh38, unlocalized/unplaced contigs and EBV were excluded. GRCh38 was split at annotated gap and centromere intervals. For each HPRC assembly, pre-QC contig lengths were read directly from .fai files. Post-QC contig lengths were generated by splitting assembly sequences at either scaffold gaps (Ns) or regions with QC_error flags (see QC Integration). NG50 was plotted in Python by sorting contig lengths from longest to shortest and recording the contig length observed across a 3.1 Gb cumulative genome-size grid. Across assemblies, the median and 10th-90th percentile bands were calculated before and after QC fragmentation.

#### Chromosome Assignment

Chromosome assignment for the assemblies was generated using a WDL pipeline (annotation/wdl/tasks/assign_chromosomes.wdl) which aligns each assembly to a reference (CHM13 v2.0) with MashMap^158^, identifies gaps and telomeres with seqtk, and produces chromosome alias and T2T chromosome assignment files.

### Pangenome Variation

#### Pangenome Growth

The growth plots were calculated using Panacus^166^ on the autosomal chromosomes of GRCh38-based MC graphs by excluding nodes covered by the GRCh38 reference. The growth curves were calculated using `panacus section-growth` with the parameters `--groupby-haplotype -l 1,2,2,2 -q 0,0,0.05,0.95`. The `-e` parameter was used to exclude nodes covered by the reference sequence. Corresponding plots for the CHM13-based MC graphs are shown in **Supplementary Fig. 13**.

For the plot, we employed various combinations of thresholds for minimum coverage and quorum, defined as the minimum ratio of the number of haplotypes in which a sequence must appear. Specifically, we set the coverage to 1 and quorum to 0% for “Cumulative all”, coverage to 2 and quorum to 0% for “Depth ≥ 2”, coverage to 2 and quorum to 1% for “Common”, and coverage to 2 and quorum to 95% for “Core”.

Panacus’ `section-growth` command calculates growth plots containing multiple groups of haplotypes (in this case HPRC1 and HPRC2). The first part of the growth curve was calculated with only the HPRC1 haplotypes as described in ref. ^166^, while the second part of the growth curve made use of a new algorithm. This new algorithm constructs a 3D histogram with the x-axis containing the number of occurrences in the first group (HPRC1) and the z-axis containing the number of occurrences in the second group (HPRC2). The y-axis contains the number of basepairs for which this combination of occurrence numbers is true. Then the growth curve could be calculated using the formula:

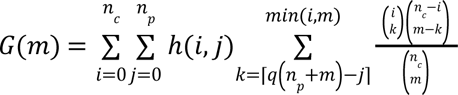

where *n_c_* is the number of haplotypes in the second group, *n_p_* in the first group and ℎ(*i*, *j*) is the 3D histogram that gives the number of basepairs for which a given pair of occurrences holds true. This approach scales to an arbitrary number of groups.

For the proportion of variants that are covered already by the current pangenome, we used the method for calculation sample coverage as described in ref.^62^, including the corresponding rarefaction procedure. This assumes that samples are drawn randomly with replacement from the population. However, since the number of the underlying population is large as compared to the sample size, drawing with replacement can be reasonably assumed. Since the variants in HPRC2 were not drawn randomly but rather in a manner that maximizes their diversity, this analysis may overestimate the extent of variation in the population and consequently underestimate the proportion that has already been covered.

#### Multi-MUM Conservation

Multi-Maximal Unique Matches (multi-MUMs) are non-extendible exact matches that are present in each sequence of a collection and unique within each sequence. They have been used to compute core genome alignments^64^ and represent perfectly conserved columns in the underlying pangenome multiple alignment. Mumemto^63^ can compute multi-MUMs efficiently for large pangenome collections, identifying collinear blocks of exact matches that form a proxy for the multiple alignment. We use multi-MUM coverage to measure the proportion of the pangenome collection that is “alignable”, i.e. the number of bases contained in collinear multi-MUM blocks. We consider two multi-MUMs to be part of a collinear block if they are adjacent in every assembly, separated by less than 100 bp, and in the same order across the pangenome (or swapped if on the reverse strand). To compute the change in multi-MUM coverage for growing pangenome collections, we computed coverage over 10 random subsets of the HPRC2 collection at various subset sizes by merging pairwise multi-MUMs from each assembly with respect to T2T-CHM13^167^.

#### Variant site composition

To enable direct comparison between the HPRC1 and HPRC2 MC graphs, variant sites on the autosomes were defined and annotated as described in ref.^13^. Variant sites were obtained from VCF files generated by `vg deconstruct`^168^. Some non-reference nodes present in the graph are absent from the corresponding VCF representation because of how `vg deconstruct` generates raw VCF output, resulting in differences between graph- and VCF-derived non-reference sequence lengths. For each variant site, `vg deconstruct` reports at most two alleles per sample. When more than two alleles are present for a sample at a site, only two are retained in the VCF. Consequently, nodes belonging to excluded alleles are not represented in the VCF, contributing to missing non-reference sequence. Large spurious variant sites with reference alleles longer than 10 Mb were removed using vcfbub^169^ with parameters `-l 0 -r 10000000`, which kept top-level sites while preserving nested child sites from filtered parent sites. Sites were then classified as small variants (<50 bp) or SVs (≥50 bp) based on the length of the longest allele at each variant site.

For SV sites, alleles were grouped to represent the underlying SV event rather than each distinct allele sequence, because additional mutations within an SV sequence, such as SNPs within an insertion, can produce technically distinct but closely related alleles. To avoid artificially splitting a single SV event into multiple categories, alleles were grouped greedily after sorting by decreasing frequency. Each allele was assigned to the first existing group containing at least one similar allele; otherwise, a new group was created. Similarity between two alleles was defined as a length difference of less than 50 bp.

After grouping, the longest allele sequence from each SV site was extracted and written in FASTA format. Sequence features, including interspersed repeats, low-complexity regions, tandem repeats, centromeric satellites, and gaps, were identified using RepeatMasker (v4.1.2-p1; NCBI/RMBLAST v2.10.0; Dfam v3.3), SDUST (v0.1; https://github.com/lh3/sdust), ETRF (commit fc059d5; https://github.com/lh3/etrf), dna-brnn (v0.1)^170^, and `seqtk gap` (v1.3; https://github.com/lh3/seqtk), respectively. SDs were defined for sites with total node length ≥1,000 bp, for which at least 20% of bases were annotated as SDs in either the reference genome or the individual assemblies. To identify homologous regions in the GRCh38 reference genome, the longest allele sequence for each SV site was aligned using minimap2 (v2.24)^171^ with parameters `-cx asm20 -r2k --cs`. Based on these annotations, SV sites were assigned to repeat categories using a modified version of `mgutils.js anno` adapted to support inputs derived from MC graphs (https://gist.github.com/wwliao/60fd10701a4040da8f93e1db0513f2c3).

#### Graph variant benchmarking

##### Variant calling

Variants were identified using 14 variant callers. Among the 14 callers, 10 were applied to PacBio HiFi reads: longcallD (v0.0.4)^172^, DeepVariant (v1.6.1)^173^, cuteSV (v2.1.1)^174^, DeBreak (v1.3)^175^, DELLY (v1.3.2)^176^, pbsv (v2.10.0; https://github.com/PacificBiosciences/pbsv), sawfish (v0.12.8)^177^, Sniffles2 (v2.5.3)^178^, SVDSS (v2.0.0)^179^, and SVIM (v2.0.0)^180^; 4 were applied to haplotype-resolved assemblies: dipcall (v0.3)^181^, PAV (v2.4.6)^14^, cuteSV-asm (v2.1.1), and SVIM-asm (v1.0.3)^182^. Here, cuteSV-asm refers to cuteSV applied to assemblies and is distinguished from its application to PacBio HiFi reads.

For assembly-based variant calling, dipcall was used to align assemblies to the reference genome (GRCh38 and T2T-CHM13 v2.0) using Winnowmap2 (v2.03)^183^ and to derive variant calls from the resulting alignments. These alignments were also used as input for additional assembly-based callers, including cuteSV-asm and SVIM-asm. Dipcall also provides per-sample BED files defining confident regions, which were used in downstream analyses. PAV follows a similar strategy but uses minimap2 (v2.26)^171^ for alignment.

For read-based variant calling, PacBio HiFi reads were aligned to the reference genome using Winnowmap2, and variants were subsequently identified using the read-based callers listed above. For SVDSS, genotypes were re-estimated using kanpig (v1.1.0)^184^ due to unreliable genotyping.

To increase sensitivity, minimum mapping quality and read support thresholds were relaxed to 5 and 3, respectively, where applicable to SV callers.

All variant calling analyses were implemented in reproducible workflows using Nextflow^185^. The workflows and index files are available at https://github.com/wwliao/hprc_release2_variant_calling, where software versions and analysis parameters are provided.

##### Variant callset merging

To construct a baseline for benchmarking graph variants, per-sample merged callsets were generated by integrating results from multiple variant callers. Existing approaches typically maintain separate merged callsets for small variants and SVs, which are evaluated using different benchmarking tools. However, this separation introduces representation-dependent inconsistencies. For example, a single large deletion may be represented either as one SV or as multiple small variants, and representations of the same event may differ across callsets. These discrepancies complicate downstream callset comparison and benchmarking. To address this, we constructed merged callsets that jointly include small variants and SVs, enabling comparison at the haplotype sequence level.

Variant callsets from all callers were converted to sequence-resolved form, and merged callsets were constructed using Aardvark (v0.10.4)^186^, which compares variants based on reconstructed haplotype sequences rather than direct matching of VCF records.

Three callers that generate joint callsets containing both small variants and SVs (dipcall, PAV, and longcallD) were used to define a backbone callset. Aardvark groups nearby variants into clusters and compares reconstructed haplotype sequences across callers. Because longcallD reports phased genotypes that are valid only within local phase blocks, these genotypes were unphased prior to merging to avoid inconsistencies during haplotype reconstruction across genomic regions. When multiple callers produced identical haplotype sequences, a predefined priority order (dipcall > PAV > longcallD) was used to select the representative variant set. If no agreement was observed among callers, variants from the highest-priority caller were retained.

To incorporate evidence from all callers, each caller-specific callset was compared to the backbone callset using Aardvark to evaluate haplotype-level concordance. Unlike the merging step, which requires identical haplotype sequences, this comparison allows partial mismatches quantified by basepair-level recall and precision. A caller was considered to support a variant if both basepair-level recall and precision exceeded predefined thresholds, which were selected based on caller type. Thresholds of 0.9 were used for joint callers (dipcall, PAV, and longcallD), 1.0 for the small-variant caller DeepVariant, and 0.5 for SV callers. For comparisons involving DeepVariant, clustering distance was reduced from 1,000 bp to 50 bp to minimize mismatches caused by nearby SVs. For SV callers, small variants from DeepVariant were incorporated prior to comparison to improve consistency.

Finally, caller support was aggregated across all comparisons to generate merged callsets. For each variant, supporting callers were recorded in the INFO/CALLERS field, and the number of supporting callers was recorded in INFO/NCALLERS. Variants were further annotated with INFO/SVTYPE and INFO/SVLEN for events ≥ 50 bp. All analyses were restricted to dipcall confident regions for each sample.

All variant merging analyses were implemented in reproducible workflows using Nextflow. The workflows and access to merged callsets are available at https://github.com/wwliao/hprc_release2_graph_variant_benchmarking.

##### Comparison of graph variants to joint ground truth

For each sample, a joint ground truth callset was derived from the merged callset by selecting variants supported by at least two callers (i.e., INFO/NCALLERS ≥ 2). This filtering step retains variants with consistent support across methods and defines the baseline for benchmarking.

Variant sites were derived from the HPRC1 and HPRC2 MC graphs using `vg deconstruct`. Large sites with reference alleles longer than 100 kb were filtered using vcfbub with parameters `-l 0 -r 100000`, which kept top-level sites while preserving nested child sites from filtered parent sites. The resulting sites were then decomposed into biallelic graph variants based on allele traversals using a modified decomposition workflow originally developed in PanGenie^29^. Unlike the original workflow, which filters sites according to the proportion of haplotypes with missing alleles before decomposition, our implementation disabled this filtering step while retaining the decomposition procedure. Finally, INFO/SVTYPE and INFO/SVLEN annotations were added using a custom script to ensure compatibility with downstream analyses.

Graph variants were then compared against the corresponding joint ground truth callset using Aardvark. Comparisons were restricted to dipcall confident regions for each sample. Performance was further evaluated across genomic contexts defined by the GIAB genomic stratifications resource (v3.6)^187^, enabling assessment in regions such as segmental duplications and tandem repeats.

All variant benchmarking analyses were implemented in reproducible workflows using Nextflow. The workflows are available at https://github.com/wwliao/hprc_release2_graph_variant_benchmarking.

#### Pantree-based pangenome coordinates

With the pantree algorithm, we traverse a MC graph with depth-first search (DFS). Standard DFS results in a spanning tree of the vertices in the graph. This tree is taken as the *reference tree*. A *reference edge* denotes a directed edge present in the reference tree; a *variant edge* denotes a graph edge absent from the reference tree. Each variant edge represents a non-reference allele.

In a MC graph, each chromosome in the reference genome corresponds to one connected component and the linear reference genome does not involve cycles. During DFS, we start from the beginning of one chromosome and always traverse the linear reference first such that the linear reference genome is always part of the reference tree. As a result, a directed variant edge ending on the linear reference genome corresponds to an alternate allele in VCF. Pantree is backward compatible with the biallelic VCF representation.

Meanwhile, the reference tree extends the traditional linear coordinate system. For example, suppose there is a long insertion to the linear reference genome with multiple SNPs on the insertion. VCF would enumerate each distinct insertion allele in the population, combined with SNPs on it, and generate a long multi-allelic record. Due to the limitation of VCF, pantree cannot report the SNPs in standard VCF, either. It instead reports the SNP alleles and the graph vertices in the VCF INFO field such that we can still count off-linear variants and calculate allele frequencies in the population.

#### GRef pangenome coordinates

Beginning with a pangenome graph and reference sample, we sought to produce a minimal covering set of forward-strand, acyclic intervals using path fragments from other samples. The end result is equivalent to rGFA as produced by minigraph^58^, except it is computed post-hoc on a base-level graph rather than during iterative construction of a structural variation graph. We used a minimum fragment length threshold, defaulting to 50bp, to avoid the computational overhead of storing thousands of tiny path segments which would not usefully serve as reference contigs. We introduced functionality in vg v1.74.0 (`vg paths -u`) to greedily compute such a cover and add it to a vg-formatted graph as a new, reference sample. This means that the existing sample-selection interface in vg tools could be used to toggle between the linear reference (e.g., T2T-CHM13) and the pangenome reference (e.g., augref_CHM13), allowing the user to decide whether or not to use the extended coordinate system without changing graph.

The cover itself is computed greedily, starting with the chosen reference sample. The method then followed the same pattern as the VCF export described in HPRC1^13^, where snarls^168^ whose end points lie on a reference path are visited independently. All haplotype paths that fully span each snarl were then enumerated, and any intervals in these paths not already in the cover were added to it. The enumeration was performed in lexicographical order on path name, ensuring that adjacent snarls choose the same paths, reducing fragmentation. Nested snarls were processed in the same manner. A second pass greedily added uncovered path intervals, and all adjacent intervals were merged together, where possible, at the end. A second pass iterated over all uncovered nodes and used the same procedure to add them to the cover. Adjacent intervals were then greedily merged where possible. Intervals on different paths were merged together if there was some path in common shared by the underlying graph segments. Once the cover was complete, it was saved to the graph as a collection of paths, including copies of the original reference paths, corresponding to a new sample. A secondary table also produced that maps each contig to its source path coordinates in the graph, along with that path’s position relative to the base reference. This table was used to produce **Fig. 3h** and **3i**.

Since the pangenome coordinate contigs are stored as reference paths in the graph, existing vg tools that operate on reference coordinates can use them without further modification. We did add support for new VCF tags, RC, RS, and RD, to help use these features to `vg call` and `vg deconstruct`. They correspond to the base reference contig’s name, start and end coordinate respectively, and complement the existing tags that denote the snarl level (LV) and parent snarl name (PS) that describe the nesting. Logic was also added to ensure that genotypes are consistent in nested sites: if an SV insertion is heterozygous, for example, then a variant within it is effectively haploid. Star alleles can optionally be used to denote this, though it is not widely supported.

#### Segmental duplication identification

Repetitive sequences in each genome assembly were annotated and masked using three complementary strategies. Tandem repeats were identified with TRF (v4.1.0)^188^ using the parameters `trf [asm.fa] 2 7 7 80 10 50 2000 -l 30 -h -ngs`. General repeats were detected with RepeatMasker (v4.1.5)^189^, using the following options, `RepeatMasker -s-e ncbi -xsmall -species human [asm.fa]`. Low-complexity regions were further characterized using WindowMasker (v2.2.22)^190^ through a two-step process including, `windowmasker -mk_counts -mem 16384 -smem 2048 -infmt fasta -sformat obinary -in [asm.fa] -out [asm.count]`, followed by interval masking `windowmasker -infmt fasta -ustat [asm.count] -dust T -outfmt interval -in [asm.fa] -out [asm.interval]`. Outputs from all three methods were combined to produce a unified annotation set, and the assemblies were soft-masked based on these annotations.

SDs were then identified from the repeat-masked assemblies using SEDEF (v1.1)^191^. Candidate duplication pairs were required to be at least 1 kbp in length, exhibit more than 90% sequence identity, and contain less than 70% satellite-derived sequence. Regions overlapping Flagger^50^ and NucFlag^57,70^ annotations were excluded. Additionally, independent validation was performed with FastCN^71^, based on re-aligning short-read sequencing data to the respective genome assemblies, resulting in the final SD callset of 459 haplotype genome assemblies, with the available short-read data. Male sex chromosomes were processed independently. False SDs not supported by read depth were filtered out.

SDs were mapped to T2T-CHM13 v2.0 using a stepwise approach. Initially, SD intervals located within 10 kbp of each other were merged to form contiguous SD chains. These chains were then filtered to retain only those situated in confidently mappable regions, defined by overlap with alignment blocks of at least 100 kbp in length. Next, the retained SD chains were projected onto candidate homologous loci, requiring the presence of at least one unique flanking sequence of 10 kbp to support positional correspondence.

#### MEI calling

To identify polymorphic variants caused by mobile element insertions, we ran L1me-aid v1.3.4 (https://github.com/Markloftus/L1ME-AID) and an custom filter (https://github.com/xzhuo/INTACT_MEI) on the MC graph SV list to generate the list of all MEIs from the HPRC2. Based on the presence of each MEI in all assembled haplotypes, we calculated their allele frequency in the human population. Intersecting their position on GRCh38 with published MEI list from the 1000G and HGSVC3 we classified them as “new” or “shared” MEIs. We also counted the total number of MEIs per individual by counting their presence on both maternal and paternal assemblies.

To characterize the landscape of MEIs, all identified MEIs were initially categorized into three classes based on their population frequency across all samples: “rare” (frequency < 1%), “polymorphic” (frequency between 1% and 95%), and “fixed” (frequency ≥ 95%). We further quantified the incremental value of expanding genomic sampling, MEI discovery was compared between two datasets: prioritized HPRC1 genomes and the HPRC2 dataset. Individual haplotypes were extracted from the MEI dataset, and their associated MEI population frequencies (derived from all superpopulations) were used for categorization. Haplotypes were then ordered to prioritize those belonging to the HPRC1 genomes at the beginning of the cumulative curve, followed by the remaining haplotypes from the HPRC2 dataset. We then investigated MEI accumulation with an emphasis on population diversity by analyzing the cumulated MEI count across different human superpopulations. For this analysis, haplotypes were primarily sorted by their 1000G assigned superpopulation^33^. Within each superpopulation block, haplotypes were further sorted by their total MEI count. These complementary analyses provide insight into MEI discovery dynamics, addressing both the impact of iterative dataset expansion and the role of population diversity in uncovering human MEI variation. All Python scripts used for these analyses are publicly available at https://github.com/asfistonlavie/HPRC_MEI.

From both phased PacBio and ONT long-read alignment to the haploid assembly, we calculated the methylation percentage on each CpG site. We used pb-CpG-tools (https://github.com/PacificBiosciences/pb-CpG-tools) with the default model for HiFi reads, and `modkit pileup` (https://github.com/nanoporetech/modkit) for ONT reads methylation calculation. We obtained the MEI coordinates on each haplotype assembly by running `rustybam liftover` (https://github.com/vollgerlab/rustybam) from the annotated MEI coordinates on the reference genome. We then extracted the sequences of all MEI from the haplotype assemblies as uBAM format and attached corresponding methylation percentage as methylation auxiliary tags using a custom script (`bam.creation.py` from https://github.com/xzhuo/modbamUtil). We then aligned all MEI sequences with methylation in uBAM format to the consensus of AluY, L1HS, SVA_F, and LTR5_Hs using minimap2^192^. At last, we extracted the methylation percentage of each CpG site from all MEI consensus to plot the MEI methylation profile (`modkit extract`).

#### VNTR growth analysis

Assemblies were mapped to the T2T-CHM13 reference using minimap2^161^ with gap parameters `-z200000,10000`. Tandem repeat (TR) sequences were annotated using vamos v3.0.6^75^ with the TR database v3.0 (efficiency value, q = 0.1) for the T2T-CHM13 genome, available at https://zenodo.org/records/16286509.

We annotated the TR sequences of the HPRC2 samples using vamos v2.2.0^75^ and the vamos v3.0 expanded motif catalog^76^. TR alleles were represented by length as the number of motifs, or by composition as the concatenated string of motif indices. We then searched for loci that showed signatures of population differentiation using 230 individuals, excluding the 2 trio children (HG00733 and NA19240), considering both length and compositional variation. While there is a copy-number (e.g., length) analog to *F_ST_*, *V_ST_* ^193^, this does not extend to multiallelic sequences that vary by composition. To uniformly compare population differentiation by length versus composition, we measured ANOVA according to length variation as well as motif-based edit distance to the major allele for each locus.

#### Inter-chromosomal matches in HPRC vs T2T-CHM13

Inter-chromosomal match quantification. We used WFMASH v0.23.0^194^ (`-p 95 -P inf`) to produce a random subset of all-vs-all pairwise haplotype alignments (25,272 pairs spanning the 466 HPRC haplotypes from 234 samples). T2T-CHM13 was then divided into non-overlapping 100 kb windows. For each window *W* on chromosome *C*, we queried the implicit pangenome graph formed by the pairwise alignments with IMPG v0.4.1 (`-x -m 5 --min-identity 0.98 --min-transitive-len 50000 --min-output-length 50000`) to obtain every haplotype contig reachable from *W* within 5 transitive hops, with each hop ≥ 50 kb and ≥ 0.98 estimated identity. Each contig was assigned to its primary chromosome *P*’; chrUn and chrM were ignored. For each *W* we counted the number of distinct chromosomes (*P*’ ≠ *C*) from which at least one contig was reachable.

#### Asat relatedness with Centrolign

Using `samtools faidx`^195^, we extracted the assembly sequence in the alpha satellites for the samples in the HGSVC and HPRC which passed QC. For each chromosome, we aligned all pairwise combinations of assemblies to each other using Centrolign in direct pairwise mode with default parameters. We used the python script `infer_tree.py` to parse the resulting CIGAR string and apply the formula 1. 0 − ((2. 0 * *matc*ℎ*es*)/(*ref*_*len* + *query*_*len*)) to calculate a distance metric for all pairs of assemblies.

#### Sequence variation at G4 loci, in different genic regions of the genome

Each HPRC haplotype assembly was annotated with G4Discovery^196^ (code available in https://github.com/saswat-km/g4Discovery.PanSN at commit 5620c226297ee98d21e2e20ca3368e928ee94fd4) to obtain G-quadruplex (G4) motifs, and with gfa^197^ (code taken from https://github.com/abcsFrederick/non-B_gfa at commit 2f5a24f08b4ed590855ca1fc871fd41bd5ae9bc4) to obtain all other non-B DNA motifs. G4discovery was run with default settings. gfa was run using the parameter `-skipGQ`. The output tsv files from gfa were converted to bed format for each of the following non-B DNA motif types: A-phased repeats (APR), direct repeats (DR), inverted repeats (IR), mirror repeats (MR), short tandem repeats (STR), and Z-DNA motifs (Z). Triplex motifs (TRI; mirror repeats likely to form triplex DNA) were extracted using awk based on sequence content (100% purine on one strand) and spacer length (maximum 10bp). The software gfa always reports results on the forward strand. All motifs except APR and G4 are symmetrical and exist on both strands. For APR, gfa searches for both A-tracts and T-tracts on the forward strand, meaning motifs on the reverse strand are also found. G4Discovery annotates G4 motifs on both strands. For population-based analysis, only populations with at least 6 haplotypes (3 samples) were used. For counting annotated non-B base pairs, overlapping motifs of the same type were first merged using bedtools merge^198^. On average, 10.4% of the HPRC2 assemblies consist of non-B DNA motifs (ranging from 9.95% to 11.20%). All haplotypes have a total non-B DNA motif content that lies in-between what is found on the previous human reference genome, hg38 (8.89%), and the present T2T reference genome T2T-CHM13 (11.50%, **Supplementary Fig. 26**, **Supplementary Table 9**), likely reflecting inter-individual variation and differing completeness of the assemblies.

The whole genome VCF file derived from PGGB graph, referenced to T2T-CHM13, for the HPRC2 was downloaded from https://garrisonlab.s3.amazonaws.com/hprcv2/vcfs/whole-genome/20250603_hprc8424.CHM13.laced.norm.vcf.gz. Following this, genes were divided into 5’ UTRs, coding sequences (CDS), introns, and 3’ UTRs, using T2T-CHM13 annotations^196^. We also included the promoter and enhancer annotations. Furthermore, the predicted G-quadruplex (G4) annotations for T2T-CHM13, derived using the G4Discovery pipeline, found at https://github.com/makovalab-psu/GreatApeT2T-G4s/tree/main/datasets/pG4s/Homo_sapiens were used to divide each of the functional regions into G4 and non-G4 regions, using `bedtools subtract`. Next, for each of these regions, a VCF was generated using BCFtools^195^ with the `view` subcommand, followed by summary statistics generated with the `stats` subcommand. Using the .vchk file, the SNP (minimum allele frequency ≥ 0.05) frequency per kb was calculated in each of these regions. We also divided the whole T2T-CHM13 genome into G4 and non-G4 regions, and similarly, using their corresponding VCFs, calculated the genome-wide SNP frequency per kb.

### Pantranscriptome

#### CAT2 Annotations

CAT2 (improved version of CAT^89^) was used to annotate genes on all of the HPRC2 assemblies using the minigraph-cactus pangenome alignment (described above) with the GENCODE v48^199^ annotations as the reference gene set. CAT2 uses multiple methods to transfer genes from the reference to each of the targets- transMap^200^ module lifts genes over pairwise chains derived from the pangenome and pairwise chains derived from minimap2^192^ alignments, Liftoff^201^ is integrated to annotate any additional genes missed by the transMap methods and AUGUSTUS^202^ gene prediction fixes gene and CDS boundaries. Furthermore, CAT was given PacBio Kinnex FLNC data to provide extrinsic hints to the Augustus PB (PacBio) module of CAT, which performs ab initio prediction of coding isoforms. CAT then combined these ab initio prediction sets with the various gene projection sets to produce the final gene sets and UCSC assembly hubs.

For the gene annotation sets, we identified the locations of frameshifting indels by iterating over the coding sequence of each transcript and looking for any gaps in the alignment. If the gap had a length that was not a multiple of 3, and its length was <30 bp long (to remove probable introns from consideration), the gap was determined to be a frameshift. We identified nonsense mutations by iterating through each codon in the coding sequence of the predicted transcripts and flagging early stop codons. We then pooled all the transcripts affected by frameshifts and early stop codons together and removed those from the list that passed these criteria (i) near the end of the transcript (based on the 50 bp rule, with modifications as described in Karczewski et al., 2020^87^) and (ii) land in an exon with non-canonical splice sites around it (**Supplementary Table 19**)

#### Ensembl and CAT gene annotation comparisons

In order to analyse the degree of concordance between the two haplotype annotation approaches we paired genes between annotation sets based on genomic overlap and retained instances with reciprocal best hit pairs for downstream transcript and CDS level comparisons with GENCODE v47. Following standardisation of chromosome names, transcript structures within each gene pair were classified into exact exon coordinate matches, same intron chain matches, partial overlap, or no match. Transcripts in genes without a reciprocal counterpart were counted as ‘gene not shared’. Across all biotype categories CAT produced more transcript features than Ensembl. For example, in protein coding genes, CAT annotated approximately 195,000 transcripts per haplotype compared to roughly 170,000 by Ensembl. These differences were shown to predominantly originate from pipeline source data differences rather than algorithmic disagreement, as CAT incorporates Kinnex data and projects GENCODE readthrough genes while Ensembl does not. Excluding these CAT specific categories and restricting to matched gene pairs raised the exact match plus same-intron-chain fraction above 86% for all major biotypes in both pipelines (**Supplementary Fig. 28a**). NA18747_hap2 and NA20762_hap1 are the two haplotypes with lower than expected gene counts.

Divergence of annotated features from their GRCh38 GENCODE v47 counterpart was assessed per matched gene pair. Representative transcript structures were compared to the corresponding reference transcript and each pair was classified as both agree with reference, both diverge from reference, Ensembl specific divergence, CAT specific divergence. The two pipelines agreed on the vast majority of divergence categorisations and pipeline specific divergence was largely biotype dependent (**Supplementary Fig. 28b**). Pipeline-specific divergence was rare and biotype-dependent: Ensembl-specific divergence was slightly elevated in snRNAs and miRNAs. CAT-specific divergence was more frequent for miRNAs and misc_RNAs. This observation is consistent with the greater difficulty projection pipelines often have in transferring short, repetitive features across assemblies.

Coding sequence changes relative to the same GENCODE reference were also analysed in the protein coding subset of the reciprocally paired genes. CDSs were categorised as exact match, in-frame shorter or longer (i.e. length change was divisible by 3), frameshift longer or shorter (i.e. length change not divisible by 3), or as coding lost. In 97.3% Ensembl CDS structure matched the reference exactly while 96.5% did in CAT annotations (**Supplementary Fig. 28c**). It was observed that in frame length changes outnumbered frameshifts and outright coding loss was rare in both pipelines (≤0.5%). Genes for which more than one assembly contained haplotype/reference CDS discordance are enumerated in **Supplementary Table 20**.

#### Gene Copy Number Variation Analyses

Gene family membership was assigned by clustering GENCODEv48 protein-coding genes on the basis of sequence homology and HGNC gene-family designations: protein sequences for the canonical isoform of each gene were aligned all-against-all with DIAMOND (v2.2.0)^203^, and families were formed by clustering at ≥80% sequence identity over ≥80% of length, with HGNC family labels used to reconcile cluster names. For each haplotype, the copy number of a family was the number of CAT2-annotated protein-coding members of that family on that haplotype.

To avoid the reference bias inherent to comparing copy number against a single linear genome, we computed the median copy number of each family across the union of all HPRC2 haploid assemblies plus GRCh38 and T2T-CHM13, stratified by whether the haplotype was X-containing or Y-containing. For each haplotype, a duplication or deletion event was recorded for any family whose copy number deviated from the cohort median; the total number of duplication and deletion events per haplotype, summed across all families, is the quantity plotted in **Fig. 5b**. Haplotypes in **Fig. 5b** are coloured by 1000 Genomes population labels, grouped into 1000G superpopulation^33^ label-based colour families (AFR yellow, AMR red, SAS purple, EAS green, EUR blue) with subpopulation shading.

We ranked gene families by the range of their per-haplotype copy number across the cohort and additionally required them to be multi-exonic in the canonical transcript to exclude calls driven by single-exon paralog clusters such as *TAF* or *USP* gene families as they could be results of assembly artifacts. The top families by this criterion are shown in **Fig. 5c**. For each gene-family locus on each haplotype, local ancestry was assigned using PCLAI, yielding per-locus AFR-like, EUR-like, SAS-like, or AMR/EAS-like genetic ancestry labels. To test whether copy number was stratified by ancestry, we fit a Kruskal-Wallis test of copy number across the four ancestry strata for each family, with effect size quantified by epsilon-squared (ε²). P-values were Bonferroni-corrected across families tested. 125 families showed significant separation in copy number by PCLAI ancestry label, but no single ancestry predominated as the high-copy ancestry across this set (AFR-like ∼30%, AMR/EAS-like ∼30%, EUR-like ∼20%, SAS-like ∼15%). Of these, only *TBC1D3* and *CCL* combined statistical significance with large effect sizes; the remainder reached significance with modest ε² values, indicating that ancestry explains only a minor fraction of their copy-number variance. This same analysis was repeated with the Ensembl annotations (**Supplementary Fig. 29**).

#### TBC1D3 Locus Analysis

Of the 462 HPRC2 haploid assemblies, we restricted the *TBC1D3* analysis to haplotypes in which both Cluster 1 and Cluster 2 were assembled with high confidence. Haplotypes were excluded if Flagger^50^ or NucFlag^57,70^ annotations flagged any misassembly within either cluster, or if the two clusters were not assembled on the same contig. This filtering yielded the 437 QC-passing haplotypes used in the cluster-specific analyses.

*TBC1D3* paralogs were extracted from the CAT2 annotation of each haplotype assembly. All annotated *TBC1D3* family members were extracted and classified by CAT2 biotype as protein-coding copies or pseudogene copies. The two principal expansion blocks on chromosome 17q12 (Cluster 1 and Cluster 2) were defined by their syntenic boundaries with GRCh38 and T2T-CHM13; copies falling outside these blocks (the dispersed copies on both arms of chromosome 17) were excluded from this analysis.

For each haplotype, the ordered sequence of *TBC1D3* protein-coding copies, pseudogene copies, and intervening segmental duplication blocks within each cluster was extracted from the CAT2 annotation coordinates. Segmental duplication content was annotated with DupMasker^97^ (available through RepeatMasker), producing the coloured arrow tracks shown. Two cluster configurations were considered identical if they shared the same ordered series of paralog identities and orientations and the same DupMasker block composition; this yielded 89 unique Cluster 1 configurations and 130 unique Cluster 2 configurations across the QC-passing haplotypes. For display in **Fig. 5d**, we selected the shortest confirmed configuration, the longest confirmed configuration, and the three most frequently observed configurations within each cluster, alongside GRCh38 and T2T-CHM13.

The total number of protein-coding *TBC1D3* copies per haplotype within each cluster was tested for ancestry stratification using the same Kruskal-Wallis procedure described above, with PCLAI local-ancestry labels assigned per cluster. Pairwise comparisons between genetic ancestry labels (shown by significance bars in the **Fig. 5d** box plots) were performed with two-sided Mann-Whitney U tests and Bonferroni-corrected for the six pairwise comparisons within each cluster. Within both clusters, those whose local ancestry, as assigned by PLCAI, are AFR-like carried significantly more protein-coding *TBC1D3* copies than those that are EUR-like, SAS-like, and AMR/EAS-like (Cluster 1: *p = 2.0 × 10^-20^, ε^2^= 0.21*; Cluster 2: *p = 9.2 × 10^-18^, ε^2^ = 0.19*). The reported statistics reflect the Kruskal-Wallis tests across all four ancestries.

#### Gene-level quantifications from Kinnex Data (N=206 Samples)

Full-length non-chimeric (FLNC) reads produced by the PacBio Iso-Seq pipeline^204^ were clustered into representative consensus sequences using the isoseq cluster tool, retaining only clusters supported by two or more FLNC reads (singletons were excluded). Representative transcript cluster sequences were then processed with IsoQuant^205,206^ against the GRCh38 reference genome and the GENCODE v44 comprehensive annotation, which assigned each cluster to an annotated gene identifier. Gene-level quantification was obtained by summing the FLNC read counts of all clusters assigned to a given gene. To enable cross-sample comparisons, gene-level counts per sample were normalized to counts per million (CPM). For each sample, we calculated the number of expressed genes (CPM ≥ 1) across three gene categories: protein-coding genes, long non-coding RNAs (lncRNAs), and all other non-coding feature types combined (**Supplementary Table 21**).

#### Detection and annotation of Reference Divergent Transcripts

For each sample, *de novo* transcript cluster sequences generated by the Iso-Seq pipeline were aligned to both the GRCh38 reference genome and the two matched personal haplotype assemblies using `minimap2` with ‘splice’ preset parameters. Reference-divergent transcripts (RDTs) were identified by comparing alignment characteristics across these targets. A transcript cluster was classified as reference-divergent if it exhibited an alignment score (ms tag) at least 50 points or 5% higher against a personal haplotype assembly than against GRCh38, and additionally aligned to at least one personal haplotype with greater than 99% sequence identity and query coverage. This criteria ensures that candidate reference divergent reads align robustly to at least one of the personal haplotype assemblies while aligning poorly to the GRCh38 reference.

Transcript clusters passing these filters were re-aligned to the matched personal haplotype assemblies using pbmm2, and transcript models were inferred using the `isoseq collapse` pipeline with sequence identity and query coverage thresholds of 99%. Collapsed transcript models were annotated by overlap with gene features from each haplotype’s LiftOff-based RefSeq annotation. Genes harboring at least one RDT were designated RDT-genes. To reduce false positives arising from loci with minimal reference-divergent signal, RDT-genes were further required to have a minimum of 10 FLNC reads supporting a reference-divergent transcript, with those reads comprising at least 10% of all FLNC reads assigned to that gene. Summary statistics for each RDT and RDT-gene, including per-locus RDT counts, haplotype incidence, and annotation labels, are provided in **Supplementary Table 22**.

#### TE-associated transcripts

FLAIR2 (v2.2.0)^207^ was used to detect transcripts from long-read RNA sequencing data of 202 LCLs. RNA reads from each individual were mapped to their corresponding personalized genomes, with each haplotype assembly processed separately, using the flair align module with minimap2. Aligned reads were corrected and high-confidence isoforms were then defined from corrected reads using the flair collapse module with recommended human parameters (“--stringent --check_splice --generate_map --annotation_reliant generate”). SQANTI3 (v5.5.1)^208^ was used to classify transcripts and perform quality control. Potential artifacts were removed using the default SQANTI3 filtering rules. TE-associated isoforms were identified by intersecting TE coordinates with isoform annotations using bedtools (v2.26.0)^198^.

### Panepigenome

#### CpG Island annotation

CpG islands were annotated for each individual haplotype assembly using the UCSC cpg_lh algorithm, which identifies maximally scoring segments based on a weighting of +17 for CG dinucleotides and −1 for all others. Candidate regions were filtered according to the standard criteria established by Gardiner-Garden and Frommer: GC content ≥ 50%, sequence length > 200 bp, and an observed-to-expected CpG ratio > 0.6 ^209^. Given the near-T2T-level contiguity of these assemblies, we performed this identification on the full sequence, including previously masked regions. Furthermore, we defined CpG shores and shelves as the 2-kb flanking regions immediately adjacent to the island boundaries and the subsequent 2-kb intervals, respectively.

#### Dinucleotide growth analysis

CpG were found and indexed in the Minigraph-Cactus derived HPRC2 graph (v2.1) using `panmethyl --index --motif CG`^102^. The same was repeated for every possible dinucleotide to count their abundance in the graph relative to T2T-CHM13 by changing `–motif` to AA, AC, AG, AT, etc. Then, graph CpG coordinates were lifted to each linear assembly using `panmethyl --lift` to obtain linear coordinates for each CpG in each assembly. CpGs overlapping regions that do not pass flagger validation were removed. Then the frequency of each CpG was counted with `collapse_flagged_cpgs.py` to obtain a frequency histogram. We ran `panacus growth -l 0,47,417` on the resulting histogram to obtain the CpG growth curve stratified by CpG frequency.

In order to contextualize CpG growth relative to other dinucleotides, for each of the 16 dinucleotides, percent growth of the pangenome over the reference (CHM13) was computed as (Pangenome - Reference) / Reference x 100, normalizing each dinucleotide to its own reference abundance, along with a pangenome-count-weighted mean across dinucleotides. An ordinary least-squares regression of percent growth on reference count was fit across the 15 non-CpG dinucleotides. Analyses were performed in R.

#### CpG growth context

The linear coordinates of graph CpGs were projected onto each haplotype assembly and intersected with assembly-specific annotations using BEDTools. Annotation sets included RepeatMasker elements, CAT gene annotations, segmental duplications, promoters, and CpG islands, shores, and shelves. Overlap patterns were summarized using ComplexUpset. Because rare segmental duplications annotate a large fraction of pangenome sequence, only segmental duplications present at a frequency greater than 1% were included in the CpG annotation analysis. To further attribute CpGs absent from the T2T-CHM13 reference to their underlying source, we intersected non-reference CpGs with PAV variant annotations generated relative to T2T-CHM13. CpGs were classified as arising from insertions, SNPs, or inversions when they could be uniquely assigned to one of these variant classes; CpGs that overlapped multiple, complex, or unresolved variant calls were retained as unassigned. Annotation-specific expansion relative to T2T-CHM13 was calculated as:

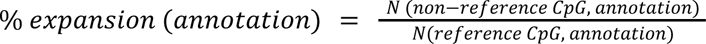

Because annotation categories were not mutually exclusive, individual CpGs could contribute to more than one annotation class.

#### Base modification calling

HiFi methylation calls for HPRC2 were harmonized with Primrose v1.4.0 by re-calling BAMs lacking methylation calls, BAMs with indeterminate methylation provenance, and BAMs previously processed with Jasmine (machine default), and by regenerating HiFi BAMs from subreads with CCS when kinetics tags were absent before Primrose methylation calling.

#### Methylation Likelihood QC

To ensure high-fidelity methylation profiling, we performed rigorous quality control (QC) on 1,158 BAM files across 232 samples. The QC pipeline comprised three stages: outlier detection, batch effect assessment, and genomic coverage verification.

First, we identified technical outliers by evaluating median and distributional metrics of ML values within raw reads. Subsequent to alignment against the T2T-CHM13 v2.0 reference genome (GCF_009914755.1_T2T-CHM13v2.0_genomic.fa) using pbmm2 (v1.14.99; `--preset HIFI --unmapped`), we retained BAM files meeting stringent thresholds: >80% of reads with mapping quality (MAPQ) >30, >90% of reads with read quality >0.99, alignment identity >99%, mean read length >10 kb, and a mapping rate >0.99. We further filtered files by assessing median methylation values and distribution profiles to mitigate technical bias, resulting in 1,111 passing BAM files.

Next, we assessed potential batch effects in genome-wide methylation levels associated with the sequencing platform (Sequel II vs. Revio), time points, and SMRTcell positions using technical replicates. A total of Five files from HG02080 (SMRTcell-1784.m64467e_220908_142429.pbccs-6.3.0.hifi_reads.primrose-1.2.0.5mC.bam and SMRTcell-1785.m64467e_220910_093757.pbccs-6.3.0.hifi_reads.primrose-1.2.0.5mC.bam) and HG002 (m84005_220827_014912_s1.hifi_reads.bam, m84005_220919_232112_s2.hifi_reads.bam, and m84011_220902_175841_s1.hifi_reads.bam) that exhibited significant laboratory-specific effects (*P* < 0.05) were excluded. Finally, we performed a genomic coverage assessment on the merged alignment files for the remaining 226 samples. Samples were retained only if genomic coverage exceeded 30X; one sample (HG00272) was excluded due to a ∼50 Mb inversion misassembly, yielding a final dataset of 1,092 BAM files across 220 samples (440 haplotypes).

Nanopore Methylation QC

Methylation quality control was performed on 929 nanopore BAM files spanning 232 samples. The workflow comprised two steps: (1) an alignment-free assessment of modification likelihoods (ML) from the unaligned BAMs, and (2) a CpG methylation rate (% mCpG) computed after minimap2 (v2.30-r1287) alignment to T2T-CHM13 (https://github.com/marbl/CHM13), using modkit pileup (v0.5.0) for mCpG aggregation. BAM files were flagged as outliers if a metric fell more than 1.5×IQR beyond the first or third quartile. This flagged 37 files for ML values and 57 for % mCpG. We then aggregated to the sample level to identify samples whose abnormalities were consistent across sequencing runs rather than confined to a single BAM. Three samples, NA20762, NA19338, and HG02583, showed consistently abnormal methylation profiles across the majority or all of their nanopore BAM files.

#### Phasing and alignment

PacBio HiFi long reads were aligned to each sample’s haplotype assembly using minimap2. The read alignments for each haplotype were compared against each other to determine which haplotype to assign the read. The first parameter considered was the mapping score and the second was the minimap2 alignment score. If both are equal, the reads were assigned by alternatively switching between haplotypes, yielding 440 haplotype-resolved CRAMs at a mean coverage of 25× (range 13–35×). Per-haplotype alignment quality was assessed from ‘samtools stats’ (sections SN, COV, MAPQ, ID, GCD, and RL) across all 440 haplotypes; indel profiles, mapping-quality distributions, and GC bias were tabulated cohort-wide to identify donors with deviant alignment behaviour. Scripts used for these analyses are available at https://github.com/twlab/HPRC2_DNA_Methylation.

#### CpG methylation calling

Aligned BAM files were sorted and indexed using samtools (v1.21). CpG methylation (5mC) levels were extracted at single-base resolution using pb-CpG-tools (v2.3.2) with the `--pileup-mode model` model which applies a machine-learning approach to estimate methylation for ML tags. Whereas panmethyl uses a “pile-up” approach and averaging to estimate methylation. In order to eliminate purely algorithmic differences in methylation estimation, pb-cpg-tools was re-run with `--pileup-mode pileup` for comparisons with panmethyl.

#### Comparing graph-based methylation calls to assembly based calls

Methylation data was remapped to the graph using `panmethyl --aligner GraphAligner --graph hprc-v2.1-mc-chm13-eval.P.gfa --bams phased_bams.csv --motif CG --code C+m` to obtain graph-based methylation estimates for every CpG. The values were joined to the assembly-based methylation values (pb-cpg-tools) using GNU join and BEDtools for comparison by leveraging the graph CpG coordinates lifted to the linear assemblies. To estimate concordance, we calculated the R^2^ between the assembly-based and graph-based methylation levels. We counted the number of off-reference CpGs methylation levels yielded by the graph for every methylome.

#### Non-reference transposable element methylation analysis

Both PacBio HiFi and ONT haplotype- phased long-reads were aligned to their respective haploid assemblies (as described above), from which we calculated the methylation percentage on each CpG site. We used pb-CpG-tools (https://github.com/PacificBiosciences/pb-CpG-tools) with the default model for HiFi reads, and modkit pileup (https://github.com/nanoporetech/modkit) for ONT reads methylation calculation. We obtained the MEI coordinates on each haplotype assembly by running rustybam liftover (https://github.com/vollgerlab/rustybam) from the annotated MEI coordinates on the reference genome. We then extracted the sequences of all MEI from the haplotype assemblies as uBAM format and attached corresponding methylation percentage as methylation auxiliary tags using a custom script (bam.creation.py from https://github.com/xzhuo/modbamUtil). We then aligned all MEI sequences with methylation in uBAM format to the consensus of AluY, L1HS, SVA_F, and LTR5_Hs using minimap2, accordingly (minimap2). At last, we extracted the methylation percentage of each CpG site from all MEI consensus to plot the MEI methylation profile (modkit extract).

#### Promoter mQTL mapping and lead variant identification

We performed variant-sequence-aware mQTL analysis using methylation profiles from 460 HPRC assemblies. To define pangenome-aware methylation phenotypes, the T2T-CHM13 reference genome was segmented into 200-bp windows and projected to the assemblies. Association testing was performed using a framework that integrates allelic imbalance and total dosage effects. Small variants were obtained from the MC v2 pangenome decomposed biallelic variant set normalized by the PanGenie pipeline and filtered at MAF > 0.01, whereas structural variants were obtained from the HPRC structural variant callset and filtered at MAF > 0.005. In the total dosage effect analysis, covariates included 30 PEER factors, 10 genetic PCs, coverage, N50, and ONT chemistry. For each region, the adjusted minimum *P* value was estimated using a β-distribution approximation derived from 1,000 permutations. Storey’s *q*-value procedure was then applied to the β-approximated *P* values across all tested regions, and significant mQTL regions were defined using a *q*-value threshold of 5%. For each promoter mQTL region, the lead variant was defined as the tested *cis* variant with the lowest nominal association *P* value within the corresponding *cis* window. Scripts used for these analyses are available at https://github.com/twlab/HPRC2_DNA_Methylation.

#### mQTL enrichment in CenSat regions

Promoter mQTLs were analyzed in T2T-CHM13 coordinates to determine whether associations with different classes of lead variants were recovered in structurally complex regions of the genome. Promoter mQTL intervals were stratified by lead variant class and compared with the T2T-CHM13 CenSat v2.1 annotation. Lead variants were grouped as single-nucleotide variants, small insertions or deletions, small complex variants, or structural variants.

For each lead variant class, the observed number of promoter mQTL intervals overlapping CenSat annotations was compared with a null distribution generated by randomly repositioning promoter mQTL intervals within the same chromosome while preserving interval width and lead variant class. This permutation was repeated 5,000 times per class. Fold enrichment was calculated as the observed overlap divided by the mean overlap across permutations. Empirical p-values were calculated as the fraction of permuted sets with overlap counts equal to or greater than the observed count, with a +1 correction, and were adjusted across lead variant classes using the Benjamini-Hochberg procedure. Scripts used for these analyses are available at https://github.com/twlab/HPRC2_DNA_Methylation.

#### Promoter CGI harmonization and PCLAI integration

All coordinates refer to ‘chm13v2.0_maskedY_rCRS.fà. CpG-island annotations were derived from the UCSC ‘cpgIslandExt’ track (30,617 islands) lifted onto T2T-CHM13 coordinates. Local ancestry via PCLAI (see **Point Cloud Local Ancestry Inference Methods**) was used to assign annotations in T2T-CHM13 coordinates to each promoter CGI.

#### mQTL lead variant allele-frequency analysis by local ancestry

Promoter mQTL lead variant positions were matched to the HPRC v2.0 minigraph-cactus T2T-CHM13-based wave VCF (hprc-v2.0-mc-chm13.wave.vcf.gz) by exact chromosome and position. Lead variants without exact VCF entries were excluded (retaining 72,700 of 80,854 lead variant rows, 62,716 of 69,966 unique positions). Phased VCF genotypes were parsed into per-haplotype allele calls, excluding the GRCh38 pseudo-sample. Per-haplotype local ancestry was assigned to each lead variant position by intersection against the HPRC2 T2T-CHM13 PCLAI release v1.0.1 with bedtools intersect (v2.30.0); HG06807 was excluded due to absence of T2T-CHM13 PCLAI annotations. Lead variants without PCLAI annotation at the lead variant position were excluded, retaining 67,562 unique lead variants.

For each retained lead variant and each PCLAI centroid, ALT allele counts were summed across called haplotypes assigned to that centroid and divided by the number of such haplotypes to give a per-cluster alternate allele frequency. Differentiation analyses were restricted to the 60,670 biallelic lead variants (single ALT allele in the matched VCF record). Per-locus population differentiation across PCLAI centroids was tested with the Genepop genic differentiation test (*Genepop* R package, struc function; dememorization = 1000, batches = 100, iterations = 5000). P-values were corrected across loci by the Benjamini–Hochberg procedure. For each biallelic lead variant retained in the per-cluster allele-frequency analysis, the number of PCLAI centroids in which the alternate allele was observed (ALT allele frequency > 0 across called haplotypes assigned to that centroid) was tabulated. Lead variants were then categorized by cluster-occupancy count (observed in one, two, three, or four PCLAI centroids). To describe variation in allele frequencies across lead variants, an alternate allele frequency matrix was constructed with biallelic lead variants as rows and the four PCLAI centroids as columns; lead variants with invariant AF across all four centroids were excluded. Principal components analysis was performed on the centered, unscaled AF matrix using R prcomp (center = TRUE, scale = FALSE).

#### PM20D1 locus analysis

PM20D1 was selected as a representative locus for the PCLAI-stratified allele-frequency and methylation analyses described above. Haplotype-resolved PM20D1 promoter methylation values were stratified by PCLAI centroid (assigned at the lead variant position as described in “mQTL lead variant allele-frequency analysis by local ancestry”) and by lead variant (rs9438393) genotype. Haplotypes were classified into hypomethylated, partially methylated, and hypermethylated states. Differences in methylation-state proportions across PCLAI centroids were tested with a likelihood-ratio G statistic computed from the methylation-state by PCLAI-centroid contingency table; significance was assessed by a sample-blocked randomization test in which PCLAI cluster labels were permuted across samples while preserving the pairing between the two haplotypes of each individual. The locus was visualized in the HPRC Panepigenome Browser using haplotype-resolved methylation, local ancestry, and genome-alignment tracks. Scripts used for these analyses are available at https://github.com/twlab/HPRC2_DNA_Methylation.

#### WashU HPRC epigenome browser

The HPRC Epigenome Browser (HPRCEB; https://epigenome.humanpangenome.org/) is a research portal centered on HPRC2 data, integrating both genetic variation and functional genomics datasets. The browser enables users to select from all currently available Release 2 samples and assay types for streamlined visualization. Individual haplotypes are displayed using genome alignment tracks^210^. At finer resolution, the browser reveals small variants, including SNP and INDEL, whereas at larger scale it displays structural variants together with RepeatMasker annotations in haplotype-specific coordinates. ISO-seq data are aligned and processed against standard reference assemblies (hg38 and T2T-CHM13), and Hi-C (i.e. Omni-C) data are processed against T2T-CHM13. In contrast, other epigenomic assays are presented in haplotype-specific coordinates to facilitate visualization of epigenomic features in the context of structural variation^103^. The portal also provides assembly quality-control metrics and additional assembly-level annotations for convenient access, thereby transforming valuable HPRC2 resources into an accessible visualization platform that can be explored in only a few clicks.

#### Fiber-seq pangenome analysis methods

In order to perform comparative analysis of chromatin actuation (Fiber-seq) across HPRC samples, we developed a pangenome graph-based approach to evaluate chromatin actuation measured in native assembly space for a set of ‘consensus peak regions’, defined as genomic regions at which at least one sample has significant chromatin actuation^211^. First, Fiber-seq reads were mapped to donor specific assemblies, and FIRE was run on each sample individually to identify a set of sample-specific FIRE peaks. To enable comparative analysis of this data, we developed a computational approach which uses a graph structure to define consensus peak regions based on these sample-specific peaks, and then evaluates actuation at assembly-specific coordinates associated with each consensus peak region. We first used minigraph-cactus to construct a pangenome graph containing 80 assemblies: 76 HPRC haplotypes (38 individuals from which we had Fiber-seq data), as well as the HG002, T2T-CHM13, and GRCh38. The --filter 0 parameter was used to prevent filtering of rare variants. Consensus peak regions were defined via an iterative approach during which all assembly-defined peaks are moved through that graph onto a common assembly (e.g. T2T-CHM13), and consensus peak regions are called at locations with at least one peak. All peaks that were not transferred to this assembly or any previously used assemblies due to an absence of a homologous sequence are then transferred to a new assembly, and consensus regions are identified in this assembly space. Once all peaks have been successfully transferred, the set of consensus peak regions are then transferred back through the graph onto each assembly. These assembly-specific coordinates are then used to pull chromatin actuation values from the original sample-specific FIRE data. We thus obtain a chromatin actuation score for all samples at each consensus peak region, enabling comparative analysis. The number of assembly-unique regions is an overestimate due to imperfect alignment at complex regions in the graph. This approach builds upon functions from the vg toolkit to interact with the pangenome graph.

### Pangenome Applications

#### SV Counting

We used the HPRC v2.0 pangenome graph constructed with Minigraph-Cactus based on T2T-CHM13 for genotyping. We ran our previously developed traversal-based decomposition

pipeline (https://github.com/eblerjana/genotyping-pipelines/tree/main/prepare-vcf-MC)^13,38,212^ to add annotations that encode nested variants to the Minigraph-Cactus output VCF containing top-level graph bubbles. We provided the resulting VCF to PanGenie (v4.2.1), as well as Illumina reads for all 3,202 1000G samples and 6 additional HPRC2 samples. PanGenie v4 implements a new preprocessing step which reduces the full set of input haplotypes to a smaller set to speed up the subsequent genotyping step. We filtered genotypes based on a previously developed regression model^13–15^ (Supplementary Material), retaining 39,366,061 of 46,996,181 SNPs + indels (< 50 bp, 83.8 %) and 832,728 of 901,029 SV alleles (≥ 50 bp, 92%). We compared our filtered genotypes for all 3,202 1000G samples to PanGenie genotypes previously generated for the same samples by the HGSVC3 based on a pangenome with 231 haplotypes^15^, as well as an Illumina-based SV callset for the 1000G cohort generated with traditional short-read based callers ^38^. The latter calls are based on GRCh38 which makes a direct comparison between all three sets difficult. We therefore compared the callsets based on the number of SVs called in each sample, focussing on rare variants with allele frequencies below 1%.

#### QV Estimation

We furthermore phased our 1000G genotypes with SHAPEIT5^213^ (v5.1.1) and constructed consensus haplotype sequences for all 3,202 1000G haplotypes by implanting phased variants into the T2T-CHM13 reference genome. We evaluated the accuracy of 16 haplotypes based on ground truth de novo assemblies from the HGSVC3^15^. We computed k-mer based QVs with Merqury^163^. We additionally applied our previously introduced QV estimation pipeline which computes local QVs within 1 Mbp windows along the consensus haplotypes based on variants called from alignments of the consensus haplotypes to ground truth assemblies. For sample HG00096, we intersected our QV windows with BISER (v4.1) ^214^ and RepeatMasker (http://www.repeatmasker.org/) annotations that we computed for the respective consensus haplotypes.

#### Challenging Medically-Relevant Genes Analysis

To calculate haplotype availability we extracted local haplotypes around 273 challenging medically relevant genes^110^, 35 HLA and 10 KIR genes, expanding locus boundaries by at most 100 kb to contain pangenomic variation appearing at the gene boundary. We then constructed global pairwise haplotype alignments and calculated phred-scaled sequence divergence (QV) using Locityper v1.4.5^28^. To obtain haplotype availability we identified, for each out-of-sample haplotype, the largest QV against any assembled haplotype from the HPRC1 and HPRC2 panels.

#### GRef Off Reference Genotyping

We obtained 30x Novaseq reads for nine GIAB samples from gs://deepvariant/benchmarking/fastq/wgs_pcr_free/30x/ and mapped them to the T2T-CHM13-based v2.1 MC graph using vg giraffe with haplotype sampling. We computed positional coverage with vg pack -Q 5 and then genotyped with vg call, using the augmented pangenome reference constructed as described previously. Only sites considered “PASS” by vg call were considered. The containing reference interval on T2T-CHM13 of each GRef variant was intersected with the HPRC annotations for genes, repeats, satellites and segmental duplications. Intervals that were more than 50% covered were labeled accordingly, and if multiple labels overlapped the one with the highest coverage was chosen. The same read mappings that were genotyped with vg call were projected to a GRef-based BAM using vg surject, restricting to contigs ≥1500bp, then called using FreeBayes v1.3.10 (Garrison et al.). The FreeBayes calls with quality below 20 were filtered out, and were compared to the vg call output with Aardvark (Holt et al, 2025).

#### RCCX Analysis

A new RCCX local pangenome was built using Minigraph-Cactus^59^ and the 464 sequences (GRCh38 as backbone, T2T-CHM13, and 462 haplotypes from HPRC2) in the region. We used the “collapse” mode of Minigraph-Cactus (“--collapse” argument) that allows self-alignments to the backbone too to create the cyclic collapsed pangenome used by Parakit^112^. The paths of each haplotype in the pangenome were then split at the RCCX region’s breakpoints to extract subpath across the RCCX module. These module subpaths were assigned to the pseudogene module or gene module based on the node similarity with the annotated reference path (here GRCh38). Of note, another approach using principal component analysis identified the same two main clusters of module subpaths. The subpath assignments were then used to color informative nodes in the pangenome: nodes specific to the pseudogene module, and nodes specific to the gene module.

To compare the added value of the HPRC dataset, we counted how many new informative nodes were included in the pangenome when integrating haplotypes from release 1 and release 2. More informative nodes result in a higher resolution to detect fusions and gene conversions.

We then used Parakit to characterize the RCCX region of each haplotype in release 2. For each haplotype, Parakit identified fusions and gene conversions from their path through the pangenome. The haplotype traversal through the informative nodes was also visualized on top of gene annotations (see Figure 7e).

#### D4Z4 array annotation and classification

The HPRC2 genome assemblies were annotated with KaryoScope^116^, an alignment-free annotation tool that classifies genomic sequences by querying k-mers against a pre-built database derived from T2T-CHM13. Each k-mer was assigned corresponding labels for chromosome identity, satellite composition, repeat classification, subtelomeric context, and gene structure. For the D4Z4 analysis, we examined the 4q subtelomere of all 462 haplotypes (231 diploid assemblies). After retaining only chromosome 4 contigs assembled telomere-to-telomere, 249 haplotypes remained. For each haplotype, we extracted a 500-kb window spanning the subtelomeric q-arm and retained four annotation categories: satellite features (β-satellite, segmental duplication-enriched sequence, and novel sequence), gene features (exonic, intronic, and intergenic), repeat features (LINE, SINE, LTR, DNA transposons, and RNA elements), and subtelomeric features (TAR1, interstitial telomeric sequence, and canonical and non-canonical telomere repeats). Hierarchy-aware smoothing was not applied; all analyses used the unsmoothed KaryoScope output^215^.

D4Z4 repeat arrays were identified with a multi-step pipeline. β-satellite fragments were extracted from the satellite annotations and merged across gaps of up to 400 bp to reconstruct individual β-satellite repeat (BSR) clusters, reflecting the canonical structure in which each ∼3.3-kb D4Z4 unit contains a single ∼2.3-kb BSR^216^. Each cluster was annotated for gene structure by computing the fraction of its span covered by exonic, intronic, and intergenic features. Clusters ≥1,500 bp with ≥50% exon coverage were designated exon-dense and treated as candidate array elements.

Arrays were defined by grouping consecutive exon-dense clusters, with gaps >10 kb marking boundaries between separate arrays. Adjacent arrays separated by a small gap (≤4 kb) or by predominantly novel sequence (>70%) were merged, unless the intervening region contained >30% intergenic content. Array boundaries were then extended to incorporate exon-dense fragments within 1,500 bp of an array edge, with trailing extension halting at intron-rich clusters indicative of terminal features.

Two terminal features were assessed distal to each array. A terminal β-satellite – a structural marker of the polymorphic 4qA allele^83^ – was defined as a BSR cluster ≥2,500 bp with <35% exon content within 3 kb of the last exon-dense element. To confirm pLAM status at the sequence level, we used BLAST to search each haplotype’s BSR region for the canonical pLAM sequence^217^ classifying each hit as functional (ATTAAA polyadenylation signal) or non-functional (ATCAAA variant). pLAM coordinates were then intersected with the KaryoScope-derived array annotations, retaining only hits within an array’s BSR region. A segmental duplication-enriched terminal, consistent with the 4qB allele, was defined as a contiguous region ≥1,000 bp with ≥50% segmental duplication-enriched density within 5 kb of the array end.

Arrays were classified as canonical with a β-satellite terminal (C+; 4qA) if a terminal β-satellite was present and the array contained ≥1 element; canonical without a β-satellite terminal (C−; 4qB) if only a segmental duplication-enriched terminal was present and the array contained ≥3 elements; or degraded (D) if no terminal was detected and the array contained ≥3 elements. C+ arrays were further annotated with BLAST-confirmed pLAM status (functional or non-functional) where available. Arrays with fewer than three elements and no terminal were classified as fragments and excluded, and haplotypes whose first array began within 1 kb of the analysis window edge were flagged as potentially truncated.

The proximal boundary of the main D4Z4 array was defined from interspersed repeat annotations (LINE, SINE, and LTR). The last interspersed element within 50 kb upstream of the first BSR cluster marked the start of the contiguous array, separating it from proximal degenerate sequences.

Per-haplotype summary statistics comprised array count, classification (C+, C−, or D), element count per array, median and mean element size, and total β-satellite coverage. Haplotypes were further annotated for clinically relevant configurations: FSHD1-like (≤10 elements with a functional pLAM, not truncated), intermediate-length (11–20 elements with a functional pLAM), multi-array, and degraded.

#### Quantification of eQTL association improvement using pangenome variant calls

We constructed a merged variant callset representing all unique variants of 430 samples from GRCh38 based and pangenome based callsets, including 1000G calls^38^, PanGenie^29^, EdgeDepth^124^, and danbing-tk^123^. Because the same variants could be reported redundantly across callsets, we identified and removed variants that were already represented in another callset. Redundant variants across callsets were assessed in two steps: variant match and genotype correlation. First, variants matched were either determined by RTG vcfeval (v3.13)^218^ as true-positive matches or by identical chromosome, position, reference allele and alternative allele after normalization with bcftools norm (left-alignment and normalization) (v1.21)^195^. When a variant matched multiple variants in another callset, the pair with the highest squared Pearson correlation coefficient (R^2^) was selected as the matched variant pair. Second, for each matched variant pair, we computed the R^2^ of genotypes (for 1000G and PanGenie) or dosages (for EdgeDepth) across the 430 samples. A variant pair was classified as redundant if it satisfied the matching criterion and had R^2^ ≥ 0.5. Matched pairs with R^2^ < 0.5 were retained as distinct variants, as the low correlation indicated they capture different information despite positional match. To assess the added value of augmenting the 1000G callset with pangenome-specific variants, we retained a single representation according to the priority order of 1000G, PanGenie, and EdgeDepth. Variants already represented in a higher-priority callset were removed from lower-priority callsets. The final merged callset comprises 1000G calls supplemented by pangenome-specific variants, including 19.17 million variants from 1000G and 9.22 million variants from pangenome-based callsets (2.18 million from PanGenie, 7.02 million from EdgeDepth and 0.02 million from danbing-tk).

We then performed eQTL analysis using the merged callset in 430 samples using tensorQTL (v1.0.8)^219^, identifying 10,169 eGenes at q value < 0.05. To quantify the improvement by the merged callset, we compared these results with an eQTL analysis performed using the same samples and 1000G calls alone. For each eGene, we identified the lead variant in each callset as the variant with the smallest nominal p-value. Because the two analyses included different numbers of variants, resulting in different multiple-testing burdens and different opportunities to identify stronger associations by chance, we compared lead marker association strength using the multiple testing corrected tensorQTL beta-approximate permutation p value. For each eGene, improvement was defined as the percent increase in -log10(beta-approximate p value) for the lead marker in the merged callset relative to the lead marker in 1000G calls alone. We then computed the proportion of eGenes exceeding each improvement threshold shown in the panel.

