## Supplementary Figures for "HPRC2: A human pangenome reference with near-complete coverage of common genetic variation"

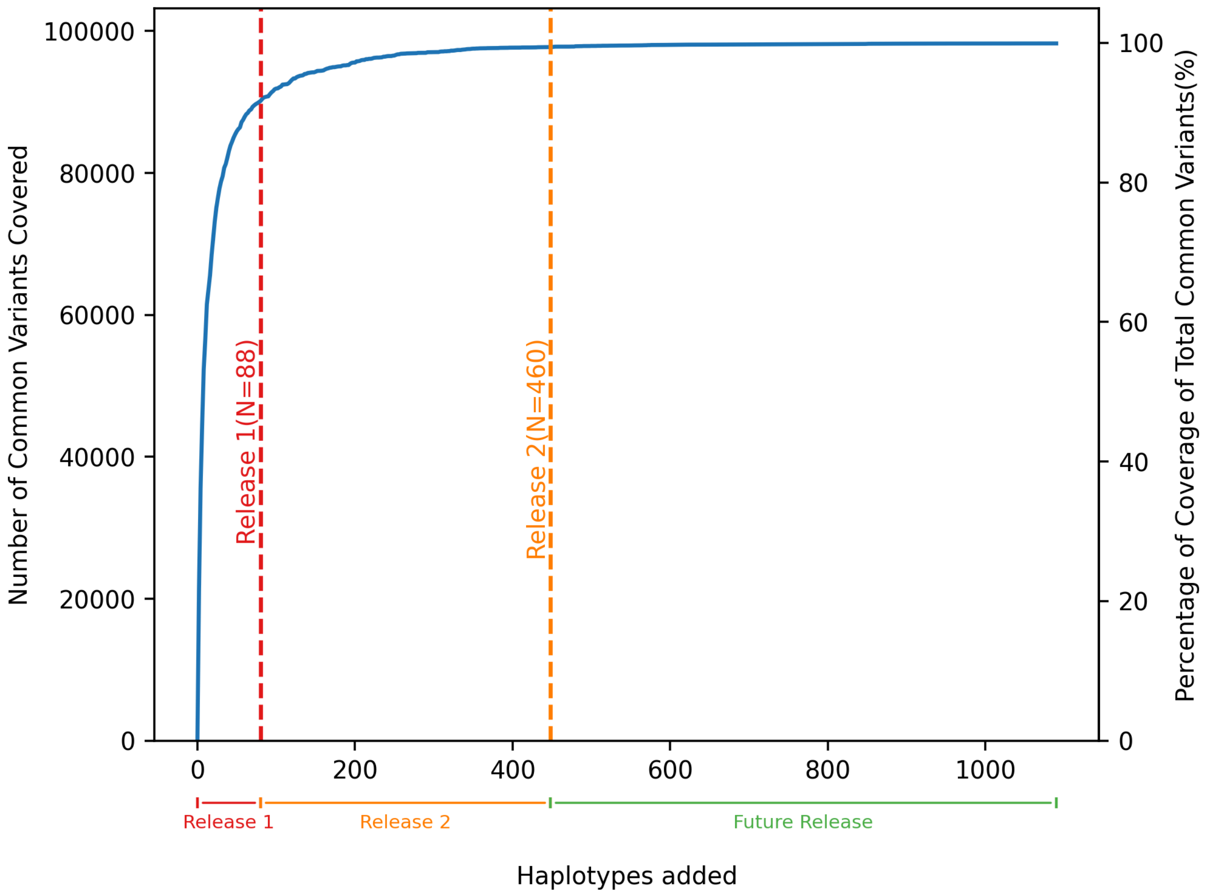


**Supplementary Fig. 1 max_var_coverage_w_hprc1.** MaxVar coverage of common variants on chromosome 22 in the All of Us Research Program cohort (N≈440,000) including HPRC1. The x-axis represents the cumulative number of haplotypes added to the reference panel, and the dual y-axes indicate the absolute number (left) and percentage (right) of common variants coverage. Vertical dashed lines mark HPRC Release 1 (N=88 haplotypes) and Release 2 (N=460 haplotypes). Unlike Figure 1b, this analysis includes Release 1 haplotypes, which were not selected by MaxVar, to capture the full variant coverage trajectory. Coverage to the right of the orange dashed line reflects projected estimates generated by continuing to run MaxVar on the All of Us samples beyond the HPRC2 collection of samples. Because including HPRC1 required rerunning the full experiment with all available common variants, this analysis was performed on chromosome 22 as a representative chromosome. The extended trajectory illustrates the steep initial gains in coverage of variants from the earliest haplotypes added, the transition into a phase of diminishing return beyond ~90% coverage, and the approach toward saturation; trends that are consistent with and reinforce the findings reported for HPRC2 onward in the main text.

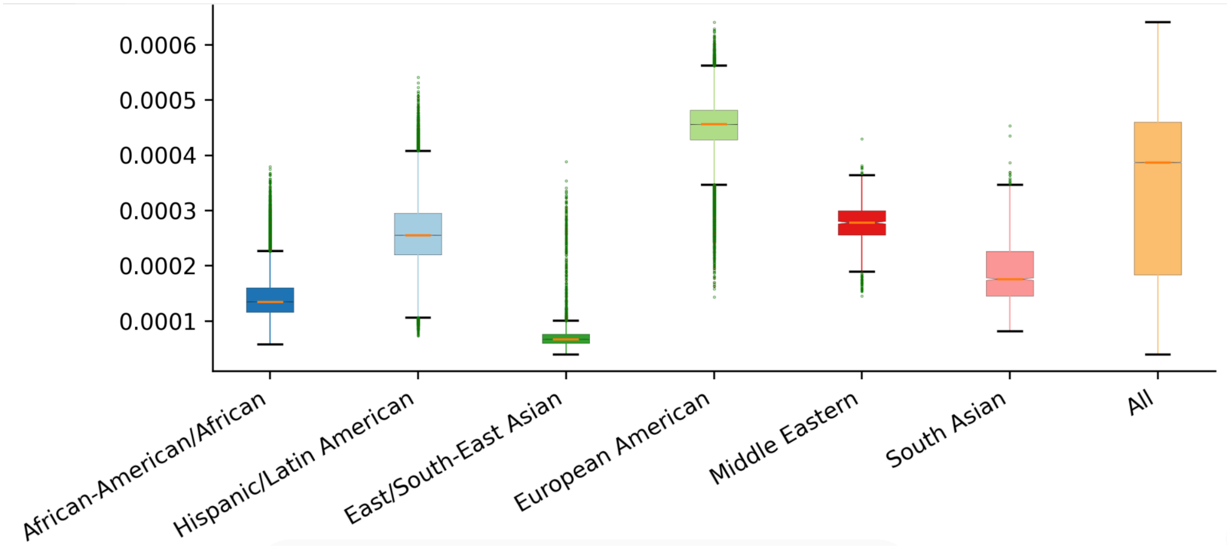


**Supplementary Fig. 2 max_var_missingness. Distribution of individual-level missingness in HPRC2 across ancestry groups present in the All of Us Research Program cohort.** For each All of Us participant, residual missingness is defined as the proportion of common variants carried by the individual (either heterozygously or homozygously) that are absent from the HPRC2 reference panel. Ancestry groups are defined and provided by the All of Us Research Program, computed as genetic-similarity bins against a combined HGDP and 1000 Genomes reference rather than self-identified. Each box displays the interquartile range, with the median shown as a horizontal line; whiskers extend to 1.5x the interquartile range, and points beyond this range of shows as individual outlines.

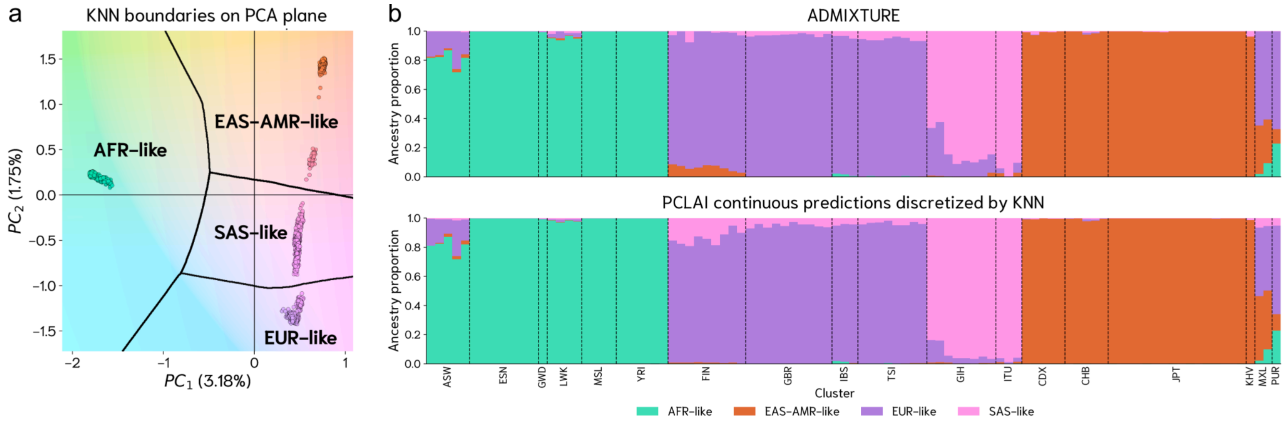


**Supplementary Fig. 3 KKN_PCLAI_vs_ADMIXTURE_comparison**: ***Concordance between discretized PCLAI genetic ancestry assignments with ADMIXTURE clusters***. **a**, Reference PCA space (PC1-PC2) with 15-nearest-neighbors (KNN) classifier decision boundaries defining four unsupervised regions used to discretize continuous PCLAI coordinates into region labels: European-like (EUR-like), South Asian-like (SAS-like), African-like (AFR-like), and East Asian-Indigenous American-like (AMR/EAS-like); **b**, Comparison of genome-wide genetic ancestry proportions inferred by ADMIXTURE at K=4 and by discretized PCLAI across the

100-sample benchmark, grouping samples by 1000G subpopulations. For PCLAI, continuous per-window predictions were assigned to the KNN-defined regions, counted genome-wide, and normalized to obtain per-sample proportions.  ADMIXTURE cluster proportions were aligned to the same PCLAI dominant genetic ancestry assignments. Discretized PCLAI ancestry assignments closely match ADMIXTURE's clustering.

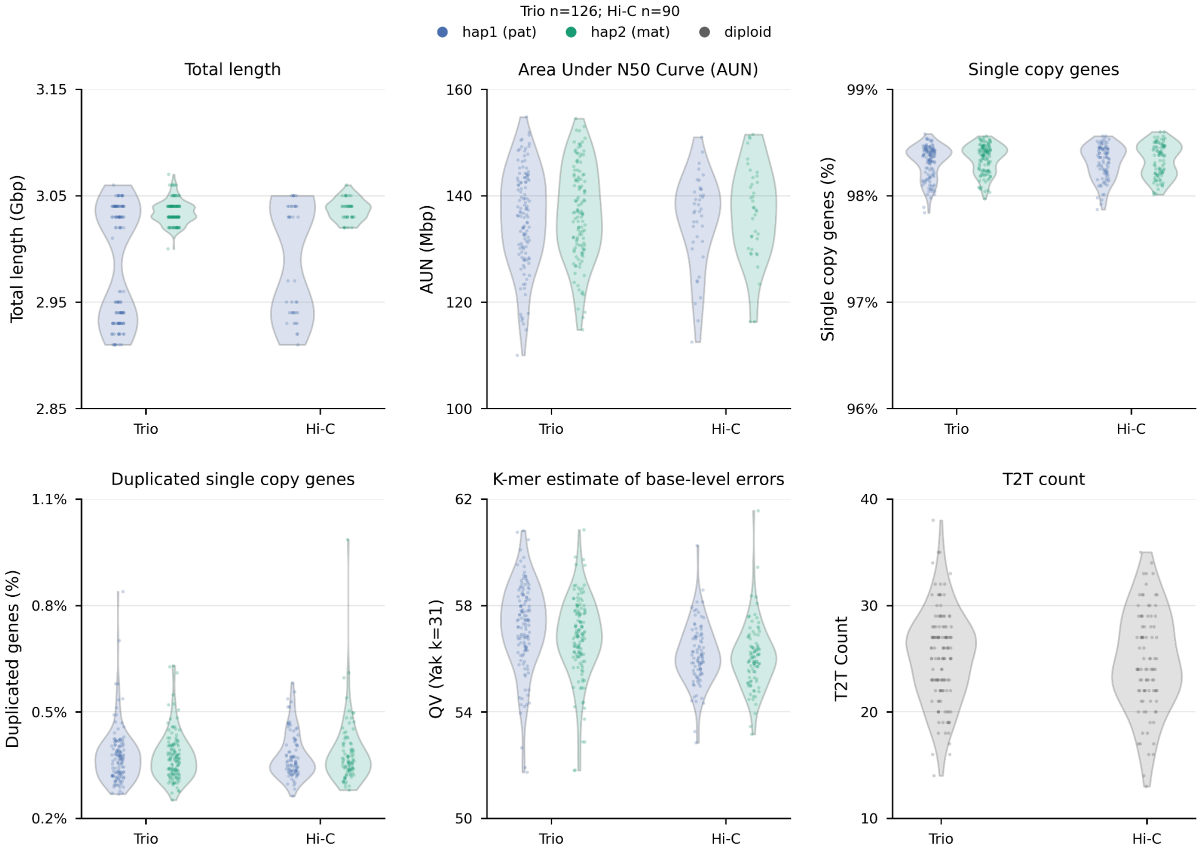


**Supplementary Fig. 4 assembly_qc_trio_vs_hic: Assembly QC metrics are similar between trio- and Hi-C-phased HPRC release 2 assemblies.** Violin plots show assembly QC metrics for HPRC2 assemblies grouped by phasing strategy. Single copy and duplicated single copy gene percentages are from asmgene. QV was estimated from yak (k=31). A two-sided Kolmogorov-Smirnov test was used for total length and two-sided Mann-Whitney U tests were used for all other metrics. Trio and Hi-C distributions were not significantly different for total length (P=0.857), AUN (P=0.949), single copy genes (P=0.818), duplicated single copy genes (P=0.611), or T2T count (P=0.500). QV was significantly higher in trio-phased assemblies than Hi-C-phased assemblies (median 57.09 vs 56.04; P = 4.17 × 10^-14).

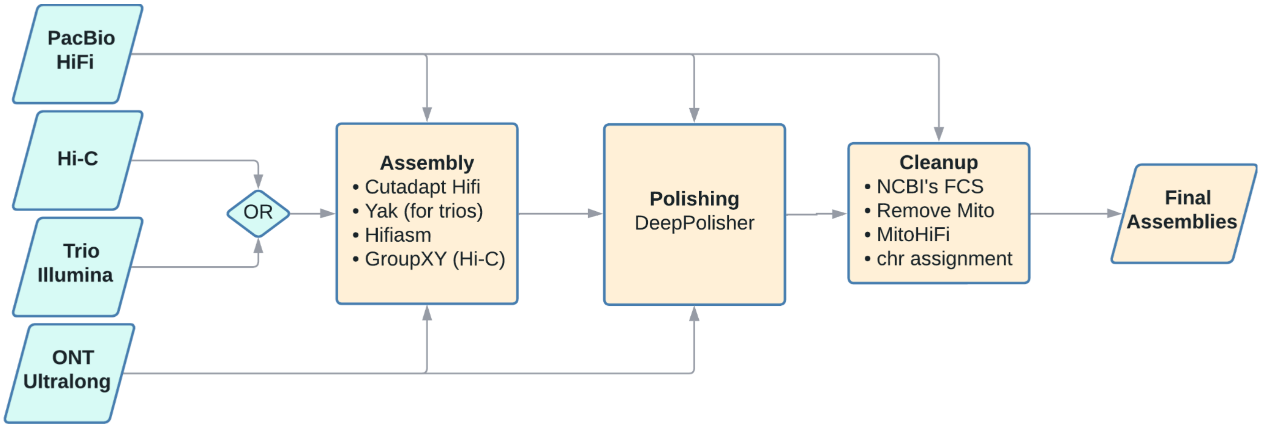


**Supplementary Fig. 5 hprc_r2_assembly_workflow: HPRC2 assembly generation workflow.** Assemblies are generated with PacBio HiFi and ONT Ultralong reads with trio or Hi-C phasing. HiFi reads have residual adapters trimmed with Cutadapt. Hi-C phased assemblies from male samples have sequences from sex chromosomes identified and grouped into haplotype 1 (chrY) and haplotype 2 (chrX). Assemblies were polished with HiFi reads aligned to the assemblies with the aid of ONT UL reads for phasing. Polished assemblies had foreign contamination such as EBV removed and the mitochondrial sequences were removed and replaced with versions from MitoHifi. Chromosome assignments for T2T contigs and scaffolds were identified for Genbank upload.

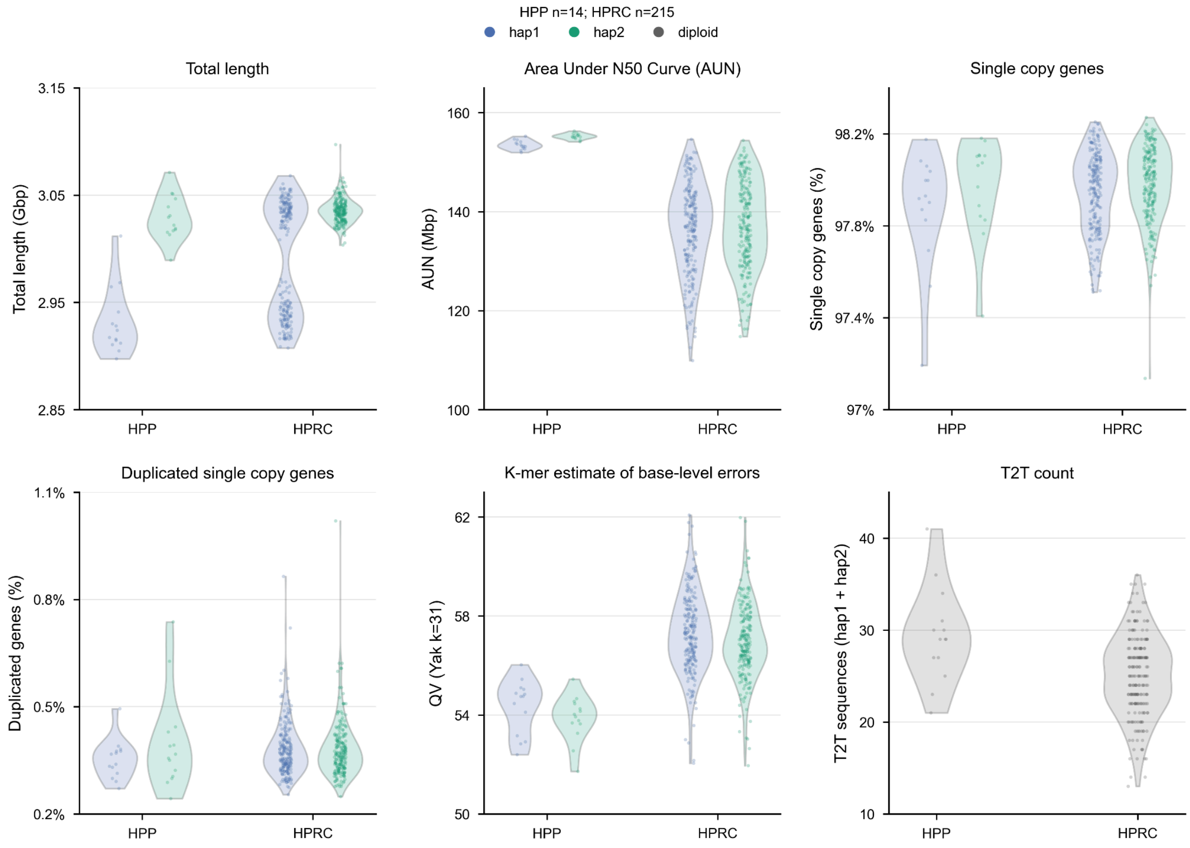

**Supplementary Fig. 6 assembly_qc_by_project: Assembly QC metrics for assemblies produced by the HPRC and HPP.** Violin plots show assembly QC metrics for HPRC2 assemblies grouped by project. Single copy and duplicated single copy gene percentages are from asmgene. QV was estimated from yak (k=31). A two-sided Kolmogorov-Smirnov test was used for total length and two-sided Mann-Whitney U tests were used for all other metrics. The KS test on assembly length was subset to male samples to compensate for HPP only including male samples which reduces the length of the chrY containing haplotype (hap1). Distributions were not significantly different for total length (P=0.199), total single copy genes (P=0.692), or duplicated single copy genes (P=0.256). HPP assemblies did have higher contiguity with statistically higher T2T counts (median 29 vs 25; P=0.00234) and AUN (median 154.58 vs 137.12 Mbp; P=1.10 x 10 ^-18). QV was significantly higher in HPRC assemblies than in HPP assemblies (median 57.05 vs 54.20; P = 6.92 × 10^-16).

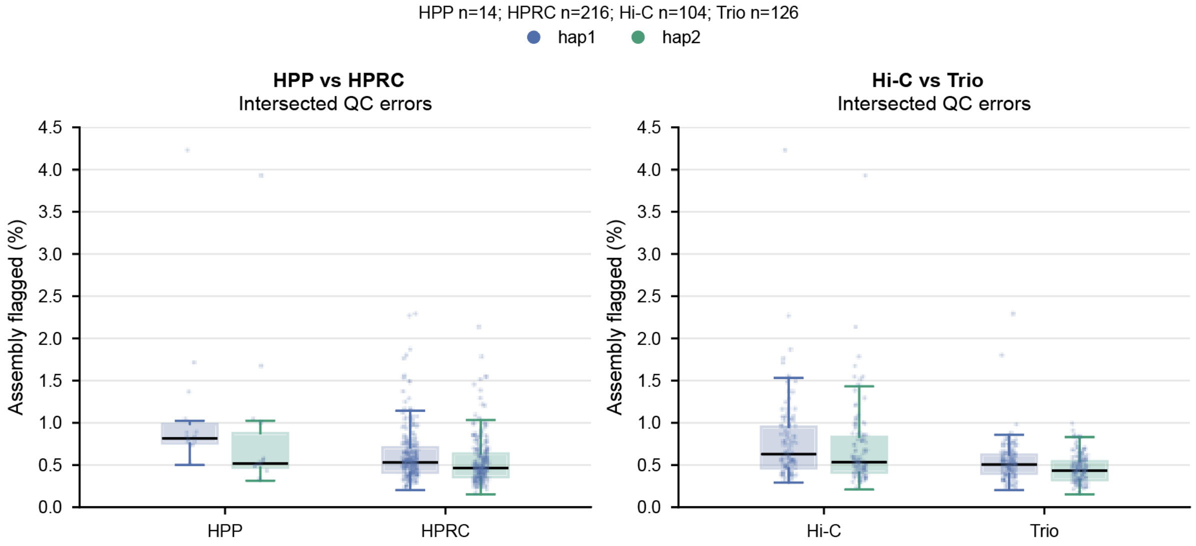


**Supplementary Fig. 7 flagged_percent_by_type: Fraction of assemblies with predicted QC flags for different producers (HPRC and HPP) and phasing strategy (Hi-C and trio).** Data points show the percent of each assembly with predicted QC errors from intersected QC bed files (see methods). Two-sided Mann-Whitney U tests were used to determine statistical significance. HPP assemblies had more assembled sequence predicted as erroneous (median 0.758% vs 0.504%; P=7.77 × 10^-5) than HPRC. HPP assemblies had some differences in workflow (assembler and post-assembly scaffolding) that led to increases in contiguity and the increase in flagged sequence may reflect insertion of more difficult regions into the assemblies. Hi-C and trio assemblies had considerable overlap in distributions, but Hi-C assemblies have more predicted error sequences (median 0.565% vs 0.469%; P=5.84 × 10^-11).

a

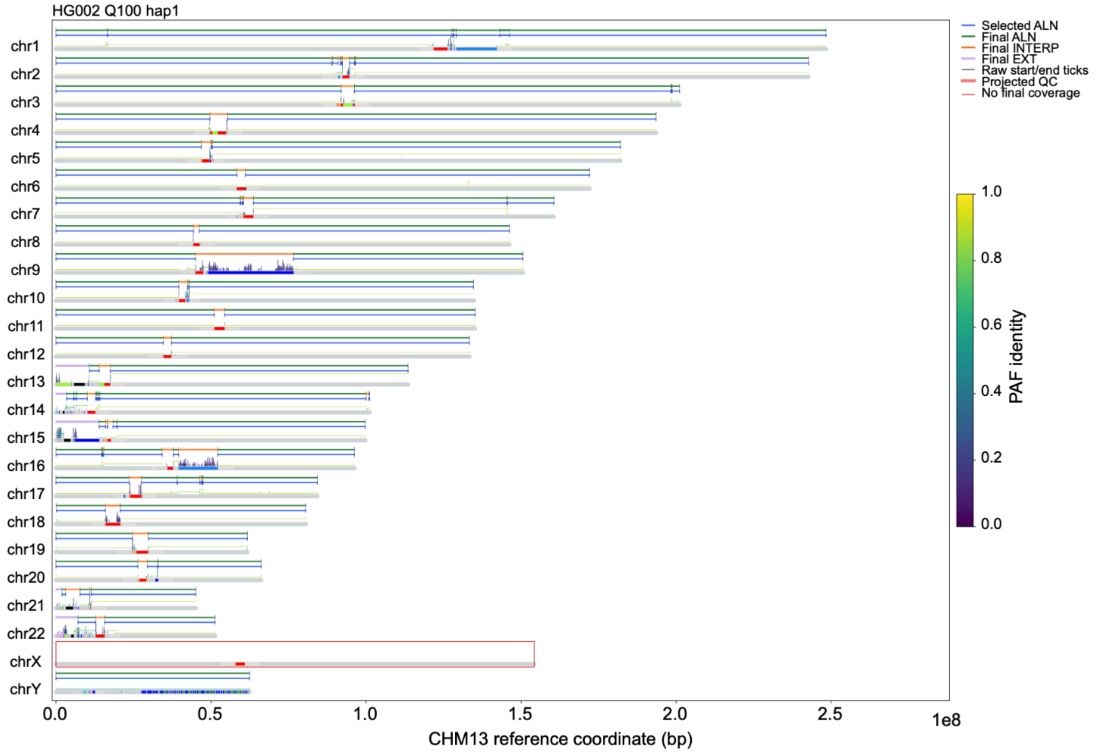


b

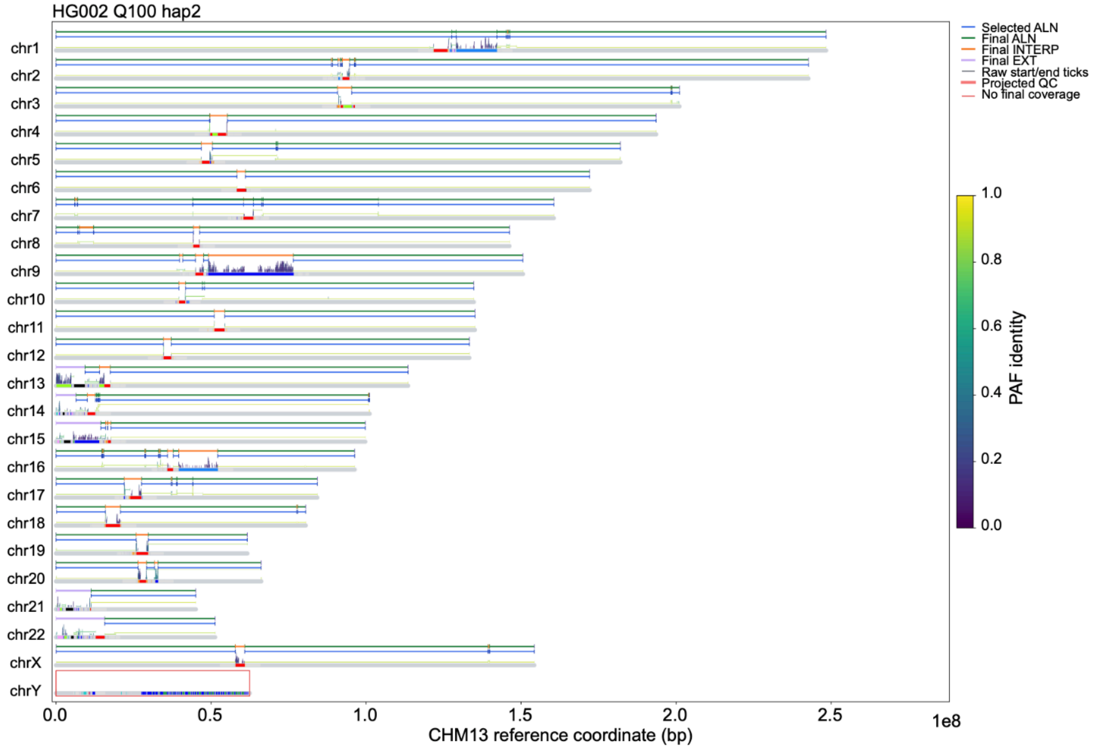


**Supplementary Fig. 8 HG002_Q100_qc_projection: HG002 Projections show complete coverage of CHM13 chromosomes.** The projection process is represented vertically for each chromosome (shown with centromeric satellite annotations). Raw Minigraph alignments between the assembly and CHM13 v2.0 are shown directly above the reference. Custom scripts filter, merge, and select alignments (shown in blue as Selected ALN). The final alignment includes interpolated alignments across large alignment gaps (Final INTERP) and extensions to the sequence ends (Final EXT). Regions without coverage are outlined in red (no QC errors were input to the projection for HG002 Q100). The paternal haplotype (hap1) shown (**a**) has no coverage of chrX as is expected for a male sample. The maternal haplotype (hap2) shown in (**b**) has no coverage of chrY.

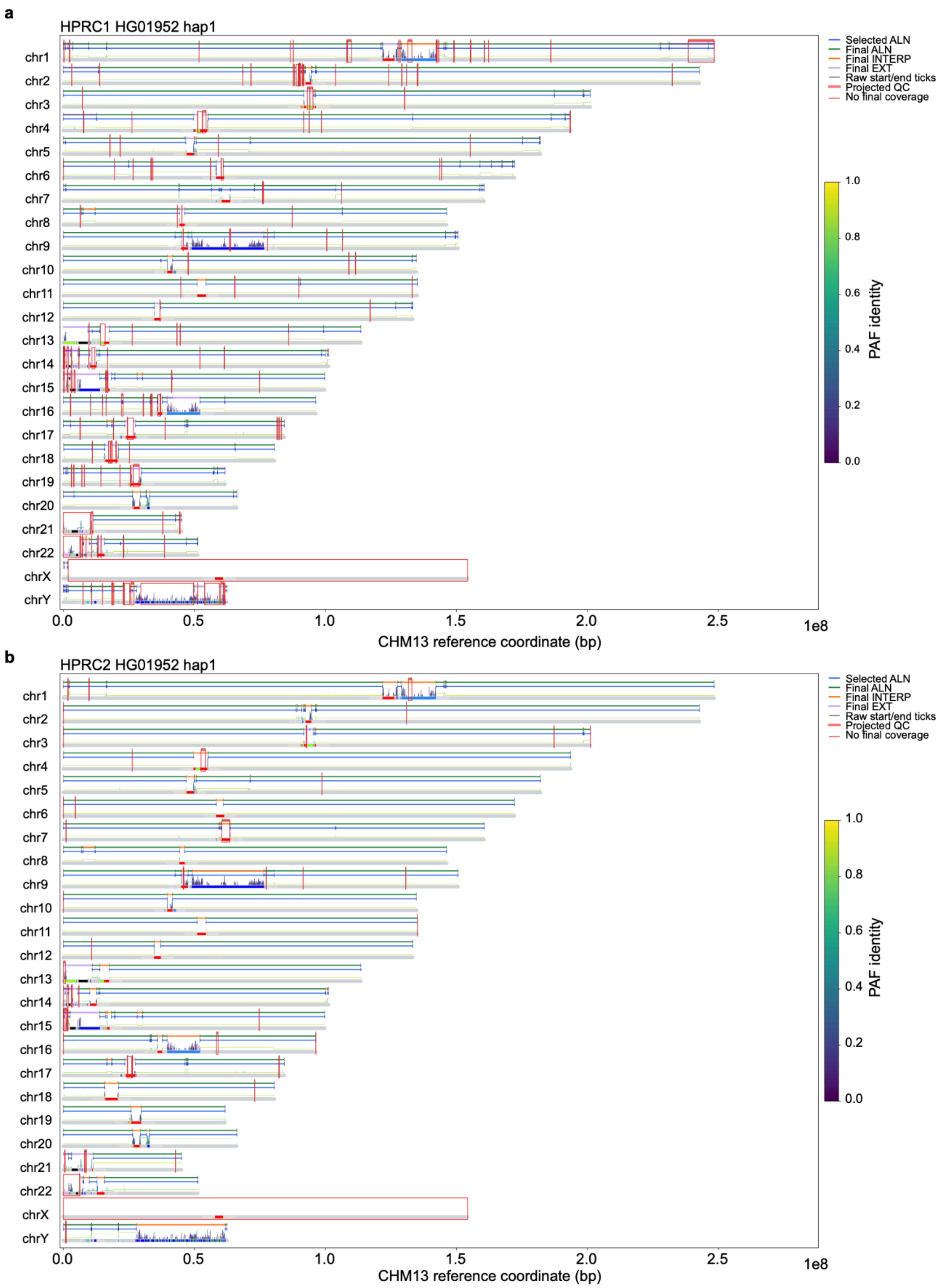


**Supplementary Fig. 9 HG01952_hap1_projected_qc. HG01952 Haplotype 1 Projected QC From HPRC1 and HPRC2 show improvements in assembly quality.** The projection process is represented vertically for each chromosome (shown with centromeric satellite annotations). Raw Minigraph alignments between the assembly and CHM13 v2.0 are shown directly above the reference. Custom scripts filter, merge, and select alignments (shown in blue as Selected ALN). The final alignment includes interpolated alignments across large alignment gaps (Final INTERP) and extensions to the sequence ends (Final EXT). Regions without coverage or with projected QC are outlined in red. The HPRC1 assembly (**a**) has more projected QC error regions than HPRC2 (**b**).

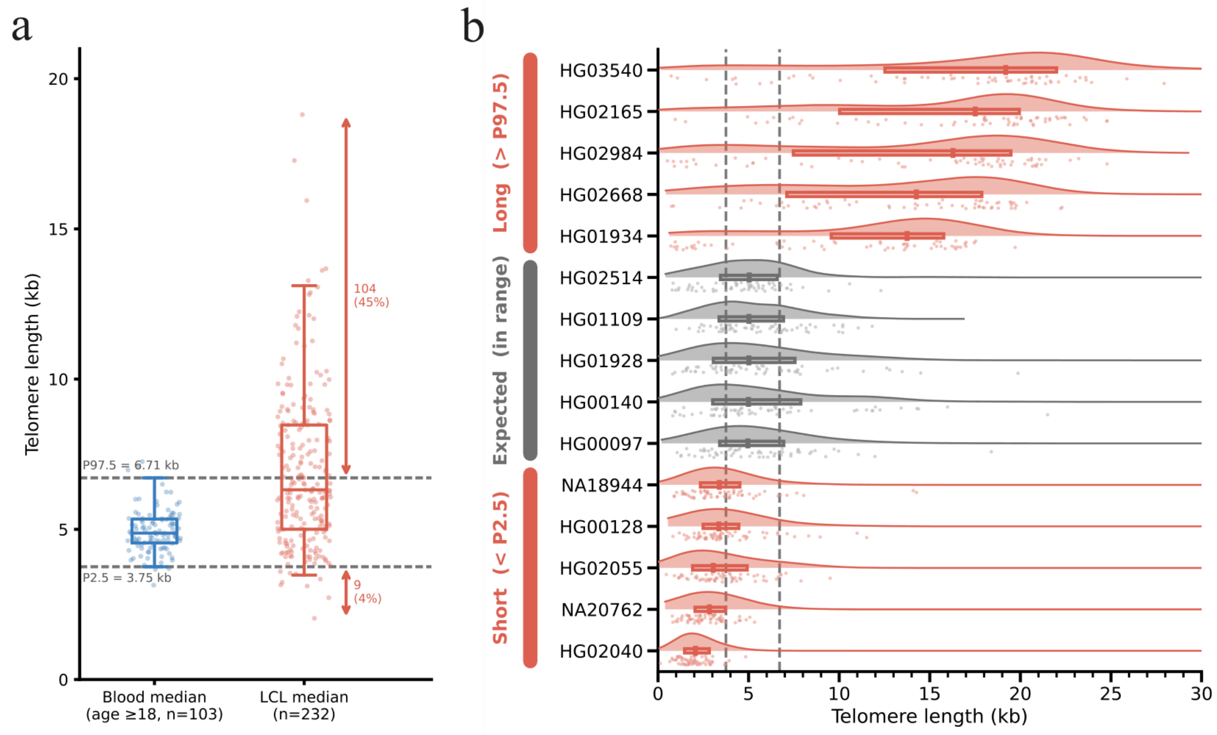


**Supplementary Fig. 10 telomere_lengths_in_lcls**. a. Comparison of ONT-based per-sample median telomere lengths from Telomere Profiling in Karimian et al. (2024) PBMC data (left) and per-sample median telomere lengths from HPRC R2 LCLs (right). Boxes indicate the 25th-75th percentile range with median and dashed lines indicate the 2.5th and 97.5th percentiles (P2.5, P97.5) of the blood telomere length distribution. b. Each panel of per read telomere lengths for a single HPRC sample (kernel density estimate, individual reads as points, and box indicating the 25th-75th percentile range with median). Samples are grouped as short (median below the 2.5th percentile of the blood reference), expected (median within the 2.5th and 97.5th percentile of blood reference), or long (median above the 97.5th percentile of the blood reference). Vertical dashed lines indicate the 2.5th and 97.5th percentile of the adult blood reference distribution from Karimian et al. (2024).

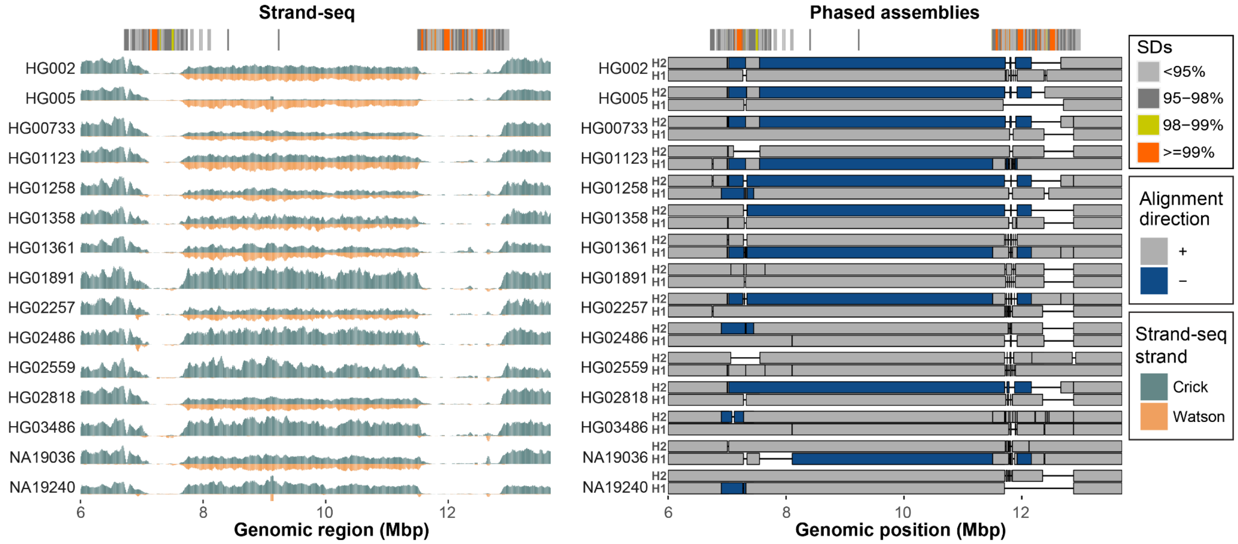


**Supplementary Fig. 11 S-SSQvalidation: 8p23.1 inversion validation.**
Left: Read-coverage profiles of Strand-seq data over the 8p23.1 region summarized as binned (bin size: 50 kbp step size: 10 kbp) read counts represented as bars above (teal; Crick read counts) and below (orange; Watson read counts) the midline. The inversion polymorphism lies between the SD blocks (colored by sequence identity) marked at the top of the plot. Here, equal coverage of Watson and Crick counts represents a heterozygous inversion (e.g HG002) as only one homologue is inverted with respect to the reference (T2T-CHM13). The homozygous inversion (e.g HG005) is visible as complete switch from Crick to Watson reads over the inverted region. Right: Alignment directionality of fully assembled haplotypes (H1 - bottom and H2 top) for 15 samples (30 haplotypes) over the 8p23.1 region. Forward ('+') oriented alignments are shown in gray while reverse oriented alignments in light green ('-'). On top there is an SD annotation (SDs) colored by sequence identity.

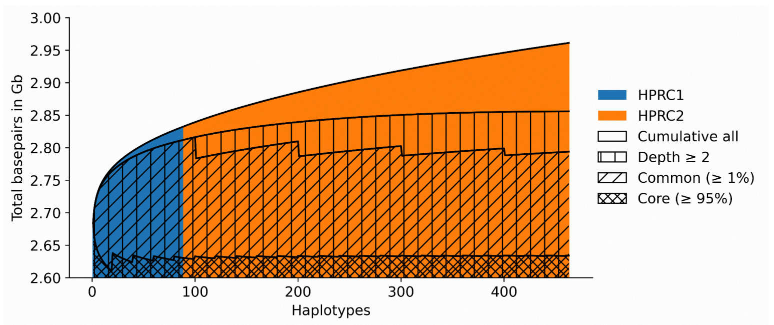


**Supplementary Fig. 12 S-panacus_autosomes_section_growth_bp_grch38: Growth plot for total basepairs in the GRCh38-based HPRC2 MC graph starting from the HPRC1 haplotypes.** Although the total number of base pairs continues to increase with each added haplotype, the amount of sequence shared by at least two haplotypes (depth ≥2) approaches saturation. This indicates that newly added sequence is predominantly rare, while the representation of common human variation is nearing completeness. The seamless and indistinguishable transition between the two groups implies that the HPRC1 and HPRC2 populations display a similar degree of genomic variability. Note that this plot only quantifies growth on the haplotypes of the pangenome graph, excluding the sequence unique to GRCh38.

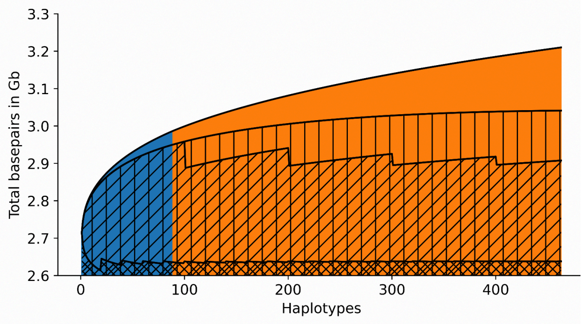


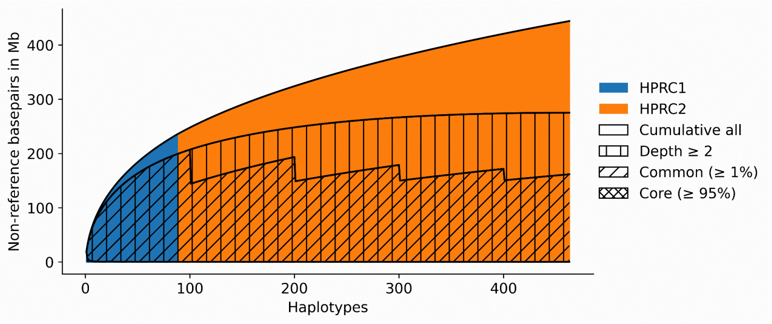


**Supplementary Fig. 13 S-panacus_autosomes_section_growth_bp_chm13: Growth plot for total (left) and non-reference (right) basepairs in the T2T-CHM13-based HPRC2 MC graph starting from the HPRC1 haplotypes.** The growth patterns closely mirror those observed for the GRCh38-based graph; however, the T2T-CHM13-based graph captures approximately 250 Mb additional total sequence, including approximately 150 Mb of additional non-reference sequence.

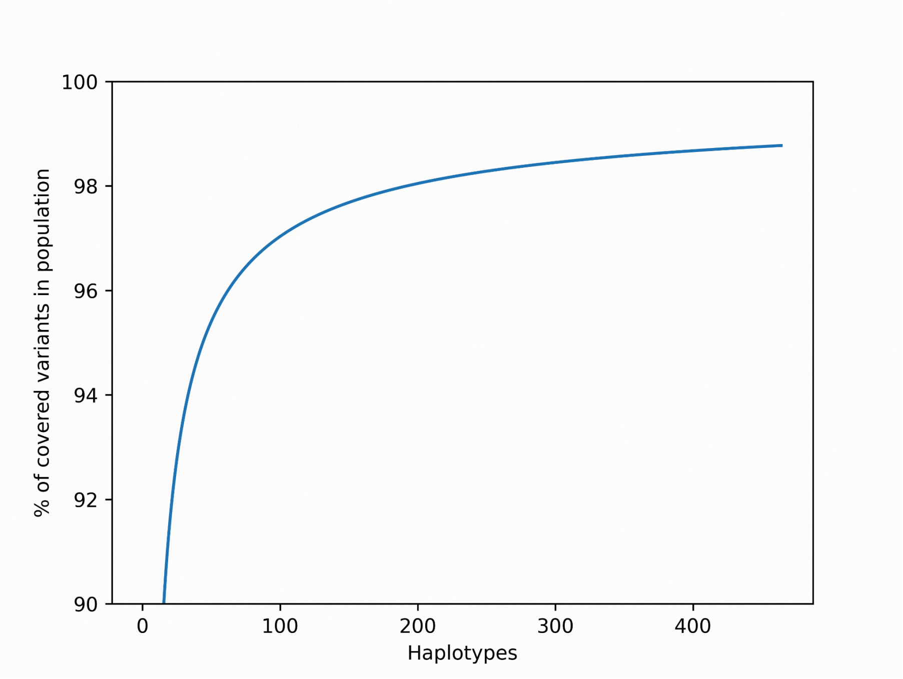


**Supplementary Fig. 14 S-variant-coverage-autosomes_hprc-v2.0-mc-chm13.wave: Population coverage of the VCFwave variants of the T2T-CHM13-based Minigraph-Cactus graph.** This plot displays the probability of a randomly sampled variant from the whole population to be already known, based on the number of haplotypes that are already part of the pangenome.

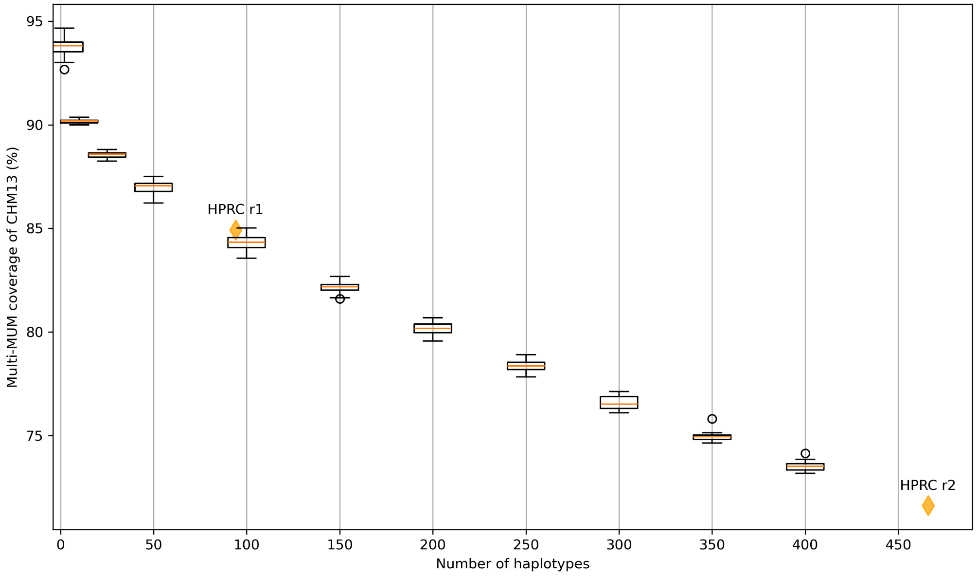


**Supplementary Fig. 15 mum_conservation. Multi-MUM coverage for increasing pangenome sizes.** Multi-MUM blocks represent the "core" pangenome, comprising regions that are syntenic across all assemblies in a collection. Coverage of the core pangenome decreases with increasing sample size due to capture of SVs that disrupts synteny. Multi-MUM blocks consist of collinear multi-MUMs separated by less than 100 bp. The proportion of T2T-CHM13 covered by collinear blocks was computed as the number of bases in collinear blocks divided by the length of T2T-CHM13.

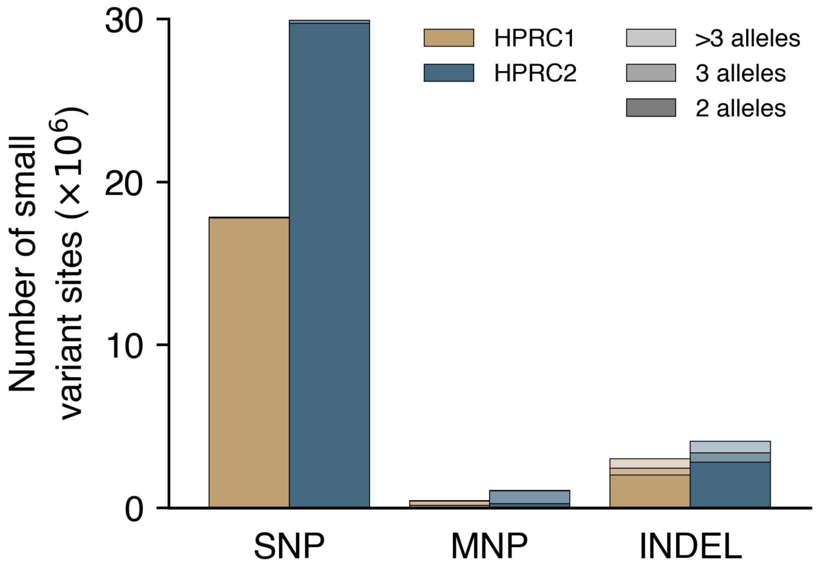


**Supplementary Fig. 16 small_variant_number_chm13. Autosomal small variant site composition in the T2T-CHM13-based HPRC1 and HPRC2 MC graphs.** Autosomal small variant sites in the T2T-CHM13-based HPRC1 and HPRC2 MC graphs, stratified by variant type and the number of alleles per site. MNP, multinucleotide polymorphism.

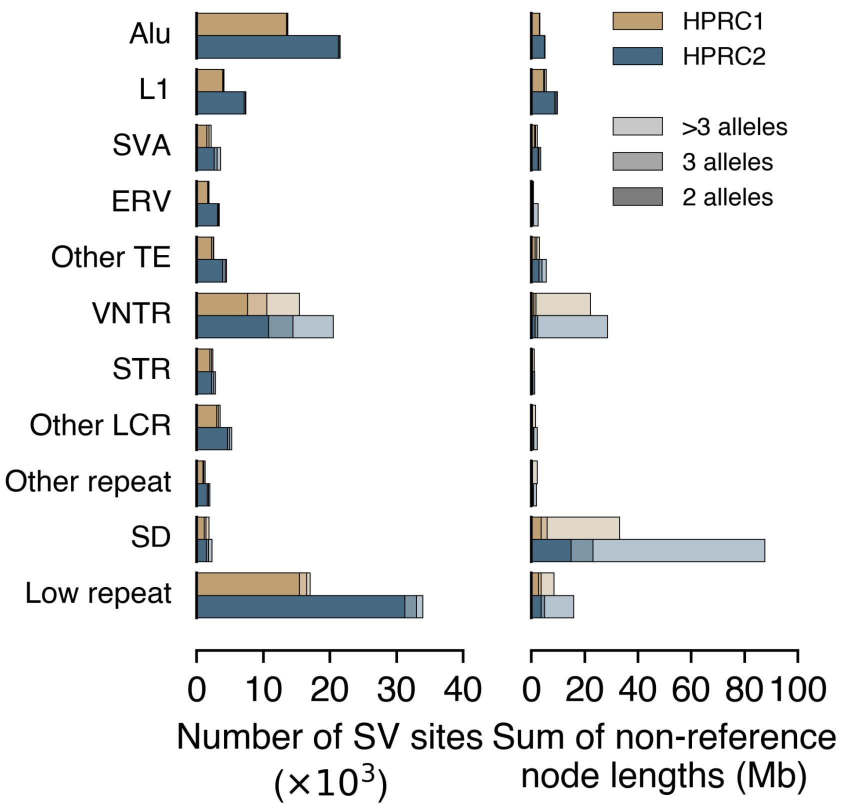


**Supplementary Fig. 17 sv_number_chm13. Autosomal SV site composition in the T2T-CHM13-based HPRC1 and HPRC2 MC graphs.** Autosomal SV sites in the T2T-CHM13-based HPRC1 and HPRC2 MC graphs, stratified by repeat class and the number of alleles per site. The left plot shows the number of SV sites, and the right plot shows the sum of non-reference node lengths. Other TE, a site involving mixed classes of transposable elements (TEs). VNTR, variable-number tandem repeat, a tandem repeat with unit motif length ≥7 bp. STR, short tandem repeat, a tandem repeat with unit motif length ≤6 bp. Other LCR, low-complexity regions involving mixed VNTR and STR categories or low-complexity regions without a clear VNTR or STR pattern. Other repeat, a site involving mixed repeat classes. SD, segmental duplication. Low repeat, a site in which only a small fraction of the longest allele is annotated as repeat sequence.

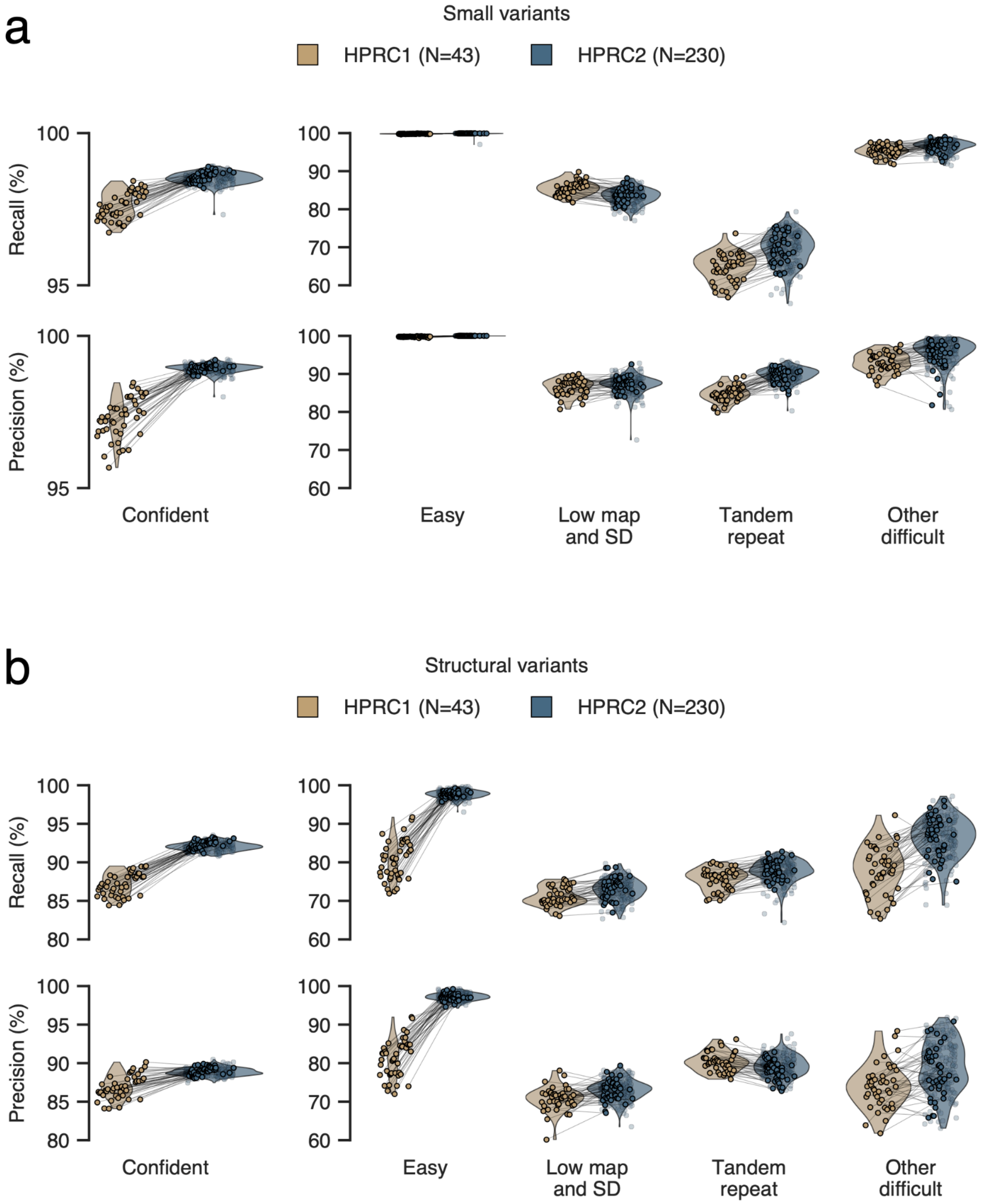


**Supplementary Fig. 18 graph_variant_benchmarking_chm13. Benchmarking variant calling performance in the T2T-CHM13-based HPRC1 and HPRC2 MC graphs. a-b,** Precision and recall of autosomal small variants (a) and SVs (b) in the T2T-CHM13-based HPRC1 and HPRC2 MC graphs relative to per-sample joint ground truth callsets. Joint ground truth callsets were generated by retaining variants supported by at least two of 14 callers in merged callsets. Comparisons were restricted to dipcall confident regions and stratified by Genome in a Bottle (GIAB) v3.6 genomic context. Each point represents one sample, and matched samples between HPRC1 and HPRC2 are connected by lines. Low map, low mappability. SD, segmental duplication. The lower recall in tandem repeat regions relative to the GRCh38-based MC graphs may reflect the fact that T2T-CHM13 contains more than twice as much tandem repeat sequence as GRCh38, resulting in a larger fraction of variants occurring in repetitive sequence where mapping and variant detection are more challenging.

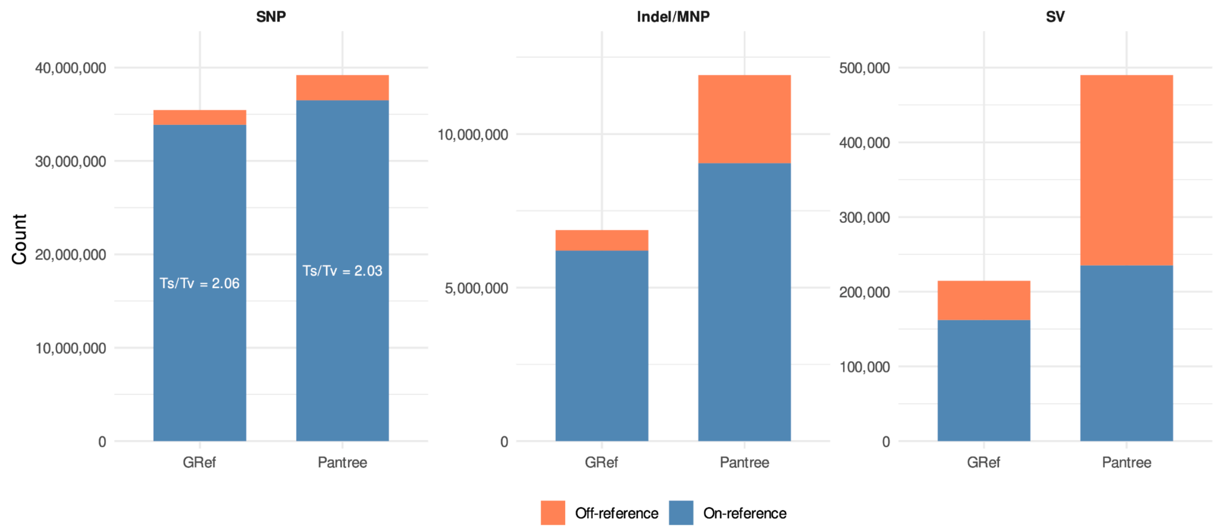


**Supplementary Fig. 19 gref_pantree. Comparison of variant counts using GRef and Pantree coordinates.** Stacked bar plots show the numbers of on-reference and off-reference SNPs, indels/MNPs, and SVs identified using the GRef and Pantree coordinates. The transition-to-transversion (Ts/Tv) ratio is shown for SNPs.

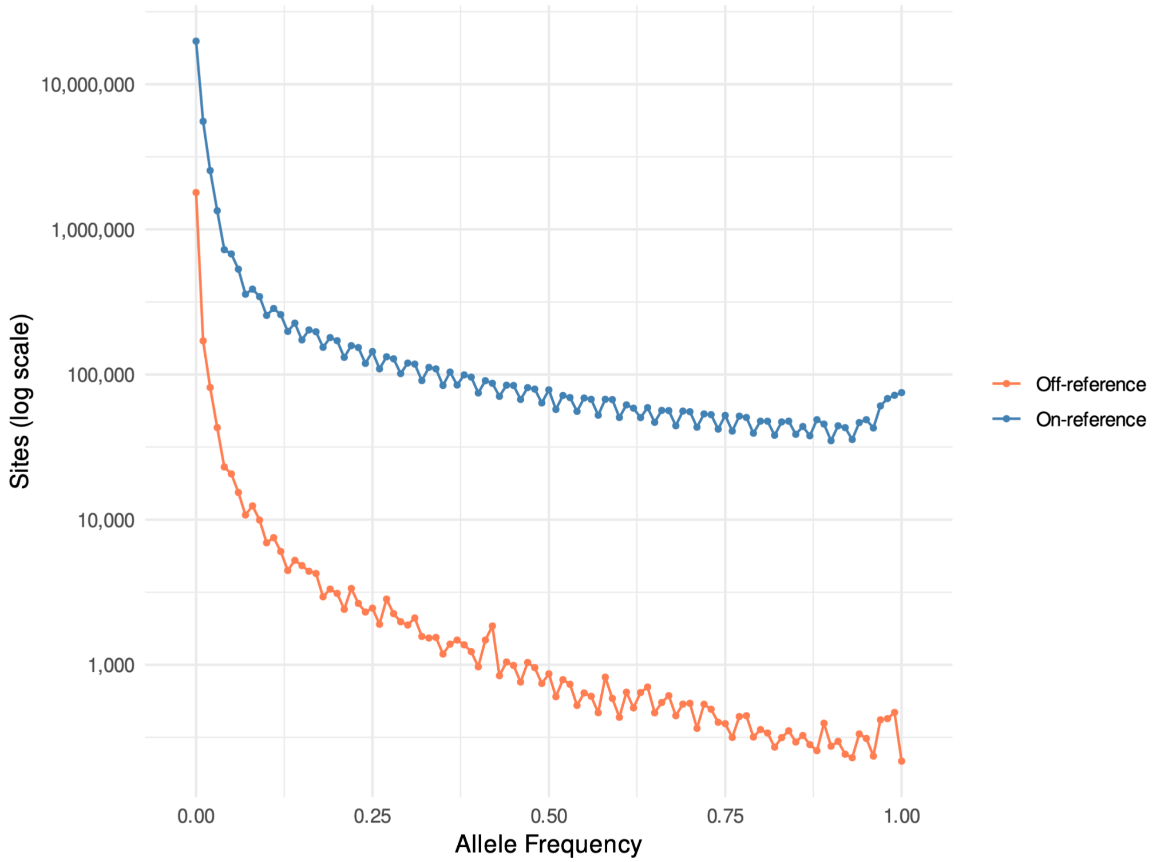


**Supplementary Fig. 19 gref_allele_freq_supp. Allele frequency distribution of variants represented by GRef coordinates.** The number of on-reference and off-reference variant sites is shown as a function of alleel frequency. The y axis is shown on a logarithmic scale.

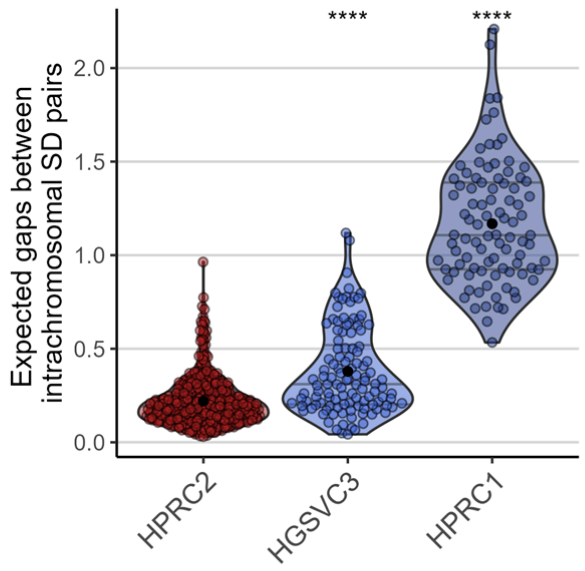

**Supplementary Fig. 20 SD1. Average number of gaps between two intrachromosomal SD pairs.** Each point represents a haplotype assembly. Two-sided Wilcoxon test significance against HPRC2 is indicated by ‘****’ (p<0.0001) on the top of the distribution.

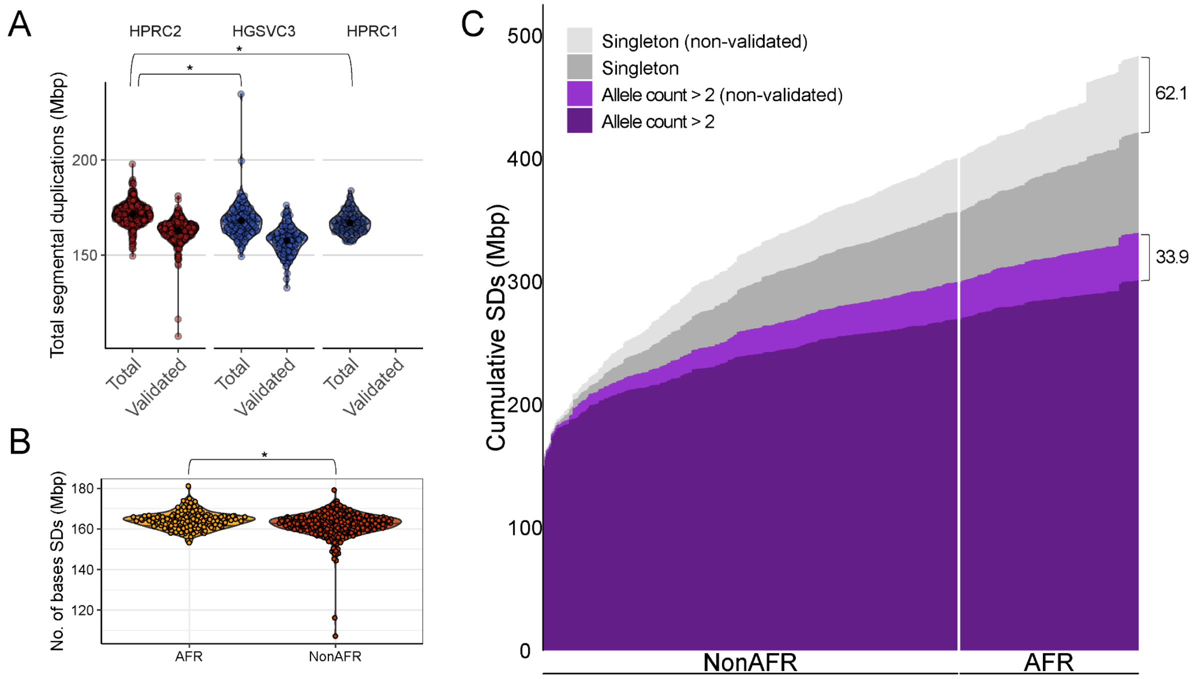


**Supplementary Fig. 21 SD2. Quantification of SD and validation by read-depth.** A) Comparison of total number of SD bases per haplotype assemblies from the current study (HPRC2), and previous studies (HGSVC3[1] and HPRC1[2]) are shown. The number of SDs validated by read depth (Methods) is indicated next to total SDs. B) Amount of validated SDs compared between AFR and non-AFR (NonAFR) populations. For A and B, two-sided Wilcoxon test significance is indicated by ‘*’ (p<0.001) on the top of the distribution. C) Cumulative curve of SDs found in the haplotype genome assemblies (n=459). The x-axis indicates haplotype assemblies while the y-axis shows cumulative sum of maximum length SDs. SDs that are singleton are indicated in grey and SDs with an allele count > 2 are purple. SDs supported and not supported by read depth are further classified by darker and lighter colors, respectively; 62.1 Mbp were non-validated by read depth for singleton SDs and 33.9 Mbp for allele count > 2.

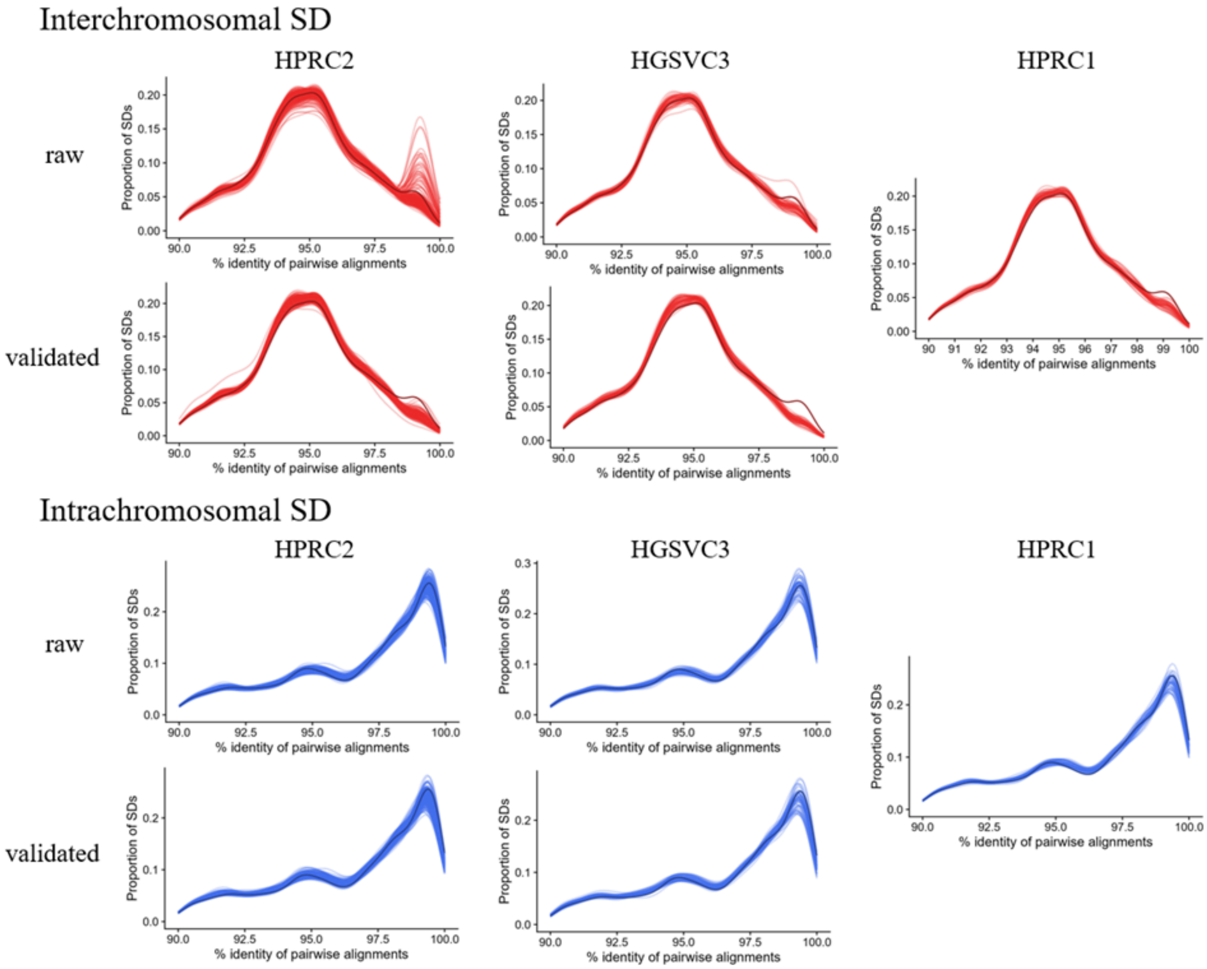


**Supplementary Fig. 22 SD3. Pairwise sequence identity distribution between SD pairs.** Inter- and intrachromosomal SD duplication are shown on the top and bottom, respectively. Each column shows different datasets, including HPRC2, HGSVC3 and HPRC1, while the row indicates before and after read-depth validation. For the genome-wide SDs, summarized in the x-axis is the sequence identity with y-axis showing density weighted by the number of bases. Indicated by dark red and blue are the distribution observed in T2T-CHM13v2.0.

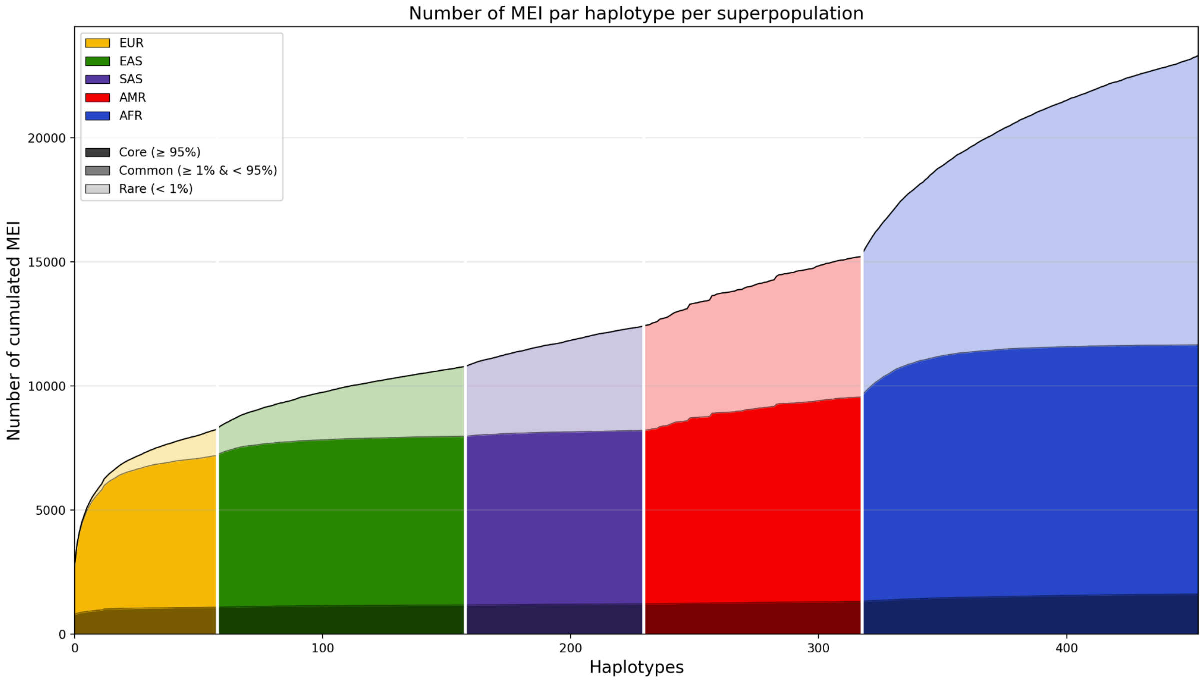


**Supplementary Fig. 23 MEI haplotype sharing and frequency across populations** Each column represents a distinct haplotype, corresponding to the five 1000G superpopulations (EUR, EAS, SAS, AMRand AFR). Numbers of cumulated MEI for each haplotype are colored according to their superpopulation and frequency across the combined dataset: e.g. dark blue correspond to “core” MEI (≥95% frequency), blue correspond to “common” MEI (≥1% and <95%), and light blue correspond to “rare” MEI (<1%).

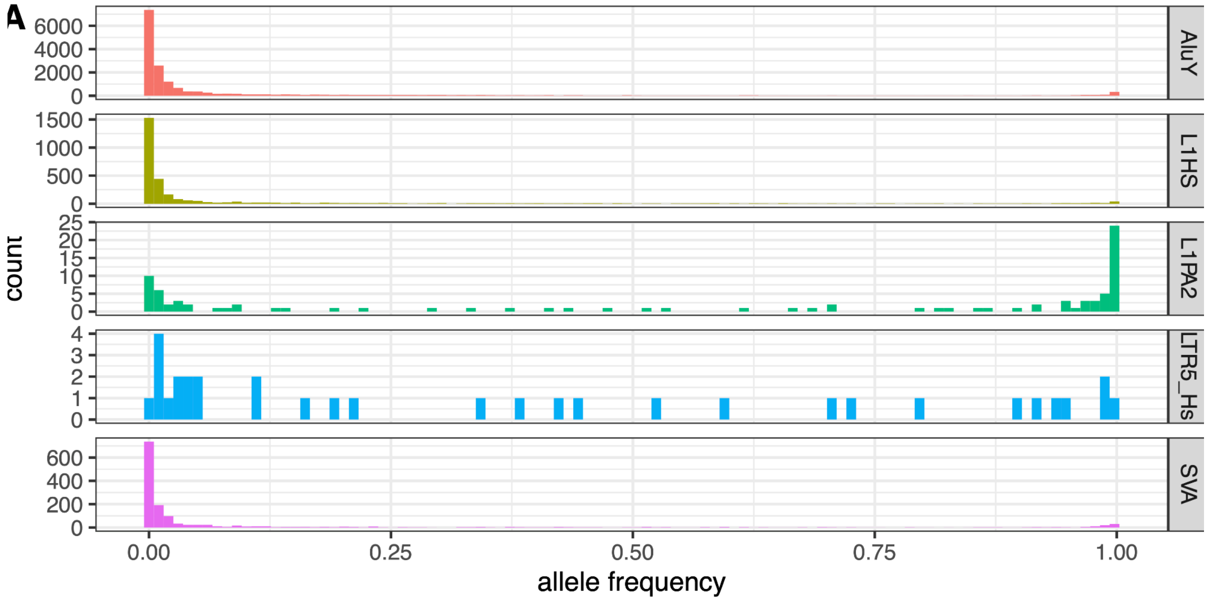


**Supplementary Fig. 24 MEI_BREAK_DOWN** Allele frequency in HPRC2 histogram of MEI families: AluY, L1HS, L1PA2, LTR5_HS, and SVA.

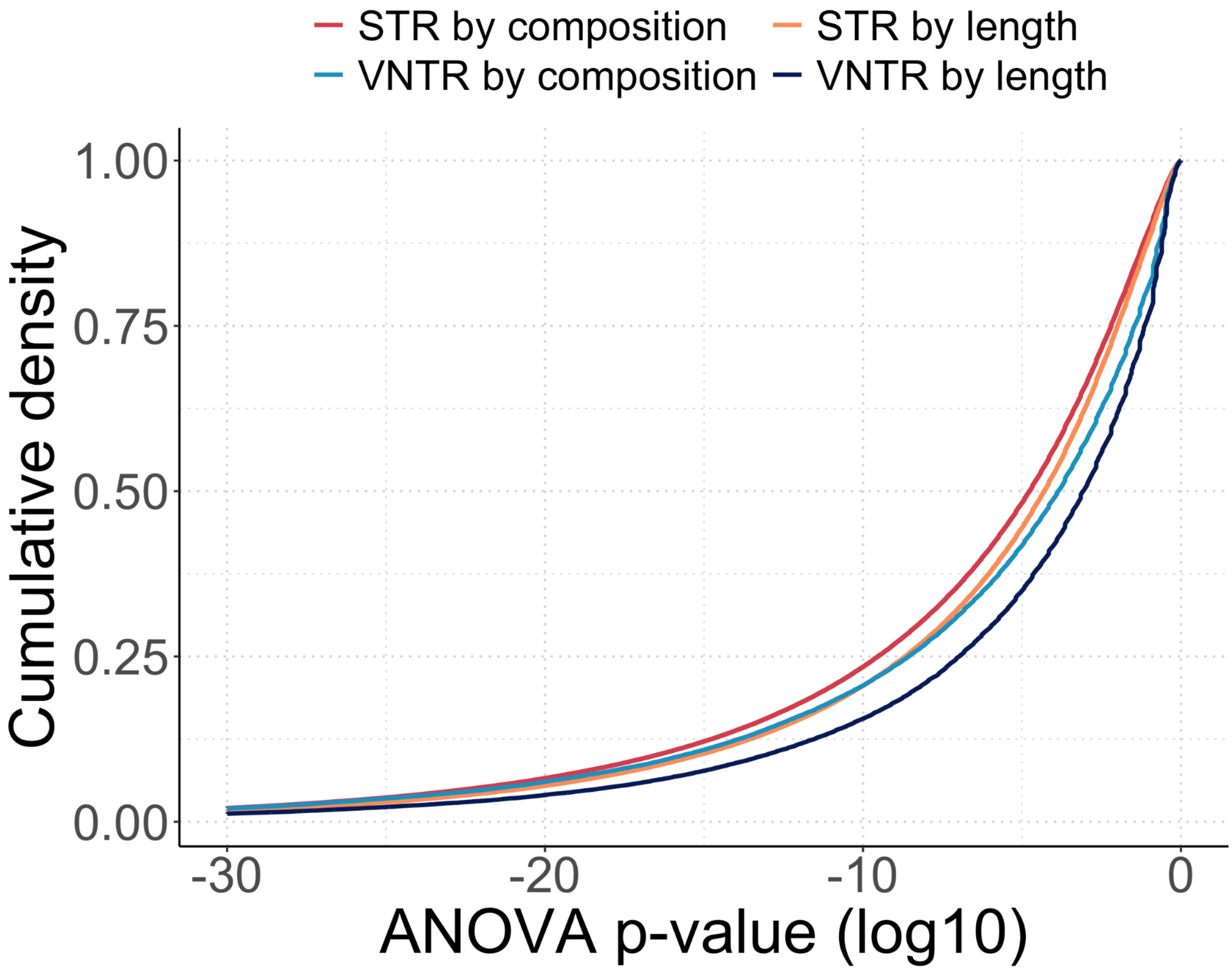


**Supplementary Fig. 25 S-TR. ANOVA of TR variability within versus between populations**. Variation was quantified by length and composition differences, shown in order of significance, p-values capped at $10^{-30}$.

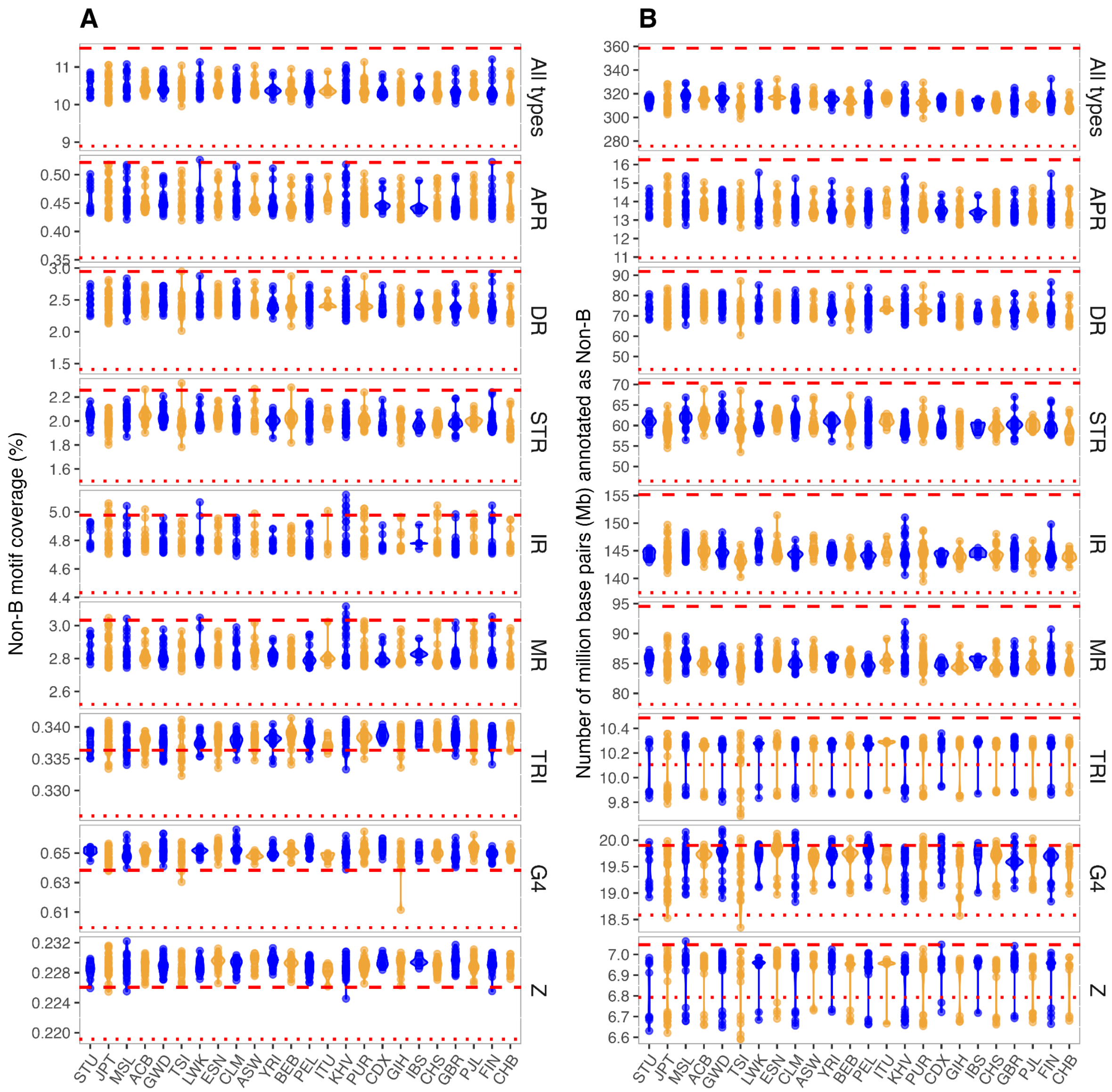


**Supplementary Fig. 26 nonB.** Non-B DNA motifs in different HPRC2 populations, sorted after overall median nonB motif density. Only populations with at least three individuals (= 6 haplotypes) are shown. Violin plots of (**A**) non-B motif coverage (in % of genome) and **(B)** number of annotated non-B base pairs. 'All' denotes all eight non-B DNA motif types combined. Red dashed lines denote the non-B motif density and base pairs in the T2T reference T2T-CHM13; Red dotted lines denote the non-B motif density and base pairs in the old human reference hg38. Population colors are alternating for visibility. Abbreviations: APR: A-phased repeats; DR: direct repeats; IR: inverted repeats; MR: mirror repeats; TRI: triplex motifs; STR: short tandem repeats; Z: Z-DNA motifs; G4: G-quadruplexes.

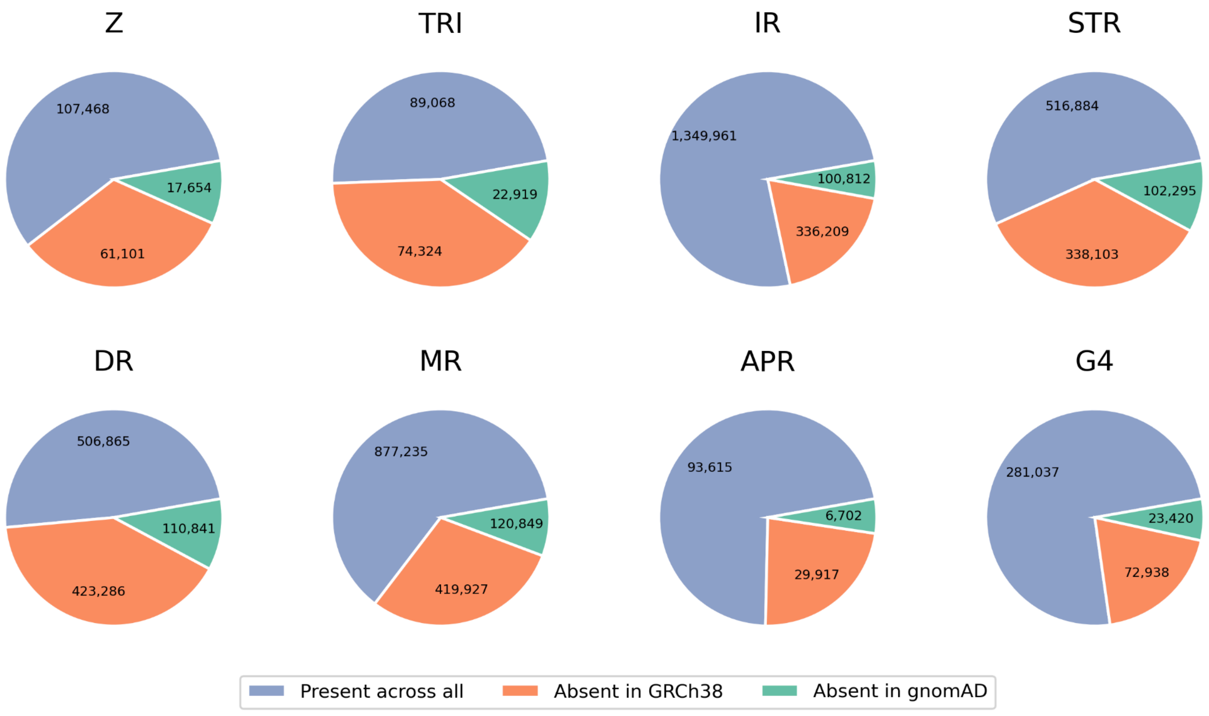


**Supplementary Fig. 27 nonB-gnomAD**. Pie plot showing SNPs at Non-B DNA motifs across the pangenome. The "present across all" represents the SNPs that are present in gnomAD database, the ones labelled "Absent in GRCh38" are the ones lost due to unmapped regions between T2T-CHM13 and GRCh38, and the ones labelled "Absent in gnomAD" are the ones that are novel in the pangenome. Abbreviations as in Supplementary Fig. nonB.

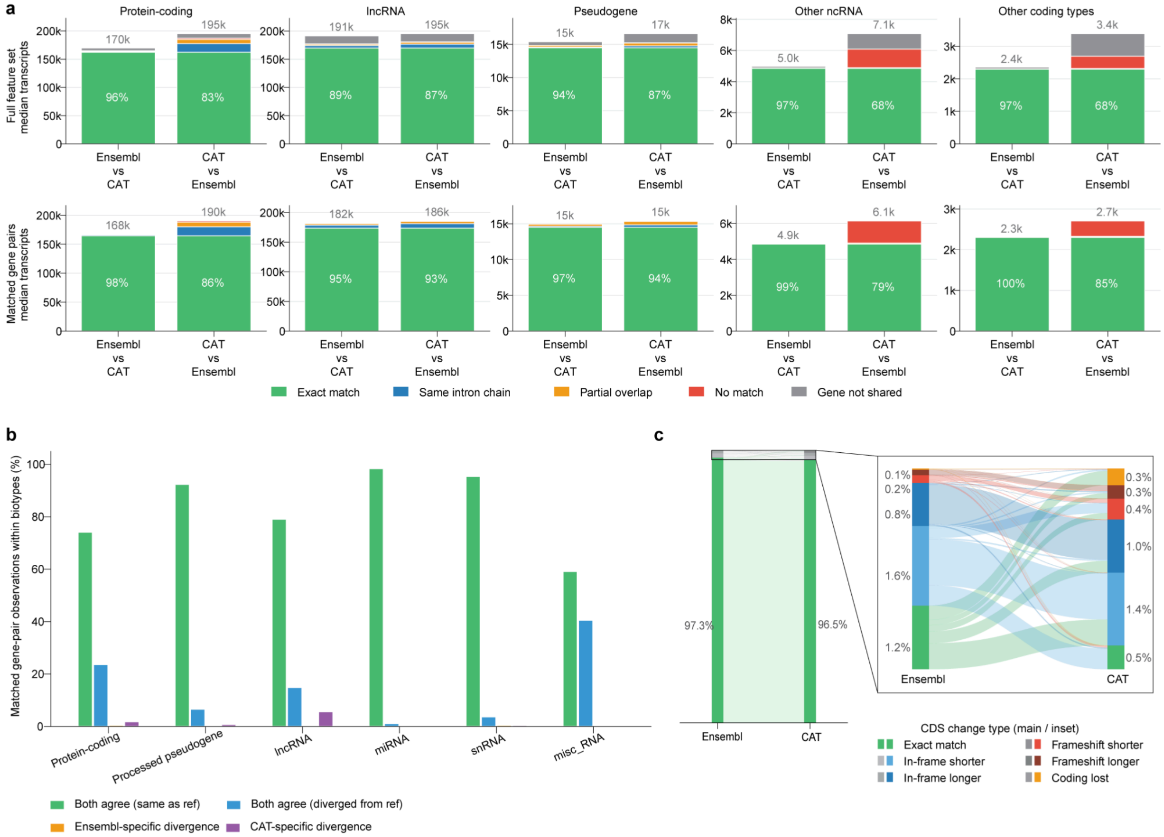


**Supplementary Fig. 28 PANTX1**. Concordance between Ensembl and CAT annotations. **(a)** Transcript-structure concordance within five biotype groups. Bars show median transcripts per assembly, partitioned by match class. Top row: full annotation sets. Bottom row: matched gene pairs after excluding CAT Kinnex models and CAT models originating from GENCODE v47 readthrough genes. Labels give the combined exact-match plus same-intron-chain percentage and median transcript count. Same intron chain: shared splice junctions, differing termini. **(b)** Reference divergence relative to GENCODE v47, by biotype. For each matched reciprocal best-hit gene pair, Ensembl and CAT representative transcripts were compared with the GENCODE v47 reference and classified as both agreeing, both diverging, Ensembl-specific divergence, or CAT-specific divergence. Within-biotype percentages. **(c)** CDS concordance with GENCODE v47 for protein-coding reciprocal best-hit pairs. Main bars: exact CDS match (97.3% Ensembl, 96.5% CAT). Inset: non-exact cases by change type. In-frame: length change divisible by three with preserved reading frame. Inset percentages are of all protein-coding pairs.

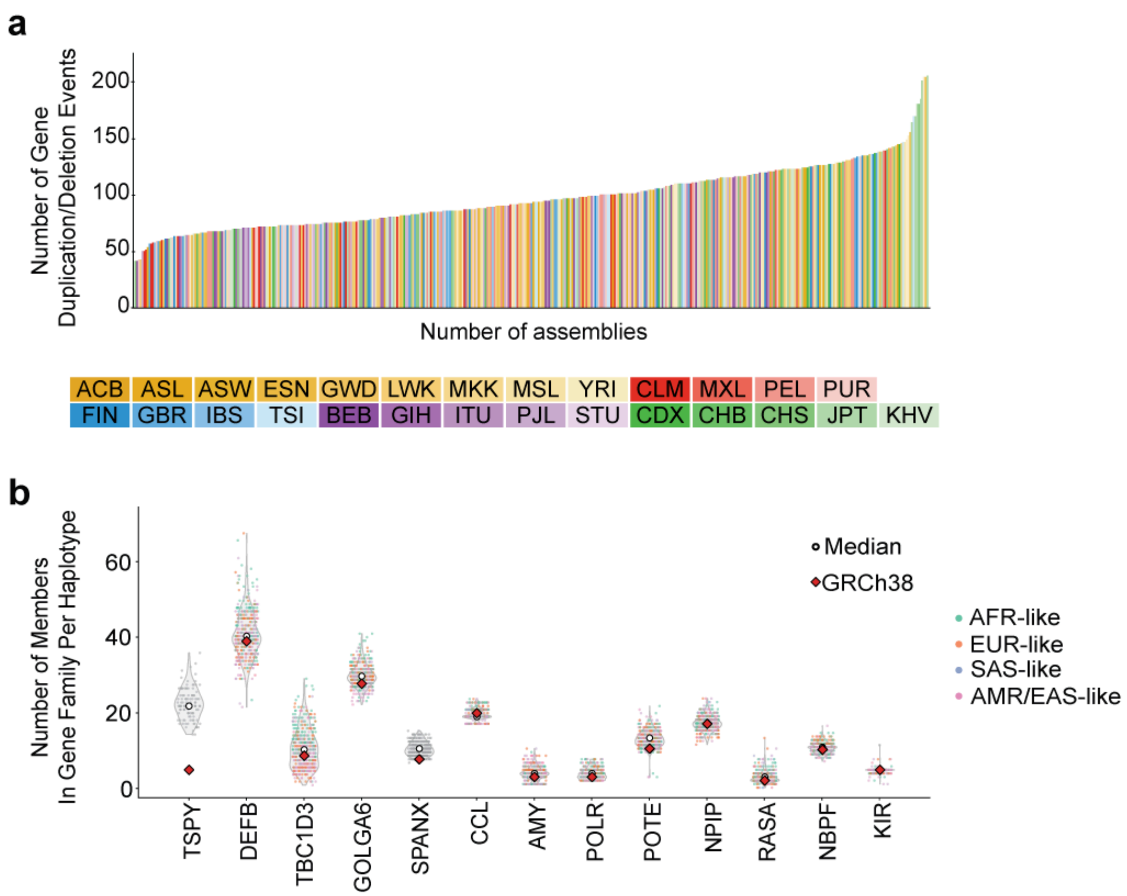


**Supplementary Fig. 29 PANTX2**. Gene CNV Analyses with Ensembl Annotations
(**a**) CNV events across each individual haplotype assembly coloured by population labels. For each gene family (see methods for definition), a deviation from the median count was considered as a variation and the summation of these variants (including both duplications and deletions) across all families for a haplotype is shown here. We are using a qualitative colour map to colour haplotypes where each colour family (yellow, red, purple, green, and blue) represent individual superpopulations and each shade within a colour family represents its subpopulation. (**b**) Counts of individual genes across each of the haplotype assemblies for the most copy-number variable gene families. Each assembly is coloured by the PLCAI derived local ancestry for the locus of the gene family.

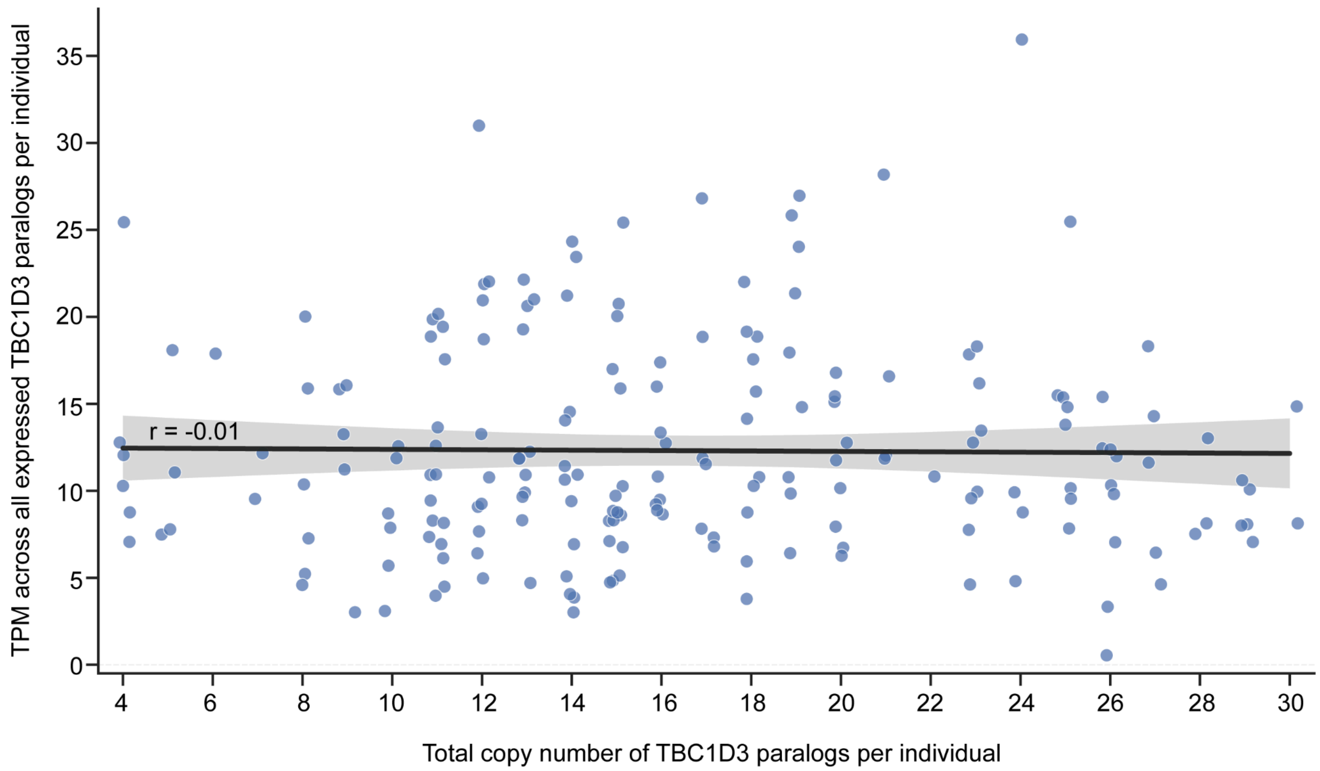


**Supplementary Fig. 30 PANTX3**. Total *TBC1D3* copy number (summed across both haplotypes) versus TPM (sum of all *TBC1D3* paralogs) from matched Kinnex samples per HPRC individual (n = 204). This shows no relationship between genomic copy number and expression (Pearson r ≈ −0.01) which is expected as only 1-2 paralogs are expressed.

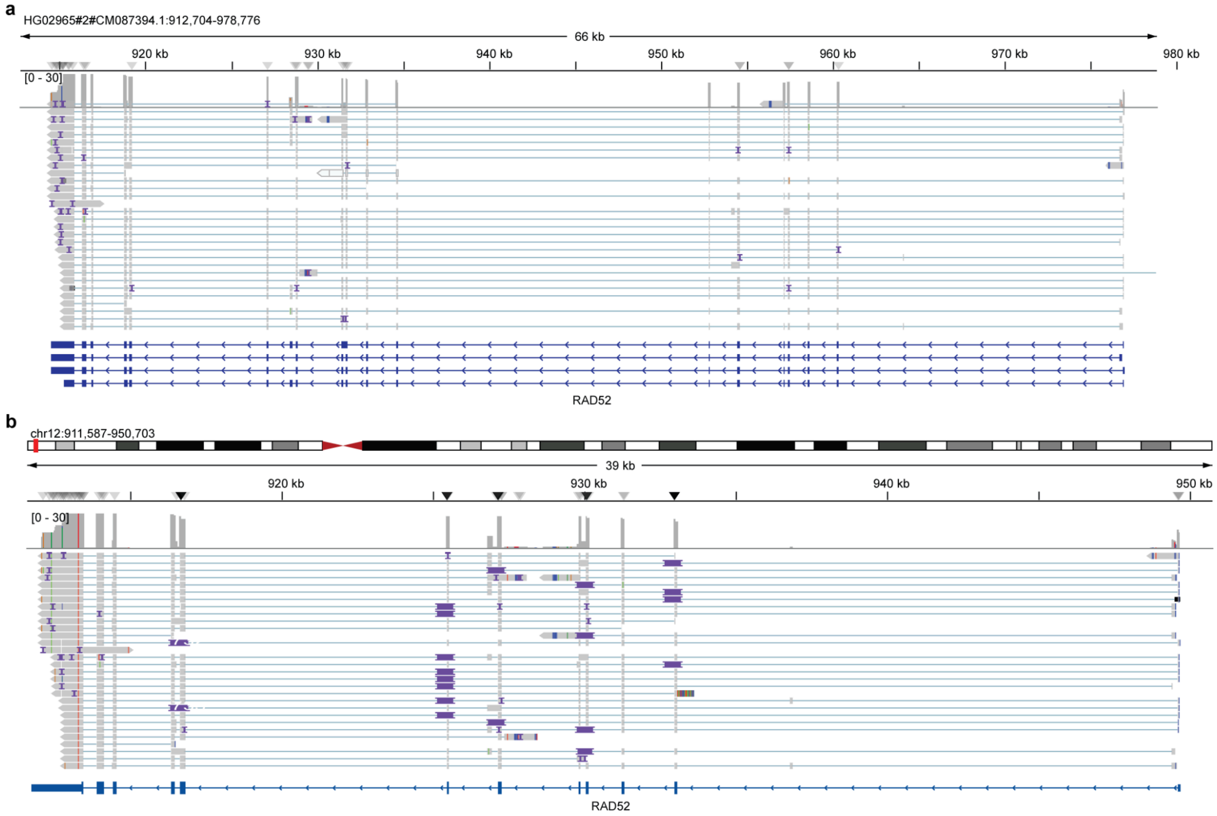


**Supplementary Fig. 31 PANTX4**. IGV snapshot showing read support at the RAD52 locus for HG02965_HAP2 and the insertion in HG38 when mapping reads from HG02965 to HG38.

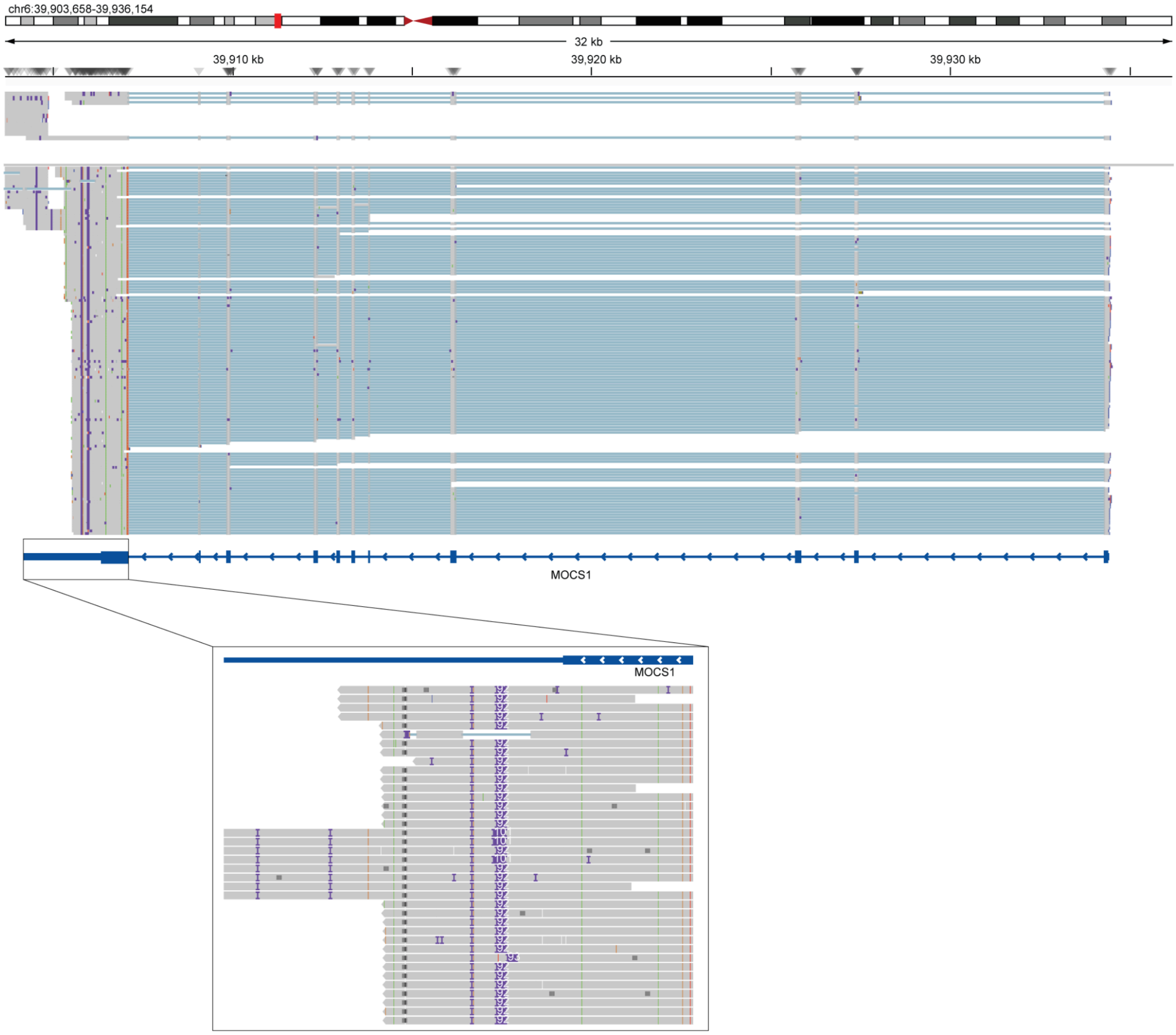


**Supplementary Fig. 32 PANTX5**. IGV snapshot showing the insertion in the 3' UTR in the MOCS1 locus in HG38 when reads from an individual that is HET ALT for the 3' UTR insertion are aligned.

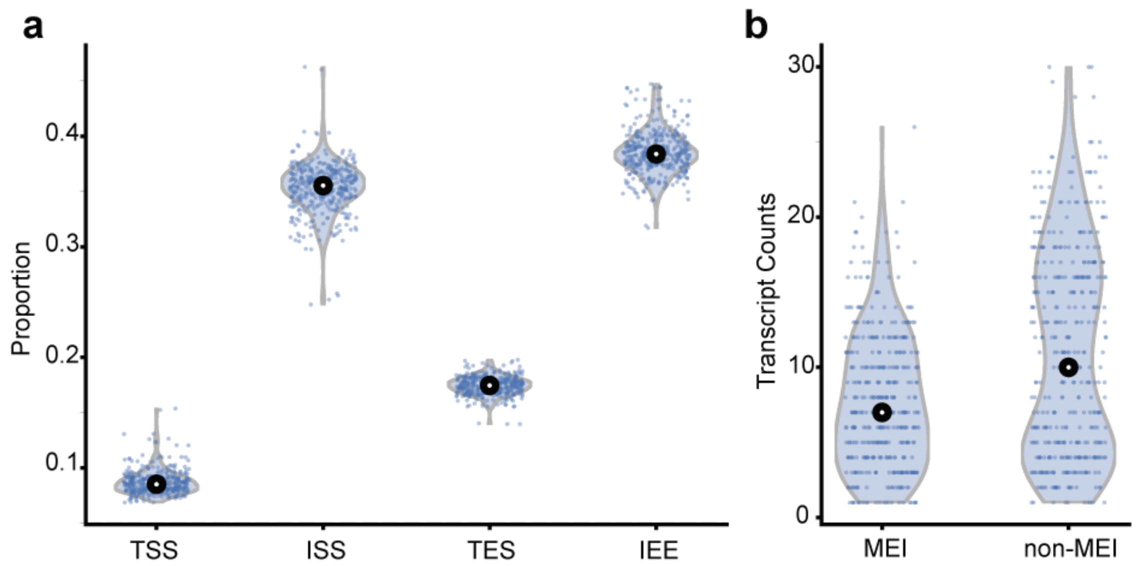


**Supplementary Fig. 33 PANTX6**. Landscape of TE-associated transcripts across individuals **(a)** Proportion of transcripts per individual by TE role (roles: TSS, ISS, TES, IEE; TSS, TE-derived transcription start site; ISS, TE-derived splice site; TES, TE-derived transcript end site; IEE, isoforms containing TE fragments that do not contribute to TSS, ISS or TES. one role per transcript using priority TSS >>ISS >> TES >> IEE). **(b)** Number of non-reference TE-associated transcripts per individual. MEI: recent/active mobile-element insertions with canonical hallmarks (e.g., poly(A) tail, target site duplication); non-MEI = non-reference TE sequences without MEI hallmarks, interpreted as not recent/inactive (e.g., older TE fragments or other rearrangements).

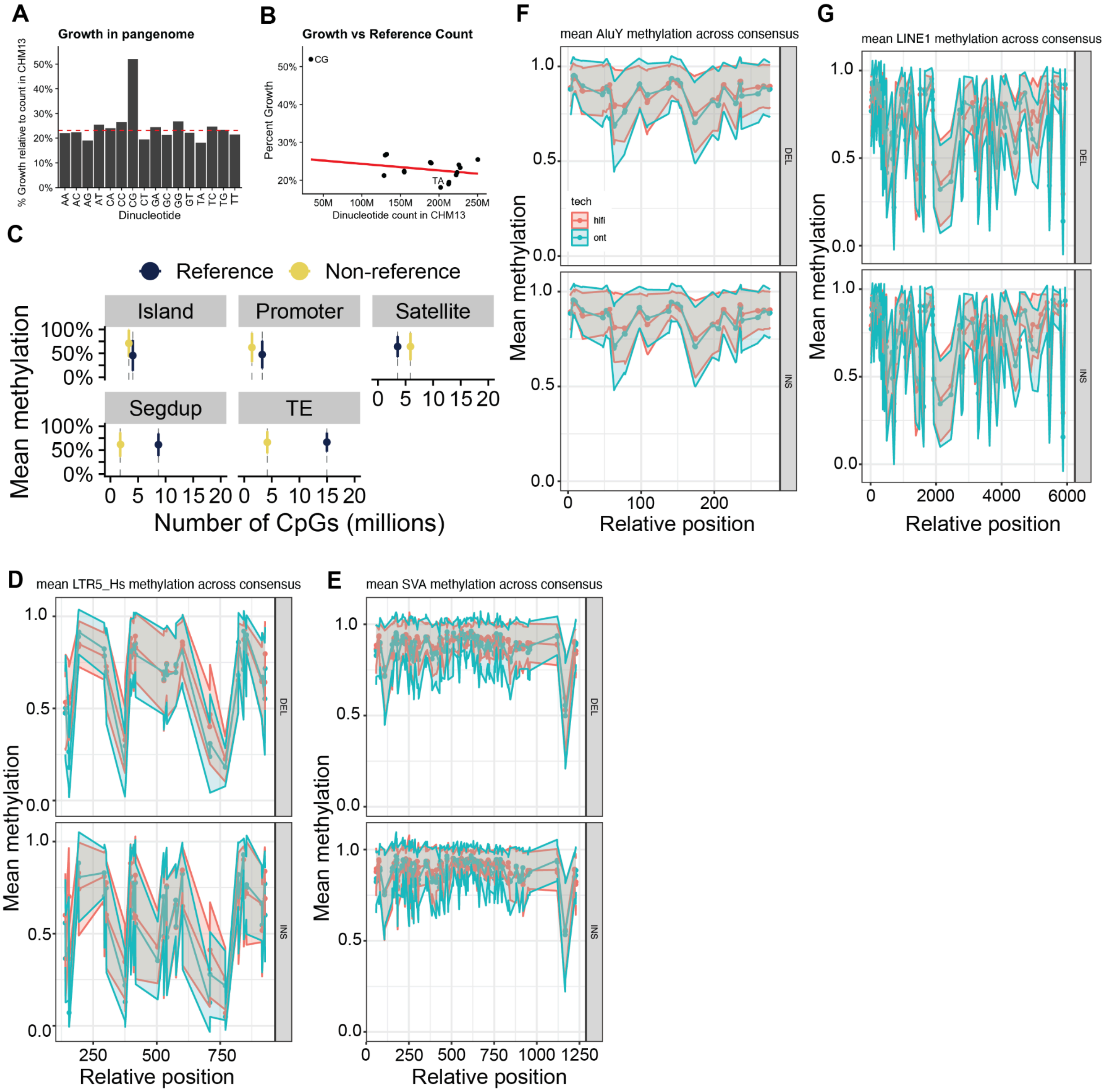


**Supplementary Fig. 34 Panepigenome1. Growth of CpG sequence space and methylation profiling over non-reference CpGs. (a)** Percent growth of the HPRC2 pangenome over T2T-CHM13 for each of the 16 possible dinucleotides. CpG (CG) is the outlier at ~52% relative to the ~22% mean across the remaining 15 dinucleotides (red dashed line). **(b)** Percent growth versus reference dinucleotide count across the 16 dinucleotides. CpG deviates from the linear relationship across the other 15 dinucleotides (red line, ordinary least-squares regression fit on the 15 non-CG points); TA labeled for reference at the lowest growth rate. **(c)** Mean CpG methylation (standard deviation error bars) of reference and non-reference CpGs stratified by genomic context. **(d-g)** Mean CpG methylation across the consensus sequences of four polymorphic transposable element (TE) families, comparing reference TEs (DEL; present in T2T-CHM13 and polymorphically deleted in some HPRC2 haplotypes) and non-reference TEs (INS; absent from T2T-CHM13 and polymorphically inserted in some HPRC2 haplotypes), holding polymorphism status constant. HiFi (red) and ONT (teal) sequencing platform-derived methylation are overlaid; lines show per-position mean methylation across MEI instances and shaded bands indicate per-position standard deviation.**(c)** AluY consensus. Mean ± SD methylation: HiFi 0.87 ± 0.05 (DEL), 0.88 ± 0.05 (INS); ONT 0.86 ± 0.07 (DEL), 0.87 ± 0.07 (INS). **(d)** L1HS consensus. Mean ± SD: HiFi 0.75 ± 0.17 (DEL), 0.77 ± 0.17 (INS); ONT 0.73 ± 0.19 (DEL), 0.75 ± 0.19 (INS). **(f)** LTR5_Hs consensus. Mean ± SD: HiFi 0.69 ± 0.17 (DEL), 0.61 ± 0.20 (INS); ONT 0.66 ± 0.22 (DEL), 0.58 ± 0.24 (INS). **(g)** SVA_F consensus. Mean ± SD: HiFi 0.90 ± 0.05 (DEL), 0.90 ± 0.06 (INS); ONT 0.88 ± 0.06 (DEL), 0.88 ± 0.06 (INS). Despite these four classes of elements being hypermethylated in general, each class has distinct patterns of CpG methylation. For example, the Internal Ribosome Entry Site (IRES) within L1HS and the canonical retroviral transcriptional termination site within LTR5 both appear to be hypomethylated relative to their surrounding elements.

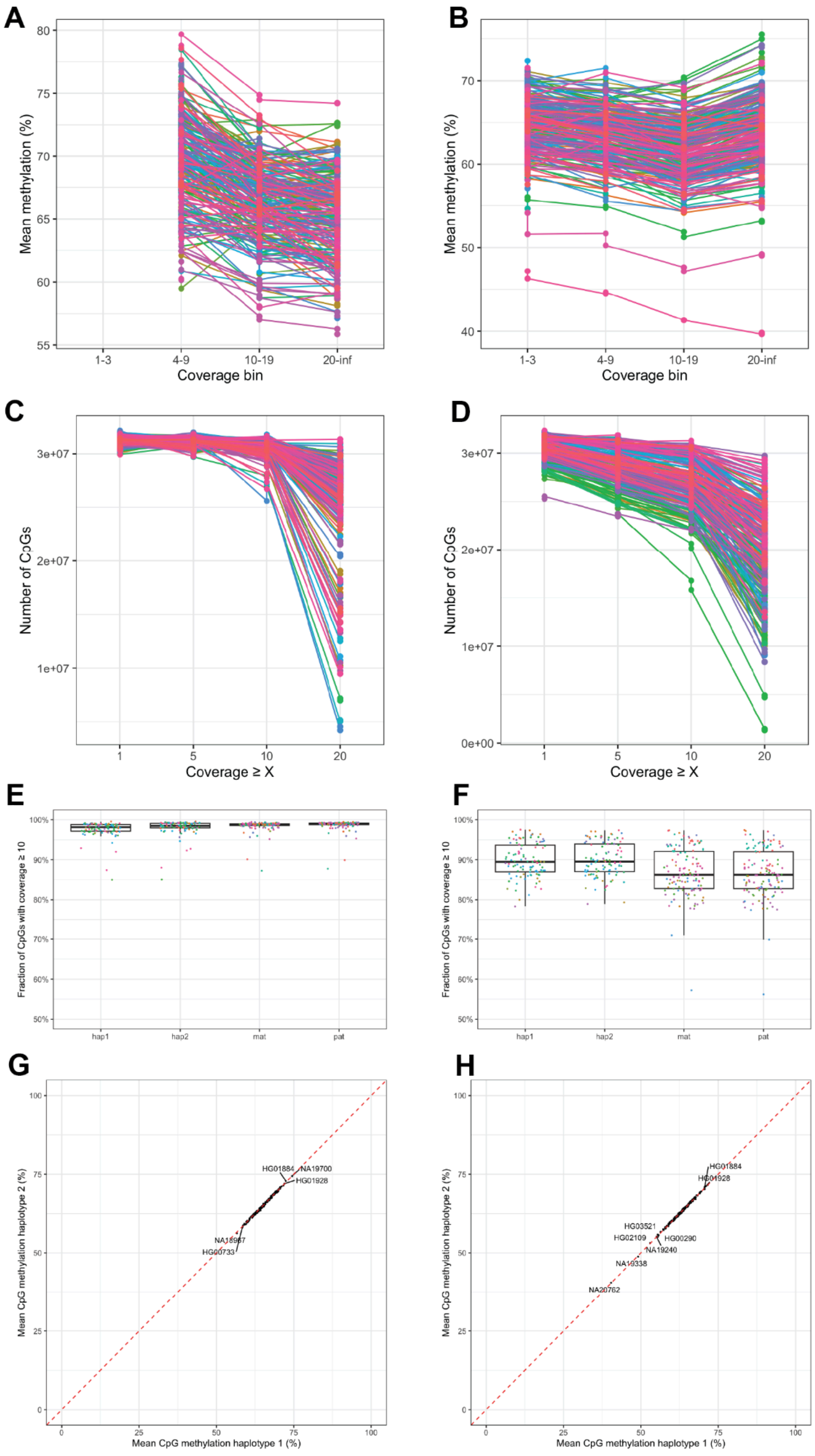


**Supplementary Fig. 35 Panepigenome2. Per-sample methylation QC across PacBio and Oxford Nanopore.** Per-sample methylation QC across 440 haplotypes, with PacBio HiFi (left column) and Oxford Nanopore (right column) shown in parallel. (**A, B**) Per-sample mean methylation across coverage bins (1–3, 4–9, 10–19, ≥20). (**C, D**) Number of CpGs retained per sample as a function of coverage threshold (≥1, 5, 10, 20). (**E, F**) Fraction of CpGs at ≥10× coverage by haplotype assignment (hap1, hap2, mat, pat). (**G, H**) Hap1 vs hap2 mean CpG methylation per donor; dashed line is identity. Per-sample QC profiles were qualitatively similar across both platforms. Median CpG completeness at ≥10× was ≈98% per haplotype on PacBio and ≈89% on ONT, the lower ONT figure consistent with the lower mean ONT coverage (≈20.8× vs ≈25× on PacBio). Coverage-dependent methylation drift was observed on both platforms, declining by approximately ten percentage points between the lowest- and highest-coverage strata. Paired hap1/hap2 mean methylation tracked the identity line tightly on both platforms.

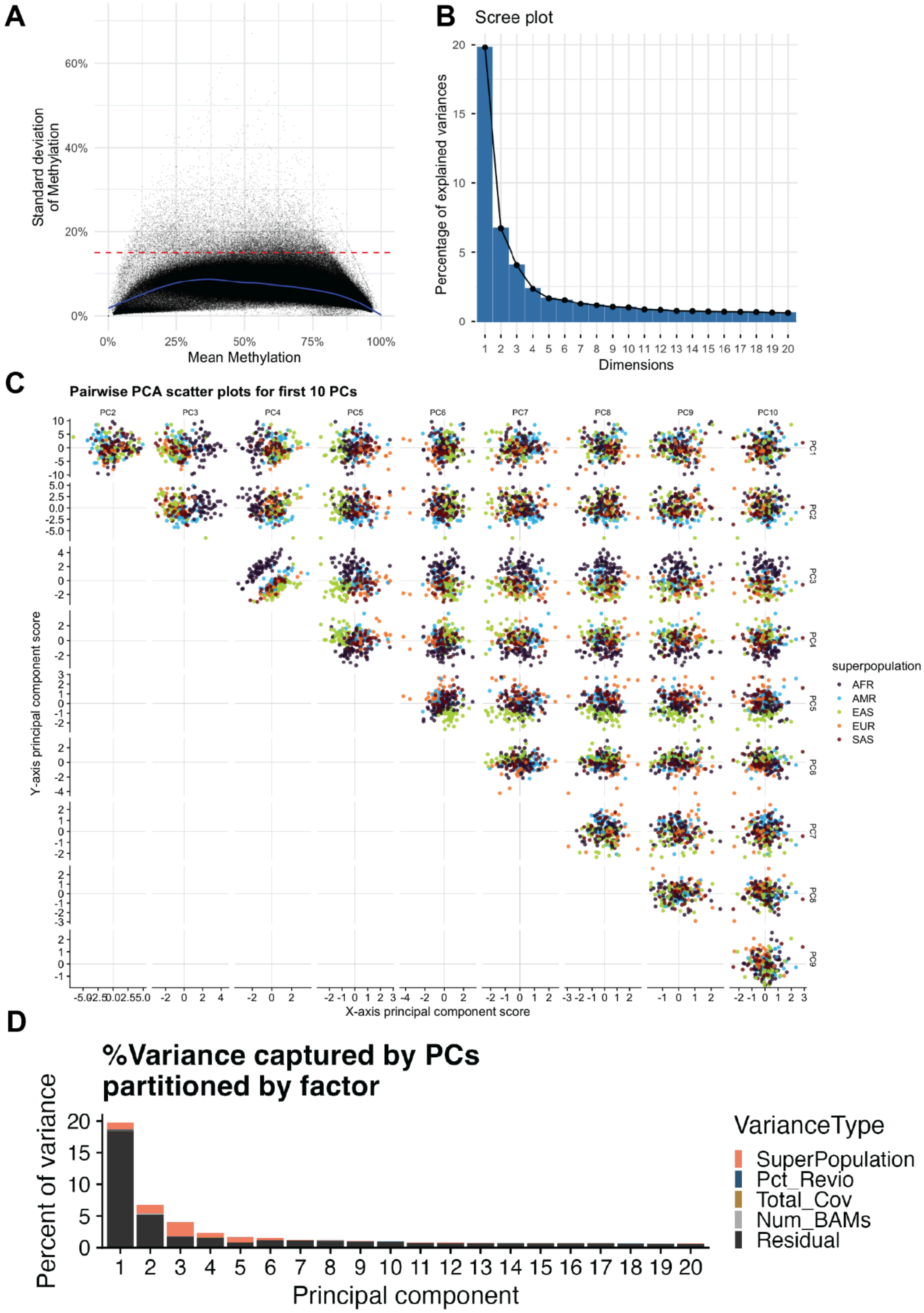


**Supplementary Fig. 36 Panepigenome3. Principal components analysis of 2-kb binned PacBio methylation.** Principal components analysis of 2-kb binned PacBio methylation across HPRC2 haplotypes. (**A**) Feature mean methylation versus standard deviation, with the SD > 0.15 threshold (red dashed line) used to retain variable features for PCA input. (**B**) Scree plot of variance explained per principal component. (**C**) Pairwise PC scatter for the first 10 PCs, colored by 1000 Genomes superpopulation. (**D**) For each principal component, the variance captured by that PC is partitioned into the fraction attributable to superpopulation, technical covariates (percent Revio, total coverage, number of BAMs per sample), and unexplained residual. PC1 captured the largest share of total methylome variance (19.8% of all variance in the data), but only 6% of PC1's own variance was attributable to superpopulation, with the remainder largely unexplained; on PC2 (which captured 6.7% of all variance) superpopulation explained 20% of PC2's variance; on PC3 (which captured 4.1% of all variance) the superpopulation share peaked at 56%. Summed across the first 20 PCs (which together capture 47.7% of all methylome variance), superpopulation accounted for 7.4% of all methylome variance, technical covariates 0.6%, and unexplained residual 39.6%.

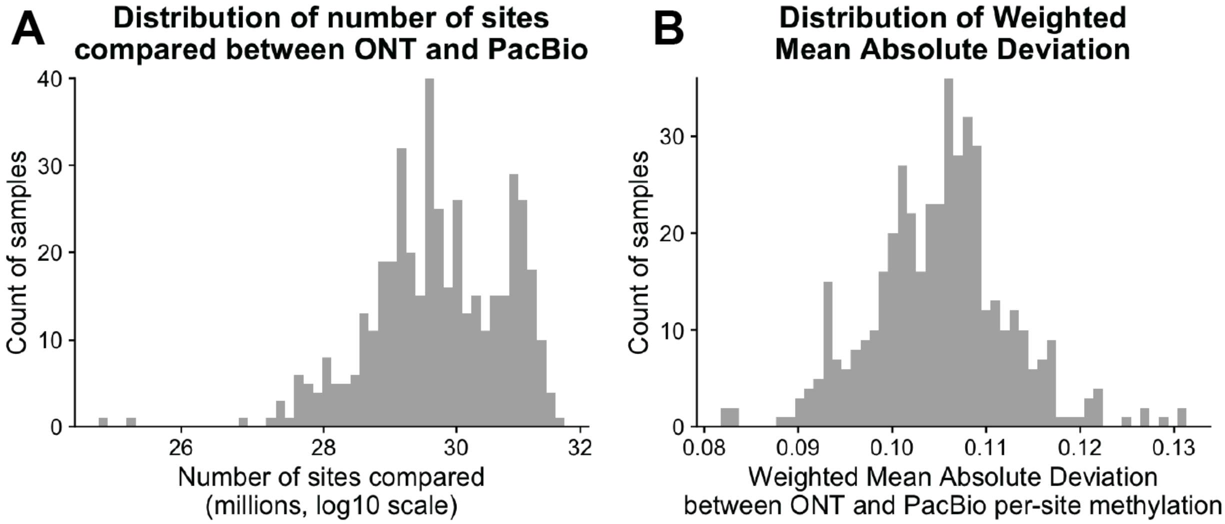


**Supplementary Fig. 37 Panepigenome4. PacBio vs Oxford Nanopore methylation concordance.** Per-CpG methylation fractions were compared at each shared site between PacBio and ONT. (**A**) Distribution of the number of shared CpG sites compared per sample. (**B**) Distribution of coverage-weighted mean absolute deviation per sample. Across ≈29–30 million shared CpGs per sample: coverage-weighted mean absolute deviation 0.100 ± 0.017; ordinary least-squares regression of PacBio on ONT, slope 0.883 ± 0.052, intercept 0.099 ± 0.036; $R^2$ 0.803 ± 0.052. The site-wise discrepancy between platforms is expected to be dominated by random error. Because our downstream analyses operate on binned methylation, we re-evaluated concordance at 1-kb bin resolution (requiring ≥10 reads per platform and ≥10 overlapping CpGs per bin); 933,724 bins per sample were retained and bin-level mean delta methylation was 1.55% (SD 1.59%). Binning averages out the random per-site error and renders the two platforms functionally equivalent for our downstream analyses. We use PacBio as the canonical call stream because it produces substantially more uniform coverage across CpGs and lower per-donor variability in CpG capture rate; ONT is retained as an orthogonal validation.

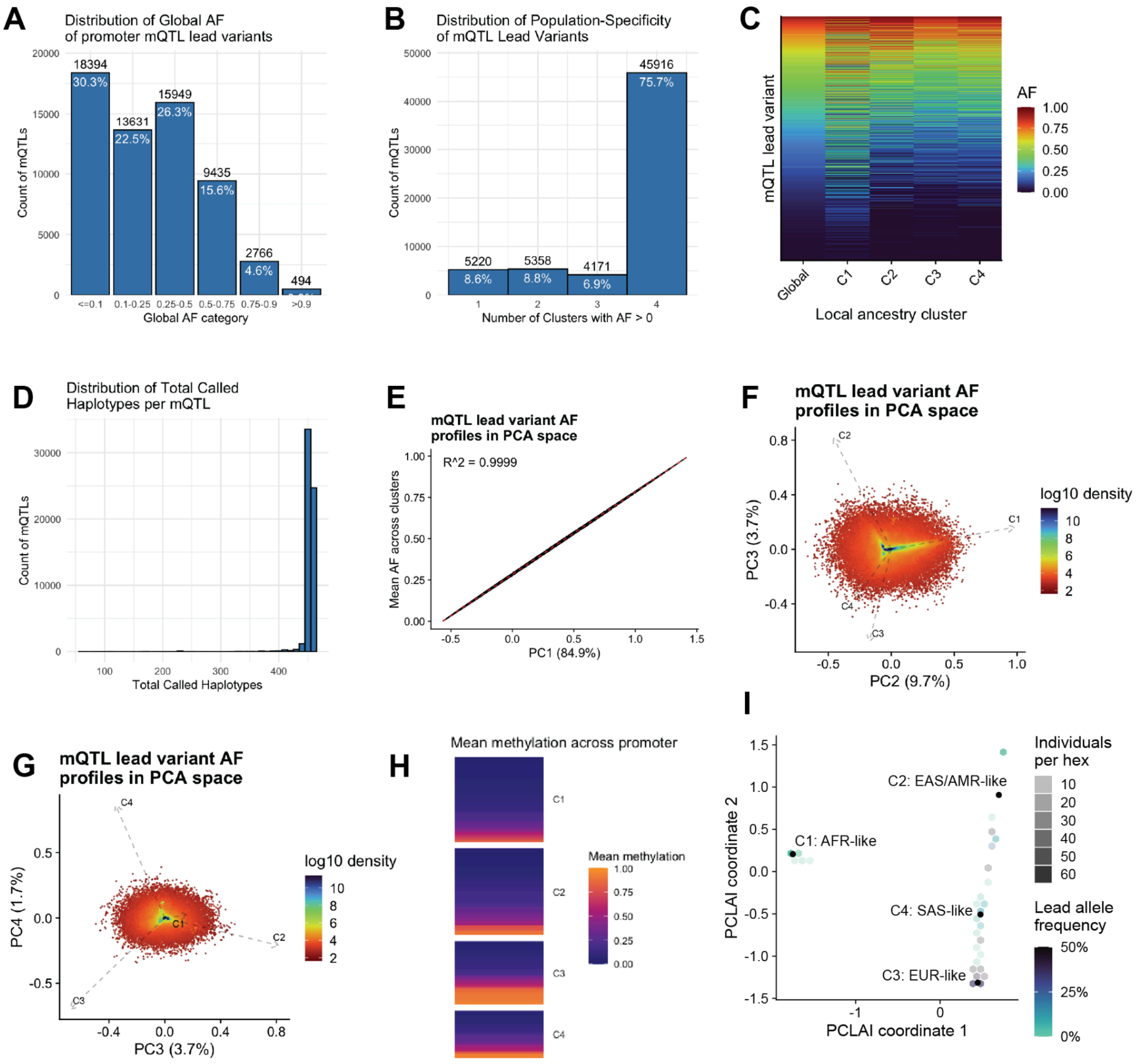


**Supplementary Fig. 38 Panepigenome5. Cluster-stratified allele frequencies of promoter mQTL lead variants.** Per-cluster allele frequencies for 60,669 biallelic promoter mQTL lead variants in the HPRC R2 panel, stratified by haplotype-resolved PCLAI local ancestry. (**A**) Distribution of global AF (per-variant mean of cluster-specific AFs) across biallelic mQTL lead variants, binned into six categories. The panel set is weighted toward rare and intermediate-frequency variants: 30.3% have global AF ≤ 0.1, 22.5% are 0.1-0.25, 26.3% are 0.25-0.5, and 15.6% are 0.5-0.75, with the remainder above 0.75. (**B**) Population-specificity of lead variants, defined as the number of PCLAI clusters with AF > 0. Most lead variants are broadly shared across clusters: 45,916 (75.7%) were observed in all four clusters, 4,171 (6.9%) in three, 5,358 (8.8%) in two, and 5,220 (8.6%) in only one. The pattern is consistent with what is expected under panel sampling, where common variants tend to be observed in all clusters and rare variants are more likely to be sampled in only a subset. (**C**) Per-variant allele-frequency profile heatmap. Rows are biallelic mQTL lead variants and columns are the global panel-average AF and the four per-cluster AFs; color indicates alternate allele frequency. Most rows track the global AF column tightly across the four cluster columns, but a subset shows visible cluster skew. Quantitatively, the fraction of lead variants with AF = 0 in a given cluster ranged from 4.97% in C1 (AFR-like) to 18.10% in C4 (SAS-like), with C2 (AMR/EAS-like, 16.09%) and C3 (EUR-like, 11.23%), while the fraction with AF > 0.5 was highest in C1 (AFR-like, 25.12%). (**D**) Distribution of total called haplotypes per mQTL across the panel. Most mQTLs are called in most of the available haplotypes, confirming that per-cluster AF estimates are based on near-complete panel sampling at each tested position. (**E**) PCA of cluster-stratified AF profiles, PC1 versus mean AF across clusters. PCA was performed on the four-column matrix of per-cluster AFs using prcomp (center = TRUE, scale. = FALSE). PC1 explained 84.9% of the variance and was almost perfectly correlated with mean AF across clusters ($R^2 = 0.9999$), so PC1 reflects how common a lead variant is in the panel rather than how it is distributed across clusters. (**F**) PCA in PC2 versus PC3 space (9.7% and 3.7% of variance, respectively), with cluster loading vectors overlaid and point density colored on log10 scale. (**G**) PCA in PC3 versus PC4 space (3.7% and 1.7% of variance). The squared loading of a cluster on a PC indicates the fraction of that PC's variance contributed by that cluster's allele frequencies. PC2 is dominated by C1 (squared loading 0.75); PC3 contrasts C2 (0.52) with C3 (0.36); PC4 contrasts C4 (0.58) with C3 (0.38). (**H**) Per-haplotype mean methylation across the PM20D1 promoter, stratified by PCLAI cluster, shown as an example connecting cluster-stratified mQTL lead-variant AF to cluster-stratified methylation phenotype distributions. Each row corresponds to a haplotype. C3 (EUR-like) shows a pronounced bimodal distribution with substantial fractions of both hypomethylated and hypermethylated haplotypes; C1, C2, and C4 are predominantly hypomethylated. The cluster pattern follows the lead variant AF directly: the PM20D1 promoter mQTL lead allele (rs9438393) is at intermediate frequency in C3 (0.356), so both ref and alt haplotypes are well-represented and the methylation distribution is bimodal, whereas the allele is rare in C1 (0.022), C2 (0.036), and C4 (0.092), so those clusters are dominated by ref haplotypes and a single methylation mode. The lead allele is approximately 16-fold more common in C3 than in C1. PCLAI clusters remain coarse partitions of a continuous local ancestry space and panel sampling across clusters is unbalanced, so per-cluster frequencies are panel-conditional rather than external population estimates.(**I**) PM20D1 lead variant (rs9438393) allele frequency across continuous PCLAI local-ancestry space. Individuals are aggregated into hexbins by their PCLAI coordinates; hex shading indicates the number of individuals per hex and hex color indicates the lead allele frequency, with the four discretized cluster centroids (C1, AFR-like; C2, AMR/EAS-like; C3, EUR-like; C4, SAS-like) annotated for reference.

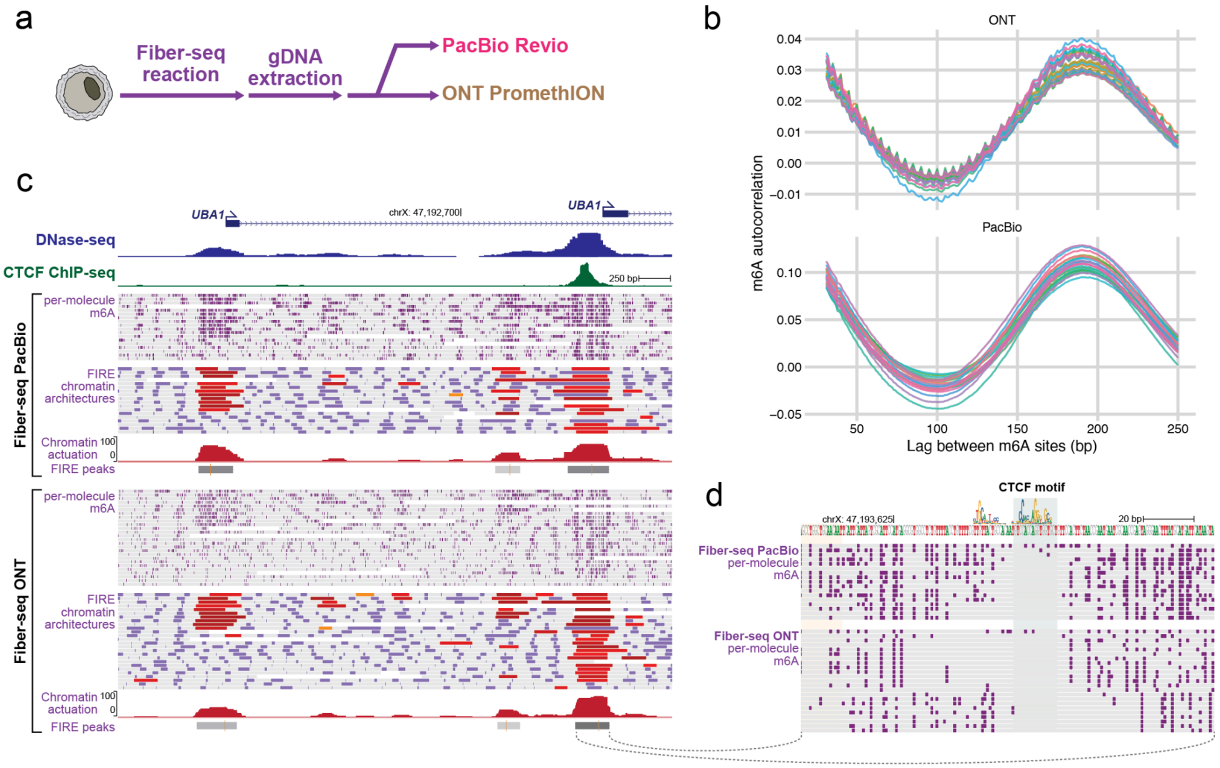


**Supplementary Fig. 39 Panepigenome6. Fiber-seq across HPRC lymphoblastoid samples.** **a)** Schematic of the experimental design. Genomic DNA from each lymphoblastoid sample was treated with the Fiber-seq m6A stenciling reaction and then sequenced on PacBio Revio, ONT PromethION, or both platforms. **b)** Per-read m6A position autocorrelation for each sample, with panels split by sequencing platform. Both PacBio and ONT recover the canonical nucleosome periodicity. **c)** Genome browser view at the UBA1 locus (chrX) with both Fiber-seq platforms shown. Tracks from top to bottom: ENCODE DNase-seq and CTCF ChIP-seq for reference, followed by PacBio Fiber-seq (per-molecule m6A, FIRE chromatin architectures with actuated FIRE elements in red, aggregate chromatin actuation, and FIRE peaks) and the matched tracks from ONT Fiber-seq. **d)** Zoom in on a CTCF binding site within the UBA1 locus (dashed line connects to the corresponding FIRE peak in panel c). The CTCF motif is shown above per-molecule m6A patterns from PacBio (top) and ONT (bottom) Fiber-seq. Both platforms resolve the protected CTCF footprint, although ONT shows lower m6A density because simplex ONT captures adenine methylation along only one strand.

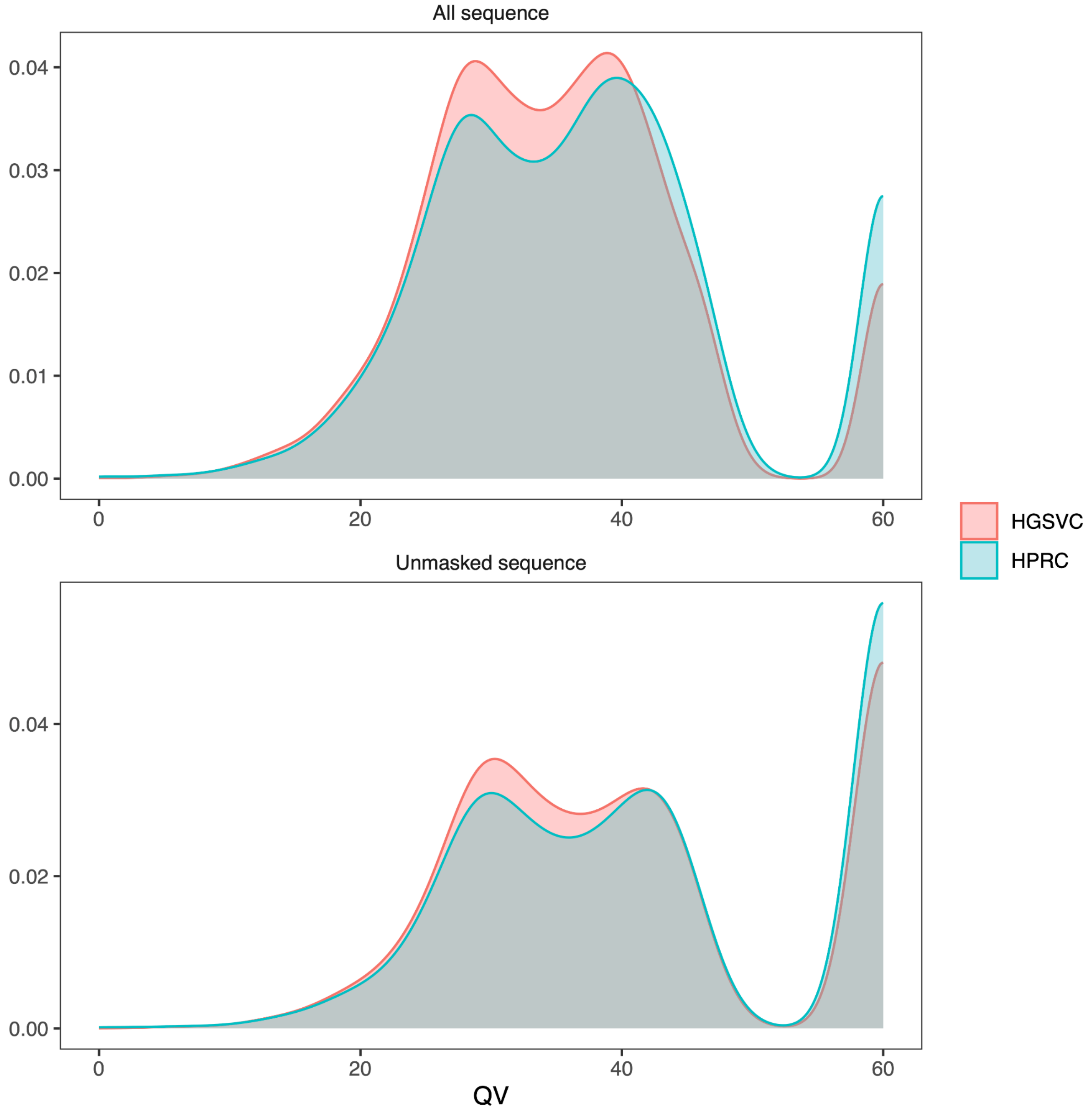


**Supplementary Fig. 40 ctyper_fig QV distributions from ctyper genotyping results on challenging medically relevant genes for the HGSVC and HPRC reference panels**. Three short-read samples were genotyped via ctyper using a leave-one-out strategy, where each sample's ground-truth genome assembly was excluded from the reference panel prior to genotyping. Genotyped sequences were aligned to their corresponding ground-truth diploid genome assemblies, with each genotyped sequence paired to its most likely source locus. Per-allele QV was calculated as $-10 \times \log_{10}$(mismatched bases / total aligned length), with zero-mismatch alleles capped at QV = 60. The top panel shows QV across the full aligned sequence; the bottom panel shows QV restricted to non–repeat-masked regions. Curves represent the density of per-allele QV values across all genotypes in each reference panel. Higher QV reflects lower mismatch rates and greater concordance with the ground-truth assemblies.

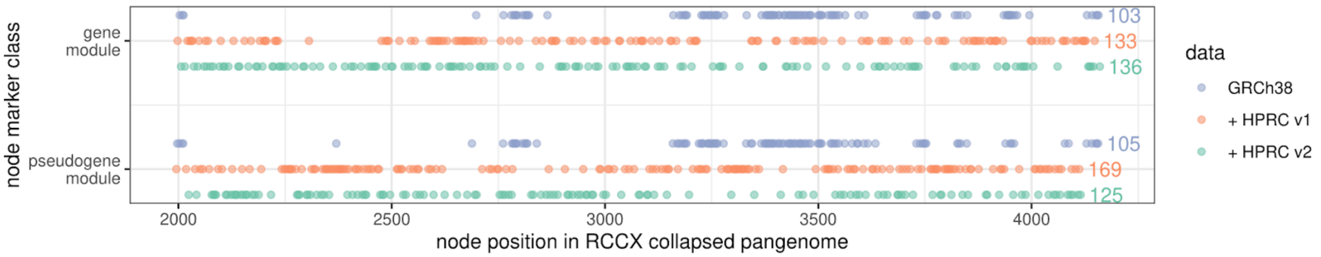


**Supplementary Fig. 41 rccx-markers**: Informative markers, either specific to the gene module (top), or pseudogene module (bottom), position across the pangenome (x-axis). Points in blue represent informative nodes that can be found when collapsing the two reference module in GRCh38. Orange points are the new informative nodes gained by further including the 94 haplotypes from HPRC release 1. Finally, green points were added thanks to the addition of new haplotypes from the HPRC2. Of note, the additional informative nodes brought by HPRC release 1 and 2 also better cover the RCCX modules.

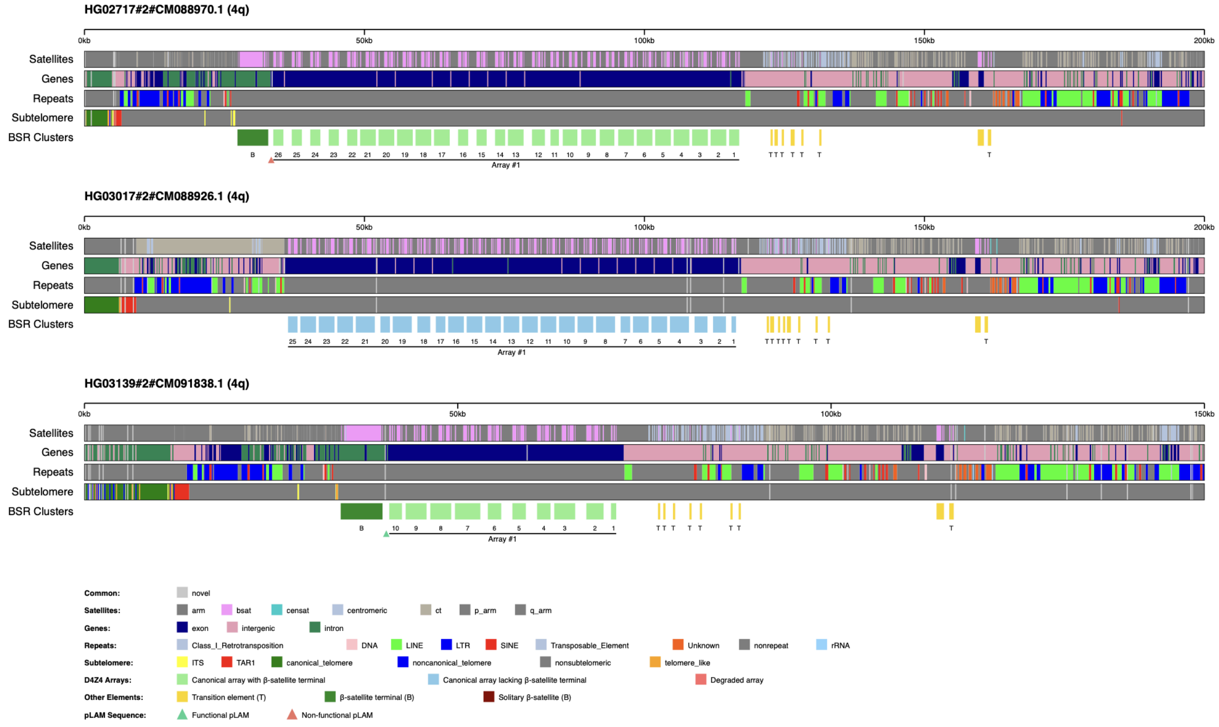


**Supplementary Fig. 42 D4Z4_SUPP**: Representative KaryoScope D4Z4 array visualizations on chr4. Each panel shows the subtelomeric region of a single chr4 haplotype with KaryoScope feature annotations. (a) HG02717 haplotype 2: a 4qA haplotype with 26 D4Z4 repeat units and terminal β-satellite, carrying a non-functional pLAM (ATCAAA). (b) HG03017 haplotype 2: a 4qB haplotype with 25 repeat units and no terminal β-satellite. (c) HG03139 haplotype 2: an FSHD1-compatible 4qA haplotype with 10 repeat units, terminal β-satellite, and a functional pLAM (ATTAAA), placing it at the upper boundary of the disease contraction range (≤10 units).

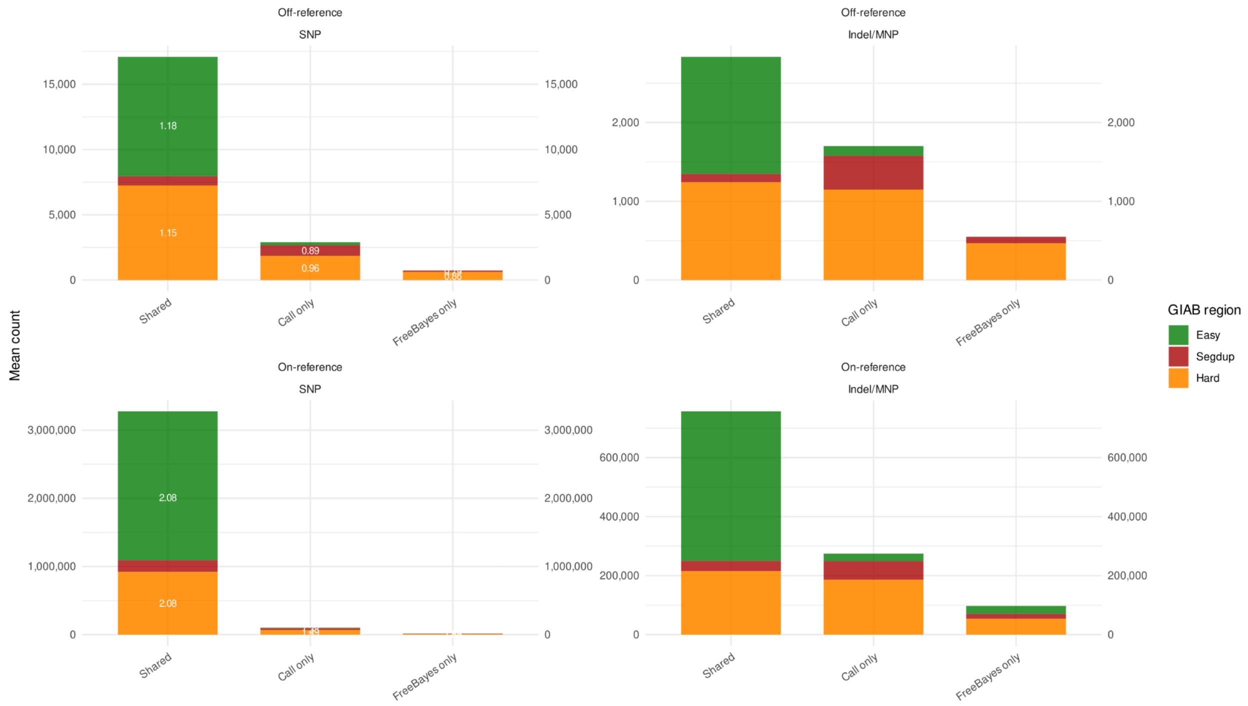


**Supplementary Fig. 43 gref_freebayes_genotyping**: Mean variant counts from 9 short read samples as genotyped by FreeBayes and vg call, stratified by (non-overlapping) GIAB regions on T2T-CHM13. Only variants in the graph are considered.

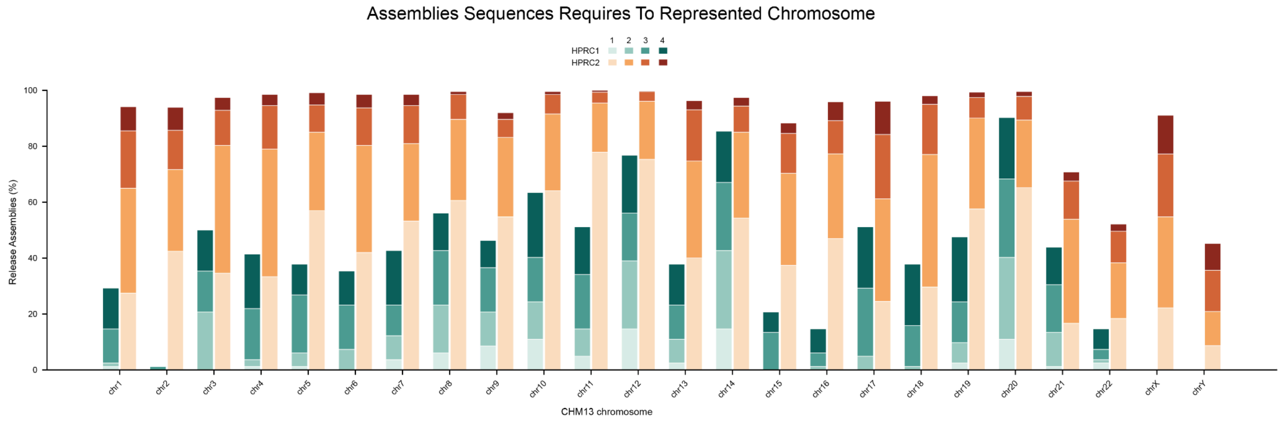


**Supplementary Fig. 44 Sequence_counts_per_chrom. HPRC2 assemblies cover CHM13 chromosomes in under 4 assembled sequences across most of the genome.** Each colored section of the bar represents the percent of the release's assemblies that represent the chromosome in that many sequences. The estimate of the number of assembly sequences are based on unique projected coverage, and in the case of HPRC2 scaffold gaps add to the estimate (see methods).
