## Supplementary Notes for "HPRC2: A human pangenome reference with near-complete coverage of common genetic variation"

### Supplemental Note 1 - Telomere Length Analysis

Conventional telomere length assays including Southern blotting, qPCR and FlowFISH, report on the mean length of all telomeres in the cell[1–3]. Long-read nanopore (ONT) sequencing now enables base pair resolution of telomere length at each individual chromosome end, providing greater accuracy[4], and enabling length determination from publicly available data sets. However, many long-read data sets rely on Epstein-Barr virus (EBV) immortalized lymphoblastoid cell lines (LCLs) as their primary data. Transformation of B cells with EBV has been shown to alter telomere lengths, as the telomeres first shorten as cells divide pre-imortalization and then are elongated after immortalization. Many LCLs are 'preimmortal' as immortalization occurs only after about 150 population doublings[5–7]. Recent work documented significant differences in telomere length estimates from T2T genome assemblies comparing blood DNA and LCL DNA from the same individual[8]. In light of this finding, we characterized telomere lengths derived from LCLs across the 232 samples comprising HPRC Release 2.

Telomere Profiling of peripheral blood mononuclear cells (PBMCs) from 103 healthy individuals aged 18-90 established a reference telomere length distribution with a 2.5th-97.5th percentile range of 3.8-6.7 kb[4]. We examined telomere length in the HPRC R2 LCL samples by first filtering sequencing reads to retain telomeric sequences, then applying the TeloNP algorithm to estimate telomere length. Whole genome BAM files were downloaded from AWS. BAM file were then filtered by a telomere k-mer screen, where we searched for 100 telomere repeats (TTAGGG/CCCTAA) within a hamming edit distance of 1, that were required to be within the first or last 1000 bp of a read. Individual read telomere lengths were quantified with the TeloNP algorithm, which defines telomere boundaries based on discontinuities in telomeric sequences G/C content (https://github.com/GreiderLab/TeloBP). The HPRC R2 release integrates data from across the current long-read sequencing landscape. Telomere length estimates from both PacBio and ONT spanned a striking range of ~2.5 to 20 kb (**Supplementary Fig. 10 telomere_lengths_in_lcls**). We found PacBio telomere length estimates were significantly shorter than ONT and given prior evidence that the PacBio platform artificially limits telomere length estimates[9–11], we used ONT data to estimate telomere lengths. We found that ~49% of LCL samples fell outside the 2.5th-97.5th percentile range of the previously published PBMC telomere length (**Supplementary Fig. 10 telomere_lengths_in_lcls**). Given these irregularities, we caution that LCL-derived telomere length estimates from the HPRC R2 dataset should not be used to draw inferences about primary-cell telomere length.

### Supplementary Note 2 - Description of DNA methylation data

This note characterizes the methylation data underpinning the HPRC2 panepigenome analyses, summarizing per-sample distributional properties, cross-platform concordance, and binned-representation principal components analyses. The methylation calling pipelines for PacBio HiFi (Primrose harmonization, pb-CpG-tools) and the methylation likelihood QC chain (1,158 of 1,092 BAMs; 220 samples; 440 haplotypes; HG00272 outlier exclusion) are described in the main Methods.

We next sought to identify genetic variation driving sample-specific methylation patterns. PacBio HiFi reads were aligned to T2T-CHM13 and methylation was called in unphased reference coordinates for this analysis. Per-CpG PacBio methylation was aggregated into fixed-width 2-kb windows against the T2T-CHM13 reference. Per-bin methylation was the coverage-weighted mean of CpG values; bins lacking coverage were dropped. A standard-deviation filter (SD > 0.15) retained the variable subset of features. Per-haplotype matrices (2,002 intervals; 221 haplotypes × features, after chrX-bin removal and HG00272 outlier exclusion) were dimensionality-reduced with prcomp (center = TRUE, scale. = FALSE). To attribute the variance captured by each principal component to explanatory factors, per-PC scores were regressed against candidate covariates (superpopulation, percent Revio, total coverage, number of BAMs per sample) and the variance explained by each covariate was tabulated per PC.

These results are consistent with what we would expect from an unsupervised PCA of genome-wide methylation. Studies that select CpGs near, or correlated with, nearby genetic variants tend to recover genetic ancestry structure among the leading PCs, because the input features are enriched for *cis*-genetic effects[12,13]. Our goal here is different. We sought to characterize how methylation varies across the genome and we restricted the analysis to bins with appreciable methylation variation across the panel so that the principal components reflect the structure of the variable methylome rather than the structure of genetic variation. In this regime, the largest axes are typically driven by tissue and cell-type composition, technical processing, and individual-level non-genetic variation, with ancestry contributing a weaker and more distributed signal[14–16]. The HPRC2 panel is composed entirely of LCLs, which removes the cross-tissue axis that typically dominates such PCAs. The variance partitioning from our PCA reflects this: across PCs 1 through 20 (~47.7% of total methylome variance), 7.4% is attributable to super population and 0.6% to the tested technical covariates, leaving 39.7% of the total methylome variance captured by these PCs but unexplained by the factors we modeled. What drives this residual is not resolvable from this analysis. Super population labels are themselves a coarse proxy for genetic background, capturing variation that sorts along the major continental axes but missing finer ancestry structure, admixture, and individual *cis*-genetic effects. The super population $\eta^2$ should therefore be read as a lower bound on the genetic contribution to methylome variance, with the remainder of the residual likely reflecting a combination of line-specific factors such as EBV transformation and culture history, and environmental exposures.

### Supplementary Note 3 - mQTL lead variant local ancestry

This note characterizes the cluster-stratified allele frequencies of promoter mQTL lead variants across haplotype-resolved PCLAI[17] local ancestry backgrounds in the HPRC R2 panel. Lead variant matching to the HPRC v2.0 minigraph-cactus T2T-CHM13 wave VCF, per-haplotype intersection with PCLAI, cluster discretization, and the allele-frequency heterogeneity testing procedure are described in the main Methods. PCLAI clusters are referenced by their reference-PCA-space centroids and assigned descriptive labels for readability: C1 (African-like, centroid (-1.743, 0.207)), C2 (East Asian-Indigenous American-like, centroid (0.695, 0.907)), C3 (EUR-like, centroid (0.445, -1.314)), and C4 (South Asian-like, centroid (0.477, -0.507)). After matching and intersection, 67,562 promoter mQTL lead variants were retained, of which 60,669 were biallelic and used for the analyses described below (55,034 SNVs, 1,897 small insertions, 2,647 small deletions, 471 small complex variants, 214 structural insertions, 405 structural deletions, and 1 inversion). Cluster-stratified allele-frequency heterogeneity was tested per variant using the Genepop genic differentiation test (struc; 1,000 dememorization, 100 batches, 5,000 iterations) with Benjamini-Hochberg adjustment across all tested mQTLs; [N] of [N_total] biallelic lead variants showed significant heterogeneity at FDR<0.05.

These observations describe the panel-level allele-frequency structure recovered for the mQTL lead variant set. They are not interpreted as evidence that mQTL lead variants are more differentiated across clusters than non-mQTL variants of comparable frequency, and were not benchmarked against a matched non-mQTL null; drift alone is sufficient to produce allele-frequency variation across populations at any genome-wide variant set. The absence of a selection signature does not diminish the biomedical relevance of the structure shown here. Every promoter mQTL lead variant in this set has a locus-resolved effect on methylation, and the methylation phenotype tracks the allele. When the lead allele is more common in one PCLAI cluster than in others, regardless of whether drift, demographic history, or selection is the underlying cause, the methylation phenotype the allele drives is correspondingly more common in that cluster. This has direct downstream consequences for any disease or trait in which the methylation phenotype contributes to gene-regulatory readout, expression of nearby genes, or risk-allele-stratified clinical features.

To formalize the cluster-stratified methylation pattern at PM20D1 (**Supplementary Fig. 39h Panepigenome6 h)**, haplotypes were stratified into three promoter methylation states (hypomethylated, partially methylated, hypermethylated) and a likelihood-ratio G statistic on the methylation-state by PCLAI-cluster contingency table was computed. Significance was evaluated against a sample-blocked randomization null in which cluster labels were permuted across samples while preserving the pairing between haplotypes from the same individual. The PM20D1 distribution differed significantly across PCLAI clusters under this test (empirical P = 0.0099)(**Supplementary Fig. 39i Panepigenome6 i)**. Specifically, 45 of 97 C3 haplotypes were hypomethylated (46.4%) and 34 of 97 were hypermethylated (35.1%), while hypermethylated fractions in the other clusters were 3.1% in C1, 7.8% in C2, and 10.8% in C4. The HPRC Panepigenome Browser companion resource displays the locus in the context of phased donor haplotype assemblies, allowing the haplotype-resolved methylation pattern to be examined alongside donor-specific sequence differences.

The PM20D1 mQTL lead variant falls within ENCODE[18] candidate enhancer EH38E3989962, approximately 36.6 kb from the PM20D1 promoter. EH38E3989962 shows signatures of regulatory activity in activated B cells, consistent with the lymphoblastoid cell line context assayed here. Chromatin interaction data support physical linkage between this enhancer and the PM20D1 regulatory domain in matched cell contexts, including Hi-C loops in B cells (ENCODE experiment ENCSR847RHU) and ChIA-PET support in WTC11 cells (ENCODE experiment ENCSR176OZL). The enhancer variant is associated with expression of multiple genes in the local regulatory neighborhood, including PM20D1, RAB29, SLC41A1, NUCKS1, and PM20D1-AS, but the magnitudes of the eQTL effects are not equivalent. The largest and most significant genotype-associated expression differences are observed for the PM20D1 sense and antisense transcriptional unit, with the strongest effect on PM20D1-AS and the next strongest on PM20D1, and these two effects run in opposite directions: the G allele is associated with increased expression of PM20D1-AS and decreased expression of PM20D1. This direction is consistent with the methylation pattern at the PM20D1 promoter, where the same allele tracks higher promoter methylation and reduced PM20D1 expression. RAB29, SLC41A1, and NUCKS1 show significant but more modest genotype-associated expression shifts. Together, these data indicate that the variant within EH38E3989962 is not simply acting on the nearest coding gene (SLC41A1) but instead defines a local regulatory haplotype whose strongest effects are centered on the PM20D1 sense and antisense transcriptional unit.

### Supplementary Note 4 - Pangenie SV Counts and QV Estimates

Our analyses are based on the T2T-CHM13 HPRC v2.0 Minigraph-Cactus graph constructed from 462 assembled haplotypes. We used the VCF output (https://s3-us-west-2.amazonaws.com/human-pangenomics/pangenomes/scratch/2025_02_28_minigraph_cactus/hprc-v2.0-mc-chm13/hprc-v2.0-mc-chm13.vcf.gz) which contains all top-level bubbles of the graph. We ran our decomposition pipeline (https://github.com/eblerjana/genotyping-pipelines/tree/main/prepare-vcf-MC, commit b2a4009) that we had introduced previously [19] to annotate each bubble allele with its nested variants. We first preprocess the VCF by filtering out variants for which more than 20% of haplotypes carry a missing allele ("."). We also convert genotypes of male samples on chrX and chrY to a homozygous representation by duplicating the haplotype with the fewest missing alleles. Our decomposition approach then detects nested variation based on the node traversals of each bubble allele through the graph and adds annotations to the input VCF encoding these nested variants ("bubble VCF"). Additionally, it produces a second, bi-allelic VCF containing a separate record for each nested allele ("callset VCF"). We use the "bubble VCF" as input to PanGenie, and the "callset VCF" to convert PanGenie genotypes for all bubbles to genotypes for the nested alleles. The VCFs are available at:

- bubble VCF (mc_filtered_ids.vcf.gz): https://zenodo.org/records/15223961/files/mc_filtered_ids.vcf.gz?download=1

- callset VCF (mc_filtered_ids_biallelic.vcf.gz): https://zenodo.org/records/15223961/files/mc_filtered_ids_biallelic.vcf.gz?download=1

The corresponding reference genome can be obtained from: https://s3-us-west-2.amazonaws.com/human-pangenomics/T2T/CHM13/assemblies/analysis_set/chm13v2.0_maskedY_rCRS.fa.gz

We ran PanGenie (v4.2.1) on all 3,202 samples from the 1000 Genomes Project[20,21] and on additional 6 HPRC v2.0 samples (HG01123, HG02486, HG02559, NA21309, HG002, HG005) using Illumina data[19]. Reads were obtained from:

- 3,202 1000G samples: http://ftp.sra.ebi.ac.uk/vol1/fastq/
- HG002: https://ftp-trace.ncbi.nlm.nih.gov/giab/ftp/data/AshkenazimTrio/HG002_NA24385_son/NIST_Illumina_2x250bps/reads/
- NA21309: https://s3-us-west-2.amazonaws.com/human-pangenomics/index.html?prefix=working/HPRC_PLUS/NA21309/raw_data/Illumina/child/
- HG01123, HG02486, HG02559: https://s3-us-west-2.amazonaws.com/human-pangenomics/index.html?prefix=submissions/30E441F3-6820-4BF6-BCF4-E64D56C8D6A4--TRUSEQ/
- HG005: https://s3-us-west-2.amazonaws.com/human-pangenomics/index.html?prefix=submissions/FFC78D9F-296E-41DD-9E50-C4B5806613EE--HPRC_PLUS_GIAB/HG005/raw_data/Illumina/child/brain-genomics/

For genotyping, we first ran PanGenie-index once on the input VCF:

```
PanGenie-index -v mc_filtered_ids.vcf -r
T2T-CHM13v2.0_maskedY_rCRS.fa -o index -t 24
```

And then genotyped each of the 3,208 samples using the commands:

```
PanGenie -f index -i <sample>_reads.fasta -o pangenie_<sample> -t 24
-j 24 -s <sample>

cat pangenie_<sample>_genotyping.vcf | python3
convert-to-biallelic.py mc_filtered_ids_biallelic.vcf.gz | bgzip >
pangenie_<sample>_genotyping_bi.vcf.gz
```

The convert-to-biallelic.py script is available at https://github.com/eblerjana/pangenie/blob/c1375ae6a4ab73b3346b7b02aff1e4f48ae66da5/pipelines/run-from-callset/scripts/convert-to-biallelic.py.

We filtered our genotypes based on a regression model we had introduced earlier[19,22,23], using the following filters:

- **ac0 fail:** a variant allele was genotyped with allele frequency of 0.0 across all samples
- **mendel fail:** the mendelian consistency across trios is less than 85 % for a variant allele. We exclude all trios with only 0/0, only 0/1 and only 1/1 genotypes.
- **gq fail:** less than 20 high quality genotypes were reported for a variant allele
- **self fail:** genotyping accuracy of a variant allele across the panel samples is less than 90%
- **nonref fail:** not a single non-0/0 genotype was genotyped correctly across all panel samples

We compared our genotyped set to PanGenie genotypes we had previously generated for the HPRC1 graph[19], the HGSVC3 project[23] and to an Illumina based SV discovery set generated with traditional, alignment-based SV callers[20,23]. Since the first and latter sets are GRCh38-based, while the other two sets are T2T-CHM13-based, we compared the callsets based on the number of SVs present in each sample. Prior to comparison, we ran `truvari collapse`[24] in order to merge similar SV alleles in the two PanGenie based sets using the parameters: `truvari collapse -r 500 -p 0.95 -P 0.95 -s 50 -S 100000`. We additionally included unfiltered PanGenie sets for HPRC1, HGSVC3 and HPRC2 genotyped sets, including sites not reliably genotypable; as well as the HPRCy2 panel VCF provided as an input to PanGenie. We show respective numbers of variants in **Fig. 7a**.

We phased our unfiltered set of genotypes with population-based phasing tool SHAPEIT5 (v5.1.1)[25]. We provided our bi-allelic "callset VCF" as a reference panel to SHAPEIT (`--reference`), after filtering out sites with missing genotyping alleles ("."), as they cannot be handled by SHAPEIT. We provided our unfiltered PanGenie genotypes across all cohort samples to `--input`. We provided pedigree information (`--pedigree`) and genetic maps (`--map`, obtained from: https://github.com/JosephLalli/phasing_T2T/tree/main/resources/recombination_maps/t2t_native_scaled_maps/). For chromosome X, we additionally provided a list of all male samples (`--haploid`). After phasing, we constructed consensus haplotypes of all samples by implanting phased variants into the T2T-CHM13 reference genome with `bcftools consensus`[26].

We applied our QV estimation pipeline[23] in order to compute variant-based QVs within windows of 1Mbp along the consensus haplotypes. We used high quality de novo haplotype-resolved assemblies from the HGSVC3 as a ground truth. We furthermore overlapped our windows with annotations from BISER[27] and RepeatMasker (http://www.repeatmasker.org/). We then plotted histograms of QVs observed across each haplotype, which each bar annotated by the fractions of overlaps with the BISER and RepeatMasker annotations (**Fig. 7c**).

### Supplementary Note 5 - ctyper Genotyping of Challenging Medically Relevant Loci

We tested how the expanded pangenome can improve genotyping accuracy in complex copy-number variable regions and challenging medically relevant genes for ctyper[28], a method that matches shared alleles between a short-read dataset and a pangenome. Using leave-one-out benchmarking on samples shared between the two cohorts, the ctyper derived genotypes with our HPRC Phase 2 assemblies (N=104) achieved higher accuracy than a pangenome constructed from HGSVC3 assemblies (N=458) for both full sequences (QV = 37.2 vs. 35.4) and unrepetitive regions (QV = 42.5 vs. 41.0).

We evaluated genotyping accuracy by benchmarking genotyped allele sequences against ground-truth diploid genome assemblies. Three samples shared between the HPRC and HGSVC panels (HG02818, NA19036, and HG002) were used for evaluation. Short-read NGS data for HG02818 and NA19036 were obtained from the 1000 Genomes Project, and HG002 data were obtained from GIAB and down-sampled to approximately 30× coverage. For each sample, genotyped alleles from pangenome matrices spanning 273 challenging medically relevant genes[29] were extracted as FASTA sequences and aligned to both haplotypes of the corresponding ground-truth assemblies using minimap2[30] with base-level alignment output.

To resolve multiple alignments and identify the most likely source locus for each genotyped allele, we applied a greedy pairing strategy. All candidate alignments were ranked by alignment score (matched bases minus mismatches), and the highest-scoring alignments were iteratively selected subject to two constraints: each allele was assigned to at most one genomic location, and overlapping alignments within the same locus group (>50% reciprocal overlap) were excluded to ensure unique pairing.
Sequence concordance for each paired alignment was quantified by parsing `cs` tags to count exact matches and mismatches. Metrics were computed over both the full aligned sequence and non-repeat-masked regions only, with repeat masking performed using Windowmasker. Per-allele accuracy was summarized as a Phred-scaled quality value (QV), defined as $-10 \times \log_{10}(\text{mismatched bases} / \text{aligned length})$, with zero-mismatch alleles capped at QV = 60. QV distributions and mismatch ratios were then compared across reference panels to assess overall genotyping accuracy.

### Supplementary Note 6 - RCCX

The RCCX pangenome built from 462 haplotypes, GRCH38, and T2T-CHM13, contains 771 informative marker nodes that are specific to the pseudogene module (with CYP21A1P) or the gene module (with CYP21A2). Parakit relies on those markers to detect fusion or gene conversions events that could be associated with a rare recessive disease, congenital adrenal hyperplasia. The number of markers and their coverage across the region affect the power and resolution of this approach. Hence, a large collection of high-quality haplotypes like the HPRC dataset is essential for producing a comprehensive pangenome for this approach. For example, a simple pangenome with just the two modules in GRCh38 would only recapitulate 208 markers.

The inclusion of the 94 haplotypes from release 1 adds 302 additional markers. Release 2 further adds 261 additional markers and improves the coverage of informative markers across the module (**Supp. Figure rccx-markers**).

This pangenome and Parakit were used to characterize the 462 haplotypes and identified three fusions overlapping the CYP21A2 gene that lead to a non-functional haplotype and could contribute to a disease phenotype if combined with a second pathogenic variant. Similarly, four haplotypes carried a gene conversion that resulted in a known pathogenic variant associated with congenital adrenal hyperplasia: Gln319Ter (c.955C>T), Met240Lys (c.719T>A), Val282Leu (c.844G>T), and In2G (c.293-13C>G). We also observed three notable haplotypes, carried by HG01071, HG02717 and HG02738, with three modules: a pseudogene module, then a module with a gene carrying Gln319Ter (c.955C>T) and a gene-to-pseudogene fusion ~1 kbp downstream of CYP21A2, and finally a functional gene module.

### Supplementary Note 7 - D4Z4

Facioscapulohumeral muscular dystrophy type 1 (FSHD1) is among the most common inherited myopathies (prevalence ~1 in 8,000–20,000)[31] and is caused by contraction of the D4Z4 macrosatellite array on chromosome 4q35 to 1–10 repeat units on a permissive haplotype, leading to chromatin relaxation and cytotoxic expression of the DUX4 retrogene in skeletal muscle[32,33]. Healthy individuals typically carry 11–100 units, and only haplotypes bearing a functional polyadenylation signal are pathogenic; the nearly identical D4Z4 arrays on chromosome 10q26 almost exclusively carry a non-functional signal variant and do not cause diseas.

Using KaryoScope (**Methods**), we resolved D4Z4 arrays across the 249 telomere-to-telomere chromosome 4 haplotypes (154 individuals) in HPRC2 (Figure 7f). The region distal to D4Z4 exists in two polymorphic forms: 4qA, which carries a terminal β-satellite repeat and the adjacent polyadenylation signal required for pathogenic DUX4 expression[32,34], and 4qB, which carries a distinct distal sequence derived from a subtelomeric transfer that replaced the ancestral β-satellite region[35] (**Supplementary Fig. 42a-b D4Z4_SUPPa-b**). Of the 239 haplotypes carrying a single D4Z4 array, 128 (53.6%) were 4qA and 111 (46.4%) were 4qB. The remaining haplotypes comprised nine with two or more arrays (six with two 4qA-type arrays; three with a degraded array alongside a canonical one) and one with no detectable array. Among the 128 single-array 4qA haplotypes, 59 (46.1%) carried a functional polyadenylation signal (ATTAAA), 68 (53.1%) carried a non-functional variant (ATCAAA), and 1 had none. Functional-signal arrays had a median of 26 repeat units (range: 8–62; mean: 28.5), non-functional arrays showed a similar distribution (median: 26; range: 7–133), and 4qB arrays, which universally lacked the signal, had a median of 22 (range: 7–59). Two of the 154 individuals (1.3%) carried a contracted allele in the FSHD1 range (≤10 units) on a single-array permissive 4qA haplotype with a functional signal: HG03470 (8 units) and HG03139 (10 units; **Supplementary Fig. 42c D4Z4_SUPPc**). Both fall at the upper end of the pathogenic range, where penetrance is incomplete — consistent with the absence of clinical ascertainment in the HPRC cohort[36].

We also found one structurally notable case: a 4qA haplotype in HG02257 carried two distinct D4Z4 arrays (a 25-unit array and a contracted 10-unit array, both with a functional signal), the shorter of which meets the FSHD1 sequence criteria. Because a contracted array in a multi-array configuration is not categorizable in the canonical single-array model for the genetic basis of FSHD1, we cannot confidently predict its effects and we do not count it among the carriers above. The pangenome exposes structural complexity like this at this locus; configurations like this are only resolvable in complete assemblies.

### Supplementary Note 8 - Variant Calling with Pangenome-Aware DeepVariant

To show that HPRC2 pangenome can boost the performance of variant calling pipelines, we trained pangenome-aware DeepVariant[37] models using data from three short read sequencing platforms, including Illumina (using both PCR-free and PCR+ library preparations with HiSeqX and NovaSeq instruments), Element AVITI (using both 500bp and 1000bp insert sizes), Roche Sequencing by Expansion Duplex (SBX-D) [38,39], and Complete Genomics DNBSEQ-T7+[39]. To create appropriate training data sets with available truth sets, we used reads from seven Genome-In-A-Bottle (GIAB) samples (HG001 to HG007). For HG001 and HG002, we used the Platinum Genomes[40] and GIAB-GRCh38-v5.0q[41] truth sets, respectively, and for the remaining samples, GIAB-v4.2.1[42] truth sets were used. These truth sets are all based on the GRCh38 reference. The reads from each platform were mapped to HPRC2 pangenome using vg giraffe (v1.71)[43,44] after diploid haplotype sampling of the graph from 32 candidate haplotypes (by using `--diploid-sampling` and `--num-haplotypes 32` in vg haplotypes). The alignments were then surjected to GRCh38. We then used bamleftalign from the freebayes package[45] to left-shift indels followed by running ABRA2[46], a local assembly-based realigner, to align reads more accurately. For Roche SBX-D data, ABRA2 realignment was skipped. Chromosomes 20, 21 and 22 were excluded during training. Chromosomes 21 and 22 were then used as validation and picking the best model to avoid overfitting to the training dataset. The results on chromosome 20, which is used as the test dataset, are explained later.

The WDL workflow for mapping reads is available from:

https://github.com/vgteam/vg_wdl/blob/62f07840ed62260e8c9e238632d288f0e41a2350/workflows/giraffe.wdl

The input JSON files used to map HG002 reads that were later used for testing DeepVariant are listed below:

- For Illumina (PCR-free Novaseq): https://s3-us-west-2.amazonaws.com/human-pangenomics/submissions/fa40d6a2-9bf6-4ebf-8f6e-798c3937db22--HPRC_V2_DEEPVARIANT_TRAINING/hprc-v2.1-mc-grch38-eval/illumina/mappings_vg_1.71/giraffe_input_jsons/HG002.novaseq.pcr-free.40x_giraffe.json

- For Element (1000bp insert size):
https://s3-us-west-2.amazonaws.com/human-pangenomics/submissions/fa40d6a2-9bf6-4ebf-8f6e-798c3937db22--HPRC_V2_DEEPVARIANT_TRAINING/hprc-v2.1-mc-grch38-eval/element/mappings_vg_1.71/giraffe_input_jsons/HG002.element.cloudbreak.1000bp_ins_giraffe.json

- For Roche SBX-D:
https://s3-us-west-2.amazonaws.com/human-pangenomics/submissions/fa40d6a2-9bf6-4ebf-8f6e-798c3937db22--HPRC_V2_DEEPVARIANT_TRAINING/hprc-v2.1-mc-grch38-eval/roche_sbx_d/mappings_vg_1.71_pangenome_consensus_caller_0.80.1/giraffe_input_jsons/HG002.roche.sbx_d.with_prune_low_cplx.YC_giraffe.json

- For Complete Genomics DNBSEQ-T7+:
https://s3-us-west-2.amazonaws.com/human-pangenomics/submissions/fa40d6a2-9bf6-4ebf-8f6e-798c3937db22--HPRC_V2_DEEPVARIANT_TRAINING/hprc-v2.1-mc-grch38-eval/dnbseq_t7_plus/mappings_vg_1.71/giraffe_input_jsons/HG002.dnbseq.t7plus.hprc_v2.1.hap32_to_dip.vg_1.71.60x_giraffe.json

Since pangenome-aware DeepVariant needs a separate sampled graph during inference, we created a haplotype-sampled graph with 32 haplotypes for each read set using vg haplotypes. The WDL workflow used for haplotype sampling is available from:

https://github.com/vgteam/vg_wdl/blob/0278b7836c7b218e062299d432f68f668e66b65a/workflows/haplotype_sampling.wdl

The input JSON files used to create the HG002 GBZ files that were later used for testing DeepVariant are listed below:

- For Illumina (PCR-free Novaseq):
https://s3-us-west-2.amazonaws.com/human-pangenomics/submissions/fa40d6a2-9bf6-4ebf-8f6e-798c3937db22--HPRC_V2_DEEPVARIANT_TRAINING/hprc-v2.1-mc-grch38-eval/illumina/gbz_files_hap32_vg_1.71/haplotype_sampling_input_jsons/HG002.novaseq.pcr-free.40x.hprc_v2.1.hap32.vg_1.71_haplotype_sampling.json

- For Element (1000bp insert size):
https://s3-us-west-2.amazonaws.com/human-pangenomics/submissions/fa40d6a2-9bf6-4ebf-8f6e-798c3937db22--HPRC_V2_DEEPVARIANT_TRAINING/hprc-v2.1-mc-grch38-eval/element/gbz_files_hap32_vg_1.71/haplotype_sampling_input_jsons/HG002.element.cloudbreak.1000bp_ins.hprc_v2.1.hap32.vg_1.71_haplotype_sampling.json

- For Roche SBX-D:
https://s3-us-west-2.amazonaws.com/human-pangenomics/submissions/fa40d6a2-9bf6-4ebf-8f6e-798c3937db22--HPRC_V2_DEEPVARIANT_TRAINING/hprc-v2.1-mc-grch38-eval/roche_sbx_d/gbz_files_hap32_vg_1.71_pangenome_consensus_caller_0.80.1/hapl

otype_sampling_input_jsons/HG002.roche.sbx_d.without_prune_low_cplx.YC_haplotype_sampling.json

- For Complete Genomics DNBSEQ-T7+: https://s3-us-west-2.amazonaws.com/human-pangenomics/submissions/fa40d6a2-9bf6-4ebf-8f6e-798c3937db22--HPRC_V2_DEEPVARIANT_TRAINING/hprc-v2.1-mc-grch38-eval/dnbseq_t7_plus/gbz_files_hap32_vg_1.71/haplotype_sampling_input_jsons/HG002.dnbseq.t7plus.hprc_v2.1.hap32.vg_1.71.60x_haplotype_sampling.json

There is a pangenome-based correction step, recommended by Roche, to obtain more accurate SBX-D alignments after running vg giraffe.This involves using a Roche-developed tool called Pangenome Consensus Caller (PCC-v0.80.1), which takes a vg BAM file as input. Using the available Duplex pair sequences, PCC processes each alignment and evaluates the consistency of each sequence with the pangenome by examining the read-to-graph alignments. It then selects the Duplex pair with highest similarity and reports it in the final BAM file. To be able to use PCC, we turned on the `--comments-as-tags` flag in vg giraffe to preserve the YC tag, which contains the information required to reconstruct the sequences of Duplex pairs. To keep the original read-to-graph mappings in the vg BAM file, we turned on the `--add-graph-aln` flag in vg giraffe and the `--graph-aln` flag in vg surject.

We followed the instructions for running PCC from: https://roche-axelios.gitbook.io/xoos/analysis-tools/pangenome_consensus_caller

The pangenome-aware models trained on vg-giraffe-mapped reads were tested on HG002 reads from three sequencing platforms which includes Illumina NovaSeq (with 50x coverage), Element (40x, 1kb insert size), Roche SBX-D (60x of both duplex and simplex sequences), and Complete Genomics DNBSEQ-T7+ (60x). The resulting call sets were then benchmarked against GIAB-GRCh38-v5.0q using aardvark GenoType variant comparison[47].

The VCF and BED files for the GIAB-GRCh38-v5.0q truth set were:

- https://ftp-trace.ncbi.nlm.nih.gov/ReferenceSamples/giab/release/AshkenazimTrio/HG002_NA24385_son/v5.0q/HG002_GRCh38_v5.0q_smvar.vcf.gz
- https://ftp-trace.ncbi.nlm.nih.gov/ReferenceSamples/giab/release/AshkenazimTrio/HG002_NA24385_son/v5.0q/HG002_GRCh38_v5.0q_smvar.vcf.gz.tbi
- https://ftp-trace.ncbi.nlm.nih.gov/ReferenceSamples/giab/release/AshkenazimTrio/HG002_NA24385_son/v5.0q/HG002_GRCh38_v5.0q_smvar.benchmark.bed

To obtain a baseline to compare pangenome-based pipelines against, we used BWA-MEM (v0.7.17, default parameters) to map HG002 short reads to the GRCh38 reference. Next, linear-reference-based DeepVariant was run on Illumina and Element BAM files and pangenome-aware DeepVariant on SBX-D and DNBSEQ-T7+ BAM files. The strong performance of the pangenome approach is so convincing that new sequencing technologies such as Roche SBX are developing primarily for the pangenome, which is why the SBX and DNBSEQ-T7+ platforms have only pangenome-aware DeepVariant models but not a linear-reference-based one. The percentage of total error reductions achieved by using pangenome in DeepVariant relative to the linear-reference-based baseline ranges from 36.0% to 52.3% for all variants, with SNPs (50%-62.7%) being more affected than InDels (15%-36.5%) (**Fig. 7g, Supplementary Table 13**).

Chromosome 20 was excluded during DeepVariant training and to ensure that the pangenome improvements are not due to overtraining, we performed a separate round of benchmarking only on chromosome 20, for both pangenome and linear-reference pipelines. The error reductions in chromosome 20 were highly comparable to the ones reported for the whole genome (**Supplementary Table 14**).

The HG002 vg BAM files used for testing with DeepVariant are listed below:

- Illumina NovaSeq:
  - https://s3-us-west-2.amazonaws.com/human-pangenomics/submissions/fa40d6a2-9bf6-4ebf-8f6e-798c3937db22--HPRC_V2_DEEPVARIANT_TRAINING/hprc-v2.1-mc-grch38-eval/illumina/mappings_vg_1.71/HG002.novaseq.pcr-free.40x_merged.positionsorted.bam
- Element:
  - https://s3-us-west-2.amazonaws.com/human-pangenomics/submissions/fa40d6a2-9bf6-4ebf-8f6e-798c3937db22--HPRC_V2_DEEPVARIANT_TRAINING/hprc-v2.1-mc-grch38-eval/element/mappings_vg_1.71/HG002.element.cloudbreak.1000bp_ins_merged.positionsorted.bam
- SBX-D:
  - https://s3-us-west-2.amazonaws.com/human-pangenomics/submissions/fa40d6a2-9bf6-4ebf-8f6e-798c3937db22--HPRC_V2_DEEPVARIANT_TRAINING/hprc-v2.1-mc-grch38-eval/roche_sbx_d/mappings_vg_1.71_pangenome_consensus_caller_0.80.1/HG002.roche.sbx_d.with_prune_low_cplx.YC_merged.positionsorted.pangenome_consensus_0.80.1.bam
- DNBSEQ-T7+:
  - https://s3-us-west-2.amazonaws.com/human-pangenomics/submissions/fa40d6a2-9bf6-4ebf-8f6e-798c3937db22--HPRC_V2_DEEPVARIANT_TRAINING/hprc-v2.1-mc-grch38-eval/dnbseq_t7_plus/mappings_vg_1.71/HG002.dnbseq.t7plus.hprc_v2.1.hap32_to_dip.vg_1.71.60x_merged.positionsorted.bam

The HG002 haplotype-sampled GBZ files (with 32 haplotypes) used for testing with pangenome-aware DeepVariant are listed below:

- Illumina NovaSeq: https://s3-us-west-2.amazonaws.com/human-pangenomics/submissions/fa40d6a2-9bf6-4ebf-8f6e-798c3937db22--HPRC_V2_DEEPVARIANT_TRAINING/hprc-v2.1-mc-grch38-eval/illumina/gbz_files_hap32_vg_1.71/HG002.novaseq.pcr-free.40x.hprc_v2.1.hap32.vg_1.71.gbz
- Element:
https://s3-us-west-2.amazonaws.com/human-pangenomics/submissions/fa40d6a2-9bf6-

4ebf-8f6e-798c3937db22--HPRC_V2_DEEPVARIANT_TRAINING/hprc-v2.1-mc-grch38-eval/element/gbz_files_hap32_vg_1.71/HG002.element.cloudbreak.1000bp_ins.hprc_v2.1.hap32.vg_1.71.gbz
- SBX-D: https://s3-us-west-2.amazonaws.com/human-pangenomics/submissions/fa40d6a2-9bf6-4ebf-8f6e-798c3937db22--HPRC_V2_DEEPVARIANT_TRAINING/hprc-v2.1-mc-grch38-eval/roche_sbx_d/gbz_files_hap32_vg_1.71_pangenome_consensus_caller_0.80.1/HG002.roche.sbx_d.with_prune_low_cplx.YC.gbz
- DNBSEQ-T7+: https://s3-us-west-2.amazonaws.com/human-pangenomics/submissions/fa40d6a2-9bf6-4ebf-8f6e-798c3937db22--HPRC_V2_DEEPVARIANT_TRAINING/hprc-v2.1-mc-grch38-eval/dnbseq_t7_plus/gbz_files_hap32_vg_1.71/HG002.dnbseq.t7plus.hprc_v2.1.hap32.vg_1.71.60x.gbz

Linear-reference-based DeepVariant can be run with:

```
docker run \
--rm \
-v ${MOUNT_DIR}:${MOUNT_DIR} \
google/deepvariant:1.10.0 \
    /opt/deepvariant/bin/run_deepvariant \
    --model_type WGS \
    --ref ${REF_FA} \
    --reads ${BAM} \
    --haploid_contigs chrX,chrY \
    --par_regions_bed ${PAR_REGIONS_BED}
    --output_vcf ${OUTPUT_DIR}/${OUTPUT_PREFIX}.vcf.gz \
    --output_gvcf ${OUTPUT_DIR}/${OUTPUT_PREFIX}.g.vcf.gz \
    --num_shards ${THREADS} \
    --intermediate_results_dir ${OUTPUT_DIR}/intermediate_results_dir
```

The PAR regions BED file is available from:
https://storage.googleapis.com/deepvariant/case-study-testdata/GRCh38_PAR.bed

Pangenome-aware DeepVariant can be run with:

```
docker run \
--rm \
-v ${MOUNT_DIR}:${MOUNT_DIR} \
--shm-size 15gb \
${DOCKER} \
    /opt/deepvariant/bin/run_pangenome_aware_deepvariant \
    --model_type WGS \
    --ref ${REF_FA} \
    --reads ${BAM} \
```

```
--haploid_contigs chrX,chrY \
--par_regions_bed ${PAR_REGIONS_BED}
--gbz_shared_memory_size_gb 15 \
--make_examples_extra_args "pileup_image_height_pangenome=100" \
--pangenome ${PANGENOME} \
--customized_model ${MODEL_CKPT} \
--output_vcf ${OUTPUT_DIR}/${OUTPUT_PREFIX}.vcf.gz \
--output_gvcf ${OUTPUT_DIR}/${OUTPUT_PREFIX}.g.vcf.gz \
--num_shards ${THREADS} \
--intermediate_results_dir ${OUTPUT_DIR}/intermediate_results_dir
```

The pangenome-aware DeepVariant models, which should be passed through the `--customized_model` flag, are available at:

- Illumina/Element model (a single model was trained for both technologies): gs://brain-genomics-public/research/pangenome_aware_dv_paper/jun_2026/xm_experiments/248871034/wu_1/best/best/
- Roche SBX-D model: gs://brain-genomics-public/experimental/exp258251756/wu_1/best/
- Complete Genomics DNBSEQ-T7+: gs://brain-genomics-public/experimental/exp264057928/wu_1/best/

### Supplementary Note 9 - Variant calling with the Sentieon pangenome pipeline

Sequencing reads were processed using the Sentieon pangenome pipeline (sentieon-cli dnascope-pangenome). Reads were first aligned to a linear reference genome (GRCh38) using Sentieon bwa (v202503.03). In parallel, a personalized (sample-specific) pangenome reference was constructed from the HPRC pangenome: k-mers from the sample's reads were counted with KMC and used to sample sample-specific haplotypes from the pangenome with the vg toolkit (vg haplotypes, v1.72.0). The resulting personalized pangenome was then reformatted with the vg toolkit (v1.72.0), extracting its haplotype sequences as a FASTA together with the corresponding graph (GFA) used for coordinate lifting.

Some reads were identified by the tool as likely to benefit from personalized reference genome re-alignment. These reads were re-aligned to the personalized reference and the resulting alignments were lifted back to GRCh38 coordinates using Sentieon (v202503.03; alignment with sentieon minimap2 and lift-over with sentieon pgutil lift). Small variants were then called from both the original linear-reference alignment and the lifted personalized-reference alignment using Sentieon DNAscope (v202503.03) with the SentieonIlluminaPangenomeRealignWGS1.2.bundle model bundle, available from Sentieon. Population allele frequency (AF) and allele count (AC) annotations derived from the HPRC pangenome were transferred onto the DNAscope output VCF. Final variant genotyping and filtering were performed with Sentieon DNAModelApply (v202503.03), also using the SentieonIlluminaPangenomeRealignWGS1.2.bundle, to produce the final filtered call set.

To compare DNAscope (linear-reference-based) and DNAscope pangenome methods, HG002 Illumina PCR-free Novaseq (35x) reads were used for mapping and variant calling. Aardvark's GenoType variant comparison was then used to benchmark the two call sets against GIAB-GRCh38-v5.0q. The results showed that with pangenome integration, the total number of errors decreased by 52.4% relative to DNAscope (**Fig. 7g, Supplementary Table 13**).

The CRAM, VCF and Aardvark summary files have been uploaded to a Sentieon public S3 bucket.

The linear-reference-based mappings and VCF files created by the Sentieon pipeline are:

- s3://sentieon-support-public/pFDAv2.35x.20260604/giabv3/HG002_pfda_giabv3_deduped.cram
- s3://sentieon-support-public/pFDAv2.35x.20260604/giabv3/HG002_pfda_giabv3_deduped.cram.bai
- s3://sentieon-support-public/pFDAv2.35x.20260604/giabv3/HG002_pfda_giabv3_deduped.cram.crai
- s3://sentieon-support-public/pFDAv2.35x.20260604/giabv3/HG002_pfda_giabv3_dnascope.vcf.gz
- s3://sentieon-support-public/pFDAv2.35x.20260604/giabv3/HG002_pfda_giabv3_dnascope.vcf.gz.tbi
- s3://sentieon-support-public/pFDAv2.35x.20260604/giabv3/HG002_pfda_giabv3_dnascope_v5.0q_summary.tsv

The pangenome-aware VCF files created by the Sentieon pipeline are:

- s3://sentieon-support-public/pFDAv2.35x.20260604/grch38/HG002_pfda_grch38_dnascope-pangenome.vcf.gz
- s3://sentieon-support-public/pFDAv2.35x.20260604/grch38/HG002_pfda_grch38_dnascope-pangenome.vcf.gz.tbi
- s3://sentieon-support-public/pFDAv2.35x.20260604/grch38/HG002_pfda_grch38_dnascope-pangenome_v5.0q_summary.tsv

The files can be retrieved with the AWS CLI (with `--no-sign-request`) or with curl on the equivalent HTTPS URLs (for example, `curl -LO https://s3.amazonaws.com/sentieon-support-public/pFDAv2.35x.20260604/giabv3/HG002_pfda_giabv3_deduped.cram`).
